# A Lethal Genetic Incompatibility between Naturally Hybridizing Species in Mitochondrial Complex I

**DOI:** 10.1101/2021.07.13.452279

**Authors:** Benjamin M. Moran, Cheyenne Y. Payne, Daniel L. Powell, Erik N. K. Iverson, Alex E. Donny, Shreya M. Banerjee, Quinn K. Langdon, Theresa R. Gunn, Rebecca A. Rodriguez-Soto, Angel Madero, John J. Baczenas, Korbin M. Kleczko, Fang Liu, Rowan Matney, Kratika Singhal, Ryan D. Leib, Osvaldo Hernandez-Perez, Russell Corbett-Detig, Judith Frydman, Casey Gifford, Manfred Schartl, Justin C. Havird, Molly Schumer

## Abstract

The evolution of reproductive barriers is the first step in the formation of new species and can help us understand the diversification of life on Earth. These reproductive barriers often take the form of “hybrid incompatibilities,” where alleles derived from two different species no longer interact properly in hybrids. Theory predicts that hybrid incompatibilities may be more likely to arise at rapidly evolving genes and that incompatibilities involving multiple genes should be common, but there has been sparse empirical data to evaluate these predictions. Here, we describe a mitonuclear incompatibility involving three genes in physical contact within respiratory Complex I in naturally hybridizing swordtail fish species. Individuals homozygous for specific mismatched protein combinations fail to complete embryonic development or die as juveniles, while those heterozygous for the incompatibility have reduced function of Complex I and unbalanced representation of parental alleles in the mitochondrial proteome. We find that the impacts of different genetic interactions on survival are non-additive, highlighting subtle complexity in the genetic architecture of hybrid incompatibilities. We document the evolutionary history of the genes involved, showing for the first time that an incompatibility has been transferred between species via hybridization. This work thus provides the first glimpse into the genetic architecture, physiological impacts, and evolutionary origin of a complex incompatibility impacting naturally hybridizing species.

## Introduction

Biologists have long been fascinated by the question of how new species are formed and what mechanisms maintain isolation between them. One key factor in the formation and maintenance of new species is the emergence of genetic incompatibilities that reduce viability or fertility in hybrids^1^. As originally described by the Dobzhansky-Muller model of hybrid incompatibility^2,3^, the unique sets of mutations that accumulate in diverging species may interact poorly when they are brought together in hybrids, given that they have never been tested against one another by selection. Due to the technical challenges of identifying these interactions^4^, only around a dozen genes involved in hybrid incompatibilities have been precisely mapped^5^ and exploration of the functional and evolutionary causes of hybrid incompatibilities has been limited to a small number of cases in model organisms^4^.

This knowledge gap leaves key predictions about the evolutionary processes that drive the emergence of hybrid incompatibilities untested. For one, theory suggests that incompatibilities should be more common within dense gene networks, both because genes involved in such interactions are expected to be tightly co-evolving and because the number of potentially incompatible genotypes explodes as the complexity of the genetic interaction increases^6,7^. Consistent with this prediction, mutagenesis experiments have highlighted the sensitivity of multi-protein interactions to changes in any of their components^6^. However, genetic interactions are notoriously difficult to detect empirically except in systems with especially powerful genetic tools^8^, and this problem is exacerbated with complex genetic interactions^9,10^. Such technical challenges may explain the rarity of incompatibilities involving three or more genes in the empirical literature^6^ (but see^8,11–14)^.

Another open question is the degree to which the genes that become involved in hybrid incompatibilities are predictable from their molecular or evolutionary properties. Researchers have proposed for decades that rapid molecular evolution will increase the rate at which incompatibilities accumulate between species^4,5,15^. While several incompatibilities identified to date involve genes showing signatures of positive selection, it is unclear how unusual rates of protein evolution are in genes involved in hybrid incompatibilities relative to the genomic background^5,15^. The mitochondrial genome, in particular, has been proposed as a hotspot for the accumulation of genetic incompatibilities^16,17^, due to substitution rates up to 25× higher than the nuclear genome in many animals^18–20^ and the potential for sexually antagonistic selection driven by its predominantly maternal inheritance^21–23^, among other factors^24^. At the same time, nuclear and mitochondrial gene products must directly interact with each other in key steps of ATP synthesis, increasing the likelihood of coevolution between these genomes^25,26^. These molecular and evolutionary factors suggest that interactions between mitochondrial- and nuclear-encoded proteins could play an outsized role in the emergence of hybrid incompatibilities^16^. Crosses in numerous species have shown that hybrid viability often depends on the identity of the maternal species^27–30^, providing indirect evidence for the prevalence of mitonuclear incompatibilities. However, the field has struggled to move beyond these coarse-scale patterns, as few studies have successfully mapped mitonuclear incompatibilities to the single gene level^31–34^ and none of those identified to date have been studied in species that naturally hybridize.

As we begin to identify the individual genes underlying hybrid incompatibilities, the next frontier is evaluating the processes that drive their evolution. Over the past two decades, it has become abundantly clear that hybridization is exceptionally common in species groups where it was once thought to be rare^35,36^. As such, we now appreciate how frequently species derive genes from their relatives, and in many cases are a patchwork of ancestry from several genomes due to historical hybridization^37–39^. The impact of these dynamics on the evolution of hybrid incompatibilities have been poorly investigated^40^ since the foundational theory in this area was developed before the ubiquity of hybridization was fully appreciated^7^.

Here, we use an integrative approach to precisely map the genetic basis and physiological impacts of a lethal mitonuclear hybrid incompatibility in swordtail fish and uncover its evolutionary history. The sister species *Xiphophorus birchmanni* and *X. malinche* began hybridizing in the last ∼100 generations in multiple river systems^41^ after premating barriers were disrupted by habitat disturbance^42^, and are a powerful system to study the emergence of hybrid incompatibilities in young species. Despite their recent divergence^43^ (∼250,000 generations; 0.5% divergence per basepair), some hybrids between *X. birchmanni* and *X. malinche* experience strong selection against incompatibilities^43,44^. One incompatibility causing melanoma has been previously mapped in this system and population genetic patterns suggest that dozens may be segregating in natural hybrid populations^43–46^. Moreover, the ability to generate controlled crosses^47,48^ and the development of high-quality genomic resources^46,49^ makes this system particularly tractable for identifying hybrid incompatibilities and characterizing their evolution in natural populations. Leveraging data from controlled laboratory crosses and natural hybrid populations, we pinpoint two nuclear-encoded *X. birchmanni* genes that are lethal when mismatched with the *X. malinche* mitochondria in hybrids, explore the developmental and physiological effects of this incompatibility, and trace its evolutionary history.

### Admixture Mapping Reveals a Lethal Mitonuclear Incompatibility

To identify loci under selection in *X. birchmanni × X. malinche* hybrids, we generated ∼1X low-coverage whole-genome sequence data for 943 individuals from an F_2_ laboratory cross and 359 wild-caught hybrid adults, and applied a hidden Markov model to data at 629,661 ancestry-informative sites along the genome to infer local ancestry (∼1 informative site per kb^45,50^; Methods, Supplementary Information 1.1.1–1.1.4). Using these results, we found evidence for a previously unknown incompatibility between the nuclear genome of *X. birchmanni* and the mitochondrial genome of *X. malinche* (Supplementary Information 1.1.5– 1.1.10). Our first direct evidence for this incompatibility came from controlled laboratory crosses (Methods, Supplementary Information 1.1.1). Because the cross is largely unsuccessful in the opposite direction, all lab-bred hybrids were the offspring of F_1_ hybrids generated between *X. malinche* females and *X. birchmanni* males and harbored a mitochondrial haplotype derived from the *X. malinche* parent species. Offspring of F_1_ intercrosses are expected to derive on average 50% of their genome from each parent species. This expectation is satisfied genome-wide and locally along most chromosomes in F_2_ hybrids (on average 50.3% *X. malinche* ancestry; Fig. S1). However, we detected six segregation distorters genome-wide^48^, with the most extreme signals falling along a 6.5 Mb block of chromosome 13 and an 4.9 Mb block of chromosome 6 (Fig. 1A; Fig. 1D).

**Fig. 1.**
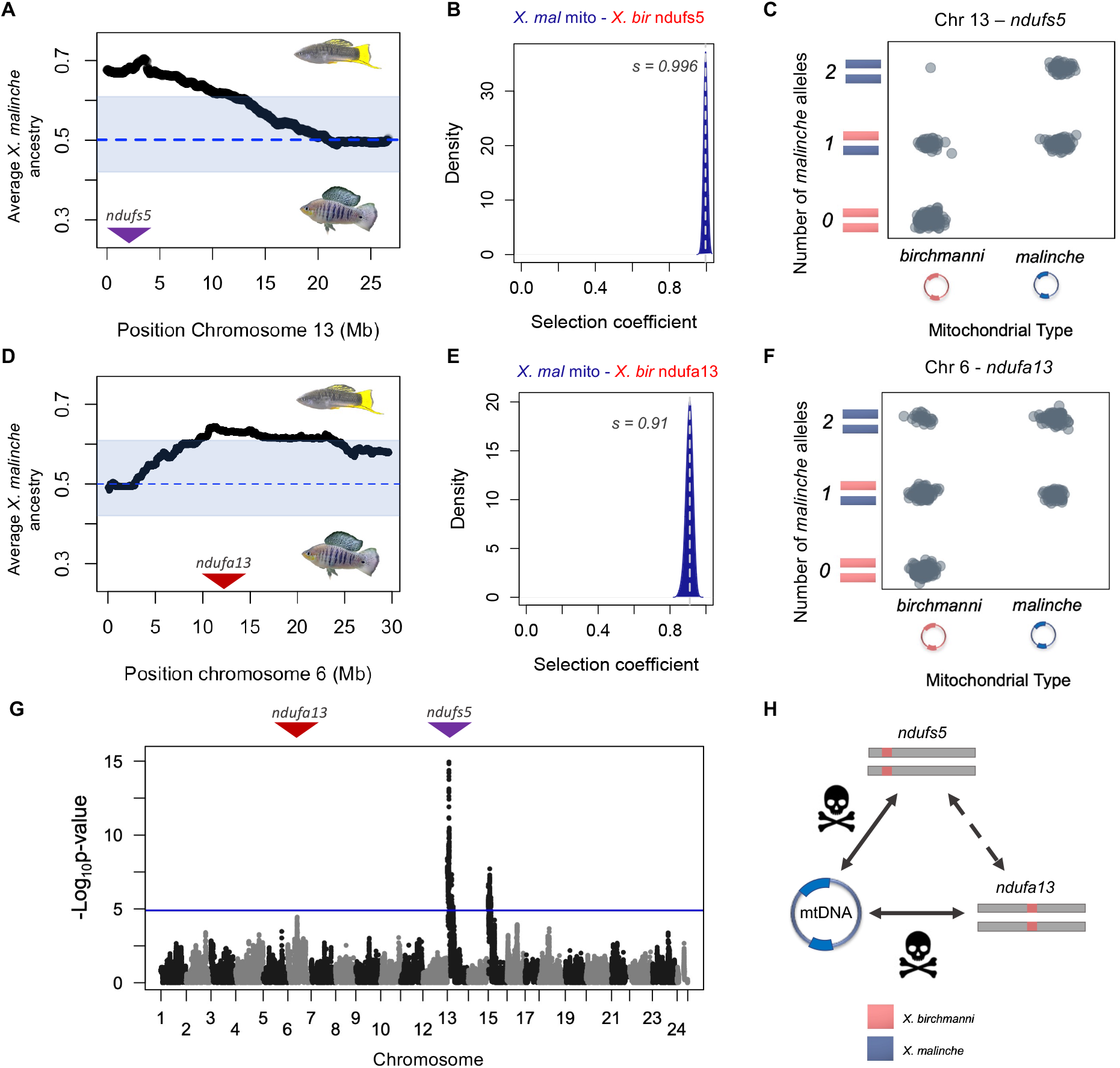
Admixture mapping pinpoints mitonuclear incompatibility in *Xiphophorus*. (**A**) Average ancestry of F_2_ hybrids on chromosome 13 reveals segregation distortion towards *X. malinche* ancestry across a large region of the chromosome. Blue envelope shows the 99% quantiles of *X. malinche* ancestry at all ancestry informative sites genome wide. Dashed line represents the expected *X. malinche* ancestry for this cross. Purple arrow points to the position of *ndufs5*. (**B)** Results of approximate Bayesian computation (ABC) simulations estimating the strength of selection on the *X. malinche* mitochondria when combined with *X. birchmanni* ancestry at *ndufs5*. Shown here is the posterior distribution from accepted simulations; vertical line and inset indicates the maximum a posteriori estimate for the selection coefficient. (**C**) Observed genotype frequencies of different genotype combinations of *ndufs5* (chromosome 13) and mitochondrial haplotypes in the admixture mapping population. (**D**) Average ancestry of F_2_s on chromosome 6, reveals segregation distortion towards *X. malinche* ancestry across a large region of the chromosome. Blue envelope and dashed line indicate 99% ancestry quantiles and expected ancestry in the cross as in (**A**), red arrow points to the position of *ndufa13*. (**E)** Results of approximate Bayesian computation (ABC) simulations estimating the strength of selection on the combination of *X. malinche* mitochondria with *X. birchmanni ndufa13*, as in (**B**). (**F**) Observed genotype frequencies of different genotype combinations of *ndufa13* (chromosome 6) and mitochondrial haplotypes in the admixture mapping population. (**G**) Admixture mapping results for associations between nuclear ancestry and mitochondrial haplotype in natural hybrids, controlling for genome-wide ancestry. Blue line indicates the 10% false positive rate genome-wide significance threshold determined by simulations. The peak visible on chromosome 15 is driven by interactions with the *X. birchmanni* mitochondria and an unknown nuclear gene and is discussed in Supplementary Information 1.1.10 and 1.2.1. (**H**) Schematic of identified interactions with the *X. malinche* mitochondrial genome from our mapping data. We detect a subtle interaction between *ndufs5* and *ndufa13* (indicated by the dashed line; see main text, Figure 2, and Supplementary Information 1.2.5).

Closer examination of genotypes in the chromosome 13 region showed that almost none of the surviving individuals harbored homozygous *X. birchmanni* ancestry in a 3.75 Mb subregion (Fig. 1C; Fig. S2; 0.1% observed vs 25% expected). This pattern is unexpected even in the case of a lethal incompatibility involving only nuclear loci (where simulations indicate that we should recover homozygous *X. birchmanni* ancestry in ∼10% of surviving individuals; Supplementary Information 1.1.1) but is consistent with a lethal mitonuclear incompatibility. Using approximate Bayesian computation (ABC) approaches we inferred what strength of selection against *X. birchmanni* ancestry in this region was consistent with the genotypes and ancestry deviations observed. We estimated posterior distributions of selection and dominance coefficients and inferred that selection on this genotype in F2s is largely recessive and essentially lethal (maximum a posteriori estimate *h* = 0.12 and *s* = 0.996, 95% credible interval *h* = 0.010– 0.194 and *s* = 0.986–0.999; Fig. 1B; Fig. S3; Methods; Supplementary Information 1.2.1–1.2.2).

The degree of segregation distortion observed in F_2_ individuals on chromosome 6 is also surprising (Fig. 1D). Only 3% of individuals harbor homozygous *X. birchmanni* ancestry in this region (compared to 0.1% in the chromosome 13 region and 25% on average at other loci across the genome; Fig 1F), which is again lower than expected for a nuclear-nuclear hybrid incompatibility (Supplementary Information 1.1.1). ABC approaches indicate that selection on homozygous *X. birchmanni* ancestry on chromosome 6 is also severe (maximum a posteriori estimate *h =* 0.09 and *s* = 0.91, 95% credible interval interval *h* = 0.01–0.21 and *s* = 0.87–0.94; Fig. 1E, Fig. S3, Supplementary Information 1.2.2). Thus, our F_2_ data show that homozygous *X. birchmanni* ancestry in regions of either chromosome 13 or chromosome 6 is almost completely lethal in hybrids with *X. malinche* mitochondria (Fig. 1H).

To formally test for the presence of a mitonuclear incompatibility involving chromosome 13 and chromosome 6, or elsewhere in the genome, we leveraged data from natural hybrid populations. Most naturally occurring *X. birchmanni* × *malinche* hybrid populations are fixed for mitochondrial haplotypes from one parental species (Supplementary Information 1.1.2, 1.1.6). However, a few populations segregate for the mitochondrial genomes of both parental types, and we focused on one such population (the “Calnali Low” population, hereafter the admixture mapping population). Admixture mapping for associations between nuclear genotype and mitochondrial ancestry (after adjusting for expected covariance due to genome-wide ancestry^44^) revealed two genome-wide significant peaks and one peak that approached genome-wide significance (Fig. 1G, Table S1–S3). The strongest peak of association spanned approximately 77 kb and fell within the region of chromosome 13 identified using F_2_ crosses (Fig. 1G). This peak was also replicated in another hybrid population (Fig. S4; Methods, Supplementary Information 1.1.5) and contains only three genes: the NADH dehydrogenase ubiquinone iron-sulfur protein 5 (*ndufs5*), E3 ubiquitin-protein ligase, and microtubule-actin cross-linking factor 1. Of these three genes, only *ndufs5* forms a protein complex with mitochondrially encoded proteins, which along with other evidence implicates it as one of the interacting partners in the mitonuclear incompatibility (Fig. 1C; see Supplementary Information 1.1.8 for analysis of other genes).

Analysis of three natural hybrid populations that had fixed the mitochondrial haplotype of one of the parental species (Fig. S5) confirmed that this region on chromosome 13 is under selection in natural hybrid populations, with the strongest signal of selection localizing precisely to the same three genes found under the admixture mapping peak (Fig. S6A; Supplementary Information 1.1.6). Moreover, comparing genotypes and phenotypes in siblings allowed us to exclude maternal effects as a driver of the chromosome 13 signal (Supplementary Information 1.1.7), and we ruled out the possibility that other confounding factors could generate the observed patterns (Supplementary Information 1.1.9).

We also identified a peak on chromosome 6 that approached genome-wide significance (Fig. 1G; Table S2; Supplementary Information 1.1.10) and fell precisely within the segregation distortion region previously mapped in F_2_ hybrids (Fig. 1D; Supplementary Information 1.1.1). This peak contained 20 genes including the mitochondrial Complex I gene *ndufa13* (Fig. S7–S8; Methods, Supplementary Information 1.1.10). Depletion of non-mitochondrial parent ancestry at *ndufa13* was unidirectional (Fig. 1F), consistent with selection acting only against the combination of the *X. malinche* mitochondria with homozygous *X. birchmanni* ancestry at *ndufa13* (see Supplementary Information 1.2.3–1.2.4). Genomic analyses in natural hybrid populations confirmed this asymmetry (Fig. S6B).

Together, these results indicate that at least two *X. birchmanni* nuclear genes cause incompatibility when they are mismatched in ancestry with the *X. malinche* mitochondria (Fig. 1H; Supplementary Information 1.2.5). These genes, *ndufs5* and *ndufa13*, belong to a group of proteins and assembly factors that form respiratory Complex I^51^ (see Table S1 for locations of the 51 annotated Complex I genes in the *Xiphophorus* genome). Complex I is the first component of the mitochondrial electron transport chain that ultimately allows the cell to generate ATP. Both nuclear proteins interface with several mitochondrially derived proteins at the core of the Complex I structure, hinting at the possibility that physical interactions could underlie this multi-gene mitonuclear incompatibility.

### Interactions with the *X. birchmanni* mitochondria

Admixture mapping analysis also identified a strong peak of mitonuclear association on chromosome 15, which we briefly discuss here and in Supplementary Information 1.1.10 and 1.2.1. This peak was associated with *X. birchmanni* mitochondrial ancestry (Fig. S9), indicating that it has a distinct genetic architecture from the incompatibility involving the *X. malinche* mitochondria and *X. birchmanni ndufs5* and *ndufa13*. Specifically, analysis of genotypes at the admixture mapping peak indicates that the *X. birchmanni* mitochondria is incompatible with homozygous *X. malinche* ancestry on chromosome 15 (Fig. 1C, Fig. S9). This region did not contain any members of Complex I, but dozens of genes in this interval interact with known mitonuclear genes (see Table S3; Supplementary Information 1.1.10). The fact that we detect incompatible interactions with both the *X. malinche* mitochondria (at *ndufs5* and *ndufa13*) and the *X. birchmanni* mitochondria (*ndufs5* and chromosome 15) in our admixture mapping results supports the idea that mitonuclear interactions can act as “hotspots” for the evolution of hybrid incompatibilities^16^.

### Lethal Effect of Incompatibility in Early Development

The combination of *X. birchmanni ndufs5* or *ndufa13* with the *X. malinche* mitochondria appears to be lethal by the time individuals reach adulthood. To investigate the developmental timing of the incompatibility, we genotyped pregnant females from the admixture mapping population and recorded the developmental stages of their embryos^52^ (swordtails are livebearing fish; see Methods). We found a significant interaction between developmental stage and *ndufs5* genotype, whereas *ndufa13* genotype did not affect developmental stage (Fig. S10–13; Supplementary Information 1.3.1). While developmental asynchrony is typically on the scale of 0–2 days in pure species^53^ (Supplementary Information 1.3.1), we observed much greater variation in broods collected from the admixture mapping population where the mitochondrial incompatibility is segregating (e.g. stages normally separated by roughly 12 days of development found in the same brood; Supplementary Information 1.3.1; Fig. 2A–B). Genotyping results revealed that embryos with homozygous *X. birchmanni* ancestry at *ndufs5* and *X. malinche* mitochondria are present at early developmental stages, but that these embryos failed to develop beyond a phenotype typical of the first seven days of gestation (the full length of gestation is 21– 28 days in *Xiphophorus*; Fig. 2A–B, Fig. 2D–E). Individuals with mismatched ancestry at *ndufs5* whose siblings were fully developed still had a detectable heartbeat but had consumed less yolk than their siblings and remained morphologically underdeveloped (Fig. 2D, Fig. S14–19). Unlike other species, in *Xiphophorus* this developmental lag could itself cause mortality, since embryos that do not complete embryonic development inside the mother fail to survive more than a few days after birth (Supplementary Information 1.3.1; Table S4). Given that Complex I inhibition lethally arrests development in zebrafish embryos^54,55^, we also tested the effects of Complex I inhibition on *X. birchmanni* and *X. malinche* fry, and found a similar level of sensitivity (Supplementary Information 1.3.2).

**Fig. 2.**
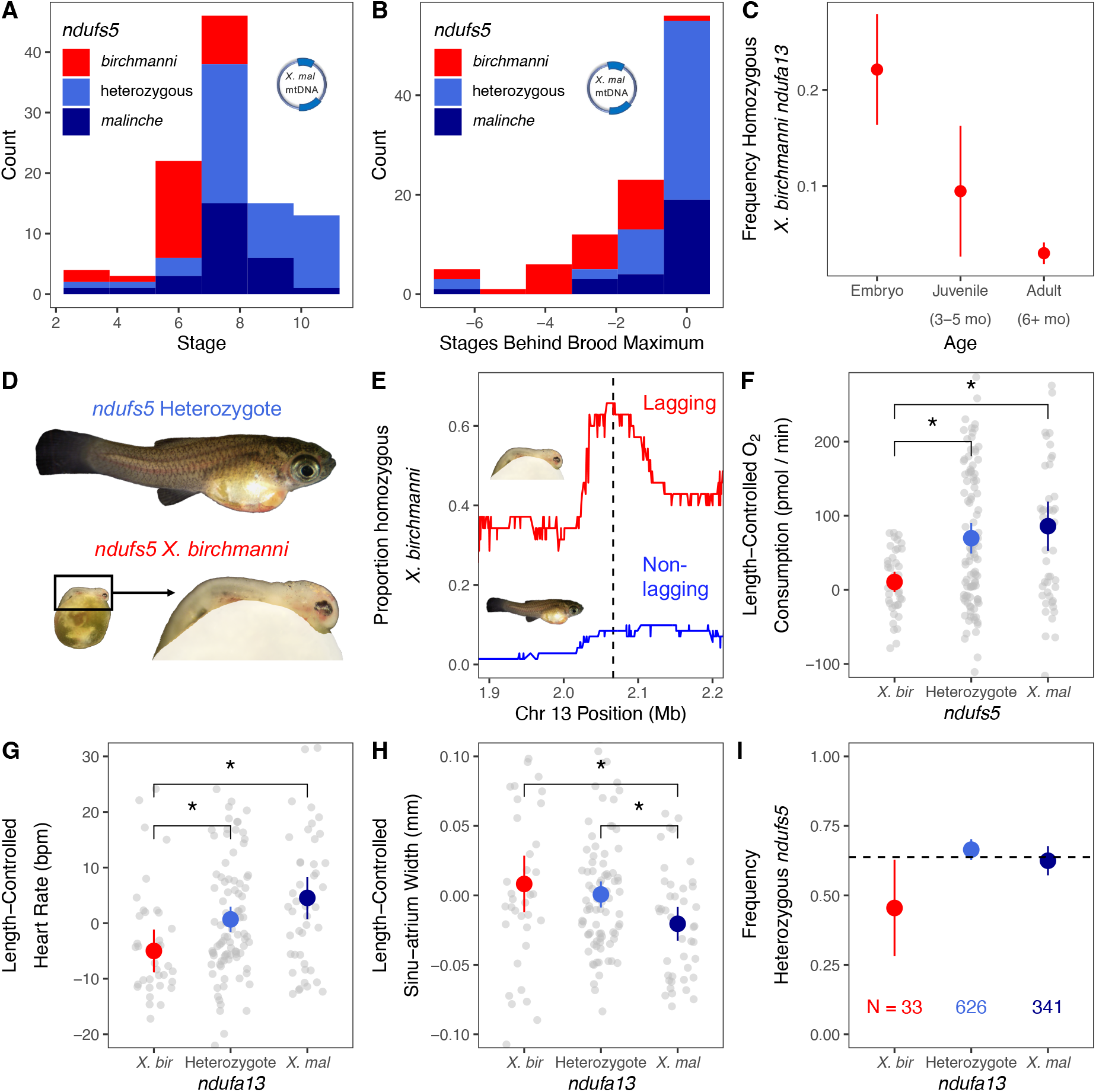
Impact of the hybrid incompatibility on *Xiphophorus* hybrid embryos. (**A**) Developmental stage and *ndufs5* genotypes of hybrid embryos with *X. malinche* mitochondria. **(B)** Lag in development of hybrid embryos with *X. malinche* mitochondria compared to the most developed embryo in their brood, as a function of *ndufs5* genotype. (**C**) Change in frequency of homozygous *X. birchmanni* ancestry at *ndufa13* in F_2_ hybrids over development. Points and lines represent the estimate ± 2 standard errors, based on observed frequency and sample size. (**D**) F_2_ hybrid siblings showing stark difference in developmental progression as a function of *ndufs5* genotype. Top photo shows entire length of heterozygote embryo, while bottom photos show lagging *ndufs5-*incompatible sibling at matched scale (left) and zoomed for detail (right). (**E**) Frequency of homozygous *X. birchmanni* ancestry along chromosome 13 in embryos with *X. malinche* mitochondria that lagged their siblings in developmental stage by ζ1 developmental stage (red) versus frequency of homozygous *X. birchmanni* ancestry in embryos that did not exhibit developmental lag (blue, see Supplementary Information 1.1.7). Dashed line indicates the location of *ndufs5*. Note that only 66% of embryos with developmental lag have homozygous *X. birchmanni* in this region, indicating that there are other causes of this phenotype. For the same analysis of chromosomes 6 and 15, where we see no clear difference in average ancestry as a function of lag status, see Fig. S11–13. (**F–H**) Respiratory and morphometric measurements in live F_2_ embryos. To control for the effect of length on other variables, residuals of each variable regressed against body length are plotted (see Supplementary Information 1.3.4-1.3.5). Gray points denote individual measurements, colored points and bars show mean ± 2 SE for each genotype, and brackets with asterisks denote significant differences based on a Tukey’s HSD test. (**F**) Relationship between *ndufs5* genotype and respiration in F_2_ hybrid embryos, as measured by Loligo plate respirometer. (**G**) Relationship between *ndufa13* genotype and heart rate in F_2_ hybrid embryos. (**H**) Relationship between *ndufa13* genotype and the width of the sinu-atrium, which peristaltically pumps blood from the yolk into the embryonic heart atrium. (**I**) Frequency of individuals heterozygous for ancestry at *ndufs5* (i.e. those with one *X. birchmanni* allele) among juveniles and adults of varying *ndufa13* genotypes. Points and lines represent observed frequency ± 2 standard errors. Dashed line represents expected frequency of *ndufs5* heterozygotes under a scenario of additive selection on *ndufs5* and *ndufa13* (see Supplementary Information 1.2.5).

In contrast to individuals with mismatched ancestry at *ndufs5,* those with *ndufa13* mismatch survived embryonic development but suffered mortality in the early post-natal period (Fig. 2C). We tracked 74 F_2_ fry from 24 hours post-birth through adulthood (Supplementary Information 1.3.3). We found that most fry with incompatible genotypes at *ndufa13* had already suffered mortality by the time tracking began, with only 7 individuals found 24 hours post-birth that were homozygous *X. birchmanni* at *ndufa13* (versus 19 expected; binomial *P* = 0.0005). No natural mortality was observed between 1 day and 3 months post-birth (one individual exhibiting distress was euthanized, and was a *ndufa13* incompatible homozygote; see Supplementary Information 1.3.3).

### Evidence for Complex Physiological and Fitness Effects in Incompatible Individuals

Our analysis of developing embryos indicates that individuals with the *ndufs5* incompatibility exhibit delayed or arrested embryonic development, whereas those with the *ndufa13* incompatibility did not. This suggests that these genes may drive lethality through partially distinct mechanisms. As such, we chose to further investigate the effects of *ndufs5* and *ndufa13* and their possible interactions using a paired genotyping and physiological approach throughout development. We sampled 235 F_2_ embryos at a range of developmental stages and measured their overall rates of respiration (Supplementary Information 1.3.4–1.3.5). We also used imaging of these embryos to track cardiovascular phenotypes as these have been associated with *ndufa13* defects in mammals^56^. We found that incompatible genotypes at *ndufs5* and *ndufa13* impacted a range of phenotypes, including heart rate, length relative to compatible siblings, and length-corrected head size (Fig. S14–22, Table S5–S7). *ndufa13* mismatch has a large effect on cardiovascular phenotypes, including heart rate and the size of the sinu-atrium (an embryo-specific heart chamber, Fig. 2G–H, Fig. S22, Table S6, S8); *ndufs5* affects only heart rate (Fig. S19, Table S6). We find initial evidence that cardiac defects persist into adulthood in surviving individuals with *ndufa13* mismatch (see Supplementary Information 1.3.3; Fig. S23). By contrast, *ndufs5* mismatch has a major impact on rates of respiration and yolk consumption during development (Fig. 2F, while *ndufa13* only affects respiration through an interaction with *ndufs5* and body length, Fig. S24–27, Table S9–11).

Naively, the separable impacts of incompatible genotypes at *ndufs5* and *ndufa13* could indicate that even though these proteins are in physical contact in Complex I (see below), they represent two distinct hybrid incompatibilities. We investigated this question by taking advantage of rare survivors of the *ndufa13* incompatibility in an expanded dataset of 1010 F_2_ hybrids. Using this dataset, we were able to identify dozens of survivors of the *ndufa13* incompatibility (3.4% of individuals) and found that genotypes at *ndufa13* and *ndufs5* were not independent (χ^2^ association test *P* = 0.032, Supplementary Information 1.2.5). Upon further investigation, we found that the majority of survivors of the *ndufa13* incompatibility had homozygous *X. malinche* ancestry at *ndufs5*, suggesting that harboring even one *X. birchmanni* allele at *ndufs5* may sensitize fry to the *ndufa13* incompatibility. Indeed, we found that individuals that had heterozygous ancestry at *ndufs5* were significantly under-represented among surviving *ndufa13* incompatible individuals (Permutation test *P* = 0.015, Fig. 2I, Supplementary Information 1.2.5). These findings highlight a subtle but significant non-additive effect of *ndufs5* and *ndufa13* on survival.

### Mitochondrial Biology in Adult Hybrids Heterozygous for the Incompatibility

Because few individuals homozygous for incompatible genotypes at *ndufs5* or *ndufa13* survive past birth, our previous experiments focused on embryos. However, the small size of *Xiphophorus* embryos prevents us from using assays that directly target Complex I. To further explore the effects of the hybrid incompatibility on Complex I function *in vivo*, we turned to adult F_1_ hybrids between *X. birchmanni* and *X. malinche* (Fig. 3A). Since F1 hybrids that derive their mitochondria from *X. malinche* and are heterozygous for ancestry at *ndufs5* and *ndufa13* are fully viable, we asked whether there was evidence for compensatory nuclear or mitochondrial regulation that might be protective in F_1_ hybrids. We found no evidence for significant differences in expression of *ndufs5* or *ndufa13* (Supplementary Information 1.3.6; Fig. S28–29) or in mitochondrial copy number (Supplementary Information 1.3.7; Fig. 3B) between F_1_ hybrids and parental species.

**Fig. 3.**
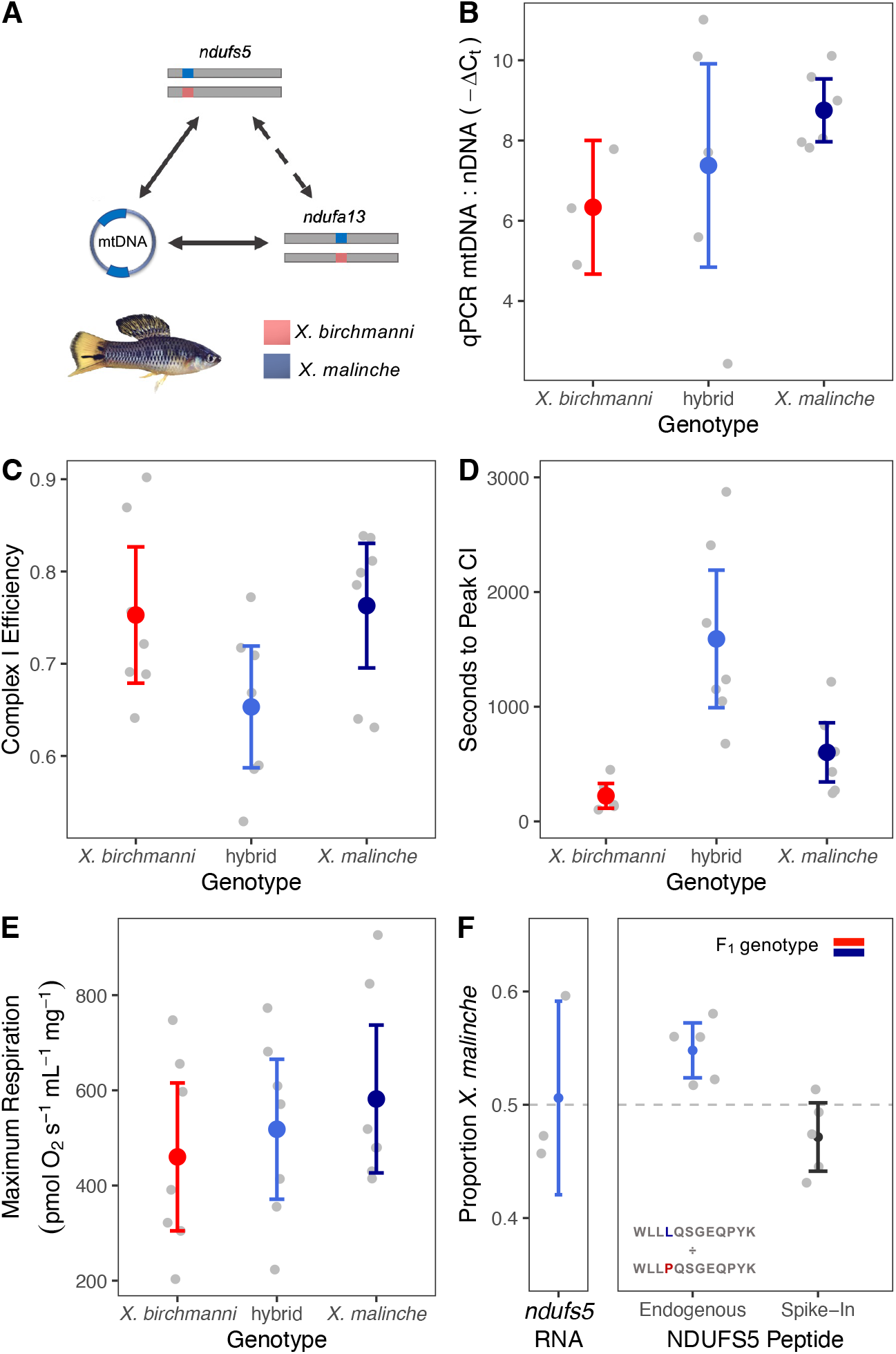
Physiological and proteomic phenotypes of viable heterozygotes harboring the hybrid incompatibility. In all panels, colored points and whiskers show the mean ± 2 standard errors, and gray points show individual data. (**A**) Schematic of ancestry at loci involved in the incompatibility in an *X. birchmanni* ξ *malinche* F_1_ hybrid. Hybrids heterozygous at both loci avoid the lethal effects of the interaction between the *X. malinche* mitochondria and *X. birchmanni* ancestry at *ndufs5* and *ndufa13* (Supplementary Information 1.2.2). (**B**) Results of rt-qPCR evaluating evidence for differences in mitochondrial copy number between genotypes in liver tissue. Ratio is derived from the difference in C_t_ between a single copy nuclear gene (*Nup43*) and the mitochondrial gene *ND1.* (**C**) Results of Oroboros O2K respirometer assay for adult *X. birchmanni*, *X. malinche*, and hybrid individuals with *X. malinche* mitochondria and heterozygous ancestry at *ndufs5* and *ndufa13* (n=7 per genotype) point to lower Complex I efficiency in heterozygous hybrids (orthogonal contrast *P* = 0.023). (**D**) Time to reach the maximum rate of Complex I-driven respiration after the addition of ADP differed between heterozygous hybrids and parental species (orthogonal contrast *P* < 0.001). Complex I-driven respiration begins with the addition of ADP, as the flow of electrons is previously limited by the inability of Complex V to relieve the proton gradient in the absence of its substrate (see Fig. S32 for example time-to-peak curves). Time to reach the peak in Complex I- and Complex II-driven respiration after the addition of succinate did not differ across genotypes (Fig. S33). (**E**) Maximum respiration rates during full O2K protocol as a function of genotype did not differ between groups despite significant differences in Complex I efficiency in heterozygotes (orthogonal contrast *P* = 0.94). (**F**) Allelic balance of *ndufs5* in the F1 hybrid proteome and transcriptome. Left subplot shows allele-specific expression of *ndufs5* in three adult F_1_ hybrids. Right subplot shows results of quantitative mass spectrometry analysis of *ndufs5* peptides in mitochondrial proteomes derived from five adult F_1_ hybrids. Data points show the proportion of area under the spectral curves contributed by the *X. malinche* allele in a given individual. Left column shows results for endogenous peptides present in F_1_ hybrids, and right column shows results for the control where heavy-labeled standards of each peptide were spiked in. Inset shows the identities of heavy-labeled peptides for each species.

With no indication of a compensatory regulatory response, we reasoned that we might be able to detect reduced mitochondrial Complex I function in hybrids heterozygous for ancestry at *ndufs5* and *ndufa13.* We quantified respiratory phenotypes in isolated mitochondria using an Oroboros O2K respirometer in adult hybrids and both parental species (Fig. S30; Methods, Supplementary Information 1.3.8). We found that Complex I efficiency was lower in hybrids compared to the two parental species (Fig. 3C, Fig. S31, orthogonal contrast *t* = -2.53, *P* = 0.023, n = 7 per genotype), and that the time required for hybrids to reach maximum Complex I-driven respiration was ∼2.5 times longer (orthogonal contrast *t* = 4.303, *P* < 0.001; Fig. 3D; Fig. S32). On the other hand, overall levels of mitochondrial respiration were unaffected by genotype (Fig. 3E, orthogonal contrast *t* = 0.078, *P* = 0.94, n = 7 per genotype; Supplementary Information 1.3.8) as were multiple independent measures of mitochondrial integrity and function (Fig. S31– 34, Supplementary Information 1.3.8–1.3.9). Together, these data point to reduced function of Complex I without broader phenotypic consequences in individuals heterozygous for incompatible alleles, as previously observed in other models of Complex I dysfunction^57,58^.

Given physiological evidence for some reduction in Complex I function in hybrids heterozygous at *ndufs5* and *ndufa13*, we predicted that there might be an altered frequency of protein complexes incorporating both *X. malinche* mitochondrial proteins and *X. birchmanni* proteins at *ndufs5* and *ndufa13* in F_1_ hybrids. To test this prediction, we took a mass spectrometry-based quantitative proteomics approach. We used stable isotope-labeled peptides to distinguish between the *X. birchmanni* and *X. malinche ndufs5* and *ndufa13* peptides in mitochondrial proteomes extracted from F_1_ hybrids (n = 5, see Methods, Supplementary Information 1.4.1–1.4.4). While endogenous *ndufa13* peptides were not observed frequently enough to quantify accurately, we found consistent deviations from the expected 50-50 ratio of *X. birchmanni* to *X. malinche* peptides for *ndufs5* in F_1_ hybrids, with a significant overrepresentation of matched ancestry at *ndufs5* in the mitochondrial proteome (*t* = 3.96, *P* = 0.016; Fig. 3F; Fig. S35; Supplementary Information 1.4.5). Since we did not observe allele-specific expression of *ndufs5* (Fig 3F; Supplementary Information 1.3.6), this result is consistent with disproportionate degradation of *X. birchmanni*-derived *ndufs5* peptides in the mitochondrial proteome or differences in translation of *ndufs5* transcripts derived from the two species.

### Paired mitonuclear substitutions at contact sites in Complex I

While we can leverage high resolution admixture mapping to pinpoint the nuclear components of the hybrid incompatibility, we cannot take this approach to distinguish among the 37 genes in the swordtail mitochondrial genome because they do not undergo meiotic recombination. To begin to explore the possible mitochondrial partners of *ndufs5* and *ndufa13*, we therefore turned to protein modeling, relying on high quality cryo-EM based structures^59–61^. Although these structures are only available for distant relatives of swordtails, the presence of the same set of supernumerary Complex I subunits and high sequence similarity suggest that using these structures is appropriate (Table S12–S13; Fig. S36–S38; Supplementary Information 1.4.6).

Barring a hybrid incompatibility generated by regulatory divergence (see Supplementary Information 1.3.6), our expectation is that hybrid incompatibilities will be driven by amino acid changes in interacting proteins^62^. We used the program RaptorX^63^ to generate predicted structures of *X. birchmanni* and *X. malinche ndufs5, ndufa13*, and nearby Complex I mitochondrial and nuclear genes, which we aligned to a mouse cryo-EM Complex I structure^59^ (Fig. 4A; Fig. S36–S38; Methods). Using these structures, we visualized amino acid substitutions between *X. birchmanni* and *X. malinche* at the interfaces of *ndufs5, ndufa13* and mitochondrially encoded genes (Fig. S39–S41). While there are dozens of substitutions in the four mitochondrially encoded genes that are in close physical proximity to *ndufs5* or *ndufa13* (Fig. S36; *nd2, nd3, nd4l,* and *nd6*), there are only five cases where amino acid substitutions in either nuclear gene are predicted to be close enough to contact substitutions in any mitochondrial gene, all of which involve *nd2* or *nd6* (Table S14, Fig. 4A; see Fig. S40 for pairwise visualizations of interacting proteins). These paired substitutions in regions of close proximity between mitochondrial- and nuclear-encoded proteins suggest that *nd2* and *nd6* are the genes most likely to be involved in the mitochondrial component of the hybrid incompatibility (Fig. 4A–B; Fig. S41–S44). While there are currently no developed approaches for transgenics in livebearing fish like *Xiphophorus* (Supplementary Information 1.3.1) and Complex I is ill-suited for *in vitro* approaches (being comprised of 45 subunits with an intricate assembly process^64^), manipulating these in-contact substitutions is an exciting direction for future work.

**Fig. 4.**
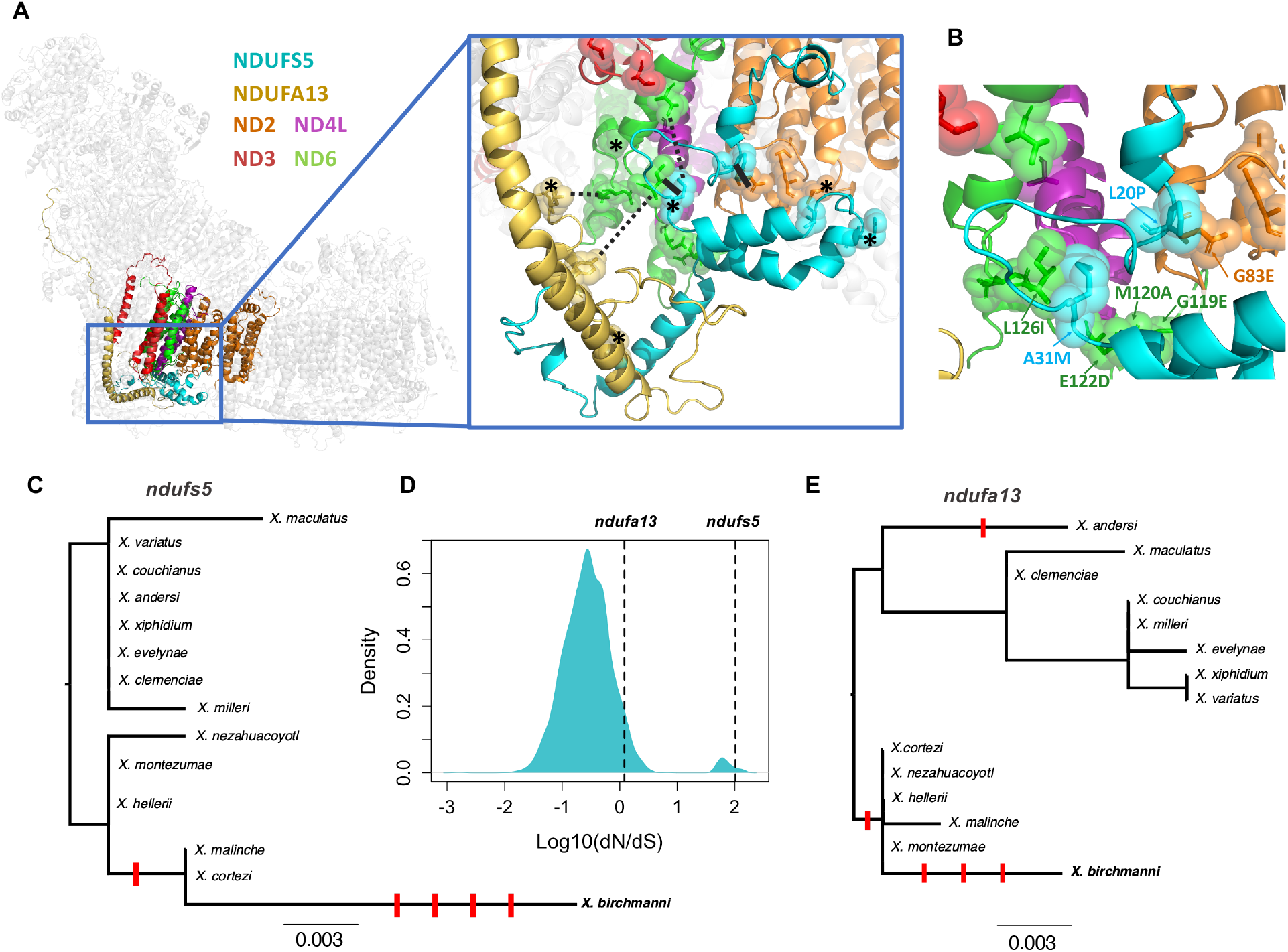
Predicted structures of *Xiphophorus* respiratory Complex I and evolutionary rates of incompatible alleles. (**A**) *Xiphophorus* respiratory Complex I structures generated by RaptorX using alignment to a template mouse cryo-EM structure. Colored protein structures include *ndufs5, ndufa13*, and the four mitochondrially encoded *nd* genes in contact with *ndufs5* or *ndufa13*. Inset shows the surface of predicted contacts between these genes. Solid black lines highlight two areas predicted to be in close contact between interspecific substitutions (alpha carbon distance :: 10 Angstrom for all models), while dashed lines show three additional areas in which there was weaker evidence for a predicted contact based on computational analyses (side chain distance :: 12 Angstrom in at least one model). Asterisks denote residues with substitutions in *X. birchmanni* computationally predicted to affect protein function (Table S16). (**B**) Detailed view of interface between *ndufs5*, *nd2*, and *nd6.* Spheres highlight substitutions between *X. birchmanni* and *X. malinche*. For substitutions predicted to be in close proximity, residues are labeled with letters denoting the *X. malinche* allele, the residue number, and the *X. birchmanni* allele, respectively (see Supplementary Information 1.4.6 and Table S16 for details). (**C**) Gene tree for *ndufs5* generated with RAxML, highlighting an excess of substitutions along the *X. birchmanni* branch. Scale bar represents number of nucleotide substitutions per site. Derived non-synonymous substitutions are indicated by red ticks along the phylogeny. Note that spacing between ticks is arbitrary. (**D**) Distribution of Log_10_ *d*_N_/*d*_S_ between *X. birchmanni* and *X. malinche* across all nuclear genes in the genome with values for *ndufs5* and *ndufa13* highlighted. (**E**) Gene tree for *ndufa13* generated with RAxML, highlighting an excess of substitutions along the *X. birchmanni* branch (as in C).

### Rapid evolution of Complex I proteins

Theory predicts that hybrid incompatibilities are more likely to arise in rapidly evolving genes^4,5,7,15^. Consistent with this hypothesis, *ndufs5* is among the most rapidly evolving genes genome-wide between *X. birchmanni* and *X. malinche* (Fig. 4C–D). Aligning the *ndufs5* coding sequences of *X. birchmanni, X. malinche*, and twelve other swordtail species revealed that all four amino acid substitutions that differentiate *X. birchmanni* and *X. malinche* at *ndufs5* were derived on the *X. birchmanni* branch (Fig. 4C). Phylogenetic tests indicate that there has been accelerated evolution of *ndufs5* on this branch (*d*_N_/*d*_S_ > 99, N = 4, S = 0, codeml branch test *P* = 0.005, Fig. 4C). Similar patterns of rapid evolution are observed at *ndufa13*, which also showed evidence for accelerated evolution in *X. birchmanni* (Fig. 4E; *d*_N_/*d*_S_ = 1.2, N = 3, S = 1, codeml branch test *P* = 0.002). While explicit tests for adaptive evolution at *ndufs5* and *ndufa13* could not exclude a scenario of relaxed selection (Supplementary Information 1.5.1, 1.5.2), our comparisons across phylogenetic scales highlight strong conservation in some regions of the proteins and rapid turnover in others, complicating our interpretation of this test (Fig. S45).

Rapid evolution of *ndufs5* and *ndufa13* could be driven by coevolution with mitochondrial substitutions, a mechanism that has been proposed to explain the outsized role of the mitochondria in hybrid incompatibilities^16,65^. Indeed, there is an excess of derived substitutions in the *X. birchmanni* mitochondrial protein *nd6*, one of the proteins that physically contacts *ndufs5* and *ndufa13* (Table S15; Fig. S42; codeml branch test *P* = 0.005). Moreover, several of the substitutions observed in both mitochondrial and nuclear genes are predicted to have functional consequences (Supplementary Information 1.5.1; Table S16), including ones predicted to be in contact between *ndufs5, ndufa13, nd2*, and *nd6* (Fig. 4A, 4B; Fig. S41).

### Introgression of genes underlying mitonuclear incompatibility

The presence of a mitonuclear incompatibility in *Xiphophorus* is especially intriguing, given previous reports that mitochondrial genomes may have introgressed between species^37^. While *X. malinche* and *X. birchmanni* are sister species based on the nuclear genome, they are mitochondrially divergent, with *X. malinche* and *X. cortezi* grouped as sister species based on the mitochondrial phylogeny^37^ (Fig. 5A–B). As we show, all *X. cortezi* mitochondria sequenced to date are nested within *X. malinche* mitochondrial diversity (Fig. 5B, Fig. S42, S46; Supplementary Information 1.5.3–1.5.4). Simulations indicate that gene flow, rather than incomplete lineage sorting, drove replacement of the *X. cortezi* mitochondria with the *X. malinche* sequence (P < 0.002 by simulation; Fig. 5C; Supplementary Information 1.5.4).

**Fig. 5.**
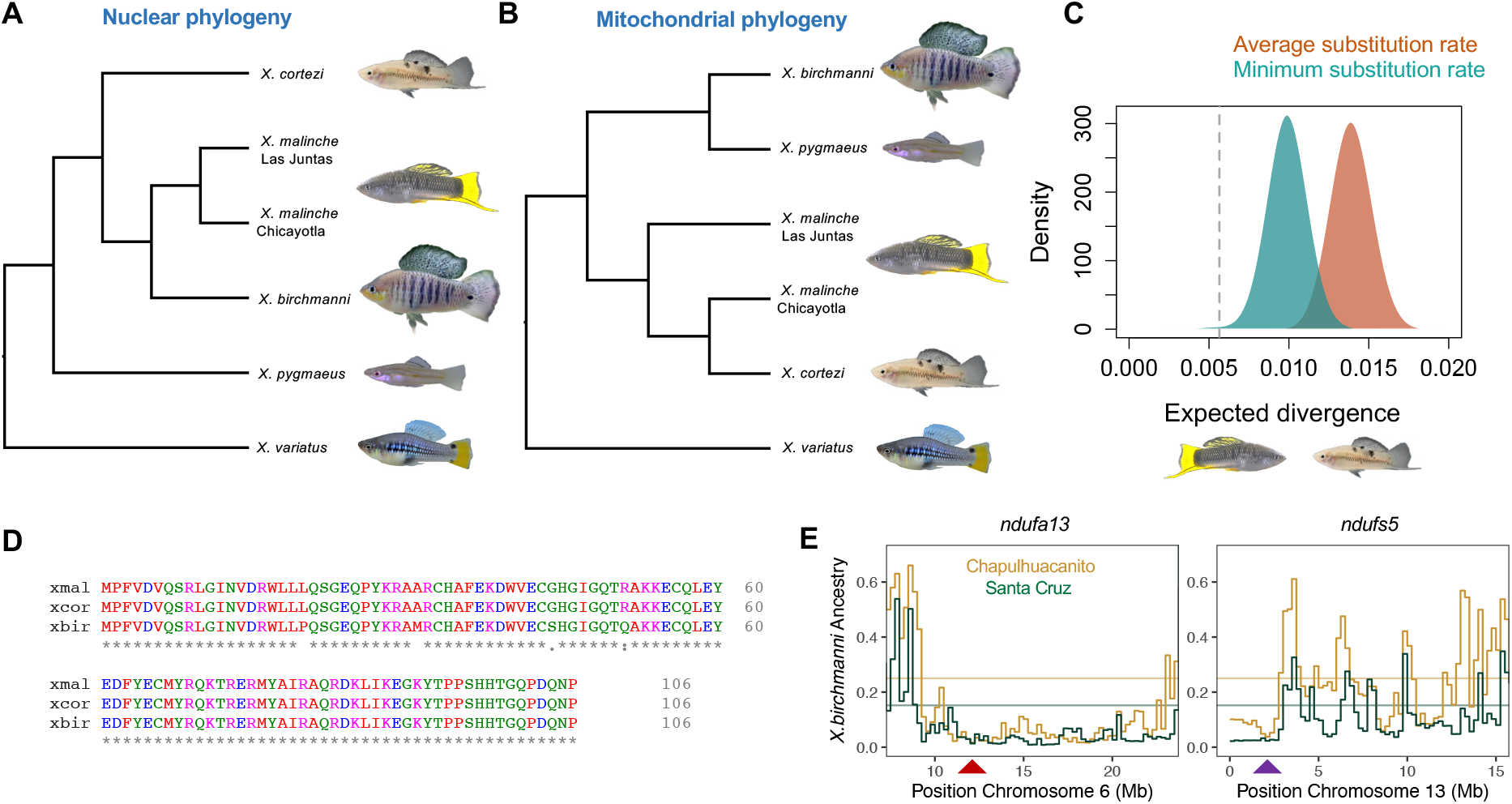
Phylogenetic analysis and ancestry mapping suggest that genes underlying the mitonuclear incompatibility have introgressed from *X. malinche* into *X. cortezi*. (**A**) Nuclear phylogeny of *Xiphophorus* species, showing that *X. birchmanni* and *X. malinche* are sister species^37^. (**B**) Phylogeny constructed from whole mitochondrial genome sequences showing that *X. cortezi* mitochondria are nested within *X. malinche* mitochondrial diversity. (**C**) Results of simulations modeling expected mitochondrial divergence between *X. malinche* and *X. cortezi* in a scenario with no gene flow. Distributions represent pairwise sequence divergence in two sets of simulations. The first set used the average mitochondrial substitution rate observed between *Xiphophorus* species (red), and the second used the minimum mitochondrial substitution rate observed (blue). The dotted line shows observed divergence between mitochondrial haplotypes in *X. malinche* and *X. cortezi*, indicating that past mitochondrial introgression is more consistent with the observed data than incomplete lineage sorting (Supplementary Information 1.5.4). (**D**) Clustal alignment of *ndufs5* sequences shows that *X. malinche* and *X. cortezi* have identical amino acid sequences at *ndufs5*, hinting at possible introgression of this nuclear gene, while *X. birchmanni* is separated from them by four substitutions. Similar patterns are observed for *ndufa13*. Colors indicate properties of the amino acid, asterisks indicate locations where the amino acid sequences are identical. (**E**) Non-mitochondrial parent ancestry is strongly depleted in natural *X. cortezi × X. birchmanni* hybrid populations fixed for the *X. cortezi* mitochondrial haplotype (Fig. S50) at *ndufs5* and *ndufa13*. Arrows show the locations of *ndufa13* and *ndufs5* on chromosome 6 and chromosome 13 respectively, which fall in the center of minor parent ancestry deserts in both independently formed populations, as expected for strong hybrid incompatibilities. Step curves show average *X. birchmanni* ancestry in 250 kb windows, and horizontal lines show the genome-wide average for each population.

The introgression of the mitochondrial genome from *X. malinche* into *X. cortezi* raises the possibility that other Complex I genes may have co-introgressed^66^. Indeed, the nucleotide sequence for *ndufs5* is identical between *X. malinche* and *X. cortezi*, and the sequence of *ndufa13* differs by a single synonymous mutation (although conservation of both genes is high throughout *Xiphophorus*; Fig. S47–S48). These identical amino acid sequences suggest that hybrids between *X. cortezi* and *X. birchmanni* are likely to harbor the same mitonuclear incompatibility we observe between *X. malinche* and *X. birchmanni*, as a result of ancient introgression between *X. malinche* and *X. cortezi* (Fig. 5D; Supplementary Information 1.5.3–1.5.5).

This inference is supported by analysis of ancestry in three contemporary *X. birchmanni* × *X. cortezi* hybrid populations^48^, two of which are completely geographically isolated, indicating that the hybridization events were independent (Fig. S49). We find that all known *X. birchmanni* × *X. cortezi* hybrid populations are fixed for the mitochondrial genome from *X. cortezi* (that originated in *X. malinche*) and also show a striking depletion of non-mitochondrial parent ancestry at *ndufs5* and *ndufa13* (Fig. 5E; Fig. S50; Supplementary Information 1.5.6). This repeated depletion is not expected by chance (Fig. 5E; Supplementary Information 1.5.6, *P* = 0.0001 by simulation) and instead indicates that selection has removed mismatched ancestry in these regions of the genome. These results suggest that the mitonuclear incompatibility observed in *X. birchmanni* × *X. malinche* is also active in hybridizing *X. birchmanni × X. cortezi* populations. This exciting finding shows for the first time that genes underlying hybrid incompatibilities can introgress together, transferring incompatibilities between related species.

## Discussion

What genetic and evolutionary forces drive the emergence of hybrid incompatibilities, especially between closely related species? Theory predicts that hybrid incompatibilities involving multiple genes should be common^6,7^, but with few exceptions^8,11–13^, they remain virtually uncharacterized at the genic level^6^. Here, we identify incompatible interactions in mitochondrial Complex I that causes hybrid lethality in lab and wild populations. Our findings in naturally hybridizing species echoes predictions from theory and studies in lab models^8,11–13^ that protein complexes may be a critical site of hybrid breakdown.

Researchers have proposed mitonuclear interactions as “hotspots” for the emergence of hybrid incompatibilities, given that mitochondrial genomes often experience higher substitution rates between species^18–20^, yet must intimately interact with nuclear proteins to perform essential cellular functions^25,26^. Our findings support this prediction, identifying incompatible interactions with both the *X. malinche* and *X. birchmanni* mitochondria. We also show that there has been exceptionally rapid evolution in both mitochondrial and interacting nuclear genes in *X. birchmanni*, which we speculate may have introduced mutations that are incompatible in hybrids (Fig. 4). Whether driven by adaptation or some other mechanism, our findings support the hypothesis that the coevolution of mitochondrial and nuclear genes could drive the overrepresentation of mitonuclear interactions in hybrid incompatibilities^25,26,65^. More broadly, our results are consistent with predictions that rapidly evolving proteins are more likely to become involved in hybrid incompatibilities than their slowly evolving counterparts^4,5,15^.

Characterizing the incompatibility across multiple scales of organization allowed us to explore the mechanisms through which it acts^67–69^. Our results suggest that in the case of the *X. malinche* mitochondria hybrid lethality is mediated through arrested development *in utero* of individuals with mismatched ancestry at *ndufs5*, while individuals with *ndufa13* mismatch have vascular defects and typically die shortly after birth. Intriguingly, individuals with *ndufa13* mismatch that do survive are much less likely to harbor any *X. birchmanni* alleles at *ndufs5* (even in the heterozygous state; Fig. 2I). Taken together, our results indicate that a subtle three-way interaction overlays two strong pairwise mitonuclear incompatibilities at *ndufs5* and *ndufa13*. Evolutionary biologists have been fascinated by the idea that hybrid incompatibilities may commonly involve three or more genes following theoretical work by Orr^7^ almost thirty years ago, but this question has been challenging to address empirically. Our results highlight how the nuances of actual fitness landscapes may defy simplifying assumptions.

Finally, this mitonuclear incompatibility provides a new case in which the same genes are involved in incompatibilities across multiple species^38,46,70^. However, tracing the evolutionary history of the genes that underlie it adds further complexity to this phenomenon: we found that introgression has resulted in the transfer of genes underlying the incompatibility from *X. malinche* to *X. cortezi*, and evidence from *X. birchmanni × X. cortezi* hybrid populations indicates that the incompatibility is likely under selection in these populations as well. The possibility that hybridization could transfer incompatibilities between species has not been previously recognized, perhaps due to an underappreciation of the frequency of hybridization. The impact of past hybridization on the structure of present-day reproductive barriers between species is an exciting area for future inquiry.

## Methods

### Biological Materials

Wild parental and hybrid individuals used in this study were collected from natural populations in Hidalgo, Mexico (Permit No. PPF/DGOPA-002/19). Artificial F_1_ and F_2_ hybrids were generated using mesocosm tanks as described previously^47^. Caudal fin clips were used as the source for all DNA isolation and for flow cytometry, and liver tissue for RNAseq, respirometry, and proteomic assays were collected following Stanford APLAC protocol #33071.

### Genotyping and local ancestry calling

Genomic DNA was extracted from fin clips and individually barcoded tagmentation based libraries were generated (Supplementary Information 1.1.3). Hybrids were genotyped with low-coverage whole genome sequencing followed by local ancestry inference across the 24 *Xiphophorus* chromosomes and the mitochondrial genome using the *ancestryinfer* pipeline^46,47,50,71^ (Supplementary Information 1.1.3–1.1.4). We converted posterior probabilities for each ancestry state to hard-calls for downstream analysis, using a posterior probability threshold of 0.9, and analyzed ancestry variation across the genome.

### QTL and admixture mapping

The regions interacting with the mitochondrial genome were first identified based on analysis of segregation distortion in 943 F_2_ hybrids generated from F_1_ crosses between *X. malinche* females and *X. birchmanni* males (Supplementary Information 1.1.1 and Langdon et al^48^). Since all hybrids in this artificial cross harbored the *X. malinche* mitochondria, we scanned for regions of exceptionally high *X. malinche* ancestry along the genome (>60% *X. malinche* ancestry), identifying one such region on chromosome 13 and one on chromosome 6 (Fig. 1; see also^48^). Evidence for interactions between these regions and the mitochondrial genome were confirmed using admixture mapping in two hybrid populations that segregated for the mitochondrial haplotype of both species (Supplementary Information 1.1.2): the Calnali Low hybrid population (N = 359) and the Chahuaco falls hybrid population (N = 244). Briefly, we used a partial correlation analysis to identify regions of the genome strongly associated with mitochondrial ancestry, after regressing out genome-wide ancestry to account for covariance in ancestry due to population structure (see ^44^ and Supplementary Information 1.1.5, 1.1.9). Significance thresholds were determined using simulations (Supplementary Information 1.1.5; see also Supplementary Information 1.1.6).

### Estimates of selection on the mitonuclear incompatibility

We used an ABC approach to estimate the strength of selection against the incompatible interaction between the *X. malinche* mitochondrial haplotype and *X. birchmanni* ancestry at the two nuclear genes involved in the hybrid incompatibility: *ndufs5* and *ndufa13* (Supplementary Information 1.2.2). For these simulations, we asked what selection coefficients (0–1) and dominance coefficients (0–1) could generate the observed deviations from the expectation of 50– 50 *birchmanni–malinche* ancestry in F_2_ hybrids at *ndufs5* and *ndufa13* after two generations of selection. We performed 500,000 simulations for each interaction and accepted or rejected simulations based on comparisons to the real data using a 5% tolerance threshold (Supplementary Information 1.2.2). We also evaluated evidence for incompatible interactions with the *X. birchmanni* mitochondrial haplotype (Supplementary Information 1.2.1–1.2.4).

### Developmental staging and genotyping of embryos

To pinpoint when in development the incompatibility between the *X. malinche* mitochondria and *X. birchmanni* nuclear genotypes causes lethality, we collected a dataset on the developmental stages of embryos with different genotype combinations. Whole ovaries were removed from pregnant females and embryos were individually dissected. Each embryo was assigned a developmental stage ranging from 1–11 based on established protocols for poeciliid embryos^52^. Unfertilized eggs were excluded from analysis. Following staging, individual embryos (N = 296) were genotyped as described above and in Supplementary Information 1.3.1. We tested for significant differences in developmental stage between siblings with compatible and incompatible genotype combinations using a two-sided two-sample t-test (Supplementary Information 1.3.1) and examined differences in ancestry between large groups of siblings that varied in their developmental stages (Supplementary Information 1.1.7). We also collected data on embryonic stage and variability between siblings in embryonic stage from both pure parental species (Supplementary Information 1.3.1). We used a different approach to pin-point the timing of *ndufa13* lethality given that it appeared to act postnatally (Supplementary Information 1.3.3).

### Embryo Respirometery and Morphometrics

To study the mechanisms of *ndufs5*- and *ndufa13*-driven lethality, we performed oxygen consumption measurements on F_2_ embryos in a Loligo plate respirometer (Supplementary Information 1.3.4). Embryo were dissected from mothers and transferred to wells of a 24-well plate, where their oxygen consumption was measured over 60 minutes. The measurement was then repeated in media dosed with 5 μM rotenone to test sensitivity to Complex I inhibition, after which the embryos were video recorded and photographed for morphometrics in ImageJ. We used linear models to test the effect of *ndufs5* genotype, *ndufa13* genotype, and individual standard length on a number of variables, controlling for batch effects (Supplementary Information 1.3.4–1.3.5).

### Mitochondrial respirometry

To further evaluate mitochondrial function in individuals heterozygous for the mitonuclear incompatibility (Supplementary Information 1.3.6–1.3.7), we conducted respirometry assays on *X. birchmanni*, *X. malinche*, and hybrid individuals that had the *X. malinche* mitochondria and were heterozygous for the nuclear components of the hybrid incompatibility (N=7 of each genotype). Mitochondria were isolated from whole liver tissue and mitochondrial respiration was quantified using the Oroboros O2K respirometry system (Fig. S30)^72^. A step-by-step description of this protocol and methods used to calculate respiratory flux control factors is outlined in Supplementary Information 1.3.8. We complemented the results of these respirometry experiments with measures of mitochondrial membrane potential using a flow cytometry-based approach (Supplementary Information 1.3.9).

### Parallel reaction monitoring proteomics

For Parallel Reaction Monitoring (PRM) with mass spectrometry, we used a similar approach to that used for respirometry to isolate whole mitochondria from five F_1_ hybrids (which harbored *X. malinche* mitochondria). This approach is described in detail in Supplementary Information 1.4.1. Briefly, we designed heavy-labeled peptides to distinguish between the *X. birchmanni* and *X. malinche* copies of NDUFS5 and NDUFA13, facilitating quantification of the peptides of interest in the mitochondrial proteome (Supplementary Information 1.4.2). Mitochondrial isolates were prepared for mass spectrometry and combined with heavy-labeled peptides in known quantities (see Supplementary Information 1.4.3), then submitted to Orbitrap mass spectrometry with separation with UPLC and parallel reaction monitoring (PRM) for ion selection. The protocol for mass spectrometry and PRM is described in detail in Supplementary Information 1.4.4.

To analyze the results, the focal peptide’s spectral peak were identified based on the peak of the heavy labeled spike-in peptide. We focused analysis on the NDUFS5 peptide WLL[L/P]QSGEQPYK since other endogenous peptides were below the expected sensitivity limits of our PRM protocol (Supplementary Information 1.4.5). We normalized the intensity of the NDUFS5 peptide based on the known spike-in quantity, and quantified the proportion of NDUFS5 in each F_1_ individual derived from *X. malinche* versus *X. birchmanni* (Supplementary Information 1.4.5). We tested whether these ratios significantly deviated from the 50-50 expectation for F_1_ hybrids using a two-sided one-sample *t*-test.

### Complex I protein modeling

Mapping results allowed us to identify *ndufs5* and *ndufa13* as *X. birchmanni* genes that interact negatively with *X. malinche* mitochondrial genes. We used a protein-modeling based approach with RaptorX (http://raptorx.uchicago.edu) to identify the mitochondrial genes most likely to interact with *ndufs5* and *ndufa13* (see Supplementary Information 1.4.6). Using the mouse Cryo-EM structure (PDB ID 6G2J) of Complex I, we identified proteins in contact with *ndufs5* and *ndufa13*, which included several mitochondrial (*nd2, nd3, nd4l, nd6*) and nuclear (*ndufa1*, *ndufa8*, *ndufb5*, *ndufc2*) genes. We then used RaptorX to predict structures for both the *X. birchmanni* and *X. malinche* versions of the proteins. In addition, we evaluated the robustness of these predictions to choice of Cryo-EM template; see Supplementary Information 1.4.6.

### Analysis of evolutionary rates

Comparison of predicted protein sequences from *ndufs5, ndufa13*, and mitochondrial genes of interest (*nd2* and *nd6*) revealed a large number of substitutions between *X. birchmanni* and *X. malinche*. We calculated *d*_N_/*d*_S_ between *X. birchmanni* and *X. malinche* for all annotated protein coding genes throughout the genome and found that both *ndufs5* and *ndufa13* have rapid protein evolution (Fig. 4D; Supplementary Information 1.5.1). Examining these mutations in a phylogenetic context revealed that many substitutions in *ndufs5*, *nudfa13*, and *nd6* were derived in *X. birchmanni*. We tested for significant differences in evolutionary rates on the *X. birchmanni* lineage and for predicted functional impacts of these substitutions; these analyses are described in Supplementary Information 1.5.1.

### Tests for ancient introgression

Previous work had indicated that the mitochondrial phylogeny in *Xiphophorus* is discordant with the whole-genome species tree^37^. Specifically, although *X. birchmanni* and *X. malinche* are sister species based on the nuclear genome, *X. malinche* and *X. cortezi* are sister species based on the mitochondrial genome. We used a combination of PacBio amplicon sequencing of 10 individuals (2 or more per species, Supplementary Information 1.5.3) and newly available whole-genome resequencing data to confirm this result and polarize the direction of the discordance by constructing maximum likelihood mitochondrial phylogenies with the program RAxML^74^. We performed similar phylogenetic analyses of the nuclear genes that interact with the *X. malinche* mitochondria (*ndufs5* and *ndufa13*; Supplementary Information 1.5.3). Combined with phylogenetic results, simulation results suggest that gene flow from *X. malinche* into *X. cortezi* is the most likely cause of the discordance we observe between the mitochondrial and nuclear phylogenies (Supplementary Information 1.5.3–1.5.4). Since *X. malinche* and *X. cortezi* are not currently sympatric, this suggests ancient gene flow between them (Supplementary Information 1.5.5).

### Contemporary hybridization between *X. birchmanni* and *X. cortezi*

To investigate the possibility that hybrids between *X. birchmanni* and *X. cortezi* share the same mitonuclear incompatibility as observed in hybrids between *X. birchmanni* and *X. malinche* (Supplementary Information 1.5.6), we took advantage of genomic data from recently discovered hybrid populations between *X. birchmanni* and *X. cortezi*^75^. Using data from three different *X. birchmanni* × *X. cortezi* populations and a permutation-based approach, we asked whether ancestry at *ndufs5* and *ndufa13* showed lower mismatch with mitochondrial ancestry than expected given the genome-wide ancestry distribution. This analysis is described in detail in Supplementary Information 1.5.6.

### Animal Care and Use

All methods were performed in compliance with Stanford Administrative Panel on Laboratory Animal Care protocol #33071.

## Data Availability

Raw sequencing reads used in this project are available under NCBI SRA Bioprojects PRJNA744894, PRJNA746324, PRJNA610049, and PRJNA745218. Mass spectrometry data are available on PRIDE (accession pending), and all other datasets necessary to recreate the results of the publication are available on Dryad (accessions pending).

## Code Availability

All new customs scripts used to generate results will be made available on Github at https://github.com/Schumerlab/mitonuc_DMI and

https://github.com/Schumerlab/Lab_shared_scripts.

## Acknowledgements

We thank Peter Andolfatto, Stepfanie Aguillon, Yaniv Brandvain, Jenn Coughlan, Hunter Fraser, Yuki Haba, Nitin Phadnis, Molly Przeworski, Ken Thompson and members of the Schumer lab for helpful discussion and/or feedback on earlier versions of this manuscript. We thank Alexa Pollock for help performing rotenone trials. We thank the Federal Government of Mexico for permission to collect fish. Stanford University and the Stanford Research Computing Center provided computational support for this project. We thank the Vincent Coates Foundation Mass Spectrometry Laboratory, Stanford University Mass Spectrometry (RRID:SCR_017801) for technical and experimental support.

## Author Contributions

B.M.M., D.L.P, and M. Schu, designed the project; B.M.M., C.Y.P., E.N.K.I., S.M.B, A.E.D, A.M, J.J.B, K.K., F.L., R.M., K.S., O.H.-P., J.C.H., A. M. and M. Schu. collected data; B.M.M., C.Y.P., Q.L.K., F.L., J.C.H., A.M., and M. Schu. performed analyses; D.L.P., T.R.G., R.D.L., C.G., R.C.-D., J.F. and M. Scha. provided expertise and technical support.

## Materials & Correspondence

Correspondence and requests for materials should be directed to B.M.M. or M. Schu.

