## Supplementary Information for "A Lethal Genetic Incompatibility between Naturally Hybridizing Species in Mitochondrial Complex I"

### 1. Supplementary Information

#### 1.1. Detection of a mitonuclear hybrid incompatibility through ancestry mapping

##### 1.1.1. <u>$F_2$ crosses between *X. birchmanni and X. malinche*</u>

We previously developed a large panel of $F_2$ hybrids to map the genetic basis of sexually selected traits in *X. birchmanni* × *X. malinche* hybrids. Because sexually selected traits are only expressed in male swordtails, we previously analyzed ancestry for 568 male $F_2$s[1], generated from $F_1$ crosses between *X. malinche* females and *X. birchmanni* males. Attempts to produce reciprocal $F_1$ crosses between female *X. birchmanni* and male *X. malinche* have been largely unsuccessful. Specifically, crosses of female *X. birchmanni* to *X. malinche* males have resulted in severely reduced offspring viability compared to the other cross direction. Offspring were born underdeveloped with only two out of 23 pregnancies producing offspring that survived past the two-week mark; median brood size was three compared to 20 in the reciprocal cross.

We revisited $F_2$ data generated from the *X. malinche* female to *X. birchmanni* male cross direction and analyzed the entire set of 943 adult $F_2$ individuals (see also Langdon et al. [2]). We performed local ancestry inference using a hidden Markov model (HMM) approach with the *ancestryinfer* pipeline[1,3–5]. We set the prior for admixture proportions to 0.5 and the prior for generations since initial admixture to 2. We used a posterior probability threshold of 0.9 to convert posterior probabilities for ancestry states into hard-calls. Past work in this system has indicated that we expect high accuracy in local ancestry inference in this cross, especially with early generation hybrids[1,4].

Due to the cross design, we expected $F_2$ hybrids to derive on average 50% of their genome from both parental species, and indeed this is what we observe. However, we identified several regions of the genome with substantial deviations from this 50-50 expectation. One region on chromosome 13 showed the most extreme deviations in ancestry, with 67% of alleles in this region derived from the *X. malinche* parent (p-value from binomial test $< 10^{-280}$). Closer examination of genotypes in this region indicated that only one of 937 successfully genotyped individuals in this region was homozygous for *X. birchmanni* ancestry. This pattern strongly deviates from expectations under Hardy-Weinberg equilibrium even when these expectations are calculated using the observed *X. birchmanni* allele frequency in this region (29% *X. birchmanni* ancestry; $\chi^2 = 79$; p-value with one degree of freedom $< 10^{-18}$).

One possible cause of deviations in expected ancestry is an excess of genotyping errors in a particular region of the genome. We found no evidence for an excess of ancestry transitions in this region in $F_2$ hybrids or switch errors in pure parental individuals compared to other regions of the genome (Fig. S51), suggesting that ancestry deviations in this region are the product of biological rather than technical processes.

Another common cause of segregation distortion in hybrids is hybrid incompatibilities that reduce viability. Notably, homozygous *X. birchmanni* ancestry was almost entirely absent from the chromosome 13 region in $F_2$ hybrids. Based on naïve expectations, we would predict that even in the case of a dominant, lethal hybrid incompatibility, some individuals with homozygous *X. birchmanni* ancestry would survive, simply because by chance some individuals would also have homozygous *X. birchmanni* ancestry at the interacting loci. In such cases, hybrid individuals would escape incompatibility by recapitulating the *X. birchmanni* genotype at both loci. To test this intuition, we performed 100 simulations using the hybridization simulator admix'em[6]. We simulated a dominant lethal nuclear-nuclear hybrid incompatibility, with each of the two interacting loci randomly placed on one of 24 chromosomes. We used admix'em to generate $F_1$ and $F_2$ hybrids that underwent viability selection as a function of their genotypes at the hybrid incompatibility. We extracted genotypes at the simulated incompatibility and counted the number of individuals with homozygous *X. birchmanni* ancestry at each locus. We recorded the number of individuals with homozygous *X. birchmanni* ancestry at each of the loci involved in the hybrid incompatibility. The results of these simulations are shown in Fig. S52.

The results of these simulations confirm our prediction that the observed data on chromosome 13 is not consistent with patterns expected for a nuclear-nuclear hybrid incompatibility. We find that in our simulations on average ~100 individuals (or ~10% of simulated individuals) have homozygous *X. birchmanni* ancestry at the loci involved in the lethal nuclear-nuclear incompatibility. This result is expected from first principles because by chance a subset of individuals will be homozygous *X. birchmanni* at both nuclear loci and escape the incompatibility. In contrast, in the real data, only one individual was homozygous for *X. birchmanni* ancestry in the chromosome 13 region (0.1% of individuals). This caused us to explore the hypothesis that our data may be more consistent with a lethal mitonuclear incompatibility, since all individuals harbored *X. malinche* mitochondria (see *Supplementary Information 1.1.5*).

Guided by the results of these simulations, we further explored whether any other regions in our dataset of $F_2$ hybrids had unexpectedly few individuals with homozygous *X. birchmanni* genotypes. These regions would also be candidates for involvement in mitonuclear incompatibilities. We focused on analysis of five previously described segregation distorters[2]. Of these regions, only one had unexpectedly high depletion of homozygous *X. birchmanni* genotypes. Across a large region of chromosome 6, only ~3% of individuals harbored homozygous *X. birchmanni* ancestry (Fig. 1D). We treated this region as an additional candidate for involvement in a mitonuclear incompatibility (see *Supplementary Information 1.1.5*; Fig. 1D).

1.1.2. <u>Collections from natural hybrid populations</u>

Consistent with our data pointing to the presence of a mitonuclear incompatibility, nearly all of the natural hybrid populations we have studied to date are fixed for one of the parental species' mitochondrial haplotypes. Invariably all populations that are fixed for mitochondrial haplotypes are fixed for the mitochondria that matches their major parent ancestry (Fig. S5; Aguazarca: N=126, fixed for *X. birchmanni*, Acuapa: N=117, fixed for *X. birchmanni*, Tlatemaco: N=126, fixed for *X. malinche*). These three populations occur in three different river systems (Fig. S5), representing independent instances of hybridization between *X. birchmanni* and *X. malinche*. Only one previously studied population, the Chahuaco Falls population on the Río Calnali (Fig. S5), which derives the majority of its genome from the *X. malinche* parental species, was segregating for both mitochondrial haplotypes (7 ± 2% *X. birchmanni* haplotype).

We were interested in identifying a hybrid population for admixture mapping with a more moderate frequency of both *X. birchmanni* and *X. malinche* mitochondrial haplotypes to maximize our power in mapping analyses. We thus sequenced samples from several additional populations on the Río Calnali between the Chahuaco Falls and Aguazarca populations (Fig. S5) to assess the frequency of the two mitochondrial haplotypes. We found that the Calnali Low population had an *X. birchmanni* mitochondrial haplotype frequency of ~60%, making it a promising target for admixture mapping. Since the Calnali Low population is separated from the Chahuaco Falls population by a stretch of river and a large waterfall (~50 m), these populations also represent semi-independent samples for mapping.

Targeting sampling efforts on the Calnali Low population, we were able to collect 359 adult individuals for subsequent mapping. We caught adult fish using baited minnow traps in the spring and summer of 2017, 2018 and summer and fall of 2021. Fish from Chahuaco Falls had been previously collected and sequenced for another study[5]. We anesthetized fish with tricaine methanesulfonate (Stanford University IACUC protocol #33071), photographed them for phenotyping, and took fin clips of the posterior dorsal quarter of the caudal fin with sterilized dissection scissors. Fin clips were stored in 95% ethanol at room temperature or 4°C until DNA extraction. An additional set of 58 fish collected from the Río Calnali in winter 2021 were housed in lab facilities for 6 months before being genotyped and used in physiological assays (see *Supplementary Information 1.3.8 Respirometry for OXPHOS Complex activity in parents and hybrids*). These fish were marked individually with color-coded elastomer tags for future identification (Northwest Marine Technologies; Stanford University IACUC protocol #33071).

##### 1.1.3. Low coverage library preparation and sequencing

We extracted DNA from fin clips using the Agencourt DNAdvance bead-based protocol (Beckman Coulter, Brea, California), following the manufacturer's instructions except that we used half reactions. We quantified and diluted the extracted DNA to 2.5 ng/ul. We then prepared tagmentation-based whole genome libraries for low coverage sequencing. Briefly, we used the Illumina Tagment DNA TDE1 Enzyme and Buffer Kits (Illumina, San Diego, CA) to enzymatically shear DNA. Sheared DNA samples were amplified in PCR reactions with dual indexed custom primers for 12 cycles. Amplified PCR reactions were pooled and purified using 18% SPRI magnetic beads. We quantified libraries with a Qubit fluorometer (Thermofisher Scientific, Wilmington, DE) and assessed library fragment size on a TapeStation (Agilent, Santa Clara, CA). We pooled libraries and sent them to Admera Health Services (South Plainfield, NJ) for sequencing 150 bp paired-end reads on a HiSeq 4000 (Illumina, San Diego, CA). All data has been deposited on the NCBI sequence read archive (SRA BioProject PRJNA744894).

##### 1.1.4. Local ancestry inference and mitochondrial haplotype inference

We previously developed accurate local ancestry inference approaches for the nuclear genome of *X. birchmanni* × *X. malinche* hybrids[4,5]. For this project, we also developed ancestry inference for the mitochondrial genome of both species and evaluated the accuracy of mitochondrial ancestry inference. To define ancestry informative sites in the mitochondrial genomes of both species, we took advantage of our *de novo* assemblies for these species and previously collected high coverage genome sequence data.

To identify contigs in our *de novo* assemblies corresponding to the mitochondrial genome, we downloaded the mitochondrial genome assembly from NCBI for *X. maculatus* (GenBank accession: AP005982.1) and used the *blastn* algorithm to identify contigs in our four *de novo* assemblies corresponding to the mitochondrial genome (one 10x based assembly for each *X. birchmanni* and *X. malinche*[5], a PacBio based assembly for *X. birchmanni*, and an Illumina-based assembly for *X. malinche*). We identified contigs that aligned to the full length of the *X. maculatus* mitochondrial genome in the *X. birchmanni* PacBio based assembly and the *X.*

*malinche* Illumina-based assembly. We also used PacBio-based amplicon sequencing to verify these sequences in several individuals (see *Supplementary Information 1.5 Phylogenetic analyses*).

We next used these mitochondrial *de novo* assemblies from *X. birchmanni* and *X. malinche* combined with population level sequence data to identify candidate ancestry informative sites that distinguished mitochondrial haplotypes of the two species. We used bwa[7], Picard tools[8], and GATK[9] as described previously[10] to map reads and perform variant calling on 26 *X. birchmanni* individuals and 4 *X. malinche* individuals that we had previously sequenced (SRP130891; SRP018918). The differences in sampling effort between the two species are reasonable given a ~4 fold difference in the level of genetic diversity in the two species (*X. birchmanni:* 0.12% per basepair, *X. malinche*: 0.03%).

Based on this analysis, we identified fixed differences between mitochondrial haplotypes in our samples of *X. birchmanni* and *X. malinche*. We next wanted to verify that these sites were ancestry informative in a larger population sample. We used low-coverage whole genome sequence data from parental individuals (124 from two parental *X. birchmanni* populations and 29 from two parental *X. malinche* populations) and evaluated the frequency of each identified ancestry informative site in this population sample. We removed sites with <98% frequency difference between *X. birchmanni* and *X. malinche* in the population sample, resulting in 24 ancestry informative sites that distinguished the two species' mitochondrial haplotypes (or 1.5 per kb). While mitochondria do not recombine in vertebrates, for the purposes of running the HMM, we set the recombination rate for the region to a low value ($1^{-9}$ cM/bp).

With mitochondrial ancestry informative sites in hand, we used our previously developed *ancestryinfer* pipeline[3,4] to infer mitochondrial ancestry in hybrids from the Tlatemaco (N=126), Acuapa (N=117), Aguazarca (N=126), Chahuaco Falls (N=244) and Calnali Low hybrid populations (N=359). As an ad-hoc quality check we determined that no individuals analyzed were called as heterozygous for ancestry (posterior probability > 0.9 for the heterozygous ancestry state) at ancestry informative sites in the mitochondria and no individuals analyzed harbored mitochondrial ancestry transitions. Moreover, we confirmed that all lab-bred $F_1$ and $F_2$ hybrids were inferred to have *X. malinche* mitochondrial ancestry as dictated by the cross design. Finally, we confirmed that the *ancestryinfer* pipeline accurately inferred mitochondrial ancestry for a set of 38 *X. birchmanni* and 8 *X. malinche* parental individuals that were not included in the training set used to define ancestry informative sites.

For analyses of mitochondrial ancestry in *X. birchmanni* × *X. cortezi* hybrids (see below) we repeated the above described steps except that we used 8 high coverage individuals in initial definition of ancestry informative sites for *X. cortezi* and 42 individuals from one *X. cortezi* source population to refine ancestry informative sites. This resulted in 35 mitochondrial ancestry informative sites that distinguished *X. birchmanni* and *X. cortezi* (or 2 per kb).

1.1.5. Admixture mapping

Mitochondrial ancestry segregates in only two of the *X. birchmanni* × *X. malinche* populations that we have sampled to date. Segregation of the mitochondria in the Chahuaco Falls and Calnali Low populations gives us the opportunity to directly map any loci that interact with the mitochondria to generate a mitonuclear hybrid incompatibility. Based on our segregation distortion results, we had good reason to believe that an interacting locus would be detected on the 5' end of chromosome 13 and the center of chromosome 6, but we scanned for interacting loci across the genome.

We performed admixture mapping using mitochondrial ancestry as the phenotype of interest. Selection that removes incompatible genotype combinations generates unexpectedly high correlations in ancestry between physically unlinked loci, and several methods can be used to scan for these incompatible interactions, particularly if hypotheses exist regarding which loci are involved[2,11]. In our case, we used a partial correlation approach to identify potential nuclear interactors with the mitochondria[11]. Specifically, we asked about the correlation between nuclear and mitochondrial genotypes after accounting for the covariance expected given each individual's admixture proportion. For computational efficiency, we thinned our set of 629,525 ancestry informative sites to retain 1 every 25 kb, yielding a set of 24,607 sites for mapping analyses. Because admixture linkage disequilibrium (LD) decays over ~1 Mb in these hybrid populations[5] we do not expect to miss any signals with this approach. For each focal ancestry informative site, we recorded the *p*-value of a partial correlation analysis between the mitochondrial and nuclear genotype at the focal site while controlling for genome-wide admixture proportion, implemented using the 'ppcor' package in R[12]. We performed this analysis separately for the Calnali Low and Chahuaco Falls populations.

To determine our significance threshold for admixture mapping, we used a simulation-based approach. Since genome-wide ancestry is strongly correlated with mitochondrial ancestry, we could not simply shuffle the phenotype (mitochondrial haplotype) onto the genotypes. Instead, we used genome-wide ancestry to simulate mitochondrial haplotypes and repeated the analysis as described above 500 times. We used the following steps to do so:

1. For each individual, we used the observed proportion of the genome derived from *X. malinche* to weight a draw from a random binomial in R. If the draw from a random binomial returned zero, we assigned that individual an *X. birchmanni* mitochondrial haplotype for the simulation. If the draw returned one, we assigned that individual a *X. malinche* haplotype for the simulation.
2. We repeated this process until all individuals had been assigned a mitochondrial haplotype.
3. With these simulated mitochondrial haplotypes, we repeated our admixture mapping procedures as described above.
4. We identified and recorded the lowest p-value between the simulated mitochondrial haplotype and nuclear genotype genome-wide in the null simulation.
5. We repeated this process 500 times.

Based on the distribution of p-values observed by chance in simulations, we set the significance threshold for our empirical data to a $-\text{Log}_{10}$ p-value threshold of 5, corresponded to a false positive rate (FPR) of approximately 10%. The threshold corresponding to a more stringent FPR of 5% was a $-\text{Log}_{10}$ p-value threshold of 6.

To delineate QTL regions for further analysis, we identified the maximum $-\text{Log}_{10}$ p-value in all regions that exceed (or approached) our genome-wide significance threshold. We conservatively treated all sites that had $-\text{Log}_{10}$ p-values that were $\pm$ 2 of the peak $-\text{Log}_{10}$ p-value in a region as part of the associated interval for further investigation. For example, for chromosome 13 the peak $-\text{Log}_{10}$ p-value was 15, and we included all sites with $-\text{Log}_{10}$ p-value greater than or equal to 13 as part of the QTL interval of interest (chromosome 13: 2.026 Mb – 2.097 Mb).

We also performed admixture mapping in the Chahuaco Falls hybrid population. While the Chahuaco Falls population segregates for mitochondrial ancestry, there are fewer individuals with *X. birchmanni* haplotypes (7% of our sample), reducing our power to map associations in this population. Nevertheless, we detected an association between mitochondrial ancestry and chromosome 13 ancestry concordant with that found in Calnali Low (Fig. S4).

1.1.6. <u>Evidence of selection from natural hybrid populations</u>

Admixture mapping in the Calnali Low population that segregates for mitochondrial haplotypes from both *X. birchmanni* and *X. malinche* identified several regions associated with mitochondrial ancestry. To evaluate whether these regions were under selection in other natural hybrid populations, we took advantage of three other *X. birchmanni* × *X. malinche* hybrid populations (Fig. S5) that have fixed mitochondrial haplotypes from one of the parental species. The Tlatemaco population has fixed the *X. malinche* mitochondrial haplotype while the Acuapa and Aguazarca populations have fixed the *X. birchmanni* mitochondrial haplotypes (Fig. S5). We reasoned that given that the mitochondrial haplotypes had fixed for one ancestry type in these populations, strong selection against mitonuclear incompatibilities should have driven the removal of incompatible genotype combinations from natural hybrid populations (Fig. 1). We thus scanned for regions within the admixture mapping quantitative trait loci (QTL) that had been purged of non-mitochondrial parent ancestry in all three of these *X. birchmanni* × *X. malinche* hybrid populations with fixed mitochondrial ancestry.

Likely due to the independent histories of recombination in each of these populations, only a small region is fixed or nearly fixed for mitochondrial-parent ancestry (defined as ≥ 95% mitochondrial parent frequency) in all three hybrid populations on chromosome 13 (Fig. S6). This region precisely overlaps with the three genes in the 77 kb admixture mapping region.

We evaluated the probability that all three hybrid populations would have purged non-mitochondrial parent ancestry in this region by chance using simulations. To do so, we permuted our real data and recorded cases in which the windows in the permuted datasets were ≥ 95% mitochondrial parent frequency in all three populations. We used two different approaches for permutation. First, we randomly drew 50 kb windows from the empirical data for each hybrid population and repeated this procedure 10,000 times. Based on this analysis, the probability that all three populations will have greater than 95% mitochondrial parent ancestry in any region by chance was very low (p-value by simulation $3 \times 10^{-4}$). However, by randomly drawing windows from our dataset we disrupt patterns of correlation in ancestry between adjacent windows due to extensive ancestry LD in hybrid populations. We thus took a second approach for permutations, where we shifted window labels independently in two of the three populations. In each permutation step, we shifted window labels by 5 Mb in population 2 and 10 Mb in population 3 (admixture LD decays over <5 Mb in all hybrid populations). For windows at the edge of a chromosome, labels were transferred to the subsequent chromosome (i.e. the last window in chromosome 1 was labeled as a window on chromosome 2). This re-labelling-based permutation maintains LD structure and allows us to ask how frequently windows with the same label had less than or equal to the observed mitochondrial parent frequency at *ndufs5* in all three permuted populations. We repeated this procedure until the labels for population 2 returned to the original labels, resulting in data from 13,503 permutations. Using this approach, we still find that the ancestry patterns we observe are unexpected by chance (P = $1.49 \times 10^{-5}$ by simulation). These simulation results indicate that this desert of minor parent ancestry within our QTL region is unexpected by chance, underscoring the fact that selection has purged non-mitochondrial parent ancestry in this region in multiple independently formed natural hybrid populations.

We also evaluated ancestry in natural hybrid populations in the region surrounding *ndufa13* on chromosome 6. In contrast to *ndufs5*, we only expect to observe selection on *ndufa13* in populations that have fixed the *X. malinche* mitochondrial haplotype, as *ndufa13* does not appear to be incompatible with the *X. birchmanni* mitochondrial haplotype (Fig. 1). Only one population, the Tlatemaco population, has fixed the *X. malinche* mitochondria. In this population, we find that *X. birchmanni* ancestry has been purged in the region surrounding *ndufa13* (Fig. S6). Compared to the distribution of ancestry at other sites across the genome in the Tlatemaco population, ancestry at *ndufa13* is unusual, falling in the upper 0.1% tail of *X. malinche* ancestry genome-wide. This extreme ancestry is suggestive of selection on *ndufa13* in this hybrid population.

##### 1.1.7. Fine mapping the chromosome 13 region using comparisons of siblings

We found that embryos with *X. malinche* mitochondria and homozygous *X. birchmanni* ancestry in the QTL region of chromosome 13 experience arrested embryonic development (Supplementary Information 1.3.1; Fig. 2). This pattern appears to be specific to ancestry in the chromosome 13 region and is not detected in other mapped mitonuclear QTL (Fig. S10-S12). Specifically, embryos with *X. malinche* mitochondria and homozygous *X. birchmanni* ancestry at the chromosome 13 region did not progress past developmental stage 7 and substantially lagged the developmental pace of their unaffected siblings (Supplementary Information 1.3.1; Fig. 2). This observation led us to explore a complementary approach for fine-mapping the mitonuclear QTL on chromosome 13.

As expected, all mothers with *X. malinche* mitochondria were either homozygous *X. malinche* or heterozygous along the associated chromosome 13 region. Of the families where we sequenced mothers and embryos, there were four mothers with *X. malinche* mitochondria, heterozygous ancestry at chromosome 13, and many developing embryos (≥20). We reasoned that if females mated with a male heterozygous for *X. birchmanni* ancestry at the chromosome 13 QTL as well, approximately 25% of their offspring would have incompatible genotypes. Moreover, comparisons among these embryos could be informative, as they shared a developmental environment and much of their genome.

We sequenced a total of 86 embryos from these broods and developmentally staged all of them (see Supplementary Information 1.3.1). We identified embryos that developmentally lagged the most developed embryo in the brood by greater than or equal to one developmental stage[13]. We then analyzed ancestry in the developmentally lagged embryos compared to their siblings with no developmental lag (9-11 embryos with developmental lag per brood). The results of this analysis are shown in Fig. 2 and indicate that these embryos have substantially higher homozygous *X. birchmanni* ancestry along a narrow region of chromosome 13 than their unaffected siblings. Strikingly, this interval almost perfectly overlapped with the admixture mapping QTL region (Supplementary Information 1.1.5), as well as the region identified by analysis of selection on hybrid populations (see Fig. S6; Supplementary Information 1.1.6), and contains only three genes. This analysis is complementary to our other mapping and population genetic approaches, and allows us to confidently focus on a small number of genes as candidates on chromosome 13 for mitonuclear interactions.

It is interesting to note that only 66% of developmentally lagging individuals were homozygous for *X. birchmanni* ancestry at 2.07 Mb on chromosome 13. This could indicate that there are other genetic or environmental factors that can cause developmental lag in *X. birchmanni* x *X. malinche* hybrids, or that some heterozygous individuals may be affected by the incompatibility. Indeed, both mechanisms are possible. The majority of individuals with lagging development that are not homozygous *X. birchmanni* at the ancestry peak on chromosome 13 were heterozygous; only one individual was homozygous *X. malinche.*

While individuals inherit their mitochondrial sequences from their mother, they also transiently inherit maternally deposited mRNAs and proteins. Past research has shown that maternal effects, mediated via deposited mRNAs, can generate abnormal developmental phenotypes in fish[14–16]. The availability of data for groups of siblings (≥20 per family) allowed us to exclude these types of maternal effects as the cause of the developmental delay phenotype. Notably, because siblings inherit the same genotypes of maternally deposited mRNAs and proteins, we would expect any developmental deficits attributable to maternal effects to be shared across siblings. Instead, we see a strong dependence on chromosome 13 ancestry, with only between 43-48% of embryos in each brood exhibiting the developmental delay phenotype.

##### 1.1.8. Evaluation of all genes in the fine-mapped interval on chromosome 13

Both admixture mapping in hybrid populations and case-control mapping in siblings pinpoint a small region of chromosome 13 associated with the mitonuclear incompatibility (see *Supporting Information 1.1.5-1.1.7*). This region contained three genes: *ndufs5*, *rnf19b*, and *macf1*. Given the key function of *ndufs5* in mitochondrial Complex I, we predicted that *ndufs5* was the gene most likely to be interacting with the mitochondria. However, we formally evaluated whether *rnf19b* and *macf1* had known mitochondrial interactions.

We used the STRING database for *Xiphophorus maculatus* to identify all known interactions of *rnf19b* and *macf1* from fusion, co-occurrence, homology, coexpression, experimental, database, or textmining data (including such data transferred from other organisms via homology). This represents a permissive set of genes that may be interacting for the purposes of our analysis. Next, we evaluated whether *rnf19b*, *macf1*, or any of the genes they interact with were annotated in the GO category GO:0005739 (cellular component – mitochondria), a category that includes both mitochondrial and mitonuclear genes. Of 1,860 annotated interactions for *macf1* and 748 annotated interactions for *rnf19b*, only one interaction involving *rnf19b* (with *park2*) was identified as containing a mitonuclear interaction.

Importantly, *rnf19b* does not directly interact with any mitochondrial or mitonuclear proteins. Its interaction with *park2,* which plays a role in mitophagy of damaged mitochondria [17,18], is mediated via an interaction with *ube2l3* (Fig. S53). Given its role, we considered *rnf19b* a less plausible candidate for mitonuclear interactions in hybrids than *ndufs5*. However, to exclude it as a candidate, we chose to investigate its possible interactions further. Because *rnf19b* (and *macf1*) are almost perfectly linked to *ndufs5,* addressing this question raised technical challenges. We reasoned that if interactions between *rnf19b* and its partners *ube213* and *park2* were driving the mitonuclear incompatibility, we would observe deviations in ancestry in $F_2$ hybrids surrounding the *ube213* and *park2* genes (as we do in the region containing *ndufs5/*

*macf1/rnf19b*, as well as in the region on chromosome 6 containing *ndufa13*; Fig. 1A, 1D) Instead, we observe no unexpected ancestry patterns in these regions in $F_2$ hybrids (Fig. S54).

We also looked more explicitly for evidence of interactions between these genes in $F_2$ hybrids, using comparisons between observed and expected two locus genotypes in a $\chi^2$ framework. We generated expected values based on independent assortment of the observed genotypes at each locus (following Langdon et al.[2]) and compared them to the observed two-locus genotype combinations in $F_2$ hybrids. We found no evidence for deviations from random assortment of genotypes at *rnf19b* and *park2* ($\chi^2 = 4.07$, $P = 0.39$), or *rnf19b* and *ube2l3* ($\chi^2 = 7.6$, $P = 0.11$). For comparison, we see a strong deviation from random assortment of genotypes at *ndufs5* and *ndufa13* ($\chi^2 = 38$; $P = 1.2 \times 10^{-7}$).

We also wanted to search for evidence that mitochondrial ancestry was non-randomly associated with ancestry at *park2* or *ube213* (*rnf19b*'s partners in its mitonuclear interactions) in our admixture mapping population. To address this question, we analyzed ancestry covariance in the Calnali low hybrid population, where individuals segregate for both *X. birchmanni* and *X. malinche* haplotypes. We did not find evidence for correlations in ancestry between the mitochondrial haplotypes and *park2* or *ube213*, after accounting for genome-wide ancestry (all $P > 0.3$). Thus, several lines of evidence indicate that interactions involving *rnf19b* are unlikely to be the cause of the mitonuclear hybrid incompatibility.

1.1.9. Exploration of potential confounding factors in our analyses

We wanted to exclude several factors that could potentially confound our mapping analyses. First, we previously mapped a hybrid incompatibility that impacts viability in *X. birchmanni* x *X. malinche* hybrids. This previously mapped hybrid melanoma incompatibility reduces viability of adults[5], and thus we considered an impact on our analysis of an embryonic lethal incompatibility unlikely. However, the melanoma incompatibility contains an X-linked gene (*xmrk*) so we wanted to carefully analyze any potential interactions with mitochondrial ancestry.

Specifically, we wanted to ask whether there was any difference in the frequency of the melanoma incompatibility in individuals with and without the mitonuclear incompatibility. Because individuals with *X. malinche* mitochondria and homozygous *X. birchmanni* ancestry at *ndufs5* and *ndufa13* largely do not survive to adulthood (Fig. 1; see Supplementary Information 1.3.1), we used embryos in this analysis. We identified embryos likely to develop melanoma if they were to survive to adulthood based on their genotypes[5]. Specifically, we identified embryos with homozygous or heterozygous *X. birchmanni* ancestry at *xmrk* and homozygous *X. malinche* ancestry at *cd97*[5]. We used a Fisher's Exact test to evaluate whether the melanoma risk genotype was more prevalent in individuals with *X. malinche* mitochondria and homozygous *X. birchmanni* ancestry at *ndufs5* or *X. malinche* mitochondria and homozygous *X. birchmanni* ancestry at *ndufa13.* We found no evidence for differences between groups based on this test (both $P > 0.5$).

Another potential confounding factor in analysis of hybrid incompatibilities, especially in natural hybrid populations, is issues arising from excess ancestry linkage disequilibrium (LD)[11]. Specifically, if unaccounted for, population structure and ancestry assortative mating[19] can increase background admixture LD, resulting in a non-conservative p-value distribution in admixture mapping studies with an excess of false positives[11]. One common driver of this problem in natural populations is ancestry assortative mating[19]. We chose to evaluate evidence of ancestry assortative mating in the Calnali Low hybrid populations used for admixture mapping. We note that ancestry assortative mating was not possible in our $F_2$ datasets because the $F_1$ parents did not vary in their ancestry. Using previously established methods[20,21], and paired mother-embryo sequencing data generated for this study, we asked whether the maternal ancestry and the inferred paternal ancestry for a given mating event differed from that expected by chance under random mating. Specifically, we randomly sampled one embryo per female, calculated the difference in ancestry between the mother and her offspring (reflecting the mother's ancestry, the paternal ancestry, and some variance introduced by recombination in the maternal and paternal gametes). We compared the observed data to expectations under random mating by randomly sampling pairs of individuals from the adult ancestry distribution and simulating their offspring's ancestry[20,21]. We found no significant differences between the observed and simulated data for the Calnali Low hybrid population (Fig. S55). This suggests that ancestry-assortative mating will not generate issues in our analysis.

More generally, our analyses of covariance in ancestry between mitochondrial and nuclear genotypes accounts for ancestry variation among individuals (i.e. r|a measure proposed by Schumer & Brandvain[11]) and simulations applied to determine significance thresholds in the $F_2$ and admixture mapping datasets preserved the ancestry structure of the real data, following best practices for reducing inflation of p-value distributions due to population structure[11] (Supplementary Information 1.1.5).

Finally, we wished to rule out any possible issues arising from assembly errors. Assembly errors can generate artifactual associations between regions of the genome simply because two regions are expected to be unlinked based on their position in the genome assembly but in reality are closely linked. This can cause the appearance of selection maintaining associations between physically unlinked sites. As a result of misassemblies, however, we expect unusual patterns of LD between a marker and its supposed neighbors[22]. To evaluate this, we selected the peak markers in the chromosome 6 and chromosome 13 QTL containing *ndufa13* and *ndufs5* and evaluated patterns of decay in admixture LD over genetic distance. We found that these markers showed typical patterns of LD decay with nearby sites (Fig. S55), suggesting that assembly errors could not explain the associations we observed involving these regions.

##### 1.1.10. Analysis of ancestry and potential interactions at other Complex I genes

We identify two genome-wide significant peaks in our admixture mapping population on chromosome 13 and 15 (at a 5% FPR threshold), and one peak on chromosome 6 that is nearly significant at a 10% FPR threshold. Although the peak on chromosome 6 did not reach genome-wide significance, we are confident in this signal because of patterns observed in $F_2$ hybrids in this region. Specifically, we are able to confirm the signals on both chromosome 6 and 13 with lab-reared $F_2$ hybrids where individuals with *X. malinche* mitochondria have a strong deficit of *X. birchmanni* ancestry in these regions (Fig. 1; Langdon et al.[2]).

Given the results of fine mapping on chromosome 13 implicating Complex I as the focal point of an incompatibility with the *X. malinche* mitochondria, we wanted to determine whether the chromosome 6 and 15 regions also contained genes involved in Complex I, and whether these genes were likely to drive the admixture mapping signal we observed on these two chromosomes.

Of the nuclear genes that are subunits or required assembly factors of mitochondrial Complex I (listed in Table S1), one falls within the chromosome 6 QTL region but no Complex I genes fall within the chromosome 15 QTL region. The associated chromosome 6 region is 361 kb and contains 20 genes, including *ndufa13*, whereas the chromosome 15 region is 3.3 Mb and contains 82 genes. Complex I gene *ndufaf4* falls nearby the chromosome 15 peak but outside the genome-wide significant region. The genes in these intervals are listed in Table S2 & Table S3.

Examination of local signals indicates that the peak on chromosome 6 precisely colocalizes with the location of *ndufa13* (Fig. S7). We next evaluated the probability that the chromosome 6 QTL would overlap with one of the 45 Complex I genes by chance. To do so, we randomly permutated the 361 kb QTL interval onto the genome, generating 1,000 null QTL intervals. We overlapped these intervals with annotated Complex I genes and asked how frequently we would expect overlap by chance. For the chromosome 6 region, overlap with any Complex I gene was unexpected by chance ($P = 0.015$ by simulation).

As noted above, in the main text we focus our analyses on interactions between the mitochondria and *ndufa13* and *ndufs5*. We make this choice for several reasons. First, both of these genes involve interactions between *X. birchmanni* nuclear ancestry and *X. malinche* mitochondrial ancestry, whereas the signal on chromosome 15 is driven by *X. birchmanni* mitochondrial ancestry (Fig. 1, Supplementary Information 1.2.1). Moreover, the driver of the chromosome 15 signal is unclear. No Complex I genes fall under the chromosome 15 peak. There are 45 genes in the peak annotated as interacting with genes that have a mitonuclear function based on the STRING database[23] but none are annotated as interacting directly with mitochondrial genes. Since we focus here on Complex I genes, we did not pursue these possible interactions further but note that it may be an interesting avenue for future work. The closest Complex I gene to the chromosome 15 peak is *ndufaf4,* which falls ~300 kb outside of the peak (Fig. S7), exhibits no nonsynonymous differences between species, no evidence of expression differences between species, and no evidence of allele specific expression in $F_1$ hybrids (see Supplementary Information 1.3). We note, however, that it is possible for *ndufa4* to be involved in mitonuclear incompatibility if there are synonymous substitutions that impact translational efficiency or mRNA stability, as has been reported in some cases[24,25].

We also performed a more targeted search for involvement of other Complex I genes found throughout the genome in mitonuclear incompatibility. We evaluated ancestry at all Complex I genes in natural hybrid populations to explore evidence for selection. At *ndufs5* and *ndufa13* we see striking depletion of non-mitochondrial parent ancestry in hybrid populations (Fig. S8), consistent with selection purging incompatible genotype combinations. We asked whether any of the remaining 45 annotated nuclear genes involved in Complex I (aside from *ndufs5*, *ndufa13*, *ndufaf4*) had unusual ancestry in hybrid populations, defined as in the lower 5% of the genome-wide distribution of non-mitochondrial parent ancestry (Fig. S8). We evaluated ancestry in two hybrid populations that have fixed *X. birchmanni* mitochondrial ancestry (Aguazarca and Acuapa) and two hybrid populations that have fixed or high frequency *X. malinche* mitochondrial ancestry (Tlatemaco and Chahuaco Falls). For Chahuaco Falls, we subset our data to exclude individuals with *X. birchmanni* mitochondrial haplotypes (7% of individuals).

We did not identify any additional genes that were depleted for non-mitochondrial parent ancestry in populations with the *X. malinche* mitochondria. For populations with the *X. birchmanni* mitochondria*, ndufb5* and *ndufaf2* were shared ancestry outliers, indicative of possible selection (Fig. S8). While we focus on the incompatibility involving *X. malinche* mitochondria in this work, these results hint that the incompatibility involving the *X. birchmanni* mitochondria may be even more complex than the admixture mapping results indicate (Fig. 1H). However, neither *ndufb5* or *ndufaf2* had a significant correlation with mitochondrial ancestry or ancestry at the chromosome 13 or 15 *X. birchmanni* associated peaks in the admixture mapping population, even at a relaxed p-value threshold (all $P > 0.16$).

#### 1.2 The strength, symmetry, and complexity of mitonuclear incompatibility selection

##### 1.2.1 Evidence for a complex *X. birchmanni* mitochondria - *X. malinche* nuclear gene incompatibility

Although we knew from our $F_2$ data that *X. malinche* mitochondria appear to be incompatible with homozygous *X. birchmanni* ancestry on chromosome 6 and chromosome 13, we did not know *a priori* whether there was also selection on the combination of *X. birchmanni* mitochondria and nuclear genes. Our admixture mapping approach allowed us to evaluate evidence for selection on mitonuclear interactions involving the *X. birchmanni* mitochondria as well. Although we do not focus on this incompatibility in the main text, we outline our findings about it below.

Visualizing the genotype combinations on chromosome 6, chromosome 13, and chromosome 15, we see that individuals with *X. birchmanni* mitochondria and *X. malinche* ancestry at *ndufa13* on chromosome 6 are viable, but that individuals with *X. birchmanni* mitochondria and homozygous *X. malinche* ancestry at *ndufs5* on chromosome 13 and at the chromosome 15 QTL are missing from our dataset (Fig. 1C; Fig. S9). This indicates the presence of mitonuclear incompatibilities between the *X. birchmanni* mitochondria and the peaks on chromosome 13 and 15. While these frequencies would not be expected by chance in the absence of population structure, we wanted to confirm that the observed two-locus genotype frequencies were unexpected conditioning on the ancestry structure of the Calnali Low hybrid population. To evaluate this, we performed simple simulations. We first recorded the observed mitochondrial ancestry for each individual from the Calnali Low population. Next, we used observed genome-wide ancestry for each Calnali Low hybrid to generate simulated nuclear genotypes at *ndufs5*. Specifically, we drew the diploid genotype for each individual from a random binomial weighted by the proportion of that individual's genome derived from the *X. malinche* parent species. We repeated this procedure 10,000 times.

Based on these simulation results, we find that the observed frequency of individuals with *X. birchmanni* mitochondria and homozygous *X. malinche* ancestry at *ndufs5* is unexpected by chance, even when accounting for ancestry structure in the population ($P < 10^{-4}$ by simulation). We replicated this simulation procedure for individuals with *X. malinche* mitochondria and obtained similar results (observed frequency $P < 10^{-4}$ by simulation). We repeated this analysis for the peak associated marker on chromosome 15 and found that the observed frequency of individuals with *X. birchmanni* mitochondria and homozygous *X. malinche* ancestry at chromosome 15 is also much lower than expected by chance (observed frequency $P < 10^{-4}$ by simulation), whereas genotype combinations do not deviate from expectations by chance in individuals with *X. malinche* mitochondria ($P = 0.79$ by simulation). The results of the simulations described in this section are summarized in Fig. S9.

As discussed in Supplementary Information 1.1.10, although there is strong evidence for a mitonuclear incompatibility between chromosome 15 and the *X. birchmanni* mitochondrial haplotype, it is unclear which gene in the large QTL region is involved in the incompatibility (Table S3). While our results indicate that *X. malinche ndufs5* is incompatible with the *X. birchmanni* mitochondria (Fig. S9), no Complex I genes are found in the chromosome 15 interval, making the mechanism of this incompatibility and the precise genes involved an exciting question for future work.

##### 1.2.2 Estimating the strength of selection against incompatible genotype combinations using lab-bred $F_2$ and natural hybrids

We were interested in inferring the strength of selection against two mitonuclear combinations: homozygous *X. birchmanni* ancestry at *ndufs5* with *X. malinche* mitochondria and homozygous *X. birchmanni* ancestry at *ndufa13* with the *X. malinche* mitochondria (see below). We were also interested in inferred the strength of selection on interactions between the *X. birchmanni* mitochondria and *X. malinche* ancestry at *ndufs5*, which is also under selection (Fig. 1C, Fig. S9), for comparison.

We first focused on the combination of homozygous *X. birchmanni* ancestry at *ndufs5* with *X. malinche* mitochondria, and took advantage of observed genotype frequencies in our $F_2$ mapping population to infer the strength of selection, using an approximate Bayesian computation approach. Because 100% of $F_2$ hybrids had *X. malinche* mitochondria, we could focus our simulations solely on the nuclear interactors. We calculated the expected frequency of each nuclear genotype at *ndufs5* from Mendelian expectations (25% MM, 50% MB, 25% BB) and randomly drew a selection coefficient against homozygous *X. birchmanni* ancestry (BB), $s$, and dominance coefficient, $h$, from a uniform prior distribution ranging from 0-1. We used the selection and dominance coefficients to modify the expected frequency of each of the three genotypes post-selection. Based on these expected frequencies we simulated sampling of 937 individuals (the number of $F_2$ individuals successfully genotyped at *ndufs5*) and counted the number of observed MB and BB genotypes in the simulation. We repeated this procedure for a total of 500,000 simulations and performed rejection sampling at a 5% threshold, based on the observed number of MB and BB genotypes in the real data from $F_2$s and the average *X. malinche* ancestry observed at *ndufs5* in $F_2$s.

We repeated this procedure for estimating the strength of selection on the combination of *X. birchmanni* ancestry at *ndufa13* with *X. malinche* mitochondria. The interactions we infer between *ndufa13* and *ndufs5* are relatively weak compared to the strength of selection on each in isolation (see *Supplementary Information 1.2.5 Detecting a complex interaction between* ndufs5, ndufa13*, and the mtDNA*), justifying our choice to model the two interactions with the *X. malinche* mitochondria independently. Simulations followed the same procedures outlined above, except that rejection sampling was based on the 932 individuals successfully genotyped at *ndufa13*. Results for these simulations are reported in the main text (Fig. 1) and in Fig. S3.

Since all $F_2$ individuals have *X. malinche* mitochondrial haplotypes, we used a different approach to evaluate the strength of selection on the incompatibility between *ndufs5* and the *X. birchmanni* mitochondria. To evaluate this question, we took advantage of the population data from Calnali Low, where we observe a few individuals with *X. birchmanni* mitochondria and homozygous *X. malinche* ancestry at *ndufs5*. We focus only on the interaction with *ndufs5* because we do not see evidence for an interaction between the *X. birchmanni* mitochondria and *ndufa13* (see *Supplementary Information 1.2.1 Evidence for a complex* X. birchmanni *mitochondria -* X. malinche *nuclear gene incompatibility*).

We again used an approximate Bayesian computation approach, this time with the forward-time population genetic simulator SELAM[26], to estimate the strength of selection against the mitonuclear incompatibility that is consistent with observed patterns in our data.

SELAM takes user-provided information on demographic history and selection to simulate admixture between populations and record ancestry for each individual along the genome under a Wright-Fisher model with selection[26]. We used a newly added selection parameter ‘N’ to simulate selection on the mitochondria and an interacting nuclear locus. Because selection on the combination of the *X. malinche* mitochondria and homozygous *X. birchmanni* ancestry at *ndufs5* will necessarily impact allele frequencies of the *X. birchmanni* mitochondria and *X. malinche* ancestry at *ndufs5*, we jointly inferred selection coefficients on the mitonuclear incompatibility in both directions. For each of 500,000 simulations, we sampled from a uniform prior distribution for a range of demographic parameters, including the number of generations since initial admixture (20-80), hybrid population size (200-2,000), and initial admixture proportion (sampled from a normal distribution centered at 0.35 with standard deviation 0.02). These values were guided by previous estimates for populations on the Río Calnali[5,20]. We also sampled from a uniform prior for parameters relevant to the hybrid incompatibility; for the two mitochondrial haplotypes we independently varied the selection coefficient when combined with an opposite ancestry nuclear locus from 0-1, and varied dominance coefficients for heterozygotes at the nuclear locus (0-1).

Simulations were accepted if the frequency of the homozygous *X. malinche* nuclear genotype-*X. birchmanni* mitotype in the simulated population was within 5% of the frequency observed in the Calnali Low data (1% frequency) and if the frequency of the homozygous *X. birchmanni* nuclear genotype-*X. malinche* mitotype was less than 0.5% frequency. Out of 500,000 simulations, approximately 500 were accepted and used to estimate posterior distributions.

Posterior distributions mirrored the prior distributions for the number of generations since initial admixture, admixture proportion, dominance coefficients, and hybrid population size. We recovered well-resolved posterior distributions for selection coefficients in both directions of the incompatibility. For the incompatibility involving the *X. birchmanni* mitochondria, which we were not able to estimate with $F_2$ data, we inferred appreciable selection on individuals with mismatched mitonuclear ancestry (maximum a posteriori or MAP estimate = 0.17, 95% confidence intervals 0.08-0.55; Fig. S3). This indicates that both combinations of mitochondrial ancestry and nuclear ancestry at *ndufs5* are under selection in hybrids.

We had already inferred selection on the incompatibility involving the *X. malinche* mitochondria using data from $F_2$ hybrids, where we infer selection to be near complete ($s \sim 1$). Here, we compare those results to the results we obtain using the Calnali Low hybrid population data to estimate selection against the same incompatible genotype combination. The posterior distribution for selection coefficient based on this inference also resulted in estimates of extremely strong selection (MAP estimate = 0.7, 95% confidence intervals 0.13-0.97; Fig. S3), albeit weaker than that inferred from the $F_2$ hybrid data. We believe that this result is expected when inference is based on data from natural hybrid populations. Specifically, in both the $F_2$ data and the Calnali Low hybrid population data, we found no individuals with *X. malinche* mitochondrial – homozygous *X. birchmanni ndufs5* ancestry. However, given a larger number of generations since initial selection in simulations of natural hybrid populations, a broader range of selection coefficients are consistent with the data.

##### 1.2.3 Power to detect asymmetric versus symmetric mitonuclear incompatibilities in genome-wide scans

We find that some of the detected interactions between mitochondrial haplotypes are "symmetric," meaning that both mitochondrial types cause incompatibilities with the nuclear genotype of the opposite species, and some are "asymmetric," meaning that only one of the mitochondrial types triggers an incompatibility (Fig. 1C; Fig. 1F; Fig. S41). We discuss the evolutionary implications of this finding in more detail in Supplementary Information 1.2.4.

An important technical consequence of these genetic patterns is that there is variation in our power to detect these two different types of signals in genome-wide admixture mapping scans. In particular, at similar strengths of selection, we expect to have less power to detect asymmetric incompatibilities compared to symmetric incompatibilities at our genome-wide FPR threshold. We demonstrate this for the case of interactions between mitochondrial haplotypes, *ndufs5*, and *ndufa13* using simulations and the inferred selection coefficients for each interaction (Fig. 1).

Focusing first on *ndufs5*, we model selection on incompatibilities with both the *X. malinche* and *X. birchmanni* mitochondrial haplotypes, guided by the simulation results presented in Supplementary Information 1.2.2. Using the observed genome-wide ancestry distributions of the individuals in the admixture mapping population, we performed the following steps to simulate individual genotypes:

1) We randomly drew a genome-wide ancestry for that individual.
2) We simulated mitochondrial ancestry by drawing from a random binomial in R, weighted by the proportion of the genome derived from *X. malinche*. Specifically, if the draw from a random binomial returned zero, we assigned that individual an *X. birchmanni* mitochondrial haplotype for the simulation. If the draw returned one, we assigned that individual a *X. malinche* mitochondrial haplotype for the simulation.
3) We repeated this procedure to simulate diploid nuclear alleles at *ndufs5*.
4) For individuals with compatible simulated genotype combinations, we retained that individual for the final simulated dataset. This included all individuals that were heterozygous at the simulated nuclear locus or had matched homozygous nuclear and mitochondrial haplotype ancestry.
5) For individuals that had homozygous *X. birchmanni* ancestry at the simulated nuclear locus and an *X. malinche* mitochondrial haplotype, we drew from a random binomial to determine whether they survived, with the probability of survival set to 1 – 0.998 (the MAP estimate for the incompatible interaction between *ndufs5* and the *X. malinche* mitochondria; Fig. 1B). If individuals survived, they were retained in the final simulated dataset, otherwise they were excluded.
6) For individuals that had homozygous *X. malinche* ancestry at the simulated nuclear locus and an *X. birchmanni* mitochondrial haplotype, we again drew from a random binomial to determine whether they survived, with the probability of survival set to 1 – 0.17 (the MAP estimate for the incompatible interaction between *ndufs5* and the *X. birchmanni* mitochondria; Fig. S3). If individuals survived, they were retained in the final simulated dataset, otherwise they were excluded.
7) We continued the simulation until 359 individuals had been added to the simulated dataset, and evaluated the association between the simulated nuclear and mitochondrial ancestry as we had for the real data.

8) We repeated this procedure 1,000 times.

Examining the p-value distribution for the simulated incompatibility with *ndufs5* (Fig. S57), we see that we expect to have excellent power to detect this incompatibility in our admixture mapping data. 96% of simulations exceed our genome-wide significant threshold.

When focusing on the incompatibility with *ndufa13*, these simulations followed the procedures outlined above except that no selection was implemented on individuals with *X. malinche* ancestry at *ndufa13* and the *X. birchmanni* mitochondria, and the selection coefficient used for the combination under selection was set to 0.91, as inferred from ABC simulations (see Fig. 1E; Supplementary Materials 1.2.2).

Examining the p-value distribution for the simulated incompatibility with *ndufa13* (Fig. S57), we see that our power to detect this interaction is substantially weaker. We detect the incompatibility at our genome-wide significance threshold in only 44% of simulations. The median $-\text{Log}_{10}$(p-value) of the association across 1,000 simulations is $2.2 \times 10^{-5}$, close to the observed chromosome 6 peak of $3 \times 10^{-5}$. Notably, the parameters for the selection coefficient used in this simulation were estimated from $F_2$ hybrids (see Supplementary Materials 1.2.2), and thus are independent of the admixture mapping results. This underscores that we have weaker power to detect the observed association with *ndufa13* in a genome-wide scan due to the architecture of the incompatibility (i.e. that the incompatibility involves only the *X. malinche* and not the *X. birchmanni* mitochondria).

We repeated the procedure described above to ask about our power to detect nuclear-nuclear incompatibilities under strong selection ($s$=1) except that instead of simulating mitochondrial ancestry in step 2, we simulated a diploid interacting allele. We performed simulations under scenarios where the nuclear-nuclear incompatibility was dominant and where it was recessive. The results of these simulations indicated that we had good power to detect selection on a dominant and recessive symmetric (detected in 100% and 89% of simulations respectively) or dominant asymmetric nuclear-nuclear hybrid incompatibility (detected in 85% of simulations) at the genome-wide significance threshold. However, we had poor power to detect recessive asymmetric nuclear-nuclear hybrid incompatibilities (detected in 3% of simulations).

This finding suggests that although we do not detect additional nuclear interactors of the loci identified in our admixture mapping scan, we cannot rule out the presence of a more complex architecture due to weak power to detect recessive nuclear-nuclear interactions. However, we have some evidence that the Complex I hybrid incompatibility involving the *X. malinche* mitochondria does not involve more than three partners. This is because scans for ancestry depletion have excellent power even at weak selection coefficients[2] and we investigated ancestry depletion for all Complex I genes in natural hybrid populations (Supplementary Information 1.1.10). We found no evidence of depletion of *X. birchmanni* ancestry at other Complex I genes (apart from *ndufs5* and *ndufa13*; Fig. S8), indicating that this subset of genes is unlikely to be under selection in hybrid populations. By contrast, we do see some evidence of selection on *X. malinche* ancestry at several Complex I genes in hybrid populations with *X. birchmanni* mitochondria (see Supplementary Information 1.1.10).

##### 1.2.4 Asymmetry in selection on mitonuclear incompatibilities and variation in incompatibility architecture

We find evidence for selection when either the *X. malinche* or *X. birchmanni* mitochondrial haplotype is combined with homozygous nuclear ancestry at *ndufs5* of the other species (Fig. 1C). This bi-directional selection is not predicted by classic theory for how hybrid incompatibilities evolve (Fig. S56). Classic evolutionary theory predicts that selection will only occur on one of the two possible combinations of heterospecific genotypes in a hybrid incompatibility (e.g. as observed for the chromosome 6 and chromosome 15 associations; Fig. 1, Fig. S9). This is because the second combination will recapitulate the ancestral genotype (Fig. S56) and hybrids with the ancestral genotype should not suffer fitness consequences[27].

How might a hybrid incompatibility like the one we report here for *ndufs5*, with selection on both heterospecific genotype combinations, arise? One possibility is that multiple substitutions impacting gene interactions have occurred in one or both lineages. In such a case, the ancestral genotype combination would no longer exist, making selection on both heterospecific genotype combinations possible. Indeed, this may be a likely explanation given the many substitutions that have accumulated between species (Fig. 4). This result highlights the importance of empirical work characterizing the architecture of hybrid incompatibilities, particularly given the prediction that there will be distinct evolutionary outcomes after hybridization as a result of asymmetric versus symmetric hybrid incompatibilities[28].

While we do not identify the classic asymmetry predicted by the DMI model, we do see asymmetry in the strength of selection on the two incompatible mitochondrial – *ndufs5* genotype combinations, as well as variation in the architecture of the genetic incompatibility. Our data suggests that the combination of *X. birchmanni* mitochondria with homozygous *X. malinche* ancestry at *ndufs5* is much less deleterious than *X. malinche* mitochondria combined with homozygous *X. birchmanni* ancestry at *ndufs5*. While the former combination is under strong selection (Fig. S3), the latter appears to be lethal (Fig. 1B).

Another notable difference in the two *ndufs5*-mitochondrial incompatibilities is their inferred genetic architecture. While individuals with *X. birchmanni* mitochondria appear to have reduced tolerance of *X. malinche* ancestry at *ndufs5* and the chromosome 15 peak, they are tolerant of *X. malinche* ancestry at *ndufa13*. By contrast, individuals with *X. malinche* mitochondria are intolerant of homozygous *X. birchmanni* ancestry at both *ndufs5* and *ndufa13* (Fig. 1H), but are tolerant of *X. birchmanni* ancestry at the chromosome 15 peak (Fig. S9).

We speculate that the observation of bi-directional selection and variation in the architecture of the mitonuclear incompatibility may highlight the importance of mitonuclear interactions as a "hotspot" for the evolution of hybrid incompatibilities between *X. birchmanni* and *X. malinche*. Indeed, mitonuclear hybrid incompatibilities are extremely common[29,30], suggesting that genes in these pathways may play an outsized role in the evolution of reproductive isolation between species. Researchers have hypothesized that the prevalence of mitonuclear incompatibilities may be due to higher mitochondrial substitution rates and tight co-evolution between mitochondrial genes and their interacting nuclear partners[31]. These mechanisms may also explain the complexity and bi-directionality of the Complex I incompatibility identified in *X. birchmanni* × *X. malinche* hybrids.

##### 1.2.5 Detecting a complex interaction between *ndufs5, ndufa13*, and the mtDNA

Above and in the main text, we characterize incompatible interactions between three unlinked regions: the *X. malinche* mitochondria, and the nuclear genes *ndufs5* and *ndufa13*. Classic theoretical work in population genetics envisioned that three locus incompatibilities would include interactions between all pairs of genes involved[32], although this is not necessarily a prediction of theoretical work on multi-locus incompatibilities from systems biology[33]. Though the high lethality of the *ndufs5* and *ndufa13* interactions with the mitochondria makes assessing *ndufs5-ndufa13* interactions difficult, we used rare survivors of the *ndufa13* incompatibility to test for evidence of three-gene interactions in our dataset.

For other experiments (see *Supplementary Information 1.3.3: Tracking the timing of ndufa13 lethality*), we generated an additional 74 $F_2$ individuals that we tracked from birth to five months of age. We noticed that 7 of these individuals were incompatible at *ndufa13* (i.e. they had homozygous *X. birchmanni* ancestry at *ndufa13* and an *X. malinche* mitochondria) but survived throughout the course of the experiment. Of these 7 surviving individuals, 5 had homozygous *X. malinche* ancestry at *ndufs5*. In other words, individuals that harbored any *X. birchmanni* ancestry at *ndufs5* were underrepresented in juvenile survivors of the *ndufa13* incompatibility (association test $\chi^2 = 5.9$, p=0.051). This depletion of heterozygotes was surprising given that we previously inferred the *ndufs5* incompatibility to be almost completely recessive (see *Supplementary Information 1.2.2*), and hints at a possible interaction between these genes in *ndufa13* incompatible individuals. As such, we sought to test for a non-additive effect of selection against the *ndufs5* and *ndufa13* incompatibilities with greater statistical power.

To do so, we assessed the relationship between *ndufs5* and *ndufa13* genotype frequencies in the full set of $F_2$ hybrids used in the study, all of whom harbored *X. malinche* mitochondria. We began with a $\chi^2$ test of independence of two-locus genotypes at *ndufs5* and *ndufa13* across all 1001 successfully genotyped individuals (excluding the one individual with *X. birchmanni* ancestry at *ndufs5*). This test showed a significant association between ancestry at *ndufs5* and *ndufa13* in surviving hybrids ($\chi^2 = 6.9$, *d.f.* = 2, $P = 0.032$), consistent with selection acting on these loci in combination. This dataset included 33 survivors of the *ndufa13* incompatibility (i.e. individuals with homozygous *X. birchmanni* ancestry at *ndufa13*, ~3% of the total individuals). Examining the observed genotype combinations in the surviving individuals, we found that the majority of survivors of the *ndufa13* incompatibility were homozygous *X. malinche* at *ndufs5*. This suggests that even one *X. birchmanni* allele at *ndufs5* may exacerbate the harmful effects of the *ndufa13* incompatibility.

To directly test the idea that individuals with the *ndufa13* incompatibility were more likely to survive if they were homozygous *X. malinche* at *ndufs5*, we first used a permutation approach. For each *ndufa13* incompatible individual, we drew a random genotype without replacement from the observed genotypes at *ndufs5* across all $F_2$ hybrids. Resampling observed genotypes accounts for selection changing the genotype frequencies at each locus in isolation, and allows us to simply ask whether the inheritance of the two loci in combination matches expectations. We repeated this procedure 10,000 times and compared the simulated two-locus genotype frequencies to the observed data. We found that the number of surviving individuals that are *ndufa13* incompatible and homozygous *X. malinche* at *ndufs5* in our real data greatly exceeds what is expected by chance ($P = 0.015$ by simulation).

As a complementary approach, we used population genetic models to ask whether individuals with incompatible genotypes at *ndufa13* had different genotypes at *ndufs5* than expected if selection were acting independently on them. To do so, we used the posterior distributions we inferred assuming independent selection on *ndufs5* and *ndufa13* (i.e. drawing $s$ and $h$ for each genotype from the posterior distributions generated in *Supplementary Information 1.2.2)* and modeled selection on each genotype with our observed sample size. We recorded the frequencies of *ndufs5* homozygotes and heterozygotes among simulated *ndufa13*-incompatible individuals, and compared them to the frequencies observed in the real data. We found that the two-locus genotype frequencies observed in the real data significantly deviated from a model of independent selection on the two loci ($P = 0.018$ by simulation). From our simulations, we expect ~65% of surviving *ndufa13* incompatible individuals to harbor one *X. birchmanni* allele at *ndufs5*, but only observe this genotype in 45% of individuals in our dataset. This complementary analysis using population genetic models again indicates that independent, additive selection on *ndufs5* and *ndufa13* incompatibilities is unlikely to generate the patterns we observe in our data.

These results indicate that while incompatible genotypes at *ndufa13* and *ndufs5* are nearly lethal or completely lethal on their own, we are better able to predict the survival of *ndufa13* incompatible individuals as a function of their genotype at *ndufs5*. Fully matched ancestry at *ndufs5* is significantly over-represented in survivors of the *ndufa13* incompatibility, indicating an interaction between these incompatibilities at the level of hybrid fitness. Importantly, this complexity is not well-described by current verbal or theoretical models of pairwise or multi-gene hybrid incompatibilities, and points to the value of empirical examples in understanding the true landscape of hybrid fitness.

The mechanisms through which genotype at *ndufs5* impacts fitness in surviving *ndufa13* incompatible individuals is an exciting area for future work. In our data collected from embryos (see *Supplementary Information 1.3.4 & 1.3.5)*, there are some hints of an interaction between these genotypes on morphological and physiological phenotypes (on head width – Table S7, on respiration and rotenone response – Table S10, S11). However, since just ~3% of $F_2$ individuals with the incompatible genotypes at *ndufa13* survive, addressing this question experimentally will be challenging and may require thousands more hybrids to generate an appropriate sample size.

We note that despite this evidence of interaction, elsewhere in the manuscript we model selection on *ndufs5* and *ndufa13* separately for simplicity (see *Supplementary Information 1.2.2. Estimating the strength of selection against incompatible genotype combinations using lab-bred* $F_2$ *and natural hybrids*). We expect that this approach is a reasonable simplification for modeling purposes since *ndufa13* incompatible homozygotes are rare in our dataset of adult individuals (~3% of individuals). In practice, weak interactions between *ndufs5* and *ndufa13* will have a minor effect from a fitness perspective since incompatibility with the *X. malinche* mitochondria at either gene is largely lethal. However, distinguishing between these possibilities could have important conceptual implications in evolutionary genetics.

#### 1.3 Organismal and cellular signatures of mitonuclear incompatibility

##### 1.3.1 Embryo staging and genotyping

The absence of adult individuals with *X. malinche* mitochondria and homozygous *X. birchmanni* ancestry at *ndufs5* (and to a lesser extent *ndufa13*) begs the question of whether these genotypes might be observed at earlier life stages. This question is of particular interest because in zebrafish, pharmacological disruption of mitochondrial Complex I leads to arrested embryonic development [34,35]. In order to evaluate this, we classified the developmental stage of embryos collected from the Calnali Low hybrid population in March 2020 prior to genotyping. Ovaries were removed from 41 ethanol-preserved gravid Calnali Low females and processed in order of increasing distance from the urogenital opening. Each egg was separated from the ovarian tissue and assigned to a developmental category from 1-11 based on criteria outlined previously[13]. Unfertilized eggs (stage <4) were not included in further analyses. Embryos which showed characteristics of two consecutive developmental stages were assigned the average of the two stages, and those that showed abnormal characteristics (e.g. embryos with significant pigmentation yet relatively short body length) were noted as abnormal and assigned the stage most consistent with their characteristics.

In broods analyzed from the Calnali Low hybrid population, we observed wide variation in developmental stage. Within-brood developmental stage ranges were greater than one in most of the fertilized broods analyzed (22 out of 38); the maximum within-brood difference was 6 stages. By contrast, variation in developmental stage was rare in broods obtained from pure parental species. We dissected and staged the eggs of 29 *X. birchmanni* females and 36 *X. malinche* females, yielding 8 and 11 fertilized broods, respectively. No broods in either parental species showed differences greater than 1 stage between embryos. These results are unexpected if the frequency of variable-stage broods (i.e. broods with a developmental range greater than 1) were identical between hybrids and parentals ($P = 0.004$ for hybrids vs. *X. birchmanni*, $P < 0.001$ for hybrids vs. *X. malinche*, Fisher's Exact Test). This suggests that large variation in developmental stage is a unique feature of hybrid broods.

Following staging, we performed low-coverage sequencing for embryos from each hybrid mother following the procedure described in *Supplementary Information 1.1.3 Low coverage library preparation and sequencing*. Specifically, for each fertilized brood, we sequenced two individuals of each developmental stage observed in the brood. In broods where incompatible mitochondrial and nuclear genotypes were identified, we performed a second round of sequencing to sample all embryos in the brood. Mitochondrial ancestry and local ancestry in the nuclear genome were determined for a total of 269 embryos using the pipeline described in *Supplementary Information 1.1.4 Local ancestry inference and mitochondrial haplotype inference*.

To determine the stage at which the *ndufs5* mitonuclear incompatibility acts, we categorized developmental stage as a function of mitochondrial and *ndufs5* ancestry (Fig. 2A). To control for variation in the average developmental stage of compatible and incompatible broods, we also calculated the difference between the stage of an embryo and the maximum stage of all the embryos in the same brood (i.e. its relative stage) as a function of nuclear and mitochondrial genotype (Fig. 2B). Among embryos carrying *X. malinche* mitochondria, we observed a significant difference in the relative stage of embryos with and without homozygous *X. birchmanni* ancestry at *ndufs5* ($P < 10^{-6}$, Welch's two-sample *t*-test), with incompatible individuals lagging behind their brood's maximum stage by an average of 2.3 stages.

We repeated this analysis focusing on *ndufa13* ancestry. In contrast to our findings with *ndufs5*, we observed no significant difference in relative stage between *X. malinche* mitotype embryos with and without homozygous *X. birchmanni* ancestry at *ndufa13* (Fig. S11, $P = 0.10$, Welch's two-sample *t*-test). This suggests that the incompatible interaction between the *X. malinche* mitochondria and *X. birchmanni* ancestry at *ndufa13* occurs later in development.

Analyses of adults from the Calnali Low hybrid population indicate that the combination of *X. malinche* ancestry at *ndufs5* and the *X. birchmanni* mitochondria is also depleted relative to expectations based on ancestry at the two loci (*Supplementary Information 1.2.1*). This genotype combination is inferred to be under moderate selection (Fig. S3; Fig. S9). However, we sampled only two embryos with *X. birchmanni* mitochondria and homozygous *X. malinche* ancestry at *ndufs5*, making it difficult to evaluate whether this incompatibility impacts developmental stage (Fig. S10).

Past research has shown that poecilid embryos that do not complete all stages of embryonic development in the maternal environment (poecilids are livebearing fish) cannot survive except in special media conditions[36,37]. We used several approaches to evaluate this in

*Xiphophorus*, the first of which used *X. birchmanni* developing embryos. Embryos were transferred from the mother into petri dishes to allow for *in vitro* development. One brood containing 6 embryos and another containing 14 embryos were removed from the mother and placed in petri dishes in a media of LB-15 and 15% bovine serum. The embryos were all late stage embryos, ranging from stage 8-9. Despite their late stage, none of the embryos survived more than 72 hours *in vitro.* Previous attempts to raise embryos from these stages in E3 medium also failed.

We also report data here for $F_1$ hybrid embryos that were naturally born prematurely. Prematurely born fry are observed with some frequency in $F_1$ and $F_2$ hybrid broods, but occur very rarely in pure parental species (*Xiphophorus* Genetic Stock Center, personal communication). We collected data from prematurely born $F_1$ hybrids broods, which do not suffer from mitonuclear hybrid incompatibilities since our data indicates that these incompatibilities are largely recessive (Fig. S3). We recorded developmental stage and survival over time of four broods of $F_1$ hybrids that were born prematurely but were alive at birth (N=25, Table S4). Although all of these individuals exceeded developmental stage 8, 96% of them died within a week of birth (84% within two days of birth). These results indicate that like other poecilids, *Xiphophorus* embryos face high mortality unless they have completed embryonic development in the maternal environment.

Importantly, these results imply that developmental delay in embryos with mismatched *X. malinche* mitochondrial and *X. birchmanni ndufs5* ancestry (Fig. 2) may itself be sufficient to cause lethality. Since embryos with this genotype combination do not appear to progress past stage 8 of embryonic development, despite normal development in their unaffected siblings (Fig. 2), these embryos will be born underdeveloped and will likely suffer high levels of mortality as described above.

###### 1.3.2 Pharmacological Inhibition of Complex I in newborn fry

As an independent line of evidence for the importance of Complex I function in *Xiphophorus* embryo viability, we tested the effects of pharmacological inhibition of Complex I with rotenone on the survival of newborn fry. Upon birth, newborn (≤ 24 hours post-partum, ~3 weeks post-fertilization) fry were evenly divided into two 2 L treatment tanks with a bubbler for aeration and submerged terracotta pots for cover. In each trial, one tank contained 2 L water to which 200 uL of 1 mM rotenone in DMSO was added, for a final concentration of 100 nM rotenone. As a vehicle control, the other was filled with water to which 200 uL DMSO had been added. Every 24 hours until the end of the trial at 48 hours, the tank water was fully replaced to maintain the concentration of rotenone, and any dead fry were removed and recorded. All tanks of fry were fed *ad libitum* twice daily with newly hatched brine shrimp during the trial. We treated 2 broods of *X. birchmanni* (30 fry total), for total samples sizes of 15 fry each. We found dramatic differences in survival between the rotenone and control treatments based on a two sample proportions test ($P$-value < $2 \times 10^{-6}$), with 93% mortality and 100% mortality in rotenone treatment conditions over 24 and 48 hours respectively, and 0% mortality in the control treatment.

We repeated this experiment at 100 nM with a pilot sample of *X. malinche* fry (data not shown) and based on these results predicted that *X. malinche* fry may be more sensitive to Complex I inhibition. We thus repeated the experiment at a lower concentration of 75 nM rotenone with both *X. birchmanni* and *X. malinche* newborn fry, collected as described above.

While *X. birchmanni* fry (N=9) suffered 33% mortality after 48 hours of rotenone treatment at 75 nM concentration, *X. malinche* fry (N=11) experienced 100% mortality after 48 hours of rotenone treatment at the same concentration. Based on a Fisher's exact test we see a species level difference in response to 75 nM of rotenone (P = 0.002). This observation raises the intriguing possibility that individuals with *X. malinche* mitochondria may already be more sensitive to Complex I dysfunction than individuals with *X. birchmanni* mitochondria. Investigating whether this explains differences in the degree of lethality in incompatible hybrids with the *X. malinche* versus *X. birchmanni* mitochondria will be an exciting direction for future work.

##### 1.3.3 Tracking the timing of *ndufa13* lethality

We find that individuals with ancestry mismatch at *ndufa13* have no evidence of severe development delay. However, our genetic data suggests that lethality occurs sometime shortly after birth in these individuals. To narrow in on this window, we collected broods of $F_2$s within a few hours of birth by checking tanks with $F_1$ breeding stocks once per day. Note that fry that are stillborn or born unable to swim are typically eaten by adults immediately after birth; adult predation of fry is common if fry do not hide in refugia.

Fry were transferred to a new tank with the number of fry recorded on the tank. These fry tanks were checked and counted twice daily for two weeks. For individuals that suffered mortality during this time period, time of death and cause of death was recorded and individuals were preserved for sequencing. All surviving individuals were sequenced after they were at least 3 months old to determine their genotype. One $F_2$ individual that appeared to be having uncontrolled spasms was euthanized on day 3 of life since mortality was expected to be imminent.

A total of 74 individuals were sequenced and their genotypes inferred at *ndufs5* and *ndufa13* as described previously. While we expect 25% of individuals to be homozygous for *X. birchmanni* ancestry at *ndufa13* given no excess mortality, we found that 9.5% of surviving individuals were homozygous *X. birchmanni* at *ndufa13* at the juvenile stage ($\chi^2$ test $P = 0.005$). By adulthood (in a separate sample), only 3% of individuals were homozygous for *ndufa13*. Moreover, the individual euthanized on day 3 described above was found to be homozygous for *X. birchmanni* ancestry at *ndufa13*.

Having identified five *ndufa13* incompatible individuals that escaped early-life lethality in our sample of tracked $F_2$ hybrids, we took the opportunity to test whether cardiovascular phenotypes that we observed in embryos (see Supplementary Information 1.3.5) were recapitulated in adults. The five *ndufa13* incompatible individuals and a control set of five *ndufa13* compatible (homozygous *X. malinche*) controls were sacrificed, slit along dorsal and ventral midline to allow formalin to penetrate the body cavity, and preserved in 10% formalin for at least 48 hours. Whole samples were dehydrated in 70% ethanol overnight before submission to Histo-Tec Laboratory, Inc. (Hayward, CA) for paraffin embedding, sectioning, and H&E staining. Starting from the left side, samples were leveled with a ~1 mm cut, then step sectioned on a sagittal plane at 125 μm intervals for a total of 12-13 slides per sample. For the atrium and ventricle independently, the slide in which the chamber showed the largest cross-sectional area was identified and used for further analysis in QuPath[38]. The perimeter of the chamber was manually outlined to measure total chamber area, after which a QuPath macro was used to randomly select three 22943 $\mu m^2$ (300 x 300 pixel) quadrats within the chamber's perimeter.

Within each quadrat, the percentage of area unoccupied by cells was measured by creating a thresholder on the blue channel with full resolution, Gaussian prefilter, a smoothing sigma of 2, and an intensity threshold of 200-215 depending on the image. The area of cardiomyocytes was manually annotated, and the area of blood cells was taken to be the remaining area within the quadrat after excluding cardiomyocytes and unoccupied space. The mean of each occupancy category was then taken across the three quadrats per individual for computing group-level statistics. After discarding samples which a blinded observer identified as uninterpretable due to damage during histological preparation (3 atria and 1 ventricle), we graphically compared the total area of each chamber, and the mean proportion of the chamber occupied by blood cells and cardiomyocytes (Fig. S23). Given that we found evidence for enlarged sinu-atria and reduced heartrates in embryos, we were intrigued to observe that *ndufa13* incompatible juveniles had dilated atria and changes in trabecular myocardium organization (Fig. S23). Collectively, slight dilation and abnormal trabeculation are early signs of heart failure. However, the differences between genotypes were only marginally significant by Welch's *t*-test (total atrium area $t = -3.14$, adjusted $df = 2.93$, $P = 0.053$; proportional area of cardiomyocytes in atrium $t = 1.93$, adjusted $df = 3.04$, $P = 0.148$).

###### 1.3.4 Respiration in whole embryos of different genotypes

While a lack of sufficient embryonic tissue prevents us from evaluating Complex I activity in individuals homozygous for incompatible genotypes at *ndufs5* and *ndufa13*, we can measure overall respiration rates in embryos of different genotypes. To do so, we used a Loligo® Systems respirometer to measure oxygen concentrations in a sealed 24-well plate with 1.7 mL wells. Following best practices for sustaining poecilid embryos *in vitro*[36,37], we performed respirometry measurements in L-15 media + 20 uM glutamate supplemented with 15% fetal bovine serum, hereafter referred to as respiration media. At the beginning of each day, respiration media was brought to 24°C in a loosely capped container while shaking at 100 rpm to ensure oxygen saturation. All measurements were carried out with the plate sealed with adhesive PCR plate tape, a silicon pad, and two metal compression blocks, on an orbital shaker set to 100 rpm.

After instrument setup, an initial calibration was performed by filling the plate with respiration media, capping, and waiting for stabilization of the oxygen saturation reading before setting this value as 100% saturation. The media was then replaced with respiration media plus 10 mg/mL sodium sulfite, the plate was capped, and the oxygen reading was allowed to stabilize before setting the 0% saturation point. A new 100% calibration was performed at the start of each respirometry trial, prior to the addition of embryos to the wells.

We crossed unmated $F_1$ females with $F_1$ males, then euthanized the pregnant females at a range of timepoints to access live embryos at varying stages of development. One brood was measured per day in one or two 24-well plates, with oxygen concentration in any empty wells used for blank corrections downstream. After the embryos were removed from the mother, ovarian tissue was removed and the embryos were placed in the respirometry plate. Oxygen saturation in each well was recorded every 15 seconds for at least 70 minutes. The embryos were then removed from the plate, and the media was replaced with fresh respiration media dosed with 5 μM rotenone. Once the oxygen reading of the empty plate stabilized, the embryos were returned to the plate for another 70 minutes of measurement under Complex I inhibition. After measurement with and without rotenone, the embryos were returned to rotenone-free media for phenotyping. Each embryo was video recorded for 30 seconds, then dechorionated and photographed from a dorsal and lateral perspective using a dissecting microscope equipped with an Amscope MU1803 camera. The embryos were then preserved in 95% ethanol prior to whole-genome sequencing and ancestry inference as described in Supplementary Information 1.1.3-1.1.4.

To estimate the oxygen consumption rate by each embryo with and without rotenone treatment, we discarded the first 10 minutes of measurement to allow stabilization, then used the Loligo MicroResp™ software to calculate the oxygen consumption rate in pmol/sec over the next 60 minutes of measurements. Oxygen consumption rates were corrected for changes in oxygen saturation unrelated to respiration by subtracting the batch-specific average rate of change observed in empty wells (n = 1-12, depending on brood size). The rotenone-sensitive proportion of oxygen consumption was calculated as the difference in respiration rate with and without rotenone divided by the rate without rotenone. Finally, a linear model was used to estimate the effects of *ndufs5* genotype, *ndufa13* genotype, and length (see following section) on corrected respiration rate, with respirometry plate included as a blocking variable. Similarly, we estimated the effects of *ndufs5, ndufa13*, and length on the proportion of respiration showing rotenone sensitivity, with respirometry plate used for blocking. Data in the rotenone-sensitivity regression were filtered to exclude proportions not between 0 and 1, which occurred on occasion due to measurement noise in embryos with extremely low (or no) respiration. Embryos which were damaged during the experiment (e.g. punctured yolks) were excluded from both regression analyses.

The full results of these analyses are reported in Supplementary Tables 10-11. Briefly, *ndufs5* genotype and length had a significant effect on respiration, and both genes had significant interaction effect on respiration with length (alone for *ndufs5* and in a combined three-way interaction in the case of *ndufa13*; Fig. S24-26). This result is consistent with the observation that *ndufa13* incompatible individuals had only slightly lower respiration, but that *ndufa13* genotype impacted change in respiration over development, in concert with *ndufs5* genotype.

We note that in rotenone trials we saw a non-significant trend towards decreased rotenone sensitivity in incompatible embryos (Fig. S27), as might be expected if Complex I function is already low in these individuals. We are cautious in interpreting this response, however, as rotenone trials were necessarily conducted several hours subsequent to baseline measurements, and *Xiphophorus* embryos have reduced viability with increasing time outside of the womb (Supplementary Information 1.3.1).

##### 1.3.5 Morphology of embryos of different genotypes

We were curious how the morphology of hybrid embryos varied as a function of genotype at *ndufs5* and *ndufa13*. From the photos taken after respirometry measurements, we used ImageJ to measure the width of the yolk along three orthogonal axes, the length of the body (excluding caudal fin rays, where applicable), and the width of the head at its widest point (Fig. S14). We were especially interested in cardiac phenotypes, given the prevalence of cardiac abnormalities in vertebrates with mitochondrial dysfunction. Choosing the two phenotypes which are most readily observable across the broadest range of developmental stages, we estimated the heart rate from the 30 second video taken of each embryo, and used ImageJ to measure the width of the sinu-atrium, a muscularized region that contracts peristaltically to pump blood from the yolk venous system into the heart atrium (Fig. S14). We analyzed the data using a linear model with sinu-atrium width, heart rate, head width, or yolk diameter as response variables, and *ndufs5* genotype, *ndufa13* genotype, length, and interactions between them as independent variables. We included brood as a blocking variable. We also performed a regression with length as a response variable, as a function of *ndufs5, ndufa13*, and the relative age of the brood (using embryo length as a proxy, see next paragraph), with brood included as a blocking variable. Embryos that were damaged in the course of the experiment (N = 18, largely due to punctured yolks) were excluded from analyses.

For the analyses described above, we were interested in accounting for the age of the brood, but were unable to control this factor directly since *Xiphophorus* store sperm and fertilization can occur any time from immediately post-mating to months after mating. To overcome this problem, we determined the upper quartile of embryo length in a given brood, which is a continuous measurement that is strongly and monotonically associated with age[13,39]. We used the upper quartile of a brood's length distribution, rather than the median or mean length, because in practice this excluded embryos lagging in developmental stage (i.e. from the *ndufs5* incompatibility). Using a single estimate of brood age is appropriate in *Xiphophorus*, where broods are fertilized simultaneously and develop synchronously[13,39,40]; lagging embryos are a phenotype associated with developmental incompatibility that does not reflect age (Supplementary Information 1.3.1).

The complete results of these analyses are reported in Supplementary Tables 5-9. Overall, we found that genotype at *ndufs5* and *ndufa13* affected a partially overlapping set of phenotypes. Incompatibility at either gene decreased head width and length relative to broodmates, but *ndufs5* had a stronger effect (Fig. S15, Fig. S20-21). Sinu-atrium width was increased in individuals incompatible at *ndufa13*, while incompatibility at either *ndufa13* or *ndufs5* decreased heart rate (Fig. S19, Fig. S22). *ndufs5* incompatible individuals had larger yolks than their compatible broodmates, consistent with decreased respiratory rates in these individuals (Fig. S16). However, they had smaller yolks than compatible individuals of the same size (Fig. S17-18).

##### 1.3.6 Expression of Complex I genes in X. birchmanni, X. malinche, and $F_1$ hybrids

While the large number of substitutions observed between species in *ndufs5, ndufa13*, and Complex I mitochondrial genes suggest that the mitonuclear incompatibility is likely driven by protein-protein interactions involving derived amino acid substitutions (Fig. 4; Table S16), we also wanted to evaluate expression of Complex I nuclear and mitochondrial genes. We generated a reference transcriptome from the annotated *X. birchmanni* genome[5] and used kallisto to pseudoalign reads and quantify abundance for all transcripts[41] from RNAseq data collected from liver tissue of *X. birchmanni*, *X. malinche*, and $F_1$ hybrids (two to five biological replicates each; Table S17). We calculated shrunken log-fold change differences in gene expression between groups using the R program DESeq2[42], using the 'ashr' adaptive shrinkage estimator[43], to evaluate whether there were differences in expression between genotypes in Complex I genes. No expression differences in Complex I nuclear or mitochondrial genes were detected between groups, even at an uncorrected p-value of 0.05. Fig. S28 shows expression levels of genes most likely to be involved directly in the hybrid incompatibility based on mapping and protein modeling results (*ndufs5*, *ndufa13*, *nd2*, and *nd6*).

Because we had access to expression data from the livers of multiple $F_1$ hybrids and *ndufs5* and *ndufa13* differ by several fixed substitutions between *X. birchmanni* and *X. malinche*, we were able to evaluate evidence for allele specific expression at these genes. To do so, we mapped reads to the *X. birchmanni* transcriptome with *bwa*[7] and called variant and invariant sites with *bcftools*[44]. Based on the allelic depth reported in the pileup file, we counted reads supporting the *X. birchmanni* and *X. malinche* alleles at each ancestry informative site >150 basepairs apart in the transcript for each individual. We used an inverted beta-binomial to test for differential expression between alleles[45]. These results are shown in Fig. S29 and Fig. 3F Because of known issues with reference bias in allele specific expression analyses, we repeated the analyses using the *X. malinche* reference transcriptome and found that the results were qualitatively unchanged.

##### 1.3.7 Rt-qPCR to explore mitochondrial copy number

While unlikely based on the expression results, we also wanted to explore evidence for differences in mitochondrial copy number between *X. birchmanni*, *X. malinche*, and $F_1$ hybrids. To evaluate this, we extracted DNA from *X. birchmanni*, *X. malinche*, and $F_1$ liver tissue, and used a qPCR-based approach to quantify mitochondrial copy number relative to single copy nuclear genes. We included 3-6 biological replicates per genotype.

We designed primers for a mitochondrial gene, *ND1*, and for a single copy nuclear gene, *Nup43*, which we identified from the BUSCO Actinopterygii database. We confirmed that this gene was single copy in *Xiphophorus* using annotations for the *X. birchmanni* and *X. malinche* genome assemblies[46]. We confirmed that all primers amplified a single fragment of the expected length before proceeding with the primer sets and evaluated rt-qPCR efficiency for both primer sets (106% for *ND1* and 105% for *Nup43* at an annealing temperature of 66°C). For each biological replicate, we ran rt-qPCR reactions in triplicate for the gene of interest and the control single-copy gene on the same plate. We prepared 10 ul reactions using iQ SYBR Green Supermix according to the manufacturer's specifications in 384 well plates (Bio-Rad, Hercules, CA). We quantified copy number using the CFX384 Touch Real-Time PCR Detection System to run two-step amplification for 45 cycles with an annealing temperature of 66°C (Bio-Rad, Hercules, CA).

We calculated delta Ct between the single copy nuclear and mitochondrial genes (Fig. 3B) and performed an analysis of variance in R to evaluate relative mitochondrial copy number differences between groups. Results of this analysis indicated that there were no significant differences in mitochondrial copy number between groups (ANOVA $F = 1.35$, $P = 0.30$).

##### 1.3.8 Respirometry for OXPHOS Complex activity in parents and hybrids

Individuals with the *X. malinche* mitochondria and homozygous *X. birchmanni* ancestry at *ndufs5* and *ndufa13* typically do not survive to adulthood, making it difficult to evaluate the physiological impact of these interactions outside of the embryonic stage. We reasoned that we might be able to detect some physiological effects of the incompatibility in heterozygotes. To do so, we quantified respiration in isolated mitochondria from *X. birchmanni*, *X. malinche*, and natural hybrids that had the *X. malinche* mitochondria and were heterozygous at *ndufs5* and *ndufa13* (N=7 each group). These hybrids were individually labeled fish from the Calnali Low hybrid population genotyped as described in *Supplementary Information 1.1.3. Low coverage library preparation and sequencing & 1.1.4. Local ancestry inference and mitochondrial haplotype inference*. We also evaluated two fish from this population that had the *X. birchmanni* mitochondria and heterozygous ancestry at *ndufs5* and *ndufa13* but treated them separately in downstream analyses. All hybrid individuals included in physiological tests had similar genome-wide ancestry (average: 45% *X. malinche*; range: 35-58% *X. malinche*). Adult fish of each genotype (weighing 1.2 – 4.1 g) were shipped overnight in breathable bags from our fish facility at Stanford University to the University of Texas in Austin. All fish arrived in good health, except for one *X. malinche* which had similar mitochondrial activity to other pure *X. malinche*, but was nonetheless excluded from statistical analyses.

Immediately upon arrival fish were anesthetized for 5 minutes in ice cold water prior to euthanasia (University of Texas IACUC AUP-2020-00237). Intact, whole livers were then dissected out and placed in ~5 mL ice-cold mitochondrial isolation buffer[47,48] (250 mM sucrose, 50 mM KCl, 25 mM $KH_2PO_4$, 10 mM Hepes, 0.5 mM EGTA, 1.5% bovine serum albumin, pH=7.4 at 20 °C). Liver tissue was homogenized by grinding with a motorized Teflon Potter-Elvehjem pestle for 30 seconds. The homogenate was strained through two layers of cheesecloth and one layer of Miracloth before being aliquoted into 2 mL microcentrifuge tubes and centrifuged for 10 minutes at 600 g and 4 °C to pellet large cellular debris. The resulting supernatant was then centrifuged for 10 minutes at 6000 g and 4 °C to pellet mitochondria. The resulting mitochondrial pellet from each tube was resuspended by pipetting in 500 µL of ice-cold mitochondrial buffer without BSA and combined into a single tube before a centrifuging again for 10 minutes at 6000 g and 4 °C. The resulting mitochondrial pellet was resuspended by pipetting in 200 µL of ice-cold mitochondrial buffer without BSA. This final mitochondrial isolate was then quantified for total protein content using a Qubit Fluorometer (ThermoFisher). Mitochondrial homogenates and isolates were kept on ice throughout the procedure.

Mitochondrial respiration was quantified using the Oroboros O2K respirometry system fitted with small volume (SV) modules. Two O2Ks with two chambers each were used, such that 4 samples were run simultaneously. In a given day, 8 samples were usually run in total, meaning that 4 samples were kept on ice for ~3 hours after isolation prior to being run, which did not affect respiratory phenotypes. Prior to adding the mitochondrial isolate, the O2K chambers were washed 3 times with dd$H_2O$ before being filled with MiR05 respiration buffer (110 mM sucrose, 20 mM HEPES, 10 mM $KH_2PO_4$, 20 mM taurine, 20 mM lactobionic acid, 3 mM $MgCl_2$ 6$H_2O$, 0.6 mM EGTA, 1 g/L BSA). Oxygen content was then air-calibrated prior to each run (a 0% calibration was performed prior to the beginning of this experiment). Following each run, chambers were rinsed with dd$H_2O$, 70% ethanol, and 100% ethanol three times each before being immersed in 100% ethanol for at least 30 min. These washes were then repeated in backwards order (ending with dd$H_2O$) before adding MiR05 and starting the next run.

We then used a multi-substrate, uncoupler, and inhibitor titration (SUIT) protocol to quantify respiration in eight distinct states (Fig. S30). This protocol was adapted from a similar protocol developed for killifish[49] (*Fundulus heteroclitus*). A detailed description of the chemicals added and associated respiratory states is presented in Table S18. A volume of mitochondrial isolate equivalent to 0.15 mg of protein was first added to the chamber, followed by the addition of malate, pyruvate, and glutamate to support mitochondrial NADH production and generate a low-flux "LEAK" respiration state facilitated by proton leak across the inner membrane in the absence of ADP (i.e., state 2 respiration, Fig. S30A). ADP was then added to drive OXPHOS via NADH dehydrogenase (Complex I, CI), dissipating the proton gradient through ATP synthase, which limited electron flow in the preceding state (i.e., state 3 respiration, Fig. S30B). Succinate was then added to supply electrons via succinate dehydrogenase (Complex II, CII), generating the maximal OXPHOS-linked respiration rate observed in the experiment (Fig. S30C). Oligomycin was then added to inhibit ATP synthase (Complex V, CV), resulting in another

"LEAK" state (i.e., state 4 respiration, Fig. S30D). Carbonyl cyanide m-chlorophenyl hydrazone (CCCP) was then added to fully uncouple mitochondria and examine the substrate oxidation capacity of the electron transport system (ETS, Fig. S30E). We then added rotenone (Fig. S30F) followed by malonate (Fig. S30G) to inhibit electron flow from Complex I and Complex II, respectively. Finally, ascorbate (ASC) and N,N,N′,N′-tetramethyl-p-phenylenediamine (TMPD) were added to fully reduce cytochrome c (CytC) and examine the maximal oxygen consumption (reductase) potential of cytochrome c oxidase (Complex IV, CIV, Fig. S30H).

Oxygen concentration was recorded every 2 seconds and oxygen flux was calculated by averaging the previous 10 seconds of data. After the addition of substrates/inhibitors/uncouplers, respiration usually reached a stable state after 2 to 20 minutes, and at least 2 minutes of stable data were recorded for each state. Exceptions were after the addition of CCCP and ASC/TMPD, when respiration usually peaked and then decreased slowly. For these states the peak value was used. Reoxygenation was performed if oxygen content in the chambers dropped below 30 nM $O_2$/mL (~25% of experiments). Chemicals were stored at −20 °C when not in use, except for pyruvate which was made fresh daily.

Respiratory flux control factors (FCFs) were calculated to capture specific aspects of respiratory control[50,51] (Table S19). FCFs are internally normalized, unitless statistics that usually range from 0 to 1 and are calculated within each O2K run. They are not dependent on starting concentrations of mitochondrial isolate and the ability to calculate them is a strength of SUIT protocols. For example, "CI efficiency" represents how much ADP increased respiration relative to the prior respiratory state in each run (analogous to the respiratory control ratio[52]). Seven different FCFs were calculated that represent the extent of respiratory control exerted by specific sites of electron entry and removal from the ETS (Table S19). When calculating the FCFs, we used the following logic to deal with three responses that were problematic in a significant fraction of the runs: 1) in 33% of the runs, after adding succinate respiration increased to a peak and then slowly decreased to a stable value (sometimes plateauing at a value lower than the ADP state, Fig. S17B). In these cases, we calculated the "CII-A" FCF by using the peak succinate value, but calculated the "CI + CII efficiency" FCF by using the lower, stable succinate value. 2) In 33% of the runs, oligomycin failed to significantly decrease respiration, despite repeated additions. We did not use these values when calculating the CI + CII efficiency FCF. 3) In 33% of the runs, CCCP failed to significantly increase respiration, despite multiple additions. We did not use these values when calculating the "ETS capacity" FCF. Finally, we calculated the maximum respiration observed after normalizing to protein input. This was taken as the highest stable value when either succinate or CCCP was added (whichever was highest).

We focused our statistical analysis on Complex I efficiency, since *ndufs5* and *ndufa13* localize to Complex I. To test the effect of ancestry heterozygosity at *ndufs5* and *ndufa13* on Complex I efficiency, we constructed an additive linear model with Complex I efficiency as the response variable, genotype as the explanatory variable, and experimental date as a blocking variable (Complex I efficiency ~ genotype + date). We then performed an orthogonal contrast between heterozygous hybrids with *X. malinche* mitochondria and parental individuals, revealing a significant difference in Complex I efficiency between heterozygotes and the two parental genotypes ($t$ = -2.53, $P$ = 0.023, n = 7 per genotype). We also calculated the time after the addition of ADP before respiration reached its maximum value in State B ("Time to peak CI", Fig. S32), and found that hybrids were significantly slower to reach this peak (orthogonal contrast $t$ = 4.303, $P$ < 0.001; Fig. 3C; Fig. S32). While Complex I efficiency can also be affected by the integrity of the mitochondrial membrane, neither measurement of LEAK state respiration (Fig. S32, $t$ = -1.213, $P$ = 0.24, n = 7) nor flow cytometry assays (Fig. S34; Supplementary Information 1.3.9) showed differences in mitochondrial membrane integrity between genotypes, pointing to reduced function of Complex I in hybrids. Likewise, the time to peak respiration after activation of Complex II by addition of succinate was similar across genotypes (orthogonal contrast $t$ = -0.705, $P$ = 0.49; Fig. S33).

To explore the possibility that limiting reaction steps downstream of Complex I are driving this observed change in Complex I efficiency, we used the same statistical framework to test for differences in other components of the electron transport chain. We analyzed the maximum oxygen consumption of Complex IV alone (known as excess CIV capacity) to investigate differences in Complex IV function. We also analyzed a metric known as CI + CII efficiency, which measures the change in respiration after the inhibition of Complex V. Since any downstream member of the respiratory chain which limits Complex I efficiency is also predicted to affect CI + CII efficiency, similarities in this ratio are suggestive of similarities in the function of Complexes III, IV, and/or V. Neither excess CIV capacity (orthogonal contrast $t$ = -1.250, $P$ = 0.23, n = 7 per genotype) nor CI + CII efficiency (orthogonal contrast $t$ = 1.44, $P$ = 0.193, n = 4 for *X. birchmanni*, 3 for hybrids, and 6 for *X. malinche*) showed significant differences between hybrids and parentals. This suggests that differences in Complex I function in hybrids are the most likely cause of observed differences in Complex I efficiency.

##### 1.3.9 Evaluation of mitochondrial membrane depolarization

The Complex I efficiency FCF is a relative measure of oxygen flux with and without Complex I-driven respiration, and is thus sensitive to changes in oxygen flux in either state. As such, the reduced Complex I efficiency detected in individuals heterozygous for the hybrid incompatibility in the previous section could reflect a decrease in Complex I activity, or an increase in proton leak due to changes in permeability of the mitochondrial membrane. To exclude this second possibility, we pursued two independent measures of mitochondrial membrane integrity. First, we calculated LEAK state respiration from our Oroboros data (Supplementary Information 1.3.8) as the rate of oxygen consumption prior to ADP addition, normalized to protein input. LEAK respiration did not differ between hybrids and parentals (orthogonal contrast $t$ = -1.213, $P$ = 0.24, n = 7 per genotype), suggesting that membrane integrity does not drive reduced Complex I efficiency in hybrids.

Second, we measured mitochondrial membrane potential in hybrid individuals heterozygous at *ndufs5* and *ndufa13* and in parental species. Mitochondrial membrane potential was measured by flow cytometry in dissociated fish fibroblast cells using TMRE (tetramethylrhodamine ethyl ester, Abcam, ab113852), with FCCP used as a positive control to test sensitivity to depolarization across genotypes. Dissociated live cells were collected from fin tissue derived from *X. birchmanni*, *X. malinche*, and $F_1$ hybrid individuals. For each biological replicate (three per genotype), fin tissue was pooled from three individuals to achieve a yield of ~500,000 cells. Each sample was washed in 100X penicillin/streptomycin for 5 minutes, and subsequently washed four times in 4X penicillin/streptomycin diluted in fish PBS (100 mM NaCl, 2 mM KCl, 6 mM $Na_2HPO_4$, 1 mM $KH_2PO_4$, pH 7.4) for 5 minutes. Tissue was shredded using a sterile scalpel and incubated in 5 mL of 0.05% Trypsin-EDTA on a shaker for 2 hours at room temperature. Trypsinization was stopped by adding fetal bovine serum at a final concentration of 20%. The cell suspension was transferred to a new tube to remove remnants of tissue and spun down for 10 minutes at 450 *g*. Cells were resuspended in fish PBS at a concentration of 1x10$^6$ cells/mL. Cells were then incubated with 20 μM FCCP or DMSO for 10 minutes at 25°C. After incubation, cells were stained with 200 nM of TMRE for 20 min at 25°C. Then, cells were spun down at 500 *g* for 5 min at 25°C and resuspended in 200μL of fish PBS with DAPI.

Cells were immediately moved to ice and analyzed by flow cytometry, using an Acea Novocyte Quanteon. TMRE measurements were performed with a 561 nM excitation laser and a 586/20 filter channel. Gates were used to identify live cells and to separate singlets from doublets. For each sample 10,000 events on the Live/Singlet gate were recorded. An additional gate to identify a TMRE+ population was drawn based on *X. malinche* disassociated cells treated with FCCP. The mean TMRE fluorescence intensity for each genotype was measured over 3 biological replicates before and after the addition of FCCP. Statistical analysis using an ANOVA showed no significant differences in baseline membrane potential (untreated median fluorescence log-transformed for normality, $F$ = 3.505, $P$ = 0.10; Fig. S34) or in response to membrane permeabilization between genotypes (proportion of untreated fluorescence remaining after FCCP treatment, $F$ = 0.196, $P$ = 0.83), suggesting that mitochondrial proton leakage does not substantially differ between parental species and hybrids. We therefore conclude that the differences in Complex I efficiency identified in our respirometry assays are unlikely to result from changes in membrane polarization, and are thus attributable to changes in Complex I activity.

#### 1.4 Mitochondrial protein quantitation and structural modeling

##### 1.4.1 Mitochondrial isolation

To isolate mitochondria for proteomic experiments, we anesthetized fish in ice cold water prior to euthanasia (following Stanford IACUC protocol #33071). We dissected out whole liver from each individual and placed it in a pre-weighed tube of 1 mL ice-cold mitochondrial isolation buffer (IB, 1 mM EGTA, 10 mM $K_2PO_4$, 1% bovine serum albumin, 250 mM sucrose, pH 7.4 at 20°C), then weighed the tube to ascertain liver weight. We transferred the liver into 5 volumes of ice-cold IB in a 2 mL Dounce homogenizer and homogenized with 5 passes with a loose-fitting pestle followed by 2 passes with a tight-fitting pestle. We transferred the homogenate to a 1.5 mL microcentrifuge tube and centrifuged at 4 °C for 5 minutes at 650 g to pellet large cellular debris. We then transferred the resulting supernatant to a fresh 1.5 mL tube and centrifuged at 4 °C for another 15 minutes at 9600 g to pellet mitochondria. We removed the supernatant and resuspended the mitochondrial pellet by gently pipetting in 5 volumes of ice-cold IB before centrifuging again at 4 °C for 15 minutes at 9600 g. We then repeated this round of discarding, resuspension, and centrifugation. Finally, we discarded the supernatant and resuspended the final mitochondrial pellet by pipetting in 5 volumes of isotonic buffer (100 mM KCl, 25 mM $K_2PO_4$, 5 mM $MgCl_2$, pH 7.4 at 20°C). This final mitochondrial isolate was then quantified for total protein content using a Pierce BCA Protein Assay Kit (ThermoFisher). We kept mitochondrial homogenates and isolates on ice throughout the procedure, and froze the mitochondrial solution at -80°C after isolation.

##### 1.4.2 Heavy labelled peptide design for protein quantitation

To test for evidence of disproportionate degradation of particular Complex I protein combinations in hybrids, we took a mass spectrometry-based quantitative proteomics approach.

Specifically, we used heavy stable isotope labeled peptides to distinguish between specific *X. birchmanni* and *X. malinche* peptides in mitochondrial proteomes extracted from $F_1$ hybrids. The reasons for this approach are two-fold. First, the labeled peptides can be used to direct the mass spectrometer to collect more data from the fraction of the sample containing the peptides of interest. Second, this approach facilitates quantification of peptides of interest, as the internal standard controls for peptide-specific ionization and detectability properties[53].

Heavy-labelled peptides are designed to mimic likely products of the protease digestion used to prepare endogenous proteins for mass spectrometry. As such, we created *in silico* trypsin digestions (cutting at every lysine and arginine residue) for the *X. malinche* and *X. birchmanni* alleles of *ndufs5* and *ndufa13*. For each gene, we selected a single peptide shared across both species' alleles, and peptides where species' alleles were distinguishable by one or more amino acid substitution (Table S20). We ordered unpurified aliquots of these peptides with $^{13}C$- and $^{15}N$-labeled arginine or lysine at the C terminus, and cysteine residues carbamoylated with iodoacetamide. High performance liquid chromatography mass spectrometry (HPLC-MS) performed by JPT (Berlin, Germany) suggested that the purity of the aliquots ranged from 37.5% to 67.7% and averaged 54.9%. Peak integration of the UV-spectra from HPLC-MS quantified a total of 16.87 nmol of peptide WLLLQSGEQPYK (NDUFS5 *X. malinche* target) and 21.74 nmol of peptide WLLPQSGEQPYK (NDUFS5 *X. birchmanni* target) in the final product used for downstream steps.

##### 1.4.3 Sample preparation for mass spectrometry

For mass spectrometry, mitochondrial isolates were prepared in 50 mM ammonium bicarbonate in the presence of protease max surfactant (Promega) followed by reduction with 10 mM DTT at 55°C for 30 min. Following reduction, proteins were alkylated with 30 mM of 2-Iodoacetamide for 30 min at room temperature in the dark. Digestion was performed with Trypsin/LysC (Promega) in a typical overnight digest at 37°C. Proteolysis was quenched using 1% formic acid and peptides were de-salted on C18 Monospin reversed phase columns (GL Sciences). The de-salted peptides were dried in a speed vacuum before reconstitution in appropriate volume for peptide quantification using the Pierce quantitative fluorometric peptide assay (Thermo Scientific, San Jose, CA). Following quantification, peptides were re-dried and re-constituted in a final reconstitution buffer (2% acetonitrile with 0.1% Formic acid) along with the peptide pool so that peptide loading could be matched across samples.

##### 1.4.4 Parallel Reaction Monitoring experiment

Mass spectrometry experiments were performed on a Q Exactive HF-X Hybrid Quadrupole – Orbitrap mass spectrometer (Thermo Scientific, San Jose, CA) with liquid chromatography performed using a Nanoacquity UPLC (Waters Corporation, Milford, MA). For a typical LC-MS experiment, a pulled-and-packed fused silica C18 reverse phase column was used, with Dr. Maisch 1.8 micron C18 beads as the packing material and a length of ~25 cm. A flow rate of 300 nL/minutes was used with a mobile phase A of 0.2% formic acid in water and mobile phase B of 0.2% formic acid in acetonitrile. Peptides were directly injected onto the analytical column using a gradient (3-45% B, followed by a high-B wash) of 80 minutes. The mass spectrometer was operated using a parallel reaction monitoring (PRM) method for ion selection. HCD fragmentation was used for MS/MS spectra generation. MS/MS resolution was

120,000 (at *m/z* 200) with an AGC target value of $1 \times 10^6$ ions, a maximum fill time of 250 ms and an isolation window of 1.0 *m/z*.

##### 1.4.5 Statistical analysis of Parallel Reaction Monitoring data

For data analysis, the .RAW data files were imported into Skyline 19.1.0.193 (MacCoss Lab, Department of Genome Sciences, UW) to generate XIC and perform peak integration (Fig. S35). Peptide identification was additionally confirmed using Byonic (Protein Metrics, Cupertino, CA). Using Skyline, we called the target peptides' spectral peaks for each individual and species version such that the window captured the signal from the heavy-labeled spike-in peptide, then applied the same retention time interval to detect the endogenous version. The spectral intensities observed in all targeted peptides except the WLL[L/P]QSGEQPYK pair were below the expected sensitivity limits of our Parallel Reaction Monitoring protocol. We therefore focused our statistical analyses on the WLL[L/P]QSGEQPYK pair. We calculated the total area under the intensity vs. retention time curve peak to represent the abundance of each peptide in the sample and used these integration values to calculate the proportion of total intensity attributable to the *X. malinche* peptide. To match the 50:50 null expectation for endogenous peptides, we normalized the intensity values for the heavy-labeled spike-in by the known ratio of the peptides which were added to the samples prior to MS. Specifically, *X. malinche* intensities were divided by 16.87 and *X. birchmanni* intensities were divided by 21.74 such that the expected proportion of adjusted *X. malinche* intensity would be 50% in the absence of differences in ionization efficiency and detectability.

With this correction, the observed proportion of peptides derived from the *X. malinche* allele was not significantly different from 50% in the heavy-labeled peptides (mean = 47%, one-sample *t*-test $P = 0.13$), but was in the endogenous peptides (mean = 55%, one-sample *t*-test $t = 3.96$, $P = 0.02$). To test the sensitivity of our results to our choice of retention intervals from which peak areas were calculated, we repeated our analysis with wide (retention time encompassing all the detectable signal from the heavy-labelled peptides) and narrow (retention time only encompassing the unimodal area around the highest intensity peak) integration intervals. Both alternatives increased the average fraction *X. malinche* in both heavy-labeled (mean = 52%) and endogenous (mean = 57%) peptides; however, the heavy-labeled peptide ratio remained statistically indistinguishable from 50% ($P = 0.10$ for both alternatives), and the ratio for endogenous peptides remained significantly greater than 50% ($P = 0.005$ for both alternatives).

##### 1.4.6 Complex I structure prediction

Since *ndufs5* and *ndufa13* both form part of Complex I of the mitochondrial electron transport chain, we hypothesized that the position of substitutions within Complex I could generate hypotheses about likely interactions between the proteins involved. We therefore set out to generate a structural model for *ndufs5*, *ndufa13*, and neighboring genes in Complex I, using Cryo-EM structures generated for mammalian model species (see below). Alignments of amino acid sequences from *X. birchmanni, X. malinche*, and mammalian model sequences are available in Supplementary File 1, and are visualized with predicted secondary structure for *X. birchmanni* and *X. malinche* in Fig. S58 and Fig. S59. The number of Complex I supernumerary subunits was identical between mammalian species and swordtails, and the lengths of these subunit was very similar (Table S13), suggesting that the use of mammalian models to predict the structure of *Xiphophorus* Complex I proteins is appropriate. Amino acid sequence identity between mammalian species and swordtails varies based on the subunit in question (Table S12), but sequence similarity (quantified by BLOSUM 45 parameters) exceeds 70% in all comparisons (Table S12).

We submitted the amino acid sequences for *ndufs5* and *ndufa13* inferred from the *X. birchmanni* and *X. malinche* genome sequences to the RaptorX protein structure prediction web server (http://raptorx.uchicago.edu), which identified the mouse Cryo-EM structure (PDB ID 6G2J) as the best-fit folding template for both sequences. To ensure that the observed model was not sensitive to the choice of template, we additionally used the MODELLER protein structure prediction package to fold *X. birchmanni ndufs5* and *ndufa13* with mouse (6G2J), sheep (5LNK), cow (5LDW), and human (5XTC) structures as a template. Predicted structures for swordtails mirror those of the mammalian templates (Fig. S37). The position of all seven substitutions observed between the two species in *ndufs5* and *ndufa13* were unchanged across templates, and we therefore consider our inferences about these two structures to be insensitive to folding template (Fig. S39). We then aligned the RaptorX models to the mouse structure (6G2J) in PyMol, and performed a second round of RaptorX structure prediction for the mitochondrial (*nd2, nd3, nd4l, nd6*) and nuclear (*ndufa1*, *ndufa8*, *ndufb5*, *ndufc2*) genes in contact with the predicted NDUFS5 and NDUFA13 structure in the mouse Cryo-EM structure. Structures were predicted for both the *X. birchmanni* and *X. malinche* version in cases where amino acid sequences differed between species (all genes but *ndufb5* and *ndufa1*). Depending on the gene, RaptorX chose mouse (6G2J), sheep (5LNK), cow (5LDW), and human (5XTC) sequences as modelling templates, but the two *Xiphophorus* species' sequences of a given gene were always matched to the same model. Alignment of *Xiphophorus* models and their templates suggests that tertiary structure is highly conserved in these genes (Fig. S37) despite divergence in amino acid sequences (Fig. S38; Table S12). We again used MODELLER to test the sensitivity of the structures to the template chosen, and found that in general the predicted structures were insensitive to the template that was used. For all templates tested, the variation between predicted structures was limited to regions without secondary structure, and in *ndufa13* and *nd2* the differences between templates were comparable to those between five alternative models with the same template (Fig. S39). Results using different templates suggest that *in silico* models are relatively insensitive to choices made in modeling the structure.

One exception to this was the case of structures predicted for *nd6.* For *nd6,* mammalian templates varied in the presence or absence of a β-hairpin and an α-helix in the coil between transmembrane helices 4 and 5, which is precisely where *nd6* contacts both *ndufs5* and *ndufa13*. Depending on the choice of mammalian model, resulting predictions for *Xiphophorus* proteins differed substantially. Specifically, four clustered substitutions in the coil of *nd6* shifted in position relative to *ndufs5* and *ndufa13* (Fig. S39, Table S14). Thus, inferences about direct contact and the distances between residues in this particular interhelix coil should be taken with caution.

Despite this uncertainty, all four substitutions in the *nd6* coil are predicted to fall within potential contact distance of *ndufs5* (α carbon distance < 10 Å) in one or more models (Table S14). Thus, we thus speculate that this cluster could a point of mitonuclear interaction between *X. birchmanni ndufs5* and *X. malinche* mitochondrial genes (Table S14). It is intriguing to note that one *nd6* substitution in particular (*malinche* M to *birchmanni* A at residue 120) mirrors the substitution found in *ndufs5* (*malinche* A to *birchmanni* M at residue 31). Investigating whether complexes composed of *X. malinche nd6* and *X. birchmanni ndufs5* suffer from steric clashes between two large methionine side chains could be an exciting direction for future work when genetic manipulation becomes possible in *Xiphophorus*.

In addition to the predicted physical interaction with *ndufs5*, the same cluster of *nd6* substitutions is potentially in contact with one *ndufa13* substitution, making this region even more interesting in light of evidence for interactions between these three proteins (Supplementary Information 1.2.5). Outside this region, one *ndufs5* substitution is predicted to directly contact a substitution in *nd2* (Fig. 4B, Fig. S40, Table S14), and two other pairs of substitutions, in *ndufa13* and *nd6* and *ndufs5* and *nd6* respectively, may also be close enough to physically interact (Fig 4A, Fig. S40-41). Based on these computational results, we predict that *nd2* and *nd6* are the genes most likely to be involved in incompatible interactions with *ndufs5* and *ndufa13* (Fig. 4A, 4B; Fig. S41-S44). Testing this question will be an exciting direction for future work when transgenic or cell culture approaches become possible in *Xiphophorus* species.

#### 1.5 Phylogenetic analyses

##### 1.5.1 <u>Analysis of evolutionary rates and intolerant mutations</u>

The nuclear and mitochondrial sequences we collected for *X. birchmanni* and *X. malinche* also allowed us to identify sequence differences between species and evaluate their evolutionary rates and possible functional effects. We used the function codeml in PAML[54] to identify nonsynonymous and synonymous substitutions in each gene in each species. We identified nonsynonymous substitutions between *X. birchmanni* and *X. malinche* and high dN/dS ratios in *ndufs5* and *ndufa13* ($dN/dS_{ndufs5} > 99$, $N_{ndufs5} = 4$, $S_{ndufs5} = 0$; $dN/dS_{ndufa13} = 1.2$, $N_{ndufa13} = 3$, $S_{ndufa13} = 1$). Although overall substitution rates were extremely high in mitochondrial-encoded Complex I genes (Fig. S42), dN/dS ratios were lower (0.085 – 0.21). After examining inferred maximum likelihood trees, we determined that branch lengths in the *ndufs5* and *ndufa13* gene tree were suggestive of different evolutionary rates across species. As a result, we also used codeml to evaluate evidence for a significantly different dN/dS ratio for these genes on the branch leading to *X. birchmanni* using a likelihood ratio test (species included: *X. birchmanni*, *X. malinche*, *X. cortezi*, *X. pygmaeus*, *X. nezahualcotoyl*, *X. montezumae*, *X. helleri*, *X. couchianus*, *X. variatus*, and *X. maculatus*). For *ndufs5*, this test supported a significantly different rate for the branch leading to *X. birchmanni* (Likelihood Ratio Test P = 0.005; Table S15). Repeating this analysis for *ndufa13*, we also found evidence for a significantly different dN/dS ratio on the branch leading to *X. birchmanni* (Likelihood Ratio Test P = 0.002; Table S15). Results for Complex I mitochondrial genes in contact with *ndufs5* or *ndufa13* are reported in Table S15, where we find significantly different evolutionary rates in *nd6* and *nd4*.

Given results indicating differences in evolutionary rates in several Complex I proteins along the *X. birchmanni* lineage, we wanted to test for evidence that dN/dS was greater than 1 in these genes, which would be suggestive of adaptive rather than neutral evolution. We used codeml to perform a likelihood ratio test comparing a model where omega was fixed at 1 on the *X. birchmanni* lineage and where it was not fixed. Based on this analysis, we could not reject a model of dN/dS = 1 on the *X. birchmanni* branch for the nuclear genes in question (Table S15), indicating that relaxed selection could explain the accumulation of substitutions along the *X. birchmanni* lineage. However, we note that our other results indicate that there is variation in constraint among sites in Complex I proteins (Fig. S45), which complicates interpretation of this test.

We also wanted to determine whether the nonsynonymous mutations we identified in *ndufs5* and *ndufa13* were likely to change protein function. To do so, we used a database of *ndufs5* (N=197) and *ndufa13* (N=118) for bony fish downloaded from NCBI's protein database. This dataset included *Xiphophorus helleri* and *X. couchianus* but not *X. birchmanni* and *X. malinche*. We aligned these peptide sequences using Clustal Omega and input the alignments into SIFT [55], which predicts the probability that an amino acid at a given position will be tolerated, assuming that the most frequently occurring amino acids at that position are tolerated. For each derived substitution in *X. birchmanni* and *X. malinche* relative to the outgroup (*X. hellerii*) we asked whether that amino acid is predicted by SIFT to be tolerated at that site in the alignment.

We found that all amino acid positions that differed between *X. birchmanni* and *X. malinche* at *ndufs5* and *ndufa13* were derived in *X. birchmanni* relative to the outgroup. Of these two out of four substitutions in *X. birchmanni ndufs5* and two of three substitutions in *X. birchmanni ndufa13* were predicted to not be tolerated by SIFT (Table S16). We repeated SIFT analyses for the four mitochondrial genes modelled using RaptorX, and found that fewer substitutions were predicted to be not tolerated despite higher substitution rates[29] (Table S16).

##### 1.5.2 Tests for coevolution of mitochondrial and nuclear genes over evolutionary timescales

Given the hypothesis that mitonuclear genes undergo compensatory coevolution driven by rapidly changing mitochondrial genes[31], we were curious whether *ndufs5* and its physical interactors bore phylogenetic signals of coevolution. To test this, we applied the GREMLIN statistical model[56] to detect coevolution signals through comparative genomics. Four separate analyses were performed through the GREMLIN web portal (http://gremlin.bakerlab.org/submit.php), pairing *ndufs5* and *nd2, ndufs5* and *nd6*, *ndufs5* and *ndufa13,* or *ndufa13* and *nd6* for analysis.

To generate the input data, we gathered *ndufs5* and *ndufa13* sequences using three iterations of PSI-BLAST with their respective *X. birchmanni* sequences as the query, and kept those within 20% of the length of the *X. birchmanni* sequence. These sequences were aligned using Clustal Omega, after which we removed sites with >25% gaps, realigned, and then retained only those sequences with >20% identity to *Xiphophorus birchmanni*. In cases where more than one sequence was available for a given taxon, we retained the one with lowest divergence from the *X. Birchmanni* gene. After these steps, the filtered alignments were 105 and 118 amino acids long for *ndufs5* and *ndufa13*, and a total of 722 *ndufs5* and 1929 *ndufa13* sequences from unique taxa were left for coevolution analysis.

We then found matching sequences for each specific comparison of *nd2* or *nd6* to *ndufs5* or *ndufa13* through a three-step process. We first performed a single PSI-BLAST iteration, then supplemented this with a BLASTP search limited to the taxa for which *ndufs5* or *ndufa13* sequences had been obtained, using the *X. birchmanni* sequence as the query. Finally, we searched for sequences in the GenBank nucleotide and protein databases from those species with *ndufs5* sequences but still missing data for *nd2, nd6,* or *ndufa13*. Where an unannotated mitochondrial sequence for a species was available, the focal gene was identified by alignment of a closely related annotated sequence, translated computationally, and retained if it contained no stop codons. These sequence alignments were then filtered with Clustal Omega alignment followed by removal of sites with >25% gaps (high mitochondrial sequence divergence meant that a 20% divergence filter could not be applied while retaining enough sequences for analysis). Discarding sites with >25% gaps left 312, 169, and 143 aligned sites from *nd2, nd6*, and *ndufa13* for comparison to *ndufs5,* respectively, and 147 sites for *nd6* when matched with *ndufa13*. The alignment steps resulted in 637, 578, and 510 sequences for *ndufs5* coevolution analysis in *nd2*, *nd6,* and *ndufa13*, respectively, and 699 for the coevolution of *ndufa13* and *nd6*. The HHBlits iterations parameter was set to 0 to perform analyses on the provided MSA, and default filters of 75% coverage and 75% gaps were used. The GREMLIN output was then aligned to the RaptorX models for each structure in GREMLIN Beta to visualize how signals of coevolution related to physical distance.

No significant signals of intermolecular coevolution were identified between *ndufs5* and *nd2* or *ndufa13* and *nd6*, with all interactions with probability >90% being intramolecular (Fig S60). *ndufs5* and *nd6,* on the other hand, showed two intermolecular contacts with probability >90%, between *ndufs5* residue Y27 and *nd6* H110, as well as *ndufs5* A78 and *nd6* M91. Notably, the inferred interactions are not in physical contact based on in our RaptorX structures (Fig. S36, S40). This counterintuitive result could be attributable to true coevolution between physically distant sites or discrepancies between the true and predicted structure of *Xiphophorus nd6*, among other factors (discussed in Supplementary Information 1.4.6).

Finally, we observed strong signals of coevolution between *ndufs5* and *ndufa13*, with three significantly coevolving locus pairs at a 95% probability threshold and another at a 90% threshold (Fig. S60). However, none of these coevolving locus pairs included sites with substitutions between *X. birchmanni* and *X. malinche* (Fig. S36). The stronger signals of coevolution between *ndufs5* and *ndufa13* compared to *ndufs5* and *nd2* or *nd6* is consistent with comparisons between *Xiphophorus* and mammalian template sequences, which suggested that the *ndufs5-ndufa13* interface is much more strongly conserved than the interface with either mitochondrial gene (Fig. S31).

Although it is intriguing that we do not detect coevolution between residues predicted to be in physical contact, we caution against strong interpretations of GREMLIN results, as our power to detect coevolution was low. The GREMLIN documentation recommends that the ratio of number of sequences to sequence length be at least 5 for accurate inference, but our exhaustive searches only resulted in a ratio of 1-2 depending on the sequence pair. Thus, the true number of coevolving sites for these proteins may be substantially greater.

As an additional test for signals of coevolution, we utilized the RaptorX-Contact server[57]. We used the same input data as described above, except that we converted ambiguous residues to gaps. RaptorX-Contact (which differs from RaptorX template-based structure prediction) uses coevolution information from a multiple sequence alignment to predict distances between residues, then feeds this information into a deep neural network trained on known structures to infer intermolecular contacts. As such, the inference of correct intermolecular contacts by RaptorX-Contact is suggestive of coevolution but is also impacted by training on known structures.

RaptorX-Contact was unable to identify the regions of contact between *nd2* and *ndufs5* or between *nd6* and *ndufa13* (Fig. S61). The majority of the *ndufa13* residues expected to contact *nd6* were discarded from analysis due to frequent gaps in the multiple sequence alignment, and RaptorX-Contact only roughly colocalized the remaining *ndufa13* residues and their *nd6* neighbors (inter-alpha carbon distance 10-20 Å, Fig. S61). In contrast, it identified the contact interface between the cluster of four substitutions in *nd6* and the region containing residue 31 of *ndufs5* (Fig. S61). Likewise, it reconstructed the interface between *ndufs5* and *ndufa13* with high accuracy (Fig. S61). These results are consistent with coevolution between *ndufs5* and *nd6* in the region where we observe contacts between interspecific substitutions, and even tighter coevolution between *ndufs5* and *ndufa13* (Fig. 4B). However, they should be viewed with caution given that RaptorX-Contact uses not only inferred coevolution information but also training datasets of known structures in generating predictions.

##### 1.5.3 Construction of phylogenies for mitonuclear interactors

We were interested in inferring the evolutionary history of the mitochondria in *X. birchmanni*, *X. malinche*, and their relatives. As a first step to infer phylogenetic relationships, we used Illumina resequencing data to infer *ndufs5, ndufa13,* and mitochondrial sequences for available species. We mapped reads to the *X. birchmanni* reference genome[5] from previously sequenced individuals for *X. birchmanni, X. malinche, X. cortezi*, *X. hellerii*, and *X. maculatus* (SRA accessions in Table S21) as well as newly sequenced individuals from *X. pygmaeus* and from a divergent lineage of *X. malinche* from the Las Juntas Tributary locality in the Rio Tuxpan drainage (SRA BioProject PRJNA744894). After mapping, we removed duplicates using Picard tools and called variants using GATK[8,9]. We filtered variants using hard-calls as described previously and removed sites with >2X or <0.5X genome-wide (or mitochondria-wide) coverage[10]. We generated individual pseudoreferences using the *X. birchmanni* assembly coordinates but for each individual we updated variant sites that passed our quality filters and masked variant and invariant sites that did not.

We then extracted mitochondrial sequences and sequences at *ndufs5* and *ndufa13* and built maximum likelihood phylogenies for each gene using RaxML[58]. As previously reported, this phylogeny revealed that *X. malinche* and *X. cortezi* are most closely related based on their mitochondrial sequences alone while *X. malinche* and *X. birchmanni* are sister species based on the nuclear genome[59]. Our inclusion of a more divergent *X. malinche* population in this analysis allowed us to infer that variation in the *X. cortezi* mitochondrial haplotype is nested within variation observed in *X. malinche*. This suggests that if gene flow is the cause of the observed mitochondrial discordance (see *Supplementary Information 1.5.4)*, the direction of introgression would have been from *X. malinche* into *X. cortezi*.

To further investigate mitochondrial phylogenetic relationships in a larger number of individuals, we performed long-read amplicon sequencing. We designed two primer sets to amplify 7 kb regions of the mitochondrial genome. We extracted DNA from fin clip tissue using a Dneasy Kit (Qiagen, Valencia, CA) and amplified both regions in ten individuals (*X. birchmanni* n = 5, *X. malinche* n = 3, *X. cortezi* n = 2) using the LongAmp Taq PCR kit (Qiagen, Valencia, CA). We purified amplicons using 18% SPRI magnetic beads and confirmed their fragment sizes using the Genomic DNA kit on the TapeStation 4150 (Agilent, Santa Clara, CA). We sent purified amplicons to Admera Health (South Plainfield, NJ) for HiFi long read library preparation and sequencing on an 8M SMRTcell (Pacific Biosciences, Menlo Park, CA). Per-amplicon coverage ranged from 180-1800X; raw data are available through NCBI's sequence read archive (SRA BioProject PRJNA744894).

We then used the PacBio Long Amplicon Analysis (pbaa; https://github.com/PacificBiosciences/pbAA) application to cluster HiFi reads within a sample before generating a consensus sequence for each cluster. For amplicons where reads did not collapse to a single cluster, we chose a random cluster consensus for further analysis. We then combined the amplicons of the two mitochondrial regions for an individual to generate a consensus sequence for the full mitochondrial haplotype, and used Clustal Omega to align these haplotypes along with the *X. birchmanni* and *X. malinche* reference mitochondrial sequences. This alignment was used to generate a whole mitochondrial phylogeny (Fig. S46), and to estimate within-species diversity in mitochondrial haplotypes ($\pi$) and pairwise sequence divergence ($d_{XY}$) of *X. birchmanni, X. malinche,* and *X. cortezi* mitochondrial genomes (Table S22).

1.5.4 Introgression of an *X. malinche* mitochondrial haplotype incompatible with *X. birchmanni ndufs5* and *ndufa13*

Our phylogenetic analysis indicates that the mitochondrial haplotypes of *X. cortezi* individuals are more closely related to *X. malinche* than either set of sequences is to *X. birchmanni*, in conflict with the species tree topology. Two major processes can generate discordance between gene trees and species trees: introgression and incomplete lineage sorting.

In nuclear genomes, lengths of discordant ancestry tracts can provide information about whether hybridization is a likely cause of discordance, as can local signatures of genetic divergence relative to expectations from the divergence time of the two species. Since mitochondrial sequences are non-recombining, we are unable to use ancestry tract lengths and instead asked whether mitochondrial haplotypes between *X. cortezi* and *X. malinche* have unexpectedly low genetic differentiation, consistent with recent gene flow. To do we used a simulation-based approach.

Mitochondrial mutation rates are thought to greatly exceed nuclear mutation rates in most animal species[60]. As a proxy for relative mitochondrial mutation rates in swordtails, we evaluated $D_{xy}$ between pairs of swordtail species for which we could calculate both nuclear and mitochondrial divergence (Table S23) and neither mitochondrial or nuclear data were suggestive of gene flow[59]. We found that mitochondrial sequence divergence between swordtail species pairs satisfying these criteria ranged from 3.1 – 5 times the observed nuclear sequence divergence. Since lower mitochondrial mutation rates are conservative for our analysis, we proceeded with the minimum (3.1) and average (4) mitochondrial to nuclear divergence ratio.

We next evaluated expected mitochondrial divergence between *X. malinche* and *X. cortezi* using SLiM[61] and the previously inferred demographic histories of *X. malinche* and *X. cortezi*[10,21]. We defined a 16.6 kb genomic element with no recombination in SliM. We used the "Y" chromosome functionality of SLiM to simulate uniparental inheritance of this genomic element. We initialized simulations with a population size of 51,500 individuals for two subpopulations 453,000 generations before the present, based on previous estimates of the long-term effective population size of *X. cortezi* and divergence times between *X. malinche*, *X. cortezi*, and *X. birchmanni*[10,21]. Pairwise sequentially Markovian coalescent (PSMC) results for *X. malinche* indicate that this species experienced a strong and persistent bottlenecked through much of its recent history[10] (Fig. S62). Thus, we simulated a reduction in population size in one lineage implemented as a strong bottleneck reducing the population to 5,000 individuals, starting 20,000 generations before the present. Using the mitochondrial to nuclear substitution ratios calculated above, and the estimated nuclear mutation rate for cichlids[62] ($3.5 \times 10^{-9}$) we performed simulations with mutation rates of $1.09 \times 10^{-8}$ and $1.4 \times 10^{-8}$ in separate sets of simulations. For each scenario we performed 500 replicate simulations and summarized pairwise mitochondrial divergence between one individual from the simulated *X. malinche* population and one individual from the simulated *X. cortezi* population.

Results of these simulations indicate that observed mitochondrial divergence between *X. malinche* and *X. cortezi* is much lower than would be expected given their split time and demographic history, even in simulations assuming a lower mitochondrial mutation rate (Fig. 5C). Combined with our phylogenetic results, this suggests the *X. malinche* mitochondria introgressed into *X. cortezi*.

Given evidence of introgression of the *X. malinche* mitochondria into *X. cortezi*, we were curious if the interacting genes in the incompatibility, *ndufs5* and *ndufa13*, had also introgressed. Maximum likelihood phylogenies of *ndufs5* and *ndufa13* had low bootstrap support (Fig. S47-48), suggesting insufficient information for phylogenetic reconstruction in this region. However, *X. cortezi* and *X. malinche* have identical nucleotide sequences at *ndufs5*, and both sequences are deeply differentiated from *X. birchmanni* at this gene (Fig. 5D). Similarly, the amino acid sequences are identical between *X. cortezi* and *X. malinche* at *ndufa13*. Divergence between *X. birchmanni* and *X. malinche* in both sequences is ~2X higher than that observed between *X. malinche* and *X. cortezi*. This pattern is suggestive co-introgression of *X. malinche ndufs5* and *ndufa13* with the *X. malinche* mitochondrial genome into *X. cortezi*.

##### 1.5.5 Genome-wide analysis of ancient *X. cortezi* × *X. malinche* admixture

Given evidence for mitochondrial introgression from *X. malinche* into *X. cortezi*, we wanted to evaluate evidence for admixture in the nuclear genome. Since *X. malinche* and *X. cortezi* are not currently known to be sympatric, we assumed that this is likely an ancient admixture event and thus chose to use site sharing approaches such as the D-statistic which can detect ancient as well as recent admixture.

To perform this analysis, we took advantage of the divergent lineage of *X. malinche* sampled from Las Juntas Tributary, described above. This individual is an outgroup to the mitochondrial clade containing other *X. malinche* populations and *X. cortezi* (Fig. 5B), and thus presumably diverged from other *X. malinche* populations prior to the introgression event. We evaluated support for two different alternative topologies to the species tree, one that groups *X. malinche* with *X. cortezi* and one that groups the Las Juntas Tributary individual with *X. cortezi*.

We performed this analysis using available pseudoreferences for samples from each population. We used a custom script to convert variant information in pseudoreference files into Admixtools format (https://github.com/Schumerlab/Lab_shared_scripts). Using the qpDstat function we calculated support for the two alternative 'ABBA' and 'BABA' topologies using a block-jackknife bootstrapping approach with a block size of 5 Mb. Based on this analysis we found evidence for gene flow between *X. malinche* and *X. cortezi* since the divergence of most *X. malinche* populations from the Las Juntas Tributary lineage, although the signal is modest (D=0.05, Z=2.1).

##### 1.5.6 Evidence for mitonuclear incompatibility in natural *X. birchmanni* × *X. cortezi* hybrid populations

Given evidence that the mitochondrial sequence of *X. malinche* introgressed into *X. cortezi*, and that *X. malinche* and *X. cortezi* share nearly identical amino acid sequences both at mitochondrial *nd2* and *nd6* genes and the nuclear genes *ndufs5* and *ndufa13*, it is possible that *X.*

*birchmanni* × *X. cortezi* will have the same mitonuclear incompatibility as is found in *X. birchmanni* × *X. malinche* hybrids. While we have been unable to produce a sufficient number of crosses in the lab of *X. birchmanni* × *X. cortezi* hybrids to evaluate this directly, we can take advantage of natural hybrid populations between *X. birchmanni* × *X. cortezi* to investigate this possibility.

Through past work we have identified three geographically distinct *X. birchmanni* × *X. cortezi* hybrid populations[21], Huextetitla, Santa Cruz, and a previously undescribed population: Chapulhuacanito. These populations derive 75-90% of their genome-wide ancestry from *X. cortezi* and based on demographic inference appear to have formed approximately 150-200 generations ago[21]. Like most *X. birchmanni* × *X. malinche* populations, all three hybrid populations are fixed for the mitochondrial haplotype from species from which they derived the majority of their genome, in this case *X. cortezi* (Fig. S50). In Huextetitla and Chapulhuacanito this means mitochondrial ancestry falls in the upper 5% and 1% of all 16 kb regions genome-wide, respectively. In Santa Cruz, mitochondrial ancestry is the most extreme observed in all 16 kb regions genome-wide (>0.01%).

While this extreme mitochondrial ancestry is striking, given the mechanisms of mitochondrial inheritance, it could also reflect sex-biased migration or assortative mating. Thus, we next turned to evaluating two-locus genotypes at the mitochondria and *ndufs5*. Out of our sample of 284 individuals, we found no individuals that harbored a genotype that is predicted to be incompatible (i.e. homozygous *X. birchmanni* at *ndufs5*). We repeated this analysis for *ndufa13* and found the same results. However, given the frequency of *X. birchmanni* ancestry nearby *ndufs5* and *ndufa13* in Huextetitla, Santa Cruz, and Chapulhuacanito, we predict that we would need to sample thousands of individuals to reject the hypothesis that the combination of *X. cortezi* mitochondria and homozygous *X. birchmanni* ancestry at *ndufs5* or *ndufa13* were absent by chance.

Nonetheless, extreme ancestry at *ndufs5* and *ndufa13* in the *X. cortezi* × *X. birchmanni* hybrid populations is itself a signal of selection on the mitonuclear DMI identified by our mapping results in *X. birchmanni* × *X. malinche*. Ancestry at *ndufs5* falls into the upper 5% tail of major parent ancestry genome-wide in the Huextetitla population, and in the upper 8% tail of major parent ancestry genome-wide in the Santa Cruz and Chapulhuacanito populations. For *ndufa13*, ancestry falls in the upper 2% tail of major parent ancestry genome-wide in the Santa Cruz population, the upper 4% tail for Chapulhuacanito, and the upper 8% tail in the Huextetitla population.

We assessed about the likelihood that all three populations would have low minor parent ancestry in the regions surrounding *ndufs5* and *ndufa13* by chance using simulations. We calculated average major parent ancestry in 0.1 cM windows in both populations. Binning the data into windows of genetic distance ensures that we do not over-sample regions of the genome with low recombination rates. Next, we randomly sampled a 0.1 cM window from each population and repeated this procedure 10,000 times. We asked how frequently we sampled a pair of windows by chance with equivalent or higher major parent ancestry to that observed at *ndufs5* and *ndufa13* in Huextetitla, Santa Cruz, and Chapulhuacanito. Based on these simulations, the co-occurrence of low minor parent ancestry at *ndufs5* and *ndufa13* in Huextetitla, Santa Cruz, and Chapulhuacanito is unexpected by chance (*ndufs5* p-value by simulation = 0.0004; *ndufa13* p-value by simulation = 0.0001).

#### 2 Supplementary Figures

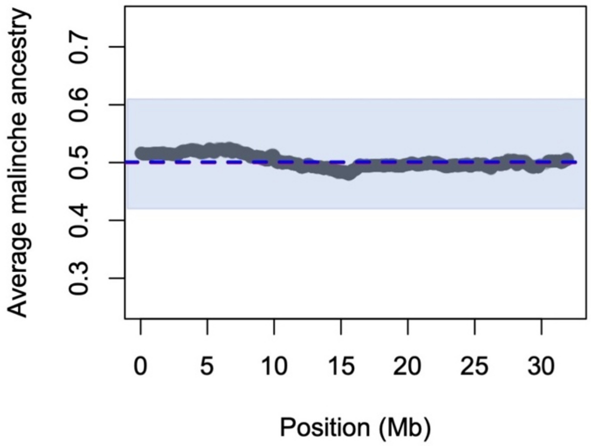

**Fig. S1.** The early generation hybrids analyzed in this project and others[1,2] are $F_1$ intercrosses. This leads to the expectation that at ancestry informative sites throughout the genome $F_2$ hybrids will have an average ancestry of 50% *X. malinche*. While we observe substantial segregation distortion on chromosome 13, genome-wide we find that individuals derive 50.3% of their ancestry from *X. malinche* and that average ancestry across most chromosomes meets expectations, as shown here for chromosome 1. Black dots indicate average ancestry across 943 individuals at ancestry informative sites, blue envelop indicates the 99% confidence intervals of ancestry genome-wide, and blue line indicates the expected ancestry across individuals for this cross.

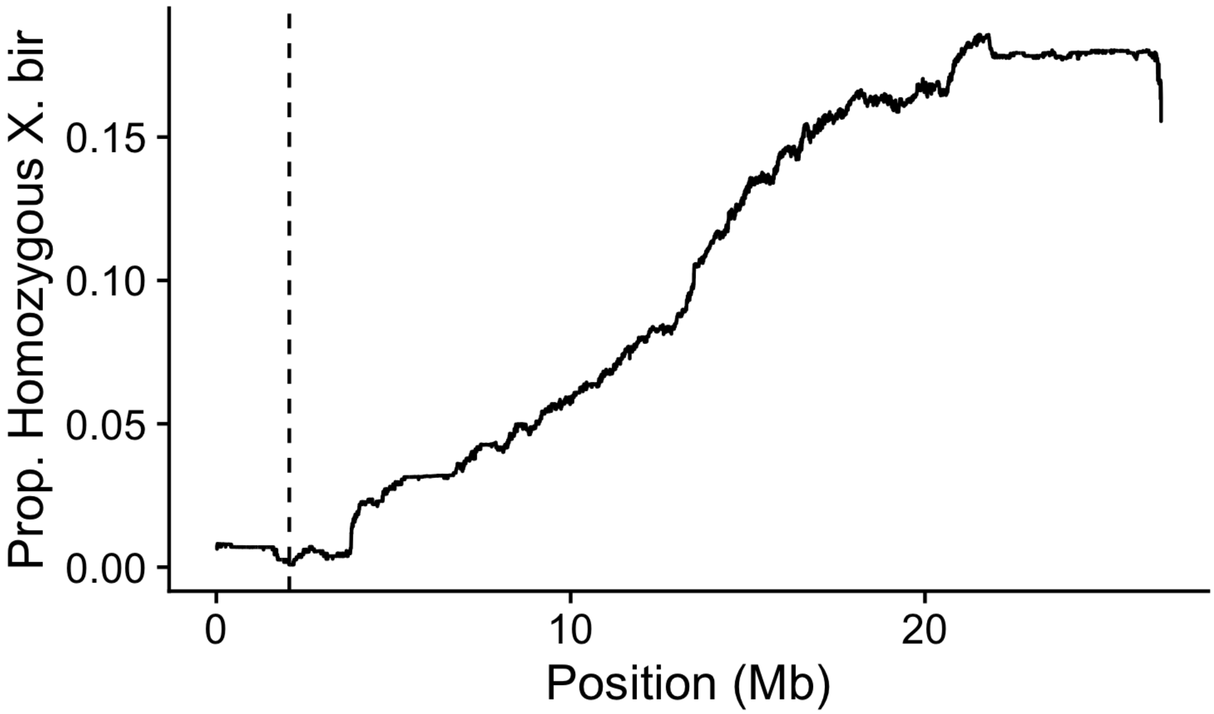

**Fig. S2.** Frequency of homozygous *X. birchmanni* ancestry in $F_2$ hybrids along chromosome 13. We find nearly 100% depletion of homozygous *X. birchmanni* ancestry at *ndufs5* (2.06 Mb, dashed line) which increases towards expected levels (25%, maximum observed on chromosome 13: 22%) with increasing distance from the incompatible site.

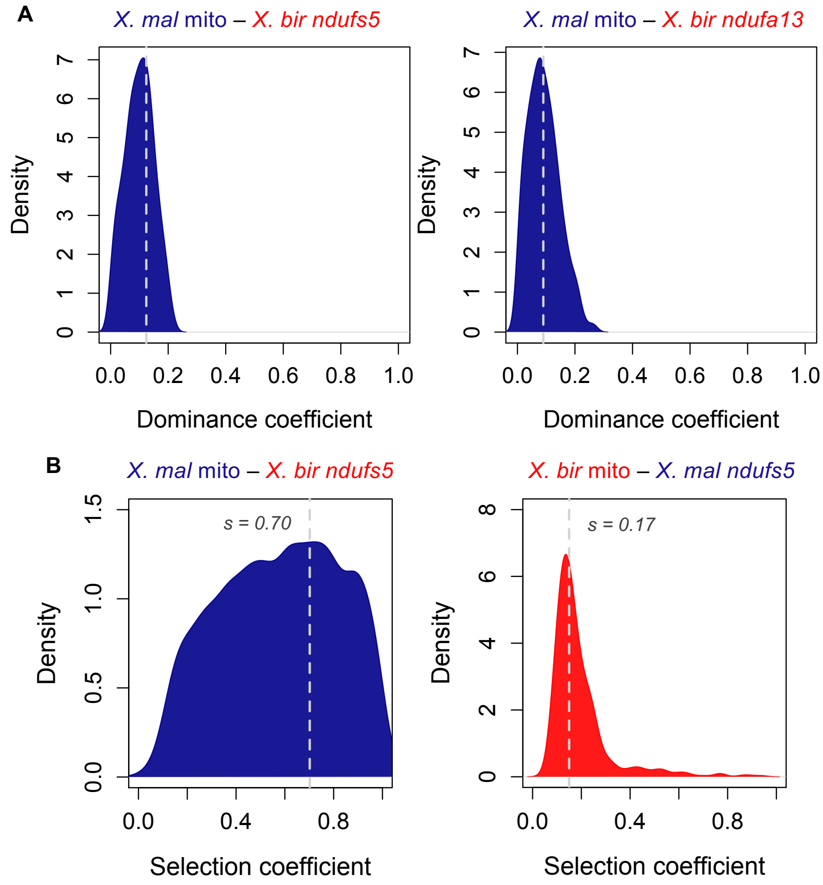

**Fig. S3.** (**A**) Posterior distributions of dominance coefficients from approximate Bayesian computation (ABC) simulations fitting observed data in $F_2$ hybrids. The simulations that resulted in the posterior distribution shown on the left modeled the *X. malinche* mitochondria – *X. birchmanni ndufs5* component of the hybrid incompatibility and the simulations that generated the posterior distribution on the right modeled *X. malinche* mitochondria – *X. birchmanni ndufa13* component of the hybrid incompatibility. Each posterior distribution shows the accepted dominance coefficients in 500 simulations and the gray dashed line indicates the maximum a posteriori estimate for the dominance coefficients of interactions involving *ndufs5* and *ndufa13* (at $h$ = 0.12 and $h$ = 0.09 respectively). (**B**) Posterior distributions of selection coefficients from ABC simulations fitting observed data in the Calnali Low hybrid population. The simulations that resulted in the posterior distribution shown on the left modeled the *X. malinche* mitochondria – *X. birchmanni ndufs5* incompatibility and the simulations that generated the posterior distribution on the right modeled the reverse *X. birchmanni* mitochondria – *X. malinche ndufs5* incompatibility. In the main text we focus on inferred selection coefficients from $F_2$ hybrids for the incompatibility involving the *X. malinche* mitochondria (see Fig. 1) but present results from fitting the Calnali Low population data here and elsewhere in the supplement (see *Supplementary Information 1.2.2 Estimating the strength of selection against incompatible genotype combinations using lab-bred F2 and natural hybrids*). Distribution shows the accepted selection coefficients in 500 simulations and the gray dashed line indicates the maximum a posteriori estimate. Note that we recover a much broader distribution and a lower maximum a posteriori estimate of the selection coefficient for the *X. malinche* mitochondria – *X. birchmanni* *ndufs5* incompatibility here compared to $F_2$ simulations (Fig. 1). We interpret this result to be driven by the fact that simulations fitting the Calnali Low population data modeled >20 generations of admixture. Thus, a range of selection coefficients are consistent with the observation that no individuals have *X. malinche* mitochondrial ancestry and homozygous *X. birchmanni* ancestry at *ndufs5* in the Calnali Low hybrid population.

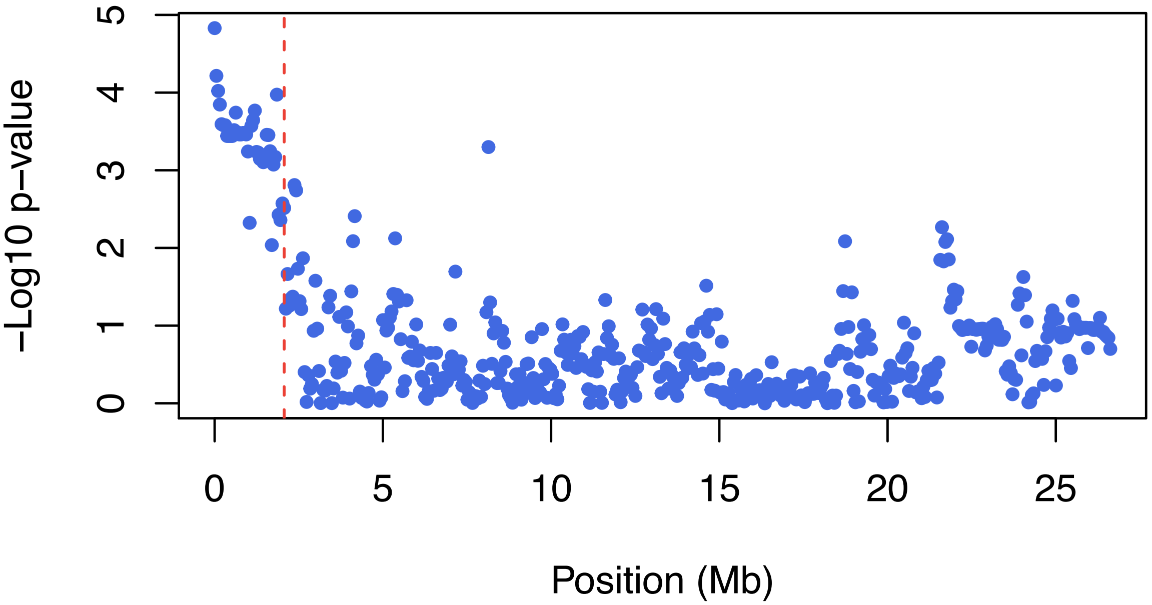

**Fig. S4.** Association between mitochondrial ancestry and nuclear genotype on chromosome 13 in the Chahuaco Falls hybrid population. Plotted here is the -$Log_{10}$(p-value) of the partial correlation between mitochondrial and chromosome 13 ancestry, after accounting for genome-wide ancestry. While the Chahuaco Falls population segregates for mitochondrial ancestry, there are few individuals with *X. birchmanni* haplotypes (~7% of our sample), suggesting that we will have lower power to map associations in this population. Nevertheless, we detect an association between mitochondrial ancestry and chromosome 13 ancestry concordant with that found in our primary mapping population (Fig. 1G; Fig S7). Red dashed line indicates the position of *ndufs5*.

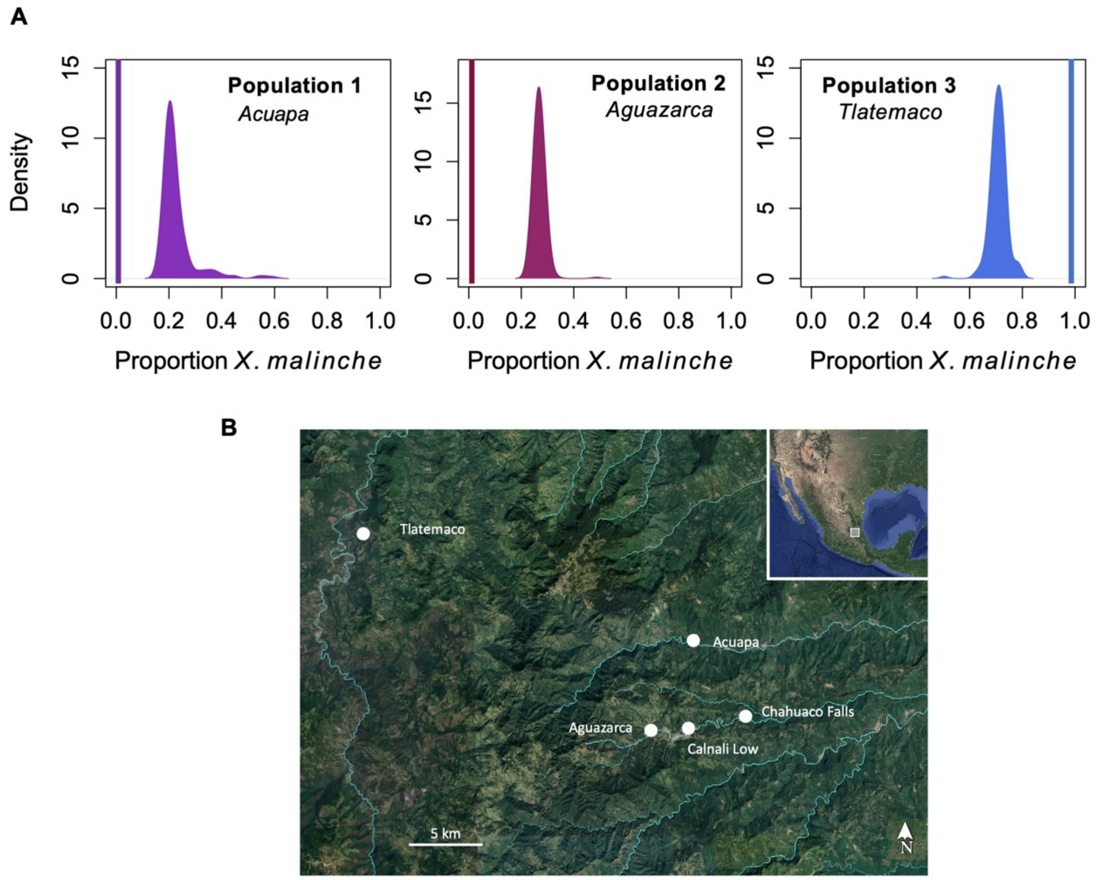

**Fig. S5**. (**A**) Average genome-wide ancestry distribution in three hybrid populations fixed for mitochondrial ancestry from a single parent species. Curves show the distribution of genome-wide ancestry proportions across individuals within each population. Vertical bars represent the mitochondrial ancestry fixed in that population. (**B**) Map of *X. birchmanni* × *malinche* hybrid sites on different river systems used in this study.

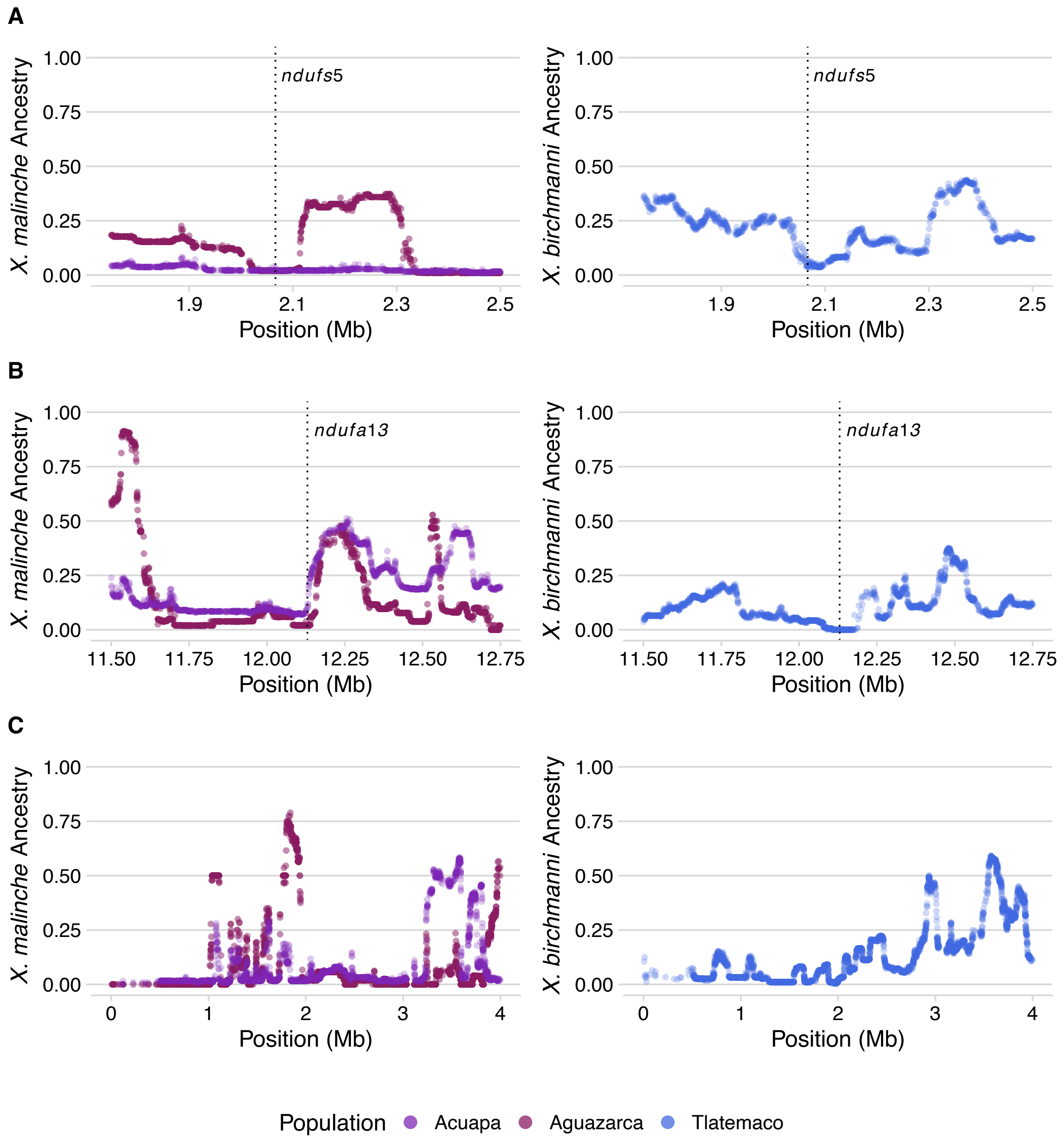

**Fig. S6**. Non-mitochondrial parent ancestry on (**A**) chromosome 13, (**B**) chromosome 6, and (**C**) chromosome 15 in natural hybrid populations fixed for *X. birchmanni* (Aguazarca, Acuapa, left column) or *X. malinche* (Tlatemaco, right column) mitochondrial haplotypes. Vertical dashed lines represent the position of *ndufs5* and *ndufa13* within the admixture mapping regions. (**A**) Non-mitochondrial parent ancestry is depleted around *ndufs5* in all natural hybrid populations that are fixed for a particular mitochondrial haplotype. This pattern is strongly suggestive of a history of selection on this region in natural hybrid populations. In (**B**), *ndufa13* is fixed for *X. malinche* ancestry only in the population with *X. malinche* mitochondria (Tlatemaco), mirroring the architecture of the genetic interaction between this gene and the mitochondria. Specifically, interactions with *ndufa13* are only expected to be under selection in combination with the *X. malinche* mitochondrial haplotype (Fig. 1C).

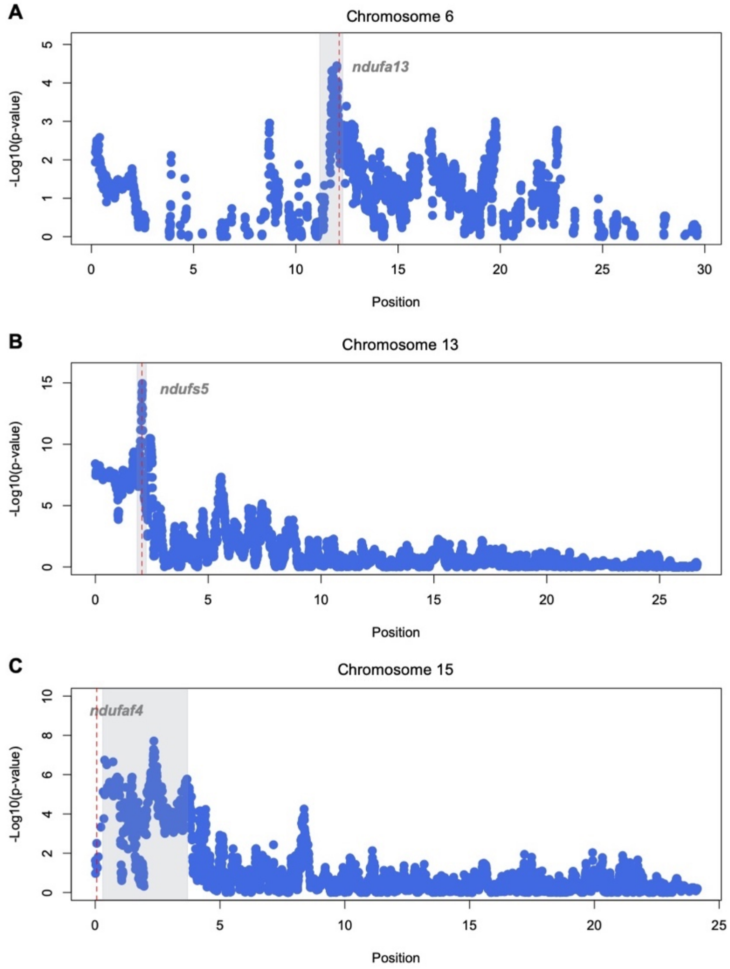

**Fig. S7.** Plots of admixture mapping results focusing on chromosomes with signals at or near the genome-wide significance threshold. Also plotted with dashed lines are the locations of Complex I genes within (**A** & **B)** or close to (**C**) each admixture mapping peak. Gray envelopes indicate the peak associated interval. Note that the Chromosome 13 envelope is widened by 250 kb to improve visibility. Significance threshold was calculated based on thinned markers as described in Supplementary Information 1.1.5 but here plotted points reflect the unthinned data.

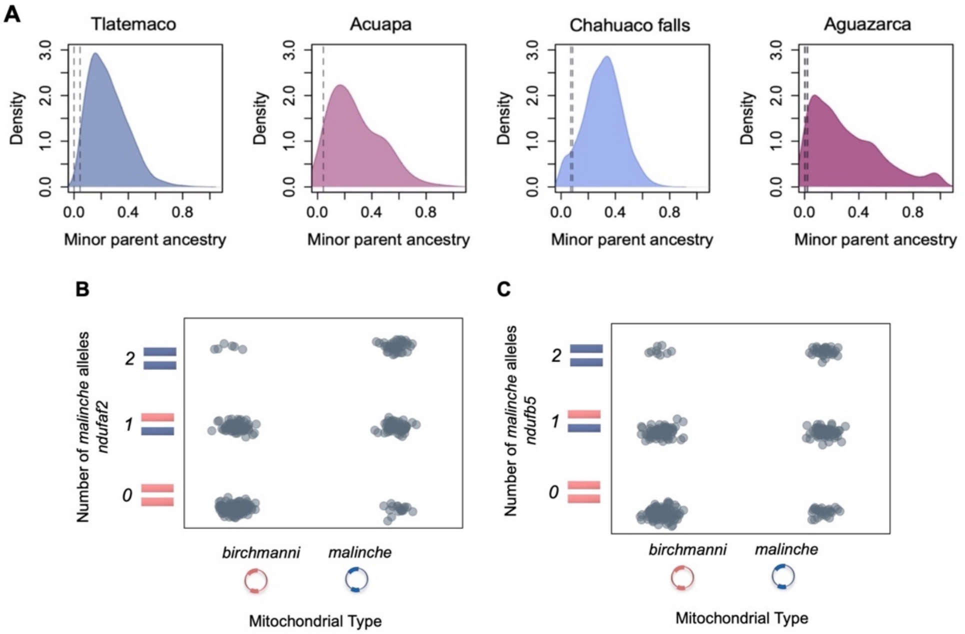

**Fig. S8. (A)** Genome-wide ancestry distribution in four hybrid populations and ancestry at Complex I genes *ndufa13* and *ndufaf4*, which show extreme ancestry in multiple populations. In each population the most common mitochondrial haplotype is that derived from the major parent ancestry. Dashed lines show ancestry in each population at *ndufa13* and *ndufaf4*. **B & C** Both *ndufaf2* and *ndufb5* show extreme ancestry in Acuapa and Aguazarca, two populations that have fixed the *X. birchmanni* mitochondrial haplotype. However, interactions between these genes and the mitochondrial haplotype are not significant in the admixture mapping population, even without a multiple hypothesis testing correction ($P = 0.13$ and $P = 0.22$ respectively).

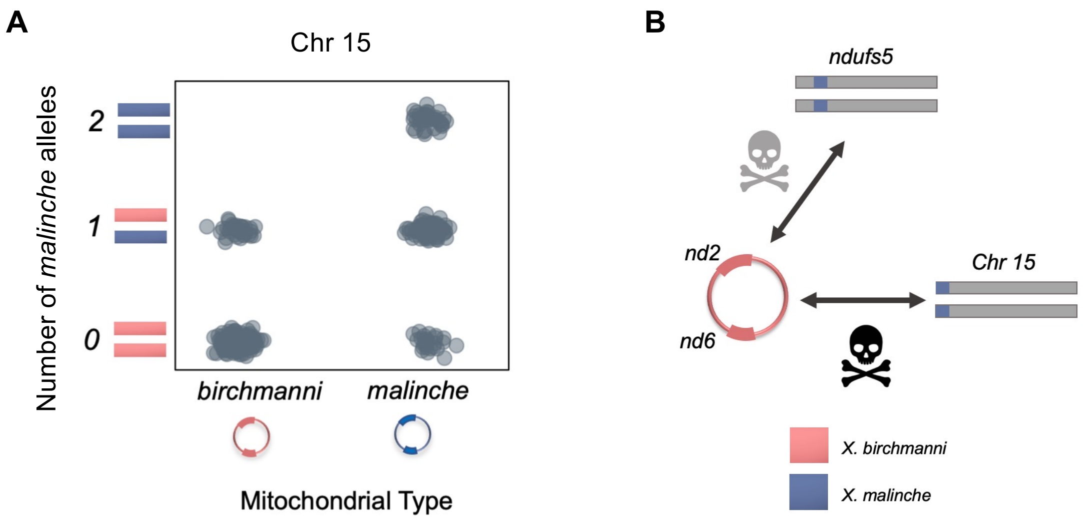

**Fig. S9.** (**A**) Observed genotype frequencies at the peak associated marker (3.37 Mb) on chromosome 15 in the admixture mapping population. (**B**) Schematic of identified interactions with the *X. birchmanni* mitochondrial genome from our mapping data and strength of selection underlying each interaction in hybrids (gray skull – moderate, black skull– near lethal). We discuss interactions with the *X. birchmanni* mitochondria in more detail in Supplementary Information 1.2.1-1.2.2.

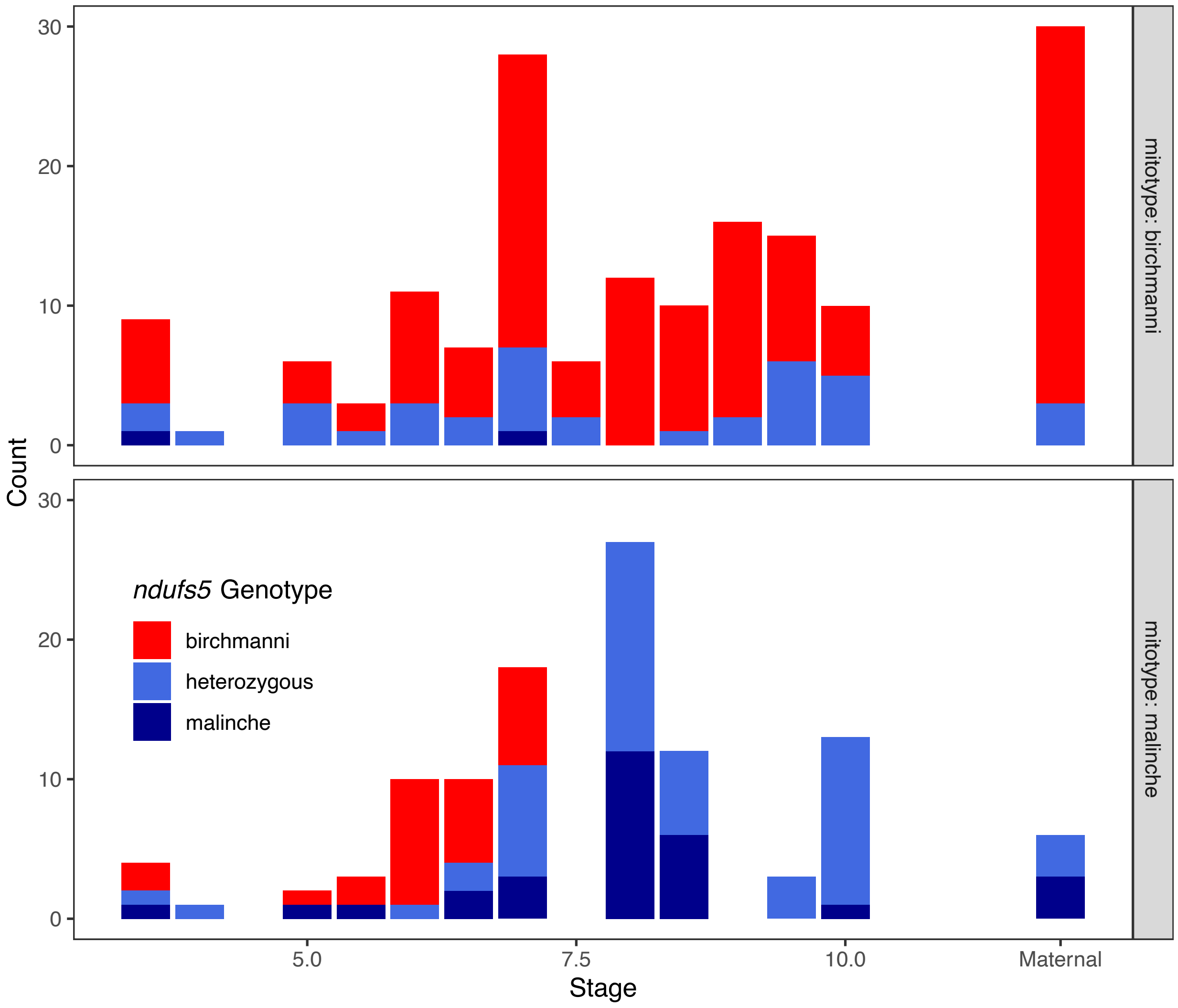

**Fig. S10**. Developmental stage and *ndufs5* genotypes of all embryos sequenced from the Calnali Low population. Genotypes of mothers are shown in the rightmost column. Panels are split by mitochondrial ancestry. Only two embryos were sampled with the incompatible genotype between the *X. birchmanni* mitochondrial genome and *X. malinche* ancestry at *ndufs5* (top). This makes us unable to evaluate whether this direction of the mitonuclear incompatibility impacts progression through embryonic development.

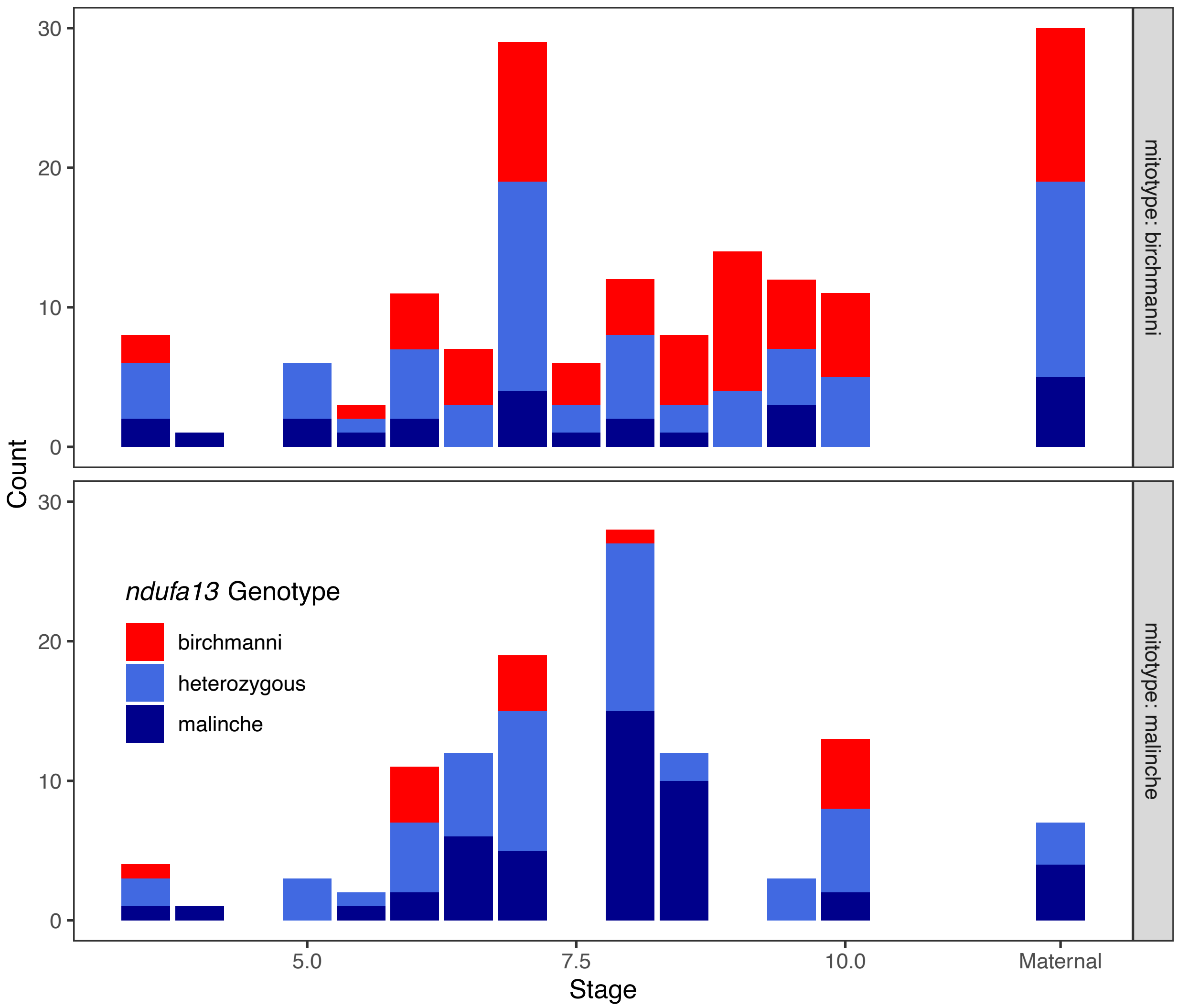

**Fig. S11**. Developmental stage and *ndufa13* genotypes of all embryos sequenced from the Calnali Low population. Genotypes of mothers are shown in the rightmost column. We do not detect a significant effect of *ndufa13* genotype on developmental stage in either embryos with *X. birchmanni* derived mitochondria (top) or *X. malinche* derived mitochondria (bottom).

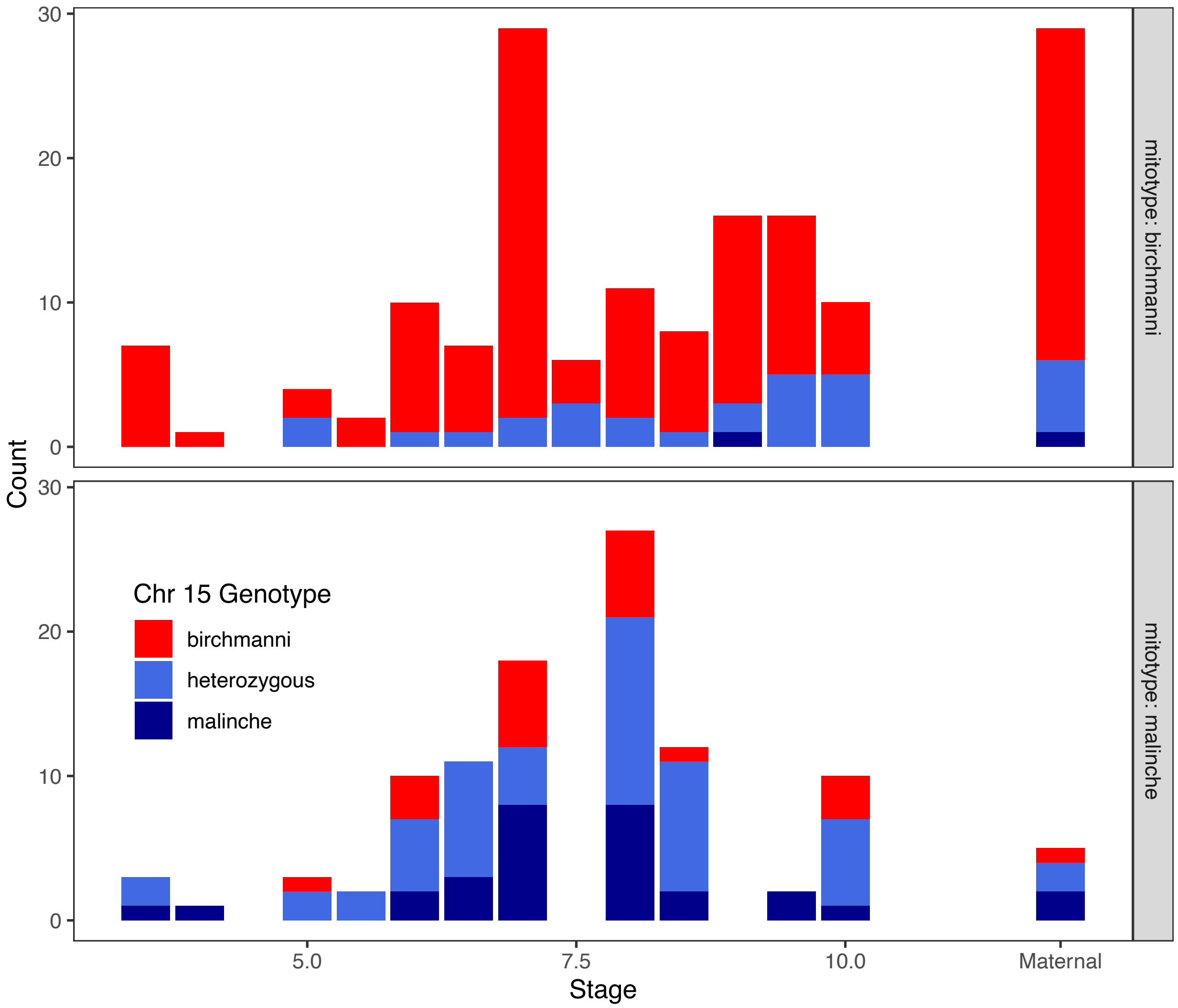

**Fig. S12**. Developmental stage and genotypes at the peak associated marker on chromosome 15 (3.37 Mb) of all embryos sequenced from the Calnali Low population. Genotypes of mothers are shown in the rightmost column. We do not detect a significant effect of chromosome 15 genotype on developmental stage in either embryos with *X. birchmanni* derived mitochondria (top) or *X. malinche* derived mitochondria (bottom).

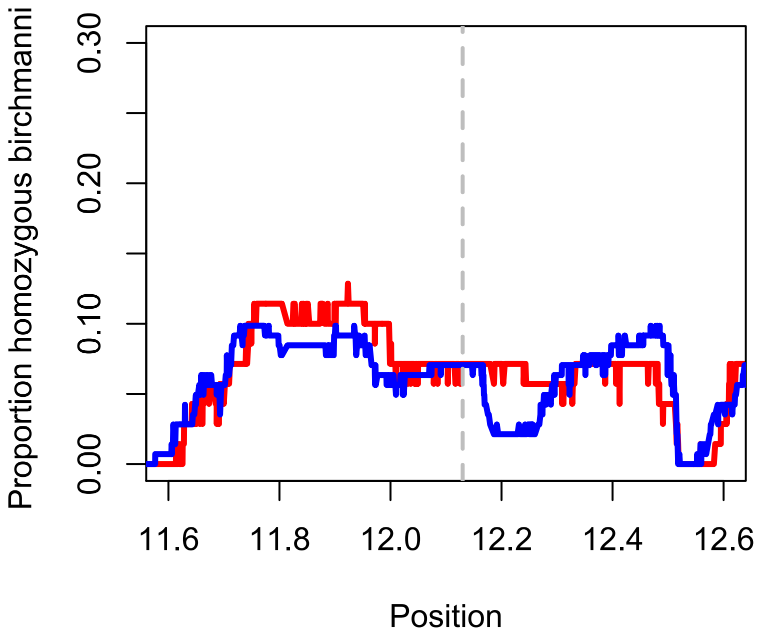

**Fig. S13.** Frequency of homozygous *X. birchmanni* ancestry along chromosome 6 in embryos with *X. malinche* mitochondria that lagged their siblings in developmental stage by ≥1 developmental stage (red) versus the frequency of homozygous *X. birchmanni* ancestry in embryos that did not exhibit developmental lag (blue, see Supplementary Information 1.1.7). Dashed line indicates the location of *ndufa13*.

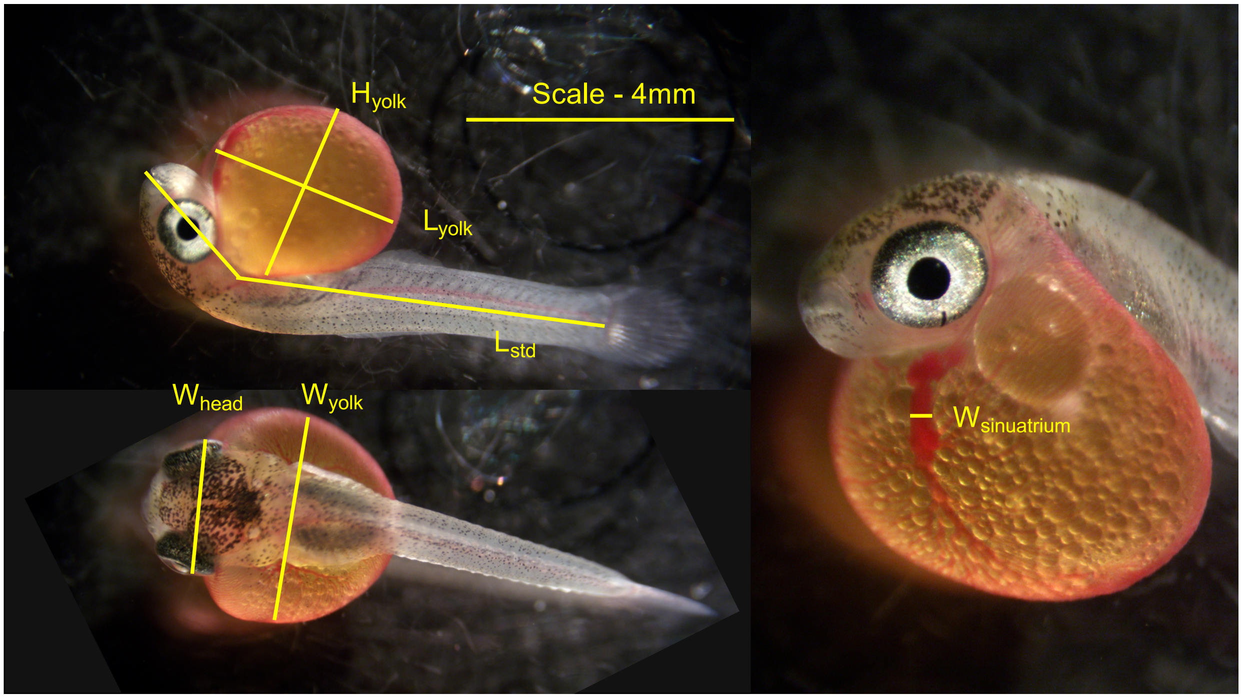

**Fig. S14**. Composite illustration of morphometric measurements taken on $F_2$ embryos after respirometry trials. To account for asymmetries in the yolk, the yolk diameter was calculated as the mean of yolk length ($L_{yolk}$), width ($W_{yolk}$), and height ($H_{yolk}$). Head width ($W_{head}$) was measured at the widest point from above, and standard length ($L_{std}$) was measured from snout to the base of the caudal rays, in one or two segments as needed to account for curvature around the yolk. Sinu-atrium width was measured at the midpoint of the area observed to contract peristaltically in video footage. The scale in each photo was set with reference to a stage micrometer.

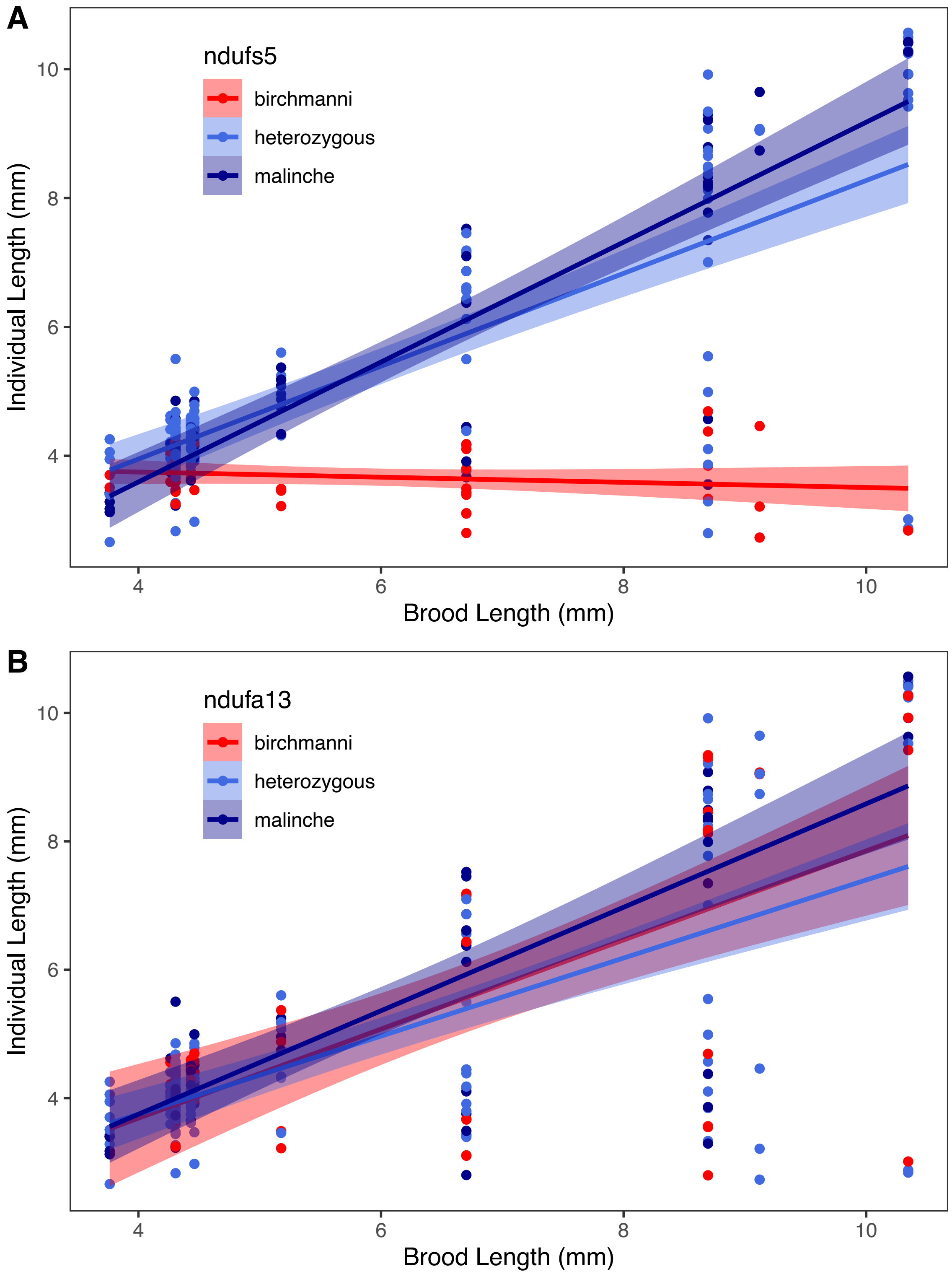

**Fig. S15**. Relationship between individual length, genotypes at (**A**) *ndufs5* and (**B**) *ndufa13*, and an estimate of the expected length for the focal brood (see Supplementary Information 1.3.5) in $F_2$ hybrid embryos. Color represents genotype, and ribbons indicate the 95% confidence intervals of the predicted slope from linear regression analysis (Table S5).

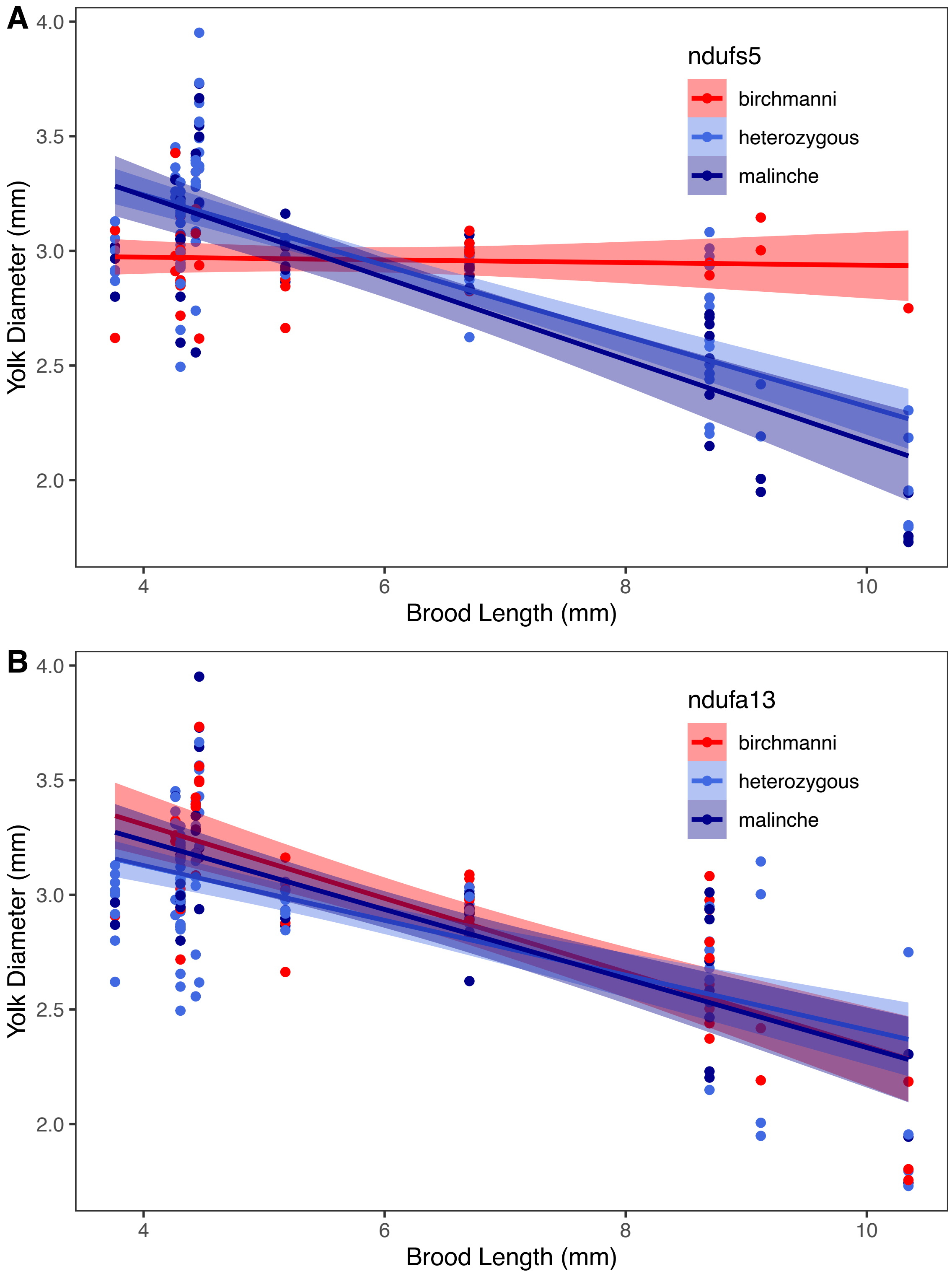

**Fig. S16**. Relationship between yolk diameter, genotype at (**A**) *ndufs5* and (**B**) *ndufa13*, and an estimate of the expected length for the focal brood (see Supplementary Information 1.3.5) in $F_2$ hybrid embryos. Color represents genotype, and ribbons indicate the 95% confidence intervals of the predicted slope from linear regression analysis (Table S9).

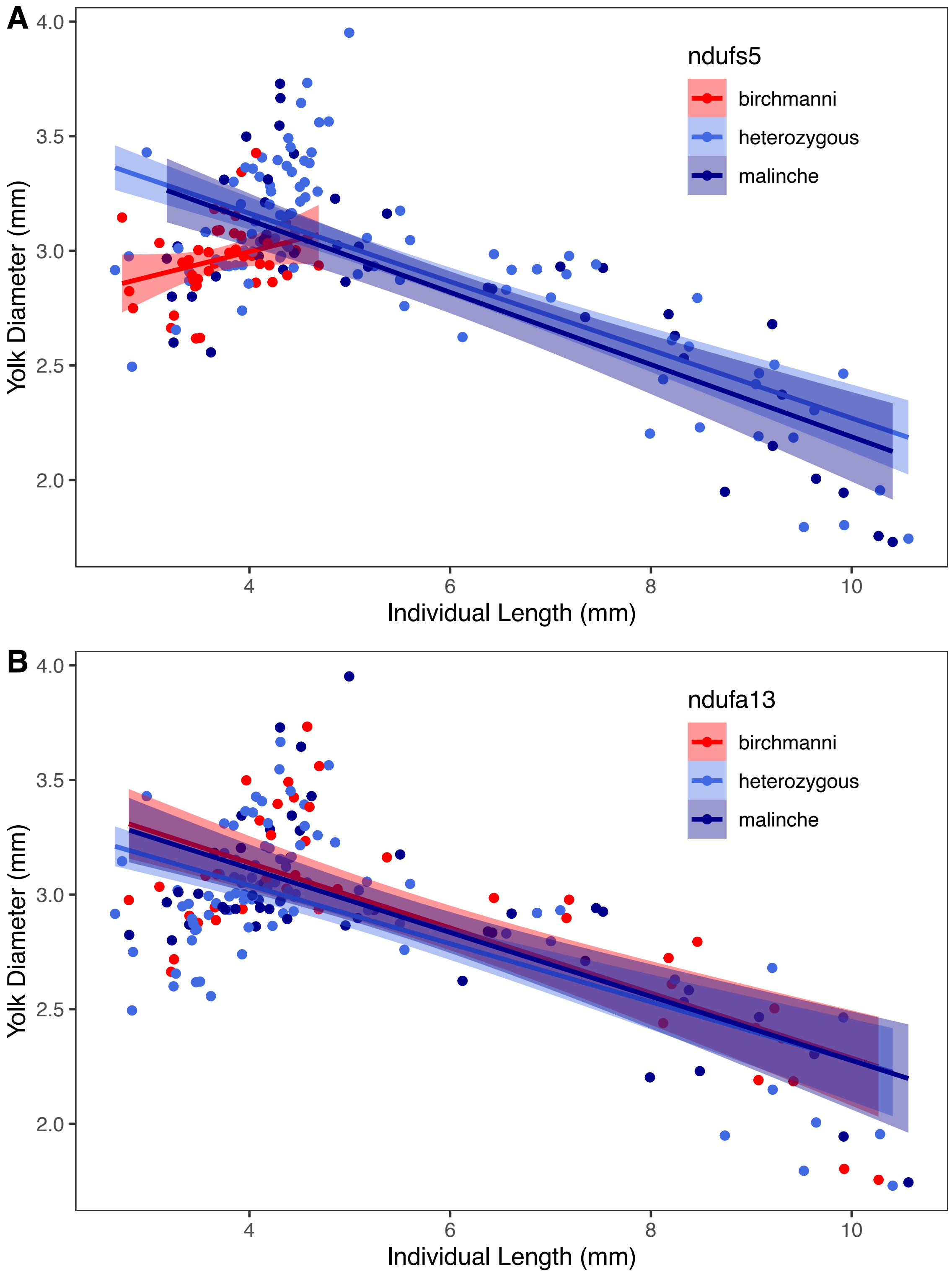

**Fig. S17**. Relationship between yolk diameter, genotype at (**A**) *ndufs5* and (**B**) *ndufa13*, and individual length (which is strongly correlated with age) in $F_2$ hybrid embryos. Color represents genotype, and ribbons indicate the 95% confidence intervals of the predicted slope from linear regression analysis (Table S9).

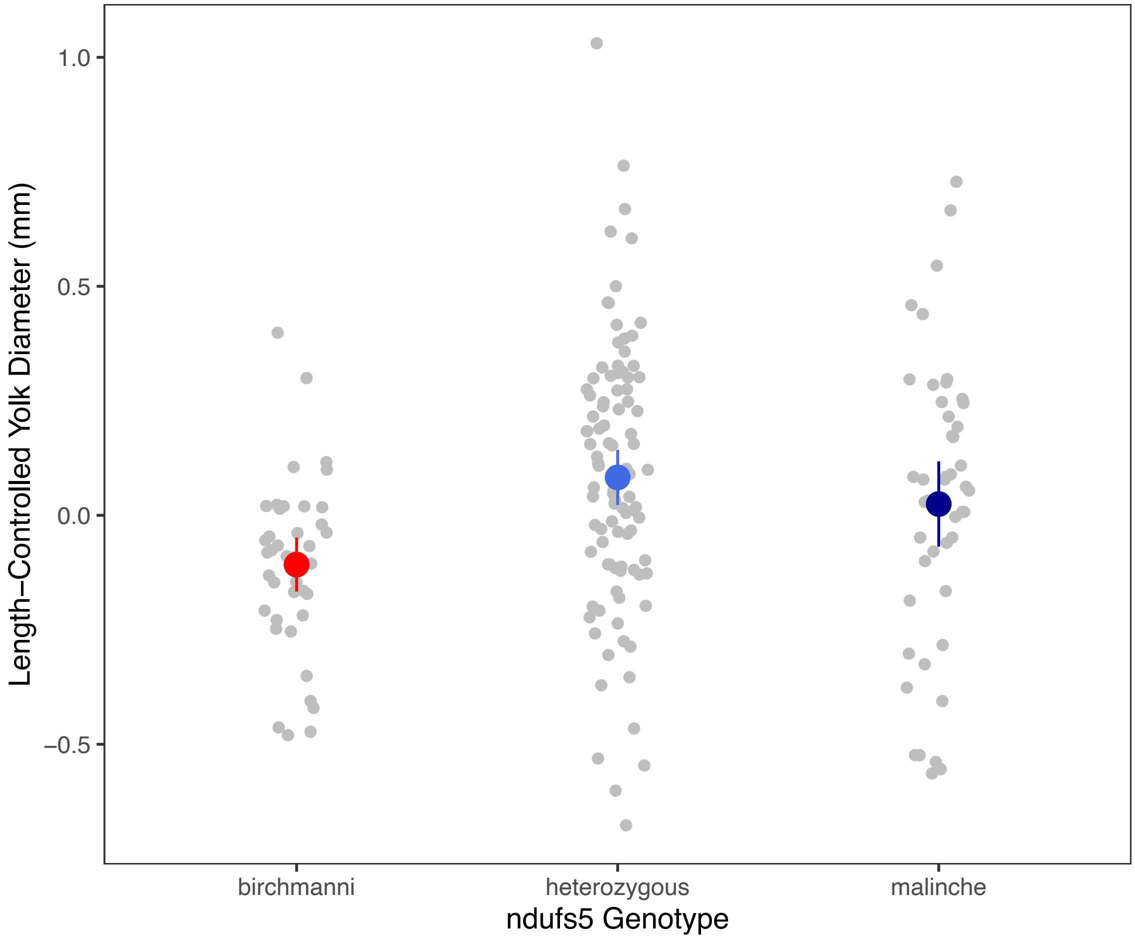

**Fig. S18**. Relationship between yolk diameter and *ndufs5* genotype in $F_2$ hybrid embryos. To control for the strong effect of length on yolk diameter, the residuals of yolk diameter after accounting for body length are plotted. Grey points represent individual measurements, colored points with vertical lines represent group mean ± 2 SE.

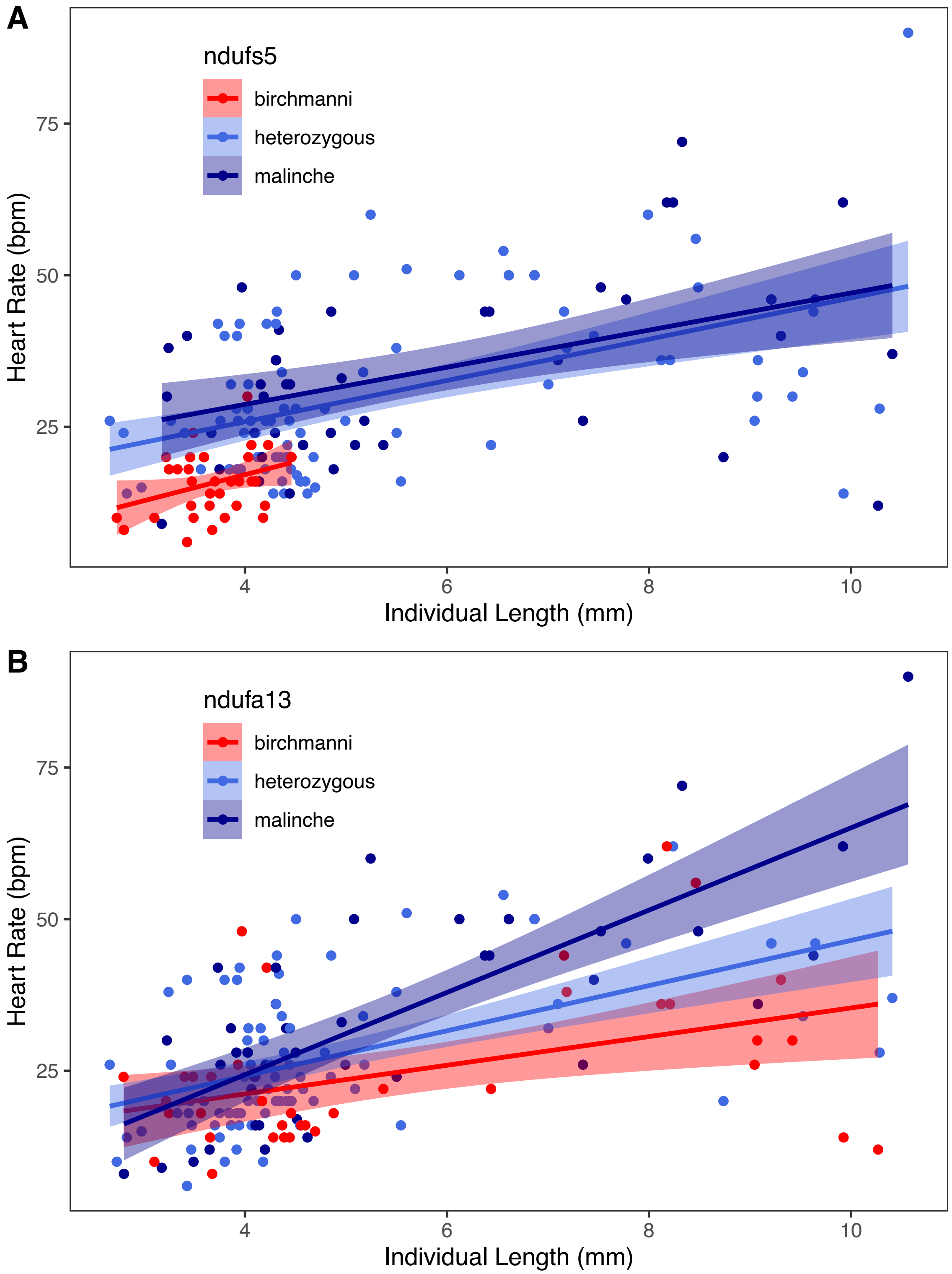

**Fig. S19**. Relationship between heart rate, genotype at (**A**) *ndufs5* and (**B**) *ndufa13*, and individual length (which is strongly correlated with age) in $F_2$ hybrid embryos. Color represents genotype, and ribbons indicate the 95% confidence intervals of the predicted slope from linear regression analysis (Table S6).

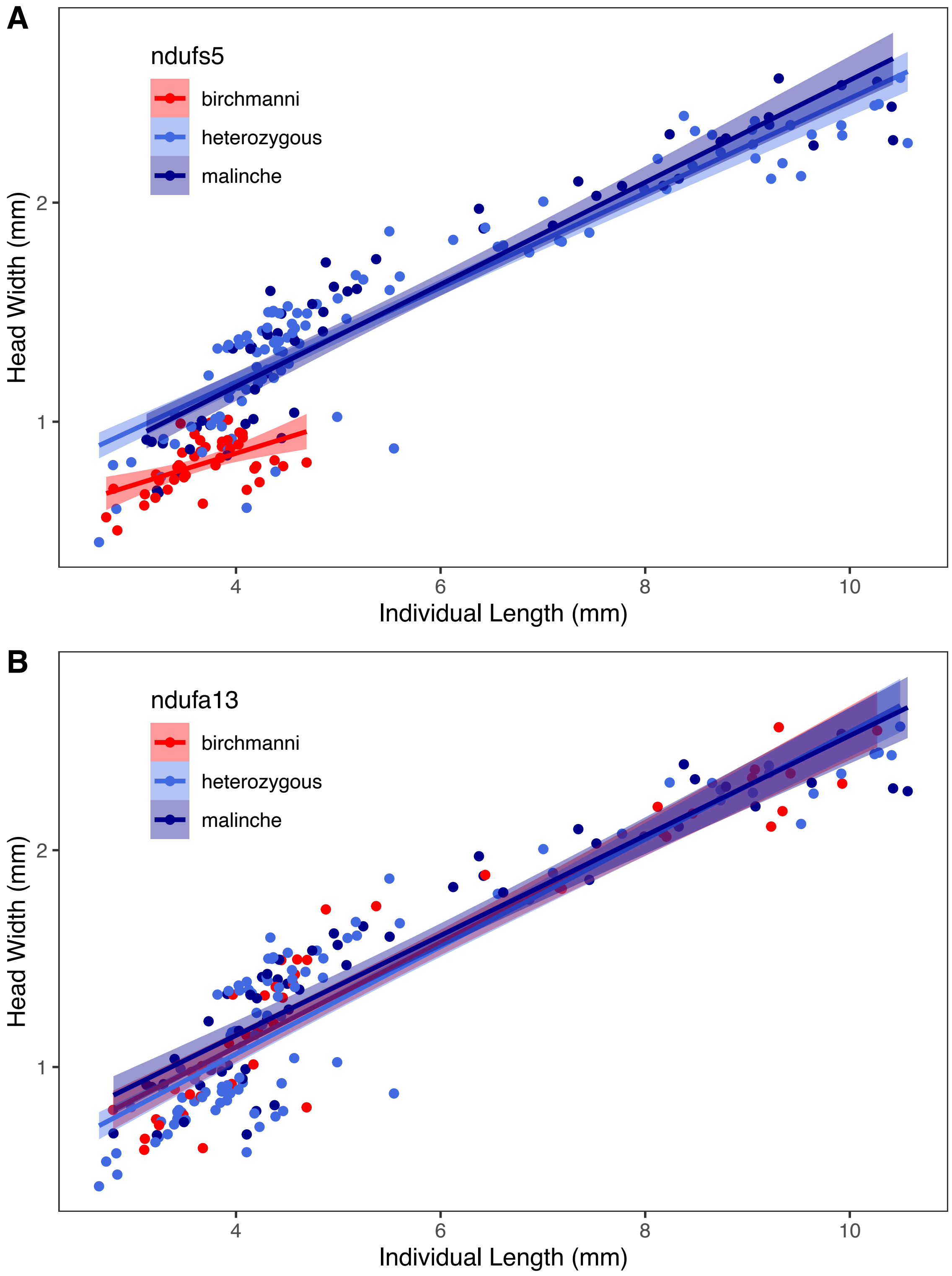

**Fig. S20**. Relationship between head width, genotype at (**A**) *ndufs5* and (**B**) *ndufa13*, and individual length (which is strongly correlated with age) in $F_2$ hybrid embryos. Color represents genotype, and ribbons indicate the 95% confidence intervals of the predicted slope from linear regression analysis (Table S7).

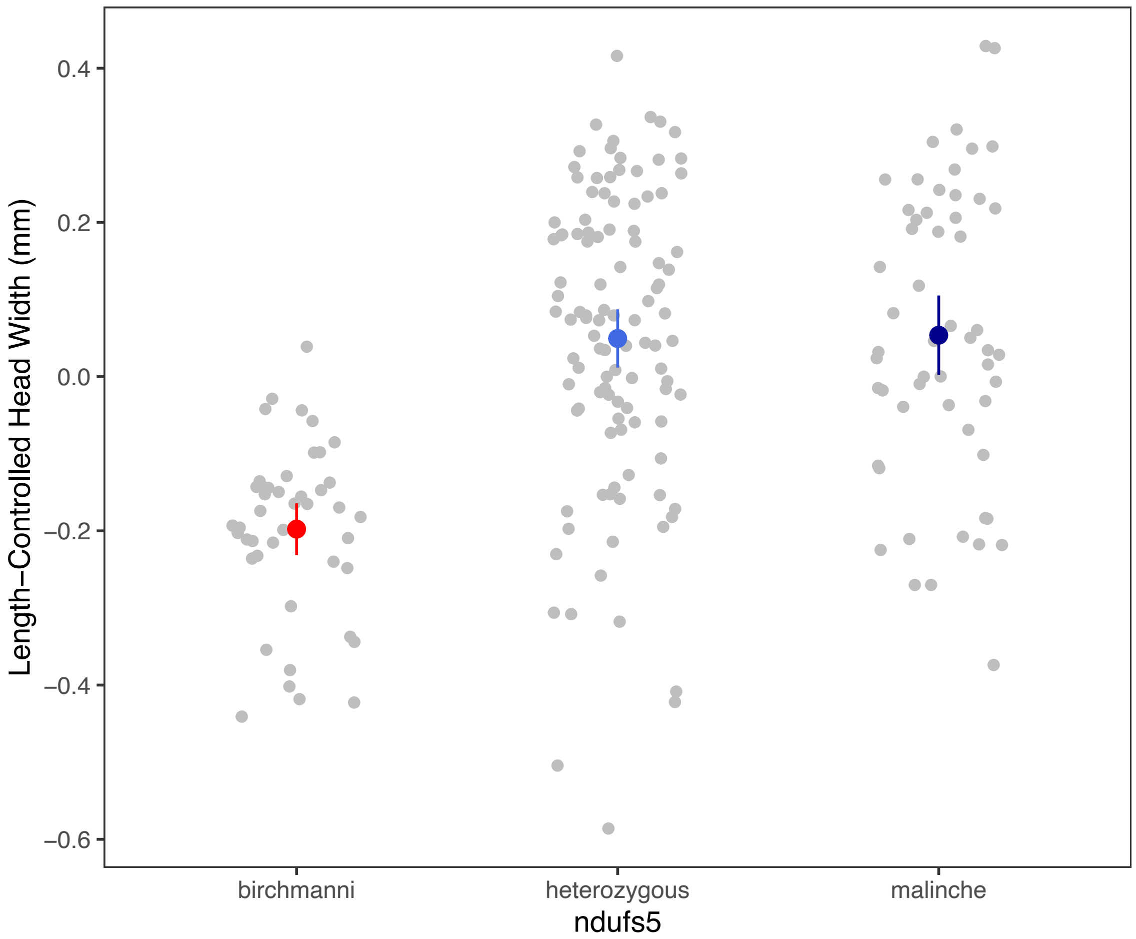

**Fig. S21**. Relationship between head width and *ndufs5* genotype in $F_2$ hybrid embryos. To control for the strong effect of developmental stage on head width, residuals of head width after accounting for body length are plotted. Grey points represent individual measurements, colored points with vertical lines represent group mean ± 2 SE.

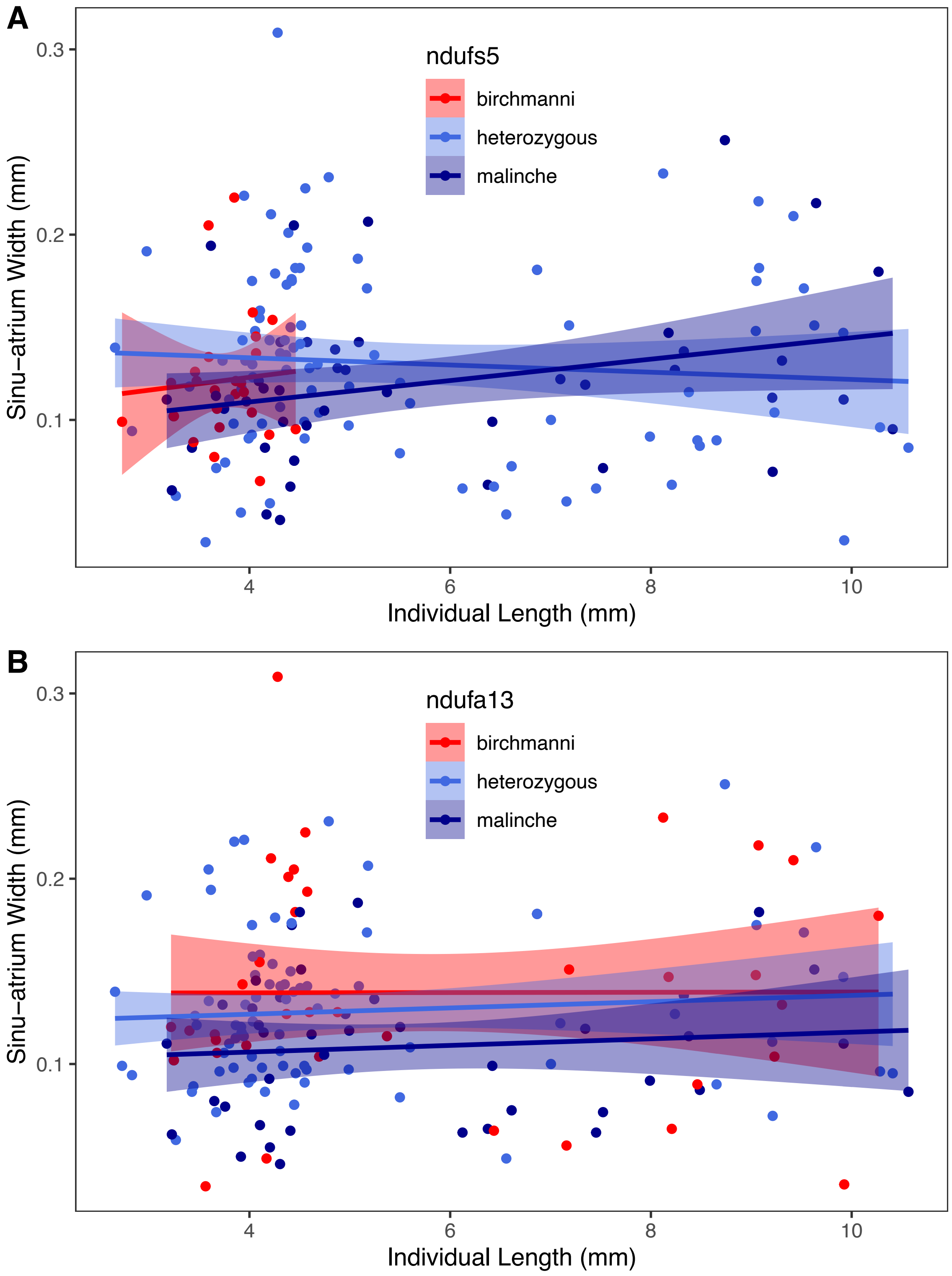

**Fig. S22**. Relationship between sinu-atrium width, genotype at (**A**) *ndufs5* and (**B**) *ndufa13*, and individual length (which is strongly correlated with age) in $F_2$ hybrid embryos. Color represents genotype, and ribbons indicate the 95% confidence intervals of the predicted slope from linear regression analysis (Table S8).

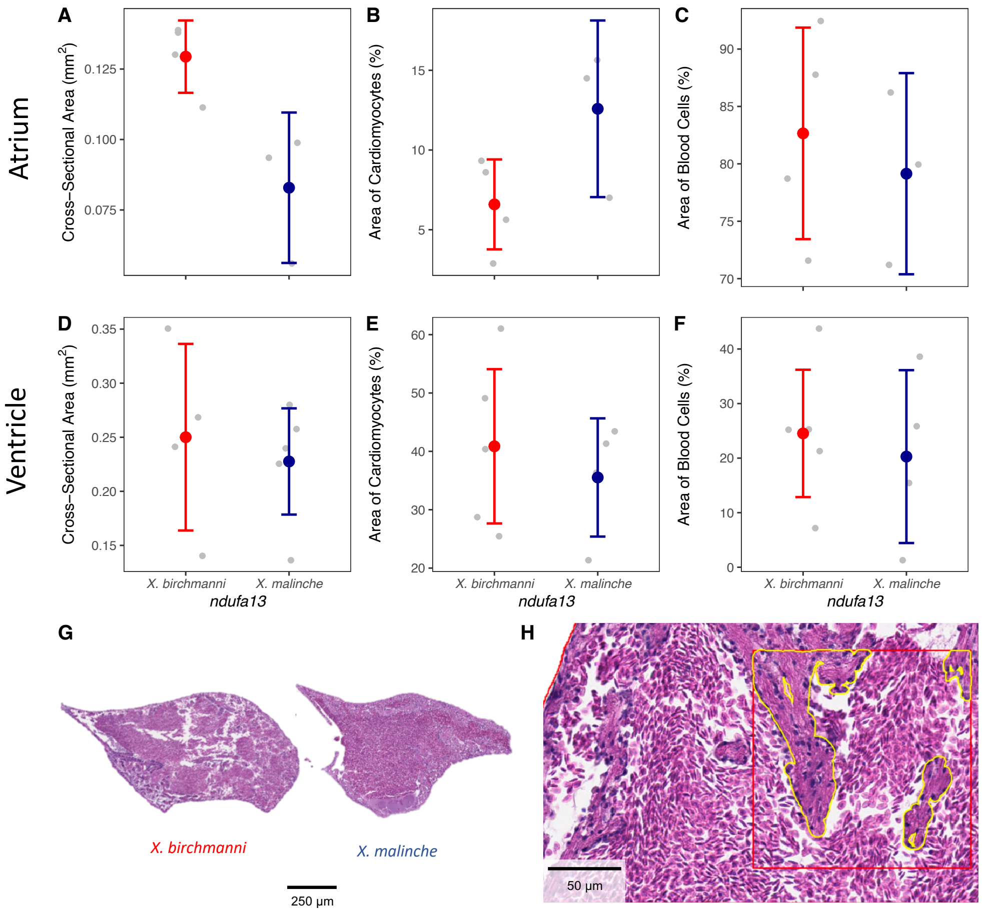

**Fig. S23**. Heart chamber morphology in juvenile $F_2$ hybrids as a function of *ndufa13* genotype. Individuals that were homozygous *X. birchmanni* at *ndufa13* (the incompatible genotype) and homozygous *X. malinche* at *ndufa13* (the compatible genotype) were raised in common laboratory conditions and sampling occurred at approximately 5 months of age. Measurements were taken from the sagittal section of largest cross-sectional area for the atrium (A-C) and ventricle (D-F), and area of occupancy for each cell class was calculated from the average of three quadrats (see Supplementary Information 1.3.3). (G) Representative images of atria from incompatible (*X. birchmanni)* and compatible (*X. malinche*) individuals. Images are from slides with maximum cross-sectional area for each individual and chamber. (H) Example of histology analysis process in a representative atrium. Red square indicates randomly placed quadrat in which occupancy was calculated, and yellow borders represent areas which were manually annotated as cardiomyocytes. Note that the epicardium is visible at top left.

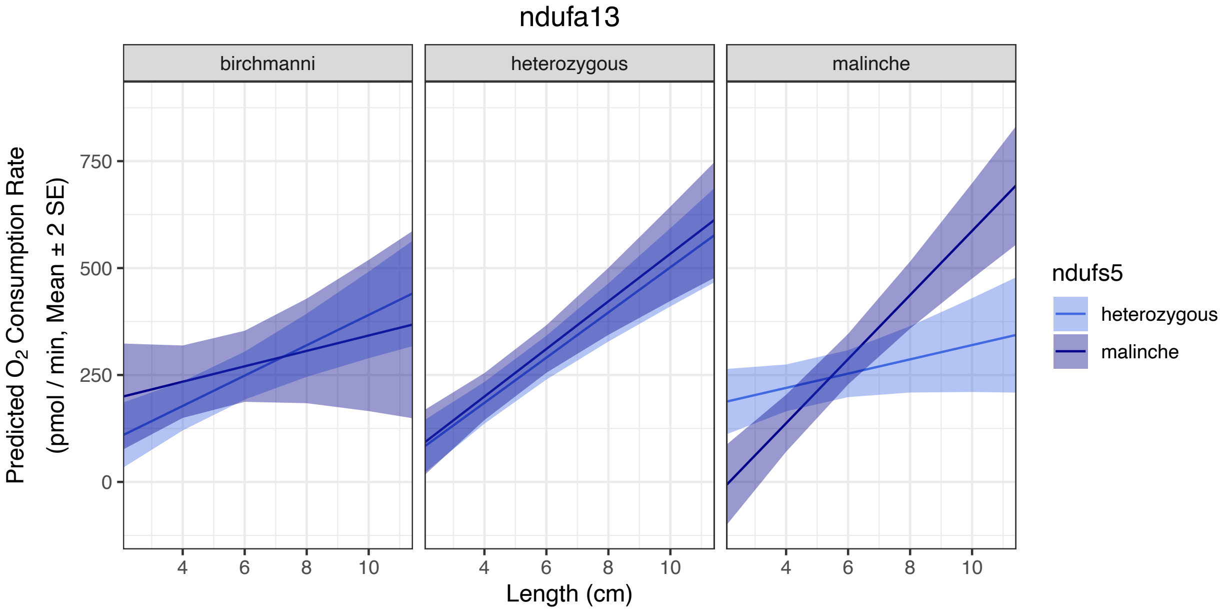

**Fig. S24**. Visualization of the statistically detected three-way interaction between body length, *ndufs5* genotype, and *ndufa13* genotype on respiration rate in $F_2$ hybrid embryos. Displayed are predicted respiration rates as a function of length (x-axis), *ndufs5* genotype (color), and *ndufa13* genotype (panel). Ribbons indicate the 95% confidence interval for predicted slope from linear regression. The differing slopes between each color and panel indicates that respiration cannot be predicted without knowing all three variables, which could hint at a complex interaction between *ndufs5* and *ndufa13* in determining the change in respiration throughout development (Table S10).

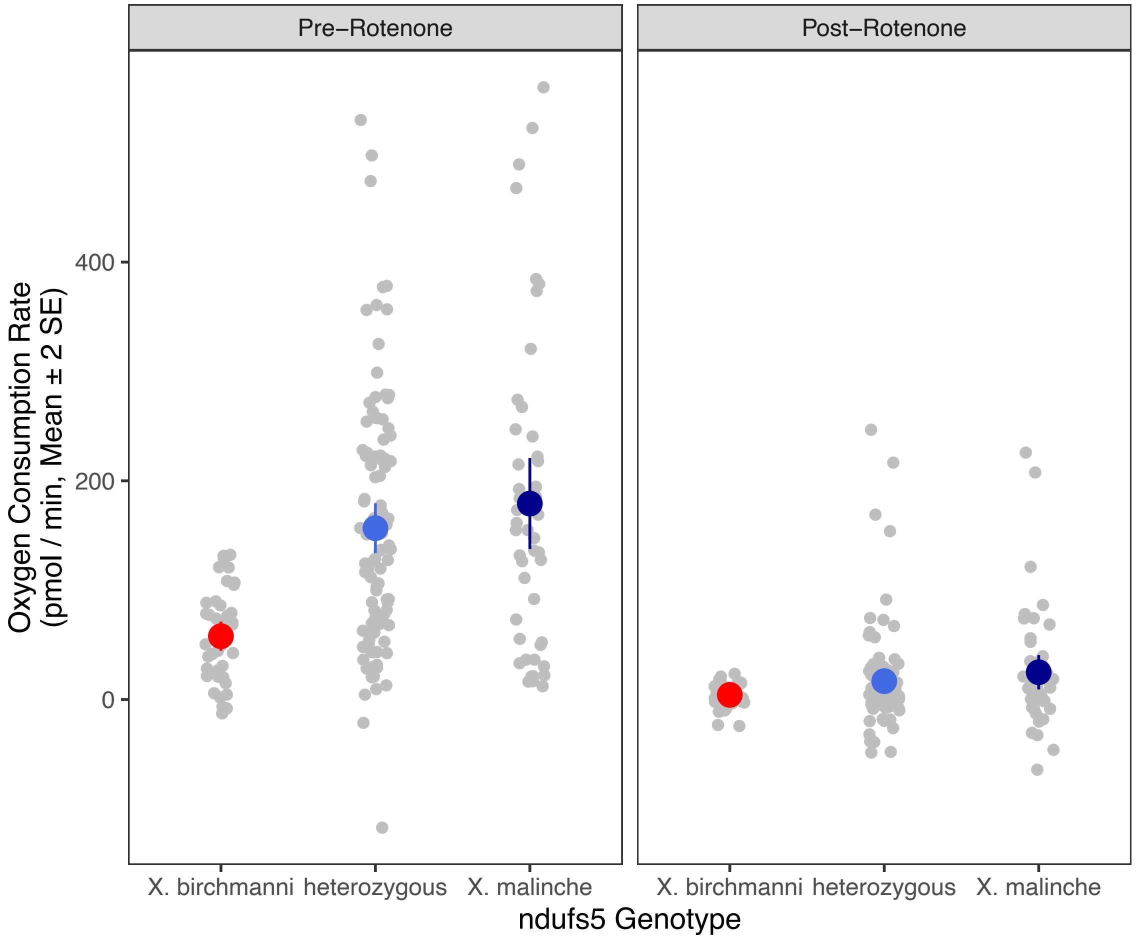

**Fig. S25**. Raw oxygen consumption data as a function of *ndufs5* genotype, before (left) and after (right) the addition of 5 µM rotenone to respirometry media. Grey points represent individual measurements, colored points with vertical lines represent the mean ± 2 SE.

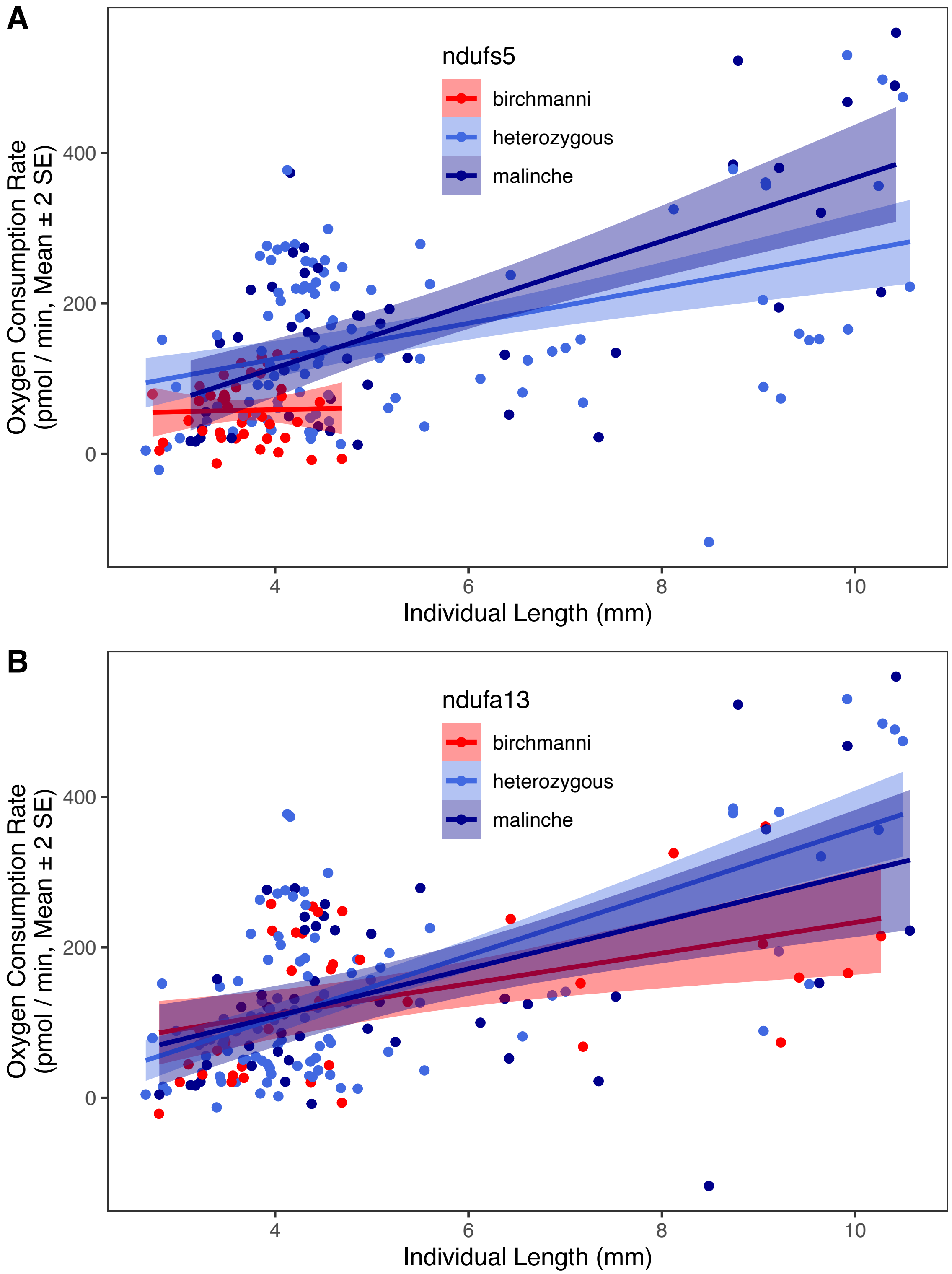

**Fig. S26**. Relationship between oxygen consumption rate, genotype at (**A**) *ndufs5* and (**B**) *ndufa13*, and individual length (which is strongly correlated with age) in $F_2$ hybrid embryos. Color represents genotype, and ribbons indicate the 95% confidence intervals of the predicted slope from linear regression analysis (Table S10).

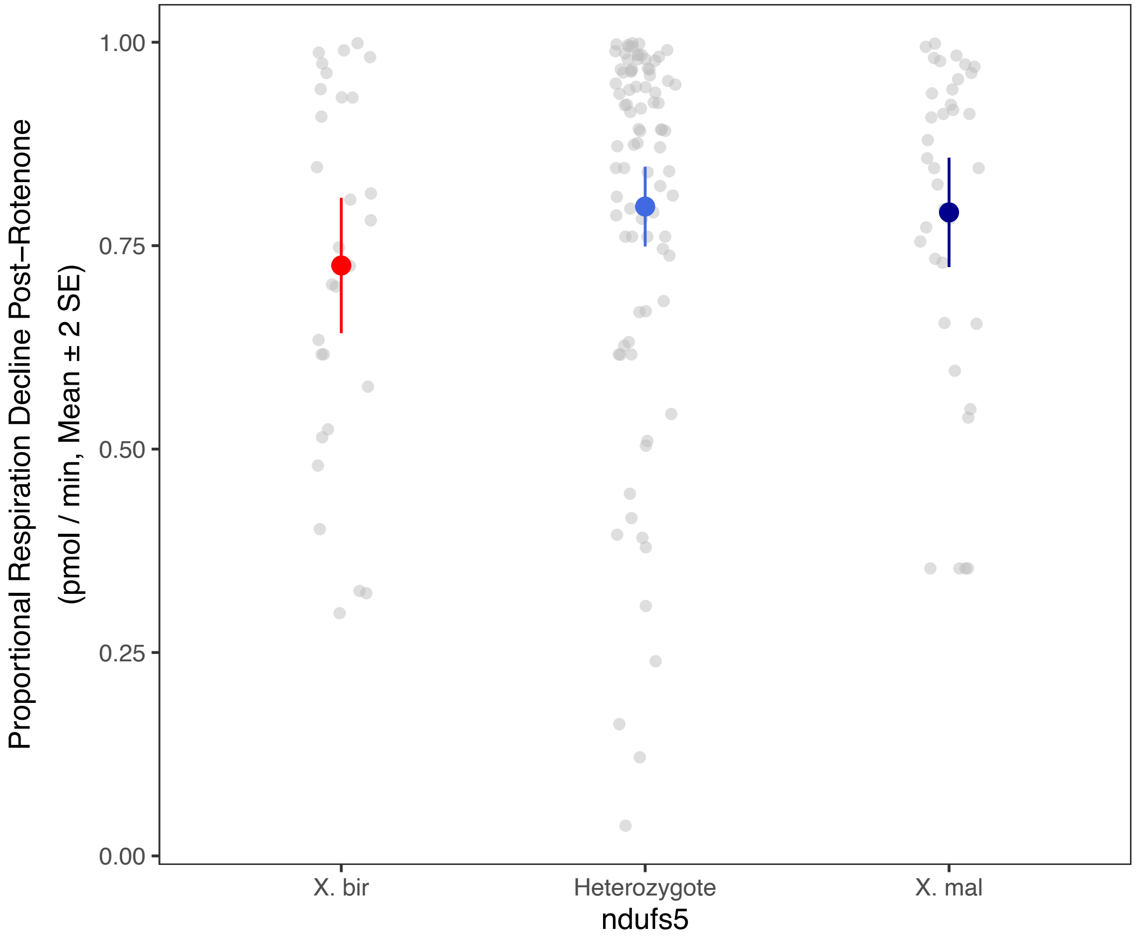

**Fig. S27**. Proportion of respiration decline between pre- and post-rotenone oxygen consumption trials as a function of *ndufs5* genotype in $F_2$ hybrid embryos. Grey points represent individual measurements, colored points with vertical lines represent group mean ± 2 SE.

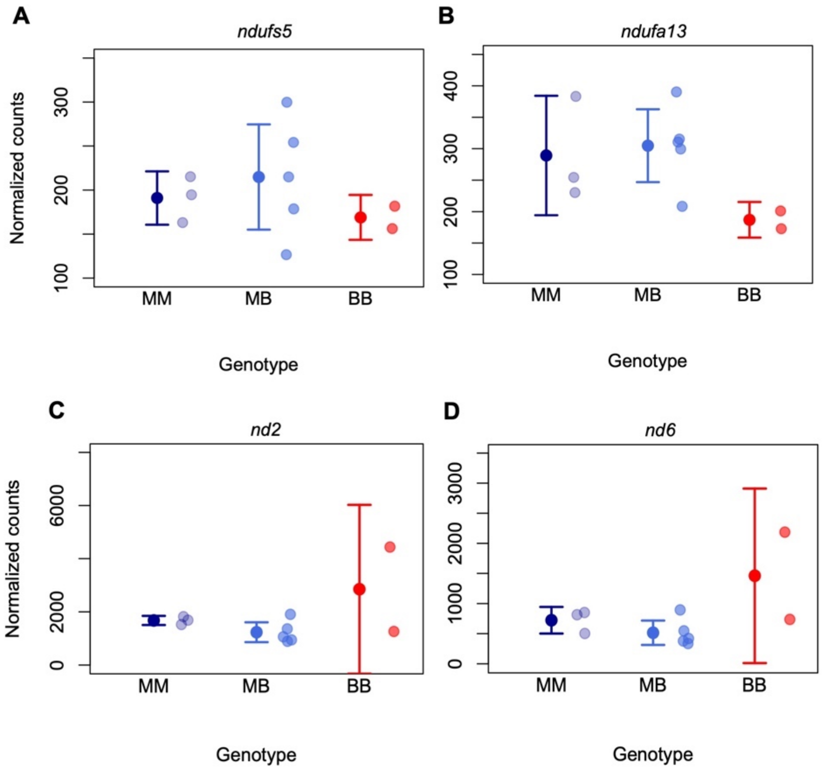

**Fig. S28.** Expression levels of genes involved in the mitonuclear hybrid incompatibility across groups based on RNAseq data collected from whole liver tissue. Plotted are normalized counts for each gene inferred using kallisto. Semi-transparent points show individual biological replicates of *X. malinche* (MM), $F_1$ hybrids (MB), and *X. birchmanni* (BB). Mean and ± 2 standard errors of the mean are shown with dots and whiskers for each genotype.

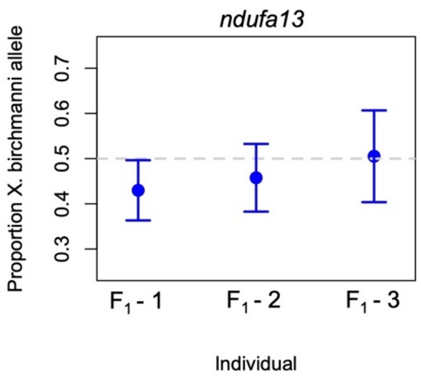

**Fig. S29.** Allele-specific expression from read counts for *X. birchmanni* and *X. malinche* *ndufa13* alleles in $F_1$ hybrids. The y-axis shows the proportion of reads in $F_1$ hybrids supporting the *X. birchmanni* allele. Differences are not significant from 50-50 expectations as evaluated by inverted beta-binomial test. Reads displayed here were mapped to the *X. birchmanni* transcriptome, but mapping to the *X. malinche* reference transcriptome did not qualitatively change results. Observed ratio is shown by the points and the error bars show two standard errors expected from the number of reads and binomial sampling of the two alleles in $F_1$ hybrids.

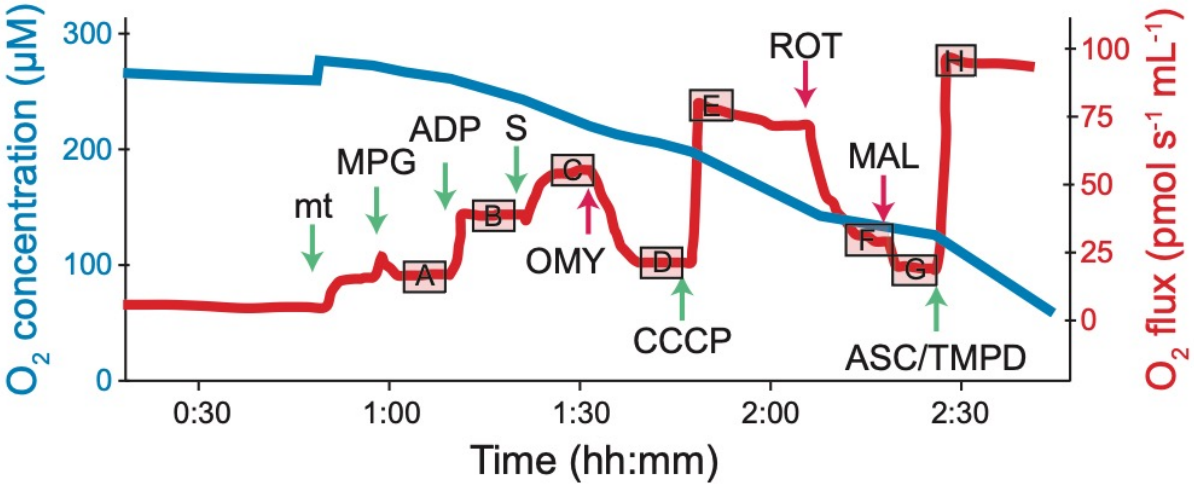

**Fig. S30.** A typical O2K run illustrating the high-resolution respiratory protocol used (data shown here are from a *X. birchmanni* male). $O_2$ concentration is shown in blue, $O_2$ flux (i.e., respiration) is shown in red. Arrows indicate the addition of substrates/uncouplers (green arrows) or inhibitors (red arrows) following Table S18. Letters in boxes (A-H) indicate the eight respiratory states examined (Table S18) and the boxes indicate the general range of data used to calculate the flux values in each state.

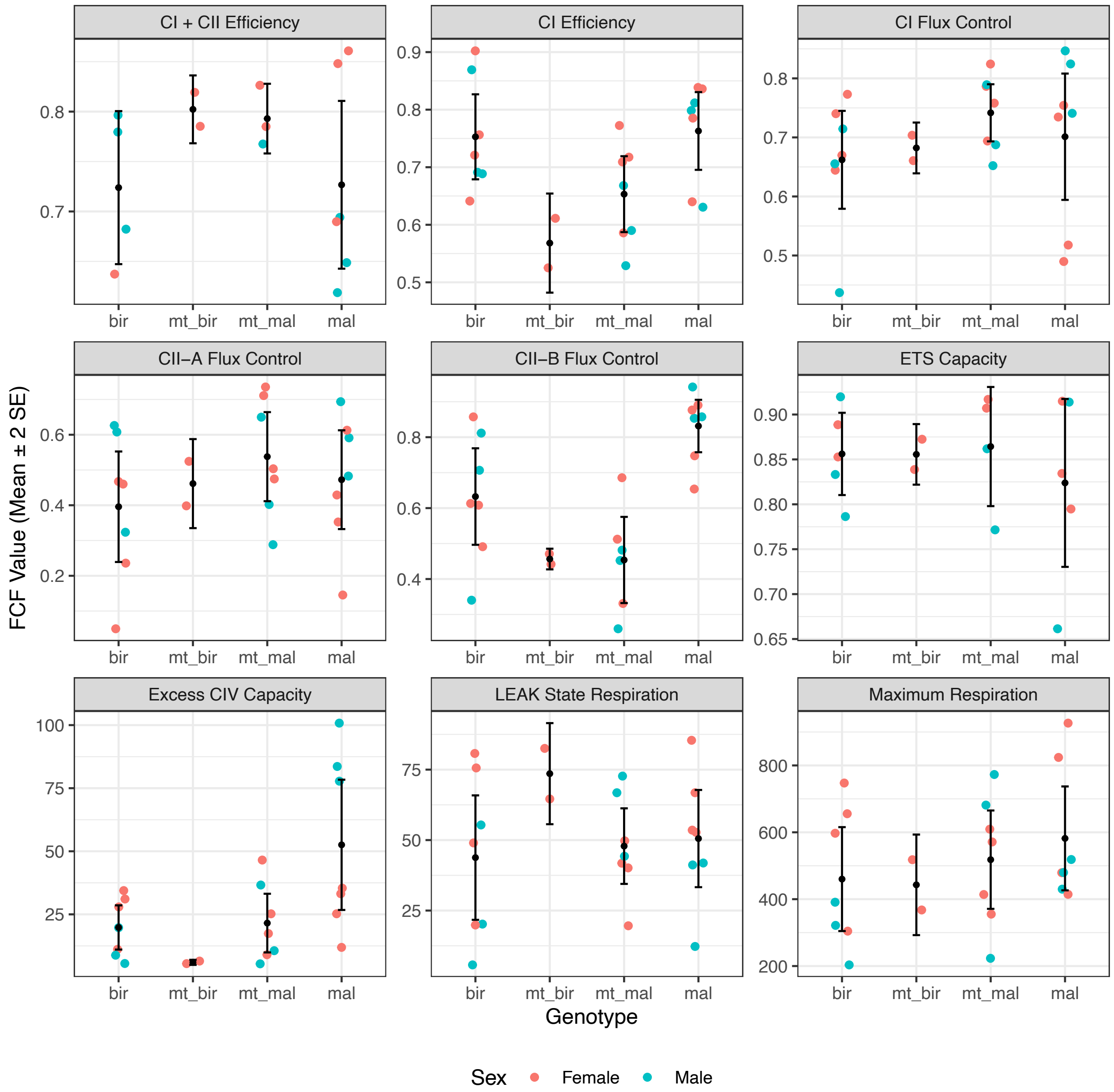

**Fig. S31.** Results of Oroboros O2K respirometry. Panels represent flux control factors (FCF) measured across pure *X. birchmanni* (bir) and *X. malinche* (mal) individuals, as well as hybrids heterozygous at *ndufs5* and *ndufa13* with either *X. birchmanni* (mt_bir) or *X. malinche* (mt_mal) mitochondrial ancestry. LEAK respiration and maximum respiration are measured in units of pmol $O_2$ $s^{-1}$ $mL^{-1}$ $mg^{-1}$, and all other FCF are unitless ratios; see Table S12 for descriptions of FCF calculations. Maximum sample sizes were n=7 for *X. birchmanni*, *X. malinche*, and hybrids with *X. malinche* mitochondria, and n=2 for hybrids with *X. birchmanni* mitochondria. Lower sample sizes for some FCF measurements indicate cases where samples were discarded due to measurement quality issues (see *Supplementary Information 1.3.8 Respirometry for OXPHOS Complex activity in parents and hybrids*). Black points with error bars show the mean and two standard errors, and colored points show data for each individual colored by sex.

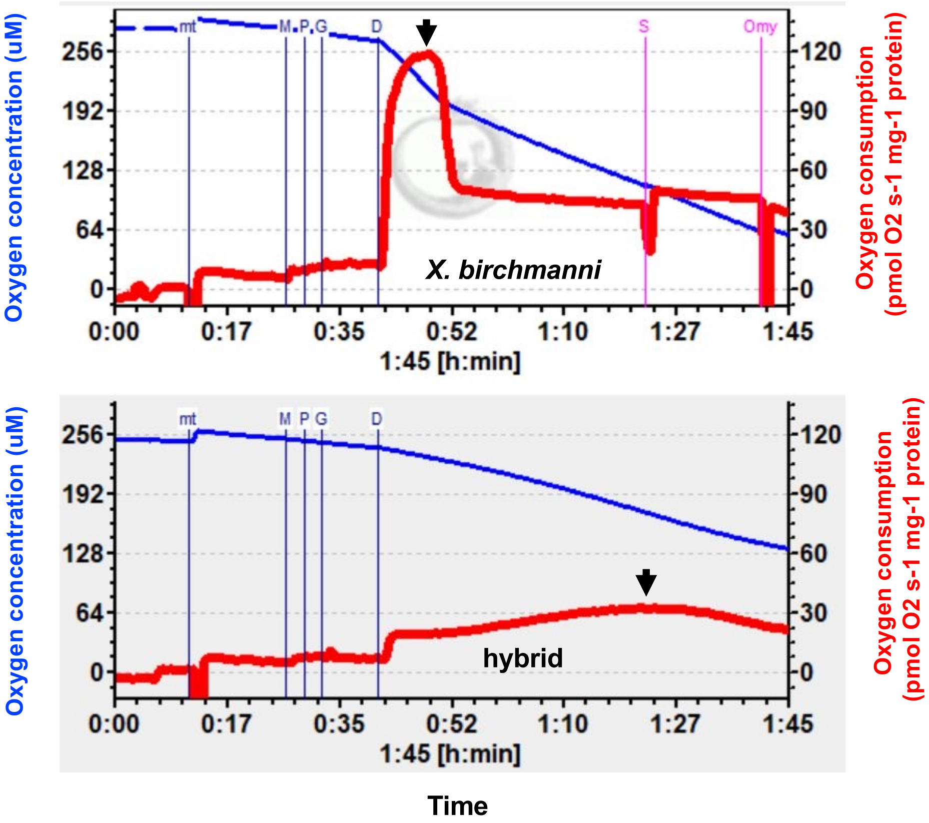

**Fig. S32.** A comparison of typical O2K runs for a *X. birchmanni female* (top) and $F_1$-hybrid male with *X. malinche* mitochondria (bottom), illustrating the difference in time to peak respiration after ADP addition. $O_2$ concentration is shown in blue, $O_2$ flux (i.e., respiration) is shown in red. Vertical lines indicate substrates and inhibitors added to each chamber after stabilization of flux: mt (addition of mitochondrial sample, standardized to 0.15 mg), M (malate), P (pyruvate), G (glutamate), D (ADP), S (succinate), and Omy (oligomycin). Black arrows indicate the time of peak respiration after ADP (D) in the *X. birchmanni* (~5 minutes after addition) and hybrid (~40 minutes after addition) mitochondria. Differences in time to post-ADP peak were significant between genotypes. Visualization is adapted from DatLab 7.3.0.3 (Oroboros, Innsbruck, Austria).

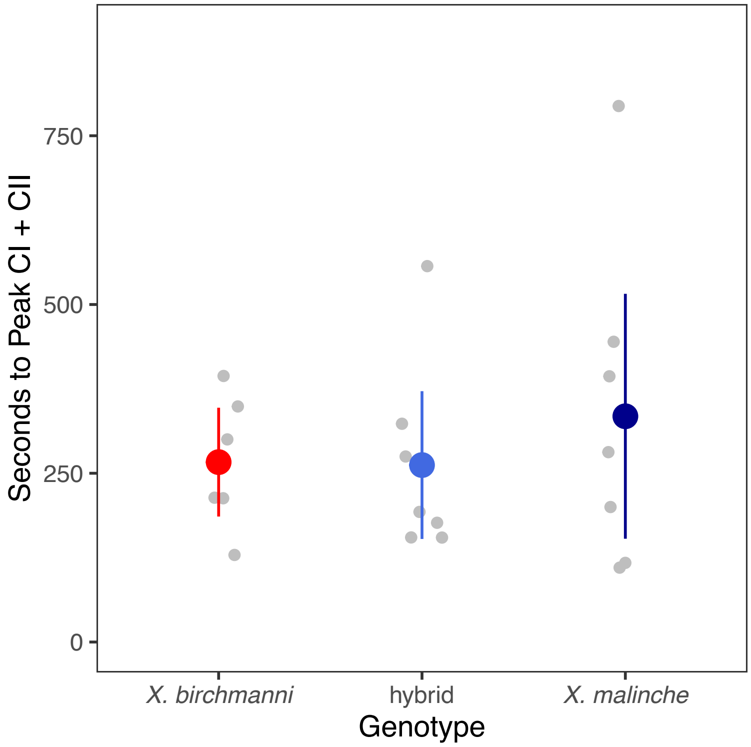

**Fig. S33.** Time to reach the peak in Complex I- and Complex II-driven respiration (State C, Table S18) after the addition of succinate did not differ across genotypes. Complex II-driven respiration begins with the addition of succinate, which is the electron-donating substrate of Complex II.

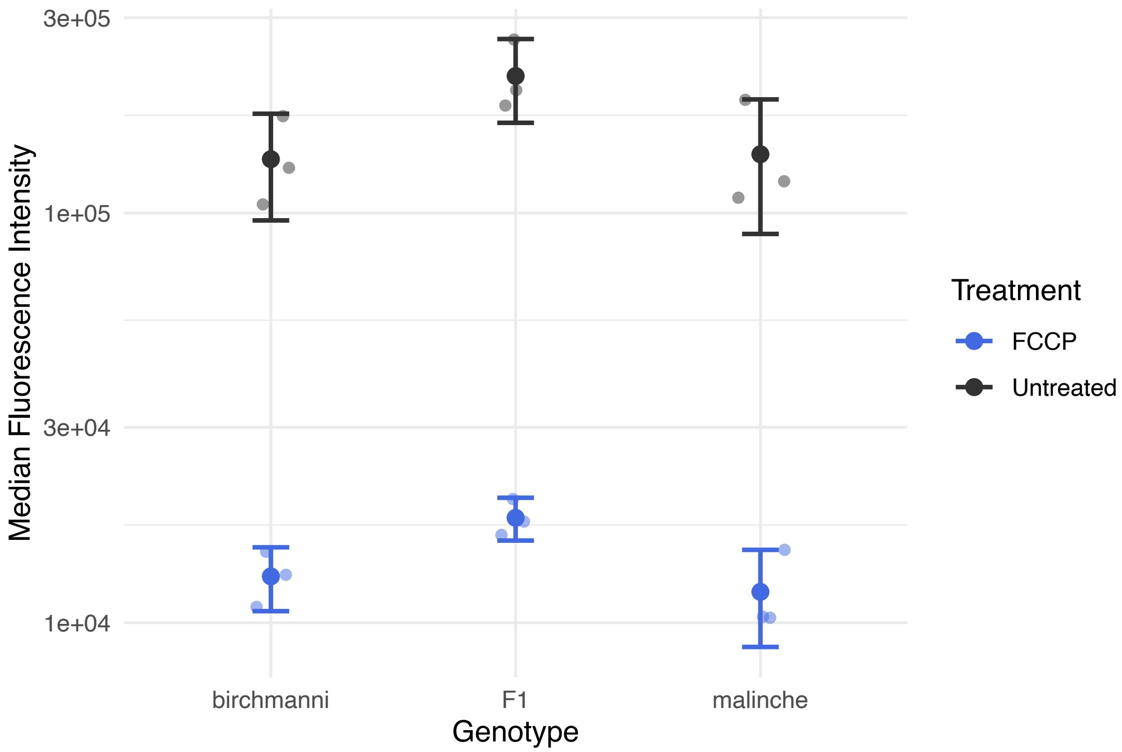

**Fig. S34.** Flow cytometry measurements of mitochondrial membrane potential in caudal fin fibroblast cells collected from *X. malinche*, *X. birchmanni*, and $F_1$ individuals. Note that the y-axis is log-scaled. Black points and bars represent mean ± 2 SE TMRE fluorescence intensity, blue points and bars represent TMRE fluorescence after addition of the uncoupling agent FCCP, with lighter points of each color showing raw data. No significant differences were observed between genotypes in untreated fluorescence (ANOVA $F = 3.505$, $P = 0.10$) or proportional change with FCCP addition (ANOVA $F = 0.196$, $P = 0.827$). Note that for statistical analysis untreated fluorescence data was log-scaled for normality.

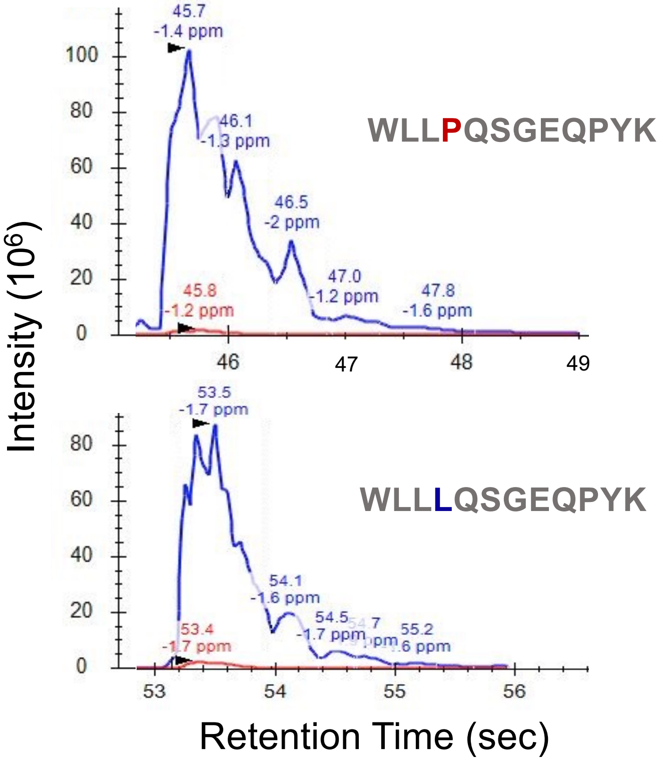

**Fig. S35.** Representative Skyline traces of the *X. malinche* and *X. birchmanni* versions of the *ndufs5* peptide from parallel reaction monitoring. Blue curves represent the spectral intensity from heavy-labeled peptides, red curves represent endogenous peptides. First and second lines of text above spectral curves represents the retention time and mass error relative to theoretical m/z, respectively, for the identified fragment. Inset shows the sequence of the *X. birchmanni* and *X. malinche* peptides being detected.

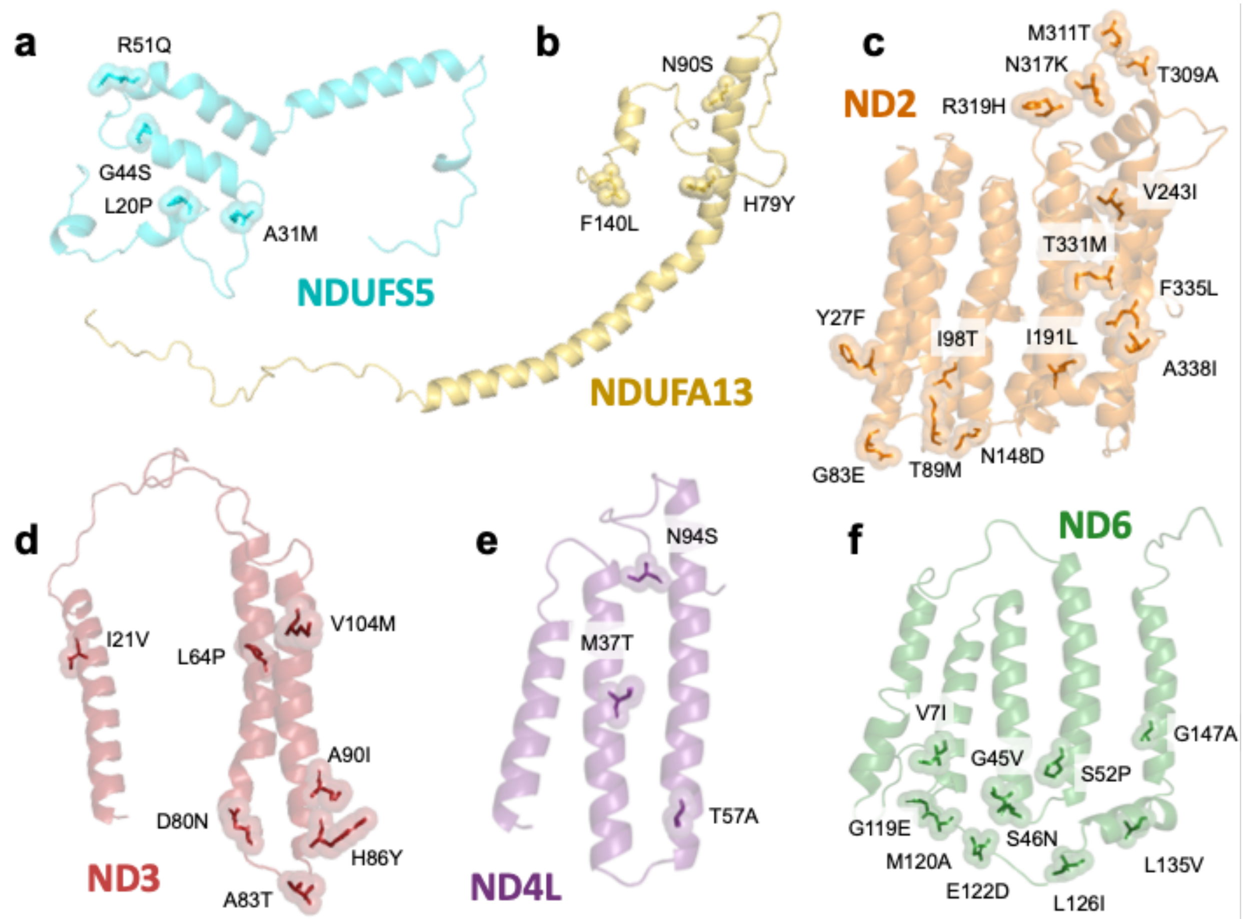

**Fig. S36.** RaptorX structure predictions for *Xiphophorus ndufs5* (a), *ndufa13* (b), *nd2* (c), *nd3* (d), *nd4l* (e) and *nd6* (f). Side chains of residues with substitutions between *X. birchmanni* and *X. malinche* are displayed as sticks surrounded by translucent spheres. Substitutions are listed in black text with single letter amino acid code for the *X. malinche* allele and *X. birchmanni* allele separated by the residue number.

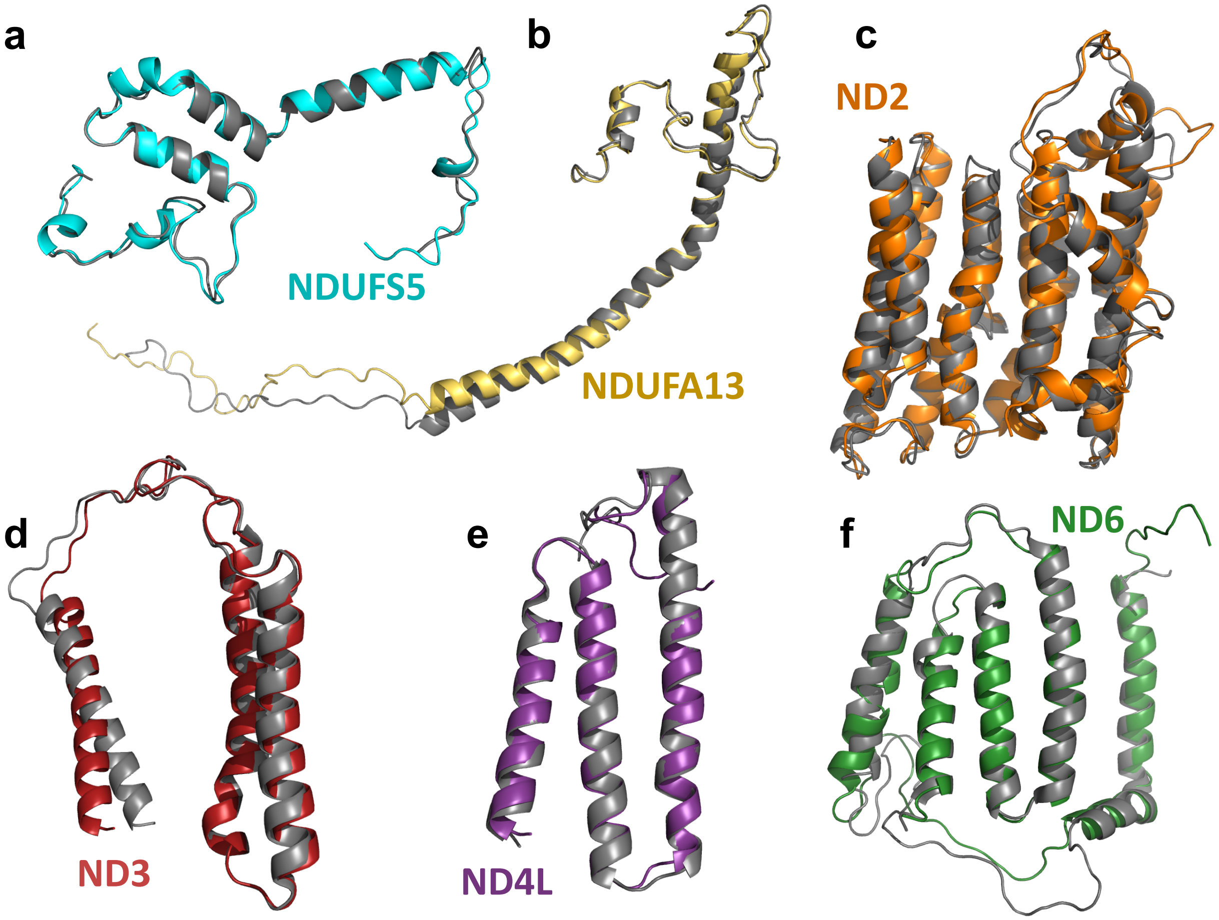

**Fig. S37.** Alignment of *X. birchmanni* RaptorX models and their mammalian template structures. Mammalian templates are displayed in gray, *X. birchmanni* models in color. Alignments were constructed in PyMol using the GUI "align" command. Model-template pairs include (a) *X. birchmanni* NDUFS5 and chain e of PDB 6G2J (mouse Complex I in the active state), (b) *X. birchmanni* NDUFA13 and chain Z of PDB 6G72 (mouse Complex I in the deactive state), (c) *X. birchmanni* ND2 and chain N of PDB 5LDW (cow Complex I, class I), (d) *X. birchmanni* ND3 and chain A of PDB 5LNK (entire sheep Complex I), (e) *X. birchmanni* ND4L and chain k of PDB 5XTC (human Complex I transmembrane arm), (f) *X. birchmanni* ND6 and chain J of PDB 5LNK (entire sheep Complex I).

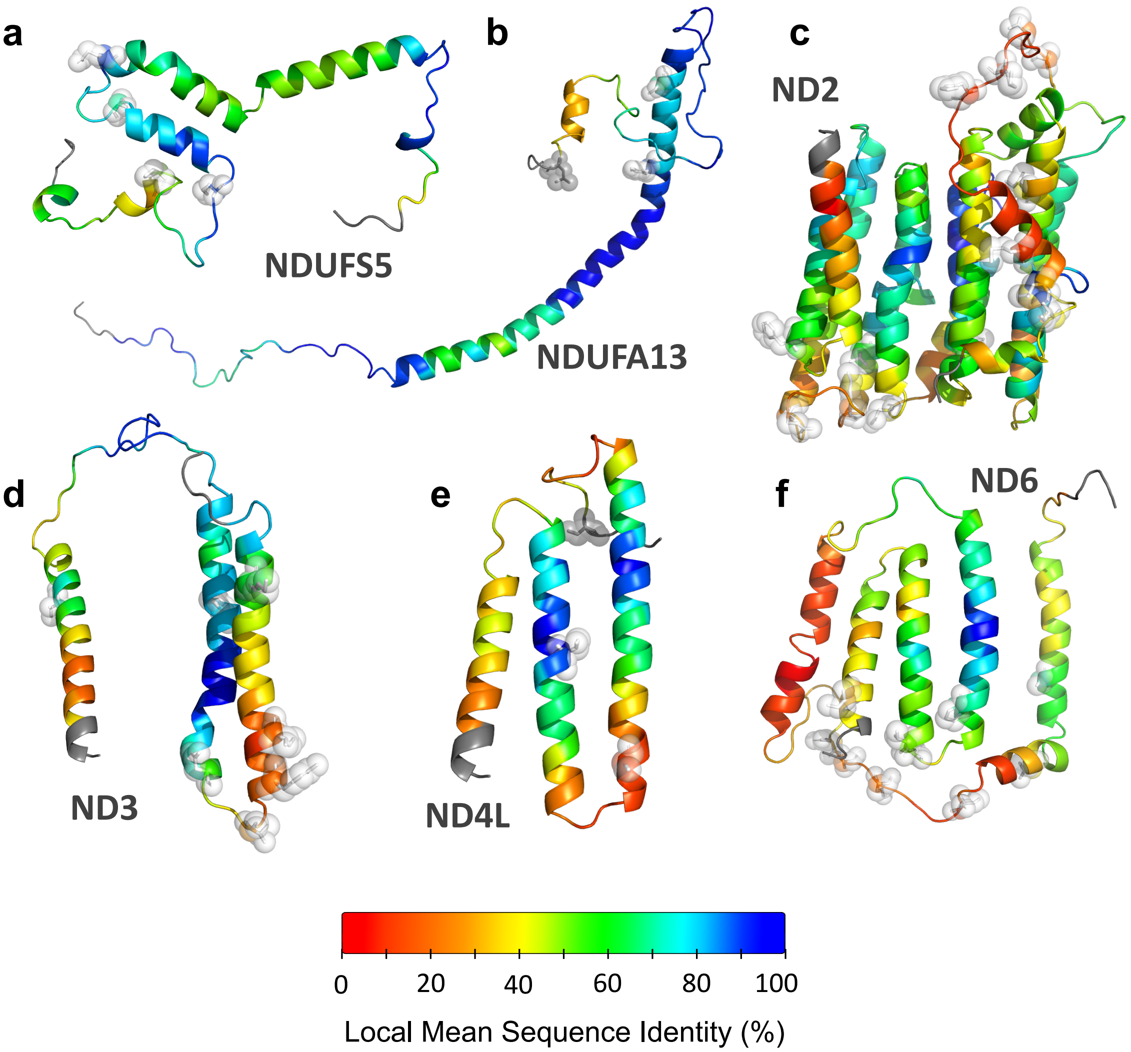

**Fig. S38.** Conservation of amino acid sequences between *X. birchmanni* Complex I genes and the mammalian models chosen by RaptorX. Color at each residue represents the mean sequence identity between the *X. birchmanni* and mammalian sequence, taken in an 11-residue window centered on the focal residue. The N- and C-terminal residues for which this metric is undefined are represented in gray. The locations of substitutions between *X. birchmanni* and *X. malinche* are highlighted by white spheres and sticks.

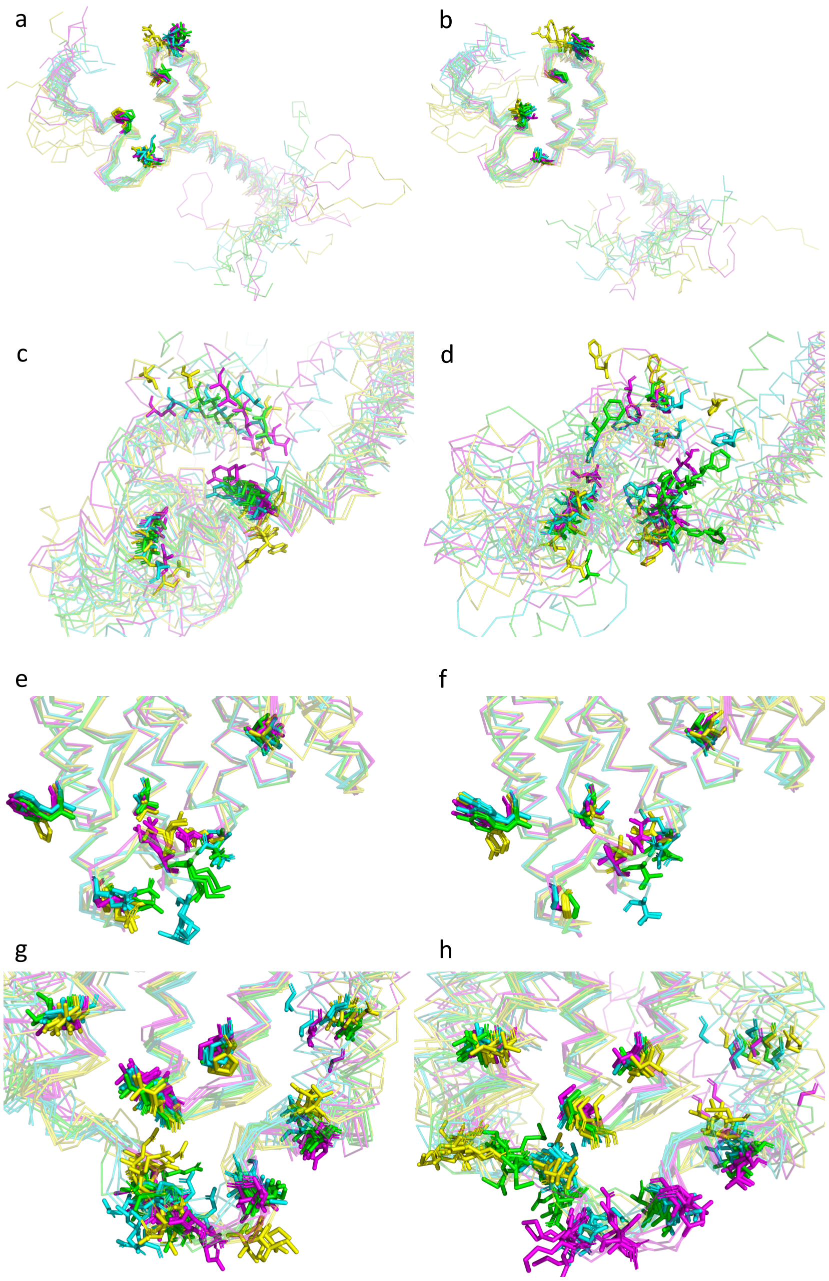

**Fig. S39.** MODELLER structure predictions for the *X. birchmanni* (a, c, e, g) and *X. malinche* (b, d, f, h) sequences for *ndufs5* (a-b), *ndufa13* (c-d), *nd2* (e-f), and *nd6* (g-h). Five alternative models were generated for each of four mammalian template structures, mouse (Protein Data Bank ID 6G2J, cyan), sheep (5LNK, magenta), cow (5LDW, yellow), and human (5XTC, green). Side chains of residues varying between species are displayed as sticks, and the main chain backbone is displayed as translucent ribbons. To make clear the difference in placement of substitutions between templates and replicate, c-h show only the region of the protein in proximity to *ndufs5*.

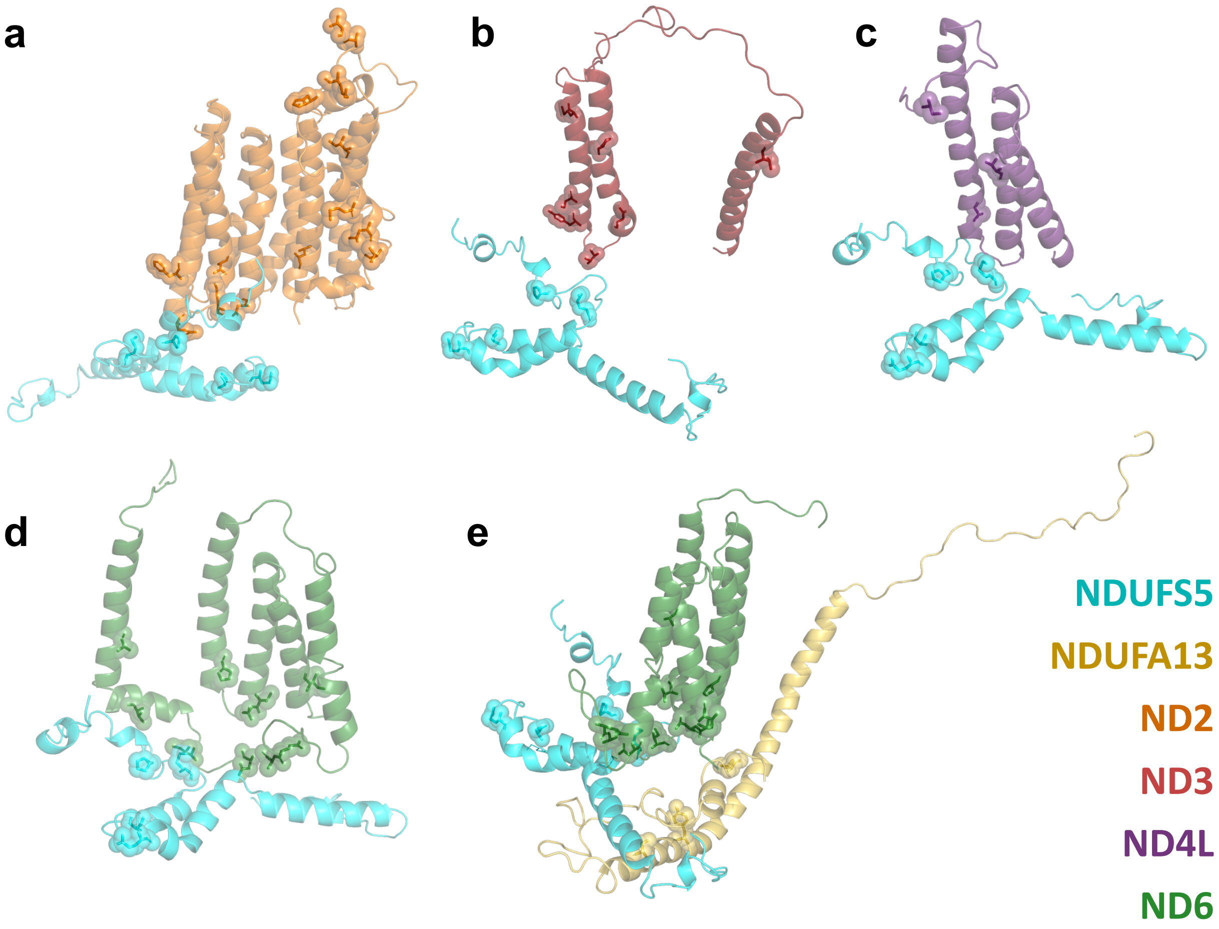

**Fig. S40.** Pairwise views of contacts between nuclear and mitochondrial subunits of respiratory Complex I. *ndufs5* is shown in cyan, in its predicted spatial position relative to (a) *nd2* (b), *nd3* (c), *nd4l* (d) *nd6*, and (e) both *nd6* and *ndufa13*. Side chains of residues with substitutions between *X. birchmanni* and *X. malinche* are displayed as sticks surrounded by translucent spheres.

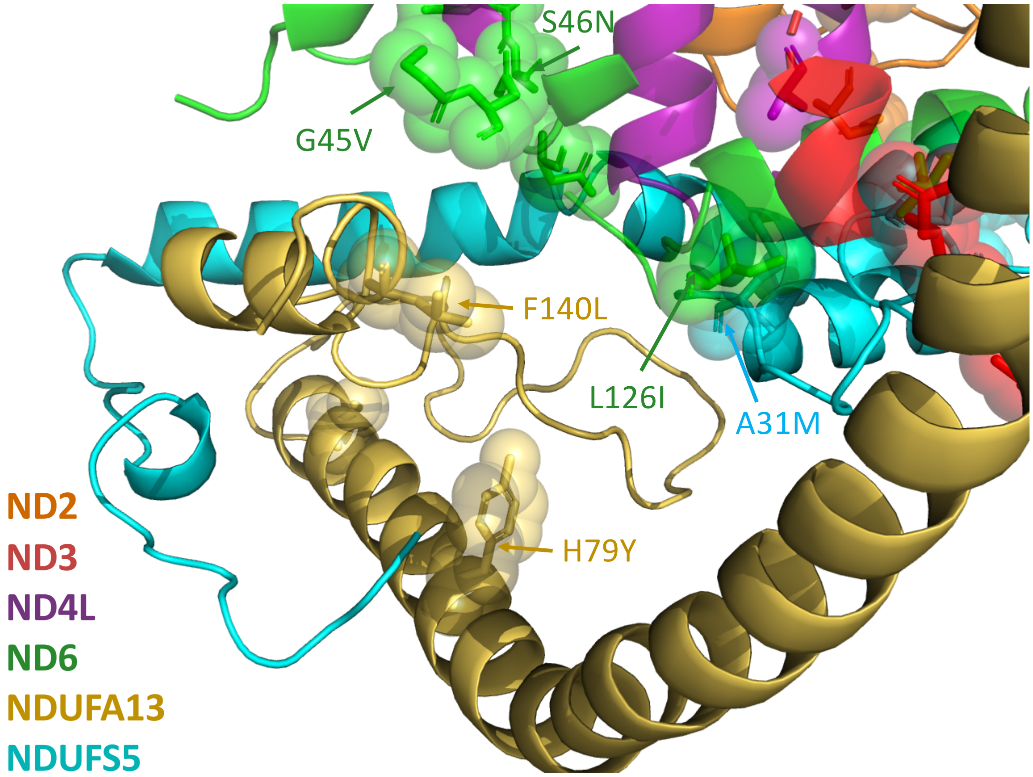

**Fig. S41**. Detailed view of protein interaction interface between *ndufa13, ndufs5* and mitochondrial protein *nd6.* Arrows highlight substitutions in *ndufa13, ndufs5* and mitochondrial proteins in proximity in the model, with alphanumeric codes denoting the *X. malinche* amino acid, the residue number, and the *X. birchmanni* amino acid, from left to right. Colors distinguish proteins, as denoted by colored protein names on right.

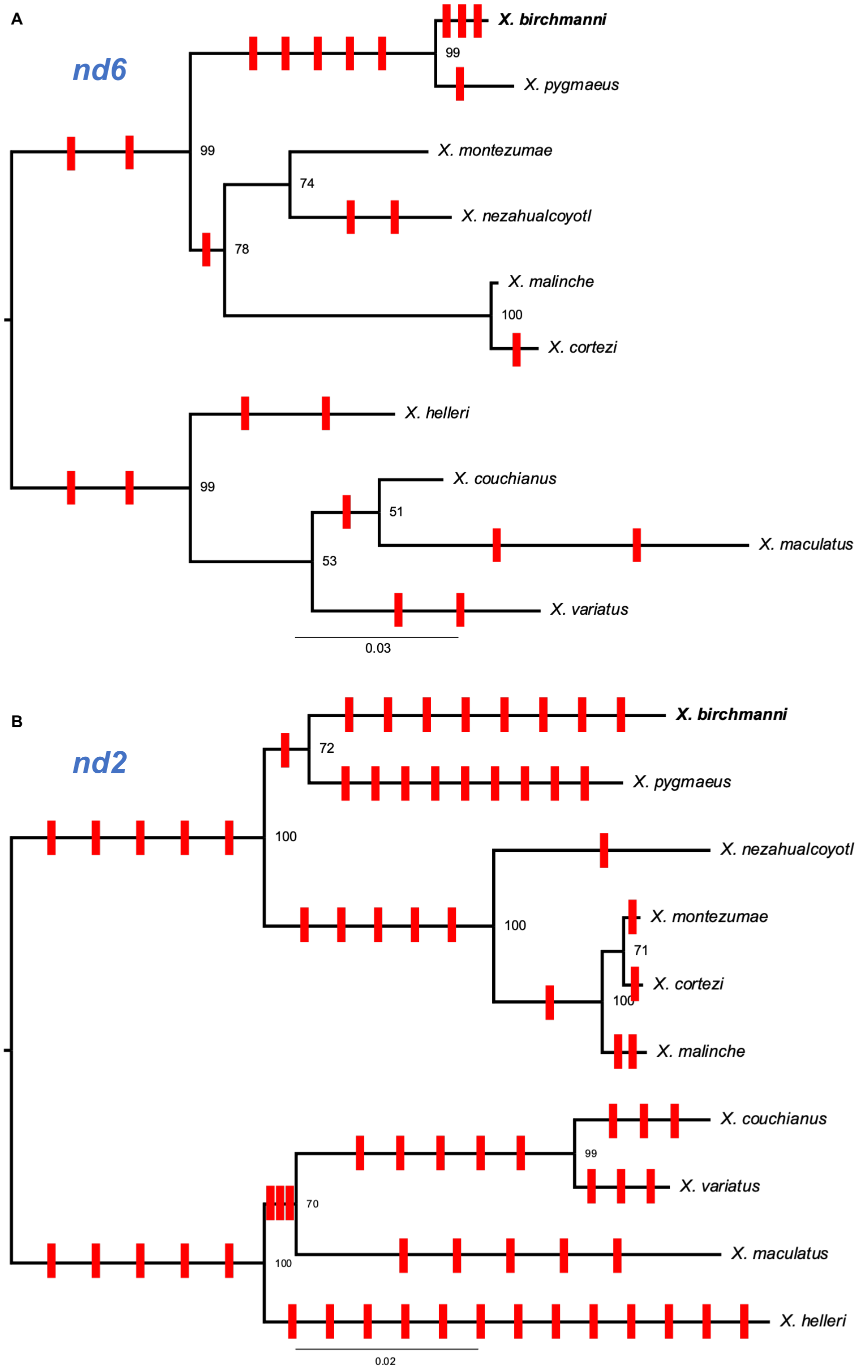

**Fig. S42.** Gene tree for (**A**) *nd6* and (**B**) *nd2* generated with RAxML, highlighting an excess of substitutions along the *X. birchmanni* branch in *nd6*. Scale bar represents number of nucleotide substitutions per site, and derived non-synonymous substitutions are indicated by red ticks along the phylogeny. Note that spacing between ticks is arbitrary and substitutions were placed on branches to maximize parsimony. In some cases, the distribution of substitutions cannot be explained by a single event, in such cases we illustrate the minimum number of events leading to observed distribution of the substitution.

```
xbir      MAPFVCSTLIISLGLGTTMTFASTHWFLAWMGIEINTLAIIPLMTQNHNPRTIEATTKYF  60
xmal      MAPFVCSTLIISLGLGTTMTFASTHWYLAWMGIEINTLAIIPLMTQNHNPRTIEATTKYF  60
xcor      MAPFVCSTLIISLGLGTTMTFASTHWYLAWMGIEINTLAIIPLMTQNHNPRTIEATTKYF  60
          *************************:**********************************

xbir      FAQATASATLLFAAVSNAFLTGEWDILQMNHPLTSTLTTLALAMKIGLAPLHSWMPEVMQ  120
xmal      FAQATASATLLFAAVSNAFLTGGWDILQTNHPLTSTLITLALAMKIGLAPLHSWMPEVMQ  120
xcor      FAQATASATLLFAAVSNAFLTGGWDILQTNHPLTSTLITLALAMKIGLAPLHSWMPEVMQ  120
          ********************** ***** ******** **********************

xbir      GVSLLTGLTLSTWQKLAPLCLIYQIQPDSPSVFTTLGLLSVIVGGWGGLNQVQLRKILAY  180
xmal      GVSLLTGLTLSTWQKLAPLCLIYQIQPNSPSVFTTLGLLSVIVGGWGGLNQVQLRKILAY  180
xcor      GVSLLTGLTLSTWQKLAPLCLIYQIQPNSPSVFTTLGLLSVIVGGWGGLNQIQLRKILAY  180
          ***************************:***********************:********

xbir      SSIAHLGWMILILPFSPPLTLLTLFTYLMMTFSLFSSFMLIRTTHINSLSTSWAKIPALT  240
xmal      SSIAHLGWMIIILPFSPPLTLLTLFTYLMMTFSLFSSFMLIRTTHINSLSTSWAKIPALT  240
xcor      SSIAHLGWMIIILPFSPPLTLLTLFTYLMMTFSLFSSFMLIRTTHINSLSTSWAKIPALT  240
          **********:*************************************************

xbir      ASIPLILLSLGGLPPLTGFLPKWLILQELTKQDLAPIATLAALSSLFSLYFYLRLSYTMT  300
xmal      ASVPLILLSLGGLPPLTGFLPKWLILQELTKQDLAPIATLAALSSLFSLYFYLRLSYTMT  300
xcor      ASVPLILLSLGGLPPLTGFLPKWLILQELTKQDLAPIATLAALSSLFSLYFYLRLSYTMT  300
          **:*********************************************************

xbir      LTMPPNNPAGTLPWRLKPHHNTLPLALTTTMTIYLLPITPAILALFTL      348
xmal      LTMPPNNPTGMLPWRLNPRHNTLPLALTTTTTIYFLPATPAILALFTL      348
xcor      LTMPPNNPTGTLPWRLNPRHNTLPLALTTTTTIYLLPATPAILALFTL      348
          ********:* *****:*:*********** ***:** **********
```

**Fig. S43.** Clustal alignment of mitochondrial gene *nd2* between *X. birchmanni*, *X. malinche* and *X. cortezi*. Colors indicate properties of the amino acid, asterisks indicate locations where the amino acid sequences are identical. Twelve of the fifteen amino acid differences between species are shared between *X. malinche* and *X. cortezi*. Phylogenetic analysis indicates that seven of these substitutions are derived in *X. birchmanni* compared to other *Xiphophorus* species.

```
xbir      MSYLMYILLSGLVLGLVAVASNPSPYFAAFGLVVVAGVGCGVLVVNGGCFLPLILFLIYL  60
xmal      MSYLMYVLLSGLVLGLVAVASNPSPYFAAFGLVVVAGVGCGVLVGSGGCFLSLILFLIYL  60
xcor      MSYLMYVLLSGLVLGLVAVASNPSPYFGAFGLVVVAGVGCGVLVGSGGCFLSLILFLIYL  60
          ******:********************.**************** .***** ********

xbir      GGMLVVFAYSAAMAAEPFPEGWGSWPTLRLMVGYVVGVGFAWGLMGKIWHEEDWLGFDEA  120
xmal      GGMLVVFAYSAAMAAEPFPEGWGSWPTLRLMVGYVVGVGFAWGLMGKIWHEEDWLGFDGM  120
xcor      GGMLVVFAYSAAMAAEPFPEGWGSWPTLRLMVGYVVGVGFAWGLMGKIWHEEDWLGFDGM  120
          **********************************************************

xbir      VDLSVIRGDVGGVSVVYSSGGWLLIGAGWVLLLTLFVVLELTRGLSRGALRAV 173
xmal      VELSVLRGDVGGVSLVYSSGGWLLIGGGWVLLLTLFVVLELTRGLSRGALRAV 173
xcor      VELSVLRGDVGGVSLVYSSGGWLLIGGGWVLLLTLFVVLELTRGLSRGALRAV 173
          *:***:********:***********.**************************
```

**Fig. S44.** Clustal alignment of mitochondrial gene *nd6* between *X. birchmanni*, *X. malinche* and *X. cortezi*. Colors indicate properties of the amino acid, asterisks indicate locations where the amino acid sequences are identical. Ten of the eleven amino acid differences between species are shared between *X. malinche* and *X. cortezi*. Phylogenetic analysis indicates that nine of these substitutions are derived in *X. birchmanni* compared to other *Xiphophorus* species.

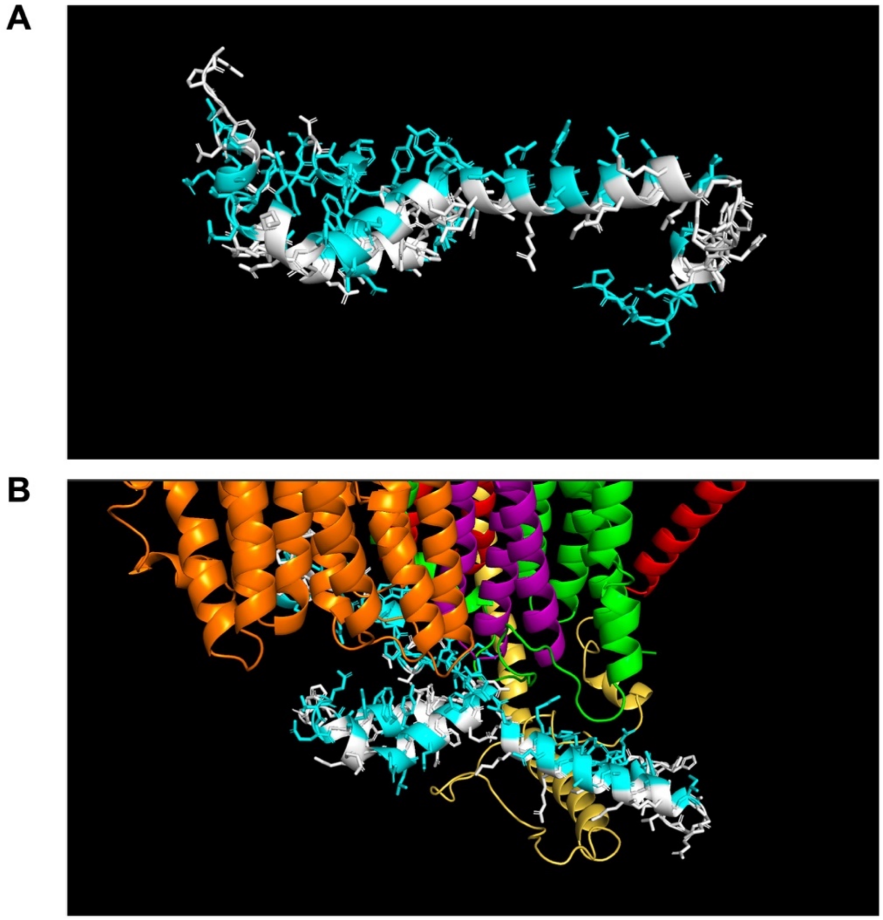

**Fig. S45.** Conservation of amino acids between mouse and *Xiphophorus* in *ndufs5* relative to their positions in the mitochondrial-nuclear protein interface of Complex I. Amino acids in *ndufs5* at the interface of mitochondrial proteins are less likely to be conserved between mouse and *Xiphophorus* (blue – not conserved, white – conserved). In **A** *ndufs5* is shown in isolation, in **B** *ndufs5* is shown with the proximate proteins in Complex I (orange - *nd2,* yellow – *ndufa13*, red – *nd3*, purple – *nd4l*, green – *nd6*). Structures of *Xiphophorus* proteins were modeled using cryoEM structures and RaptorX as described in Supplementary Information 1.4.6.

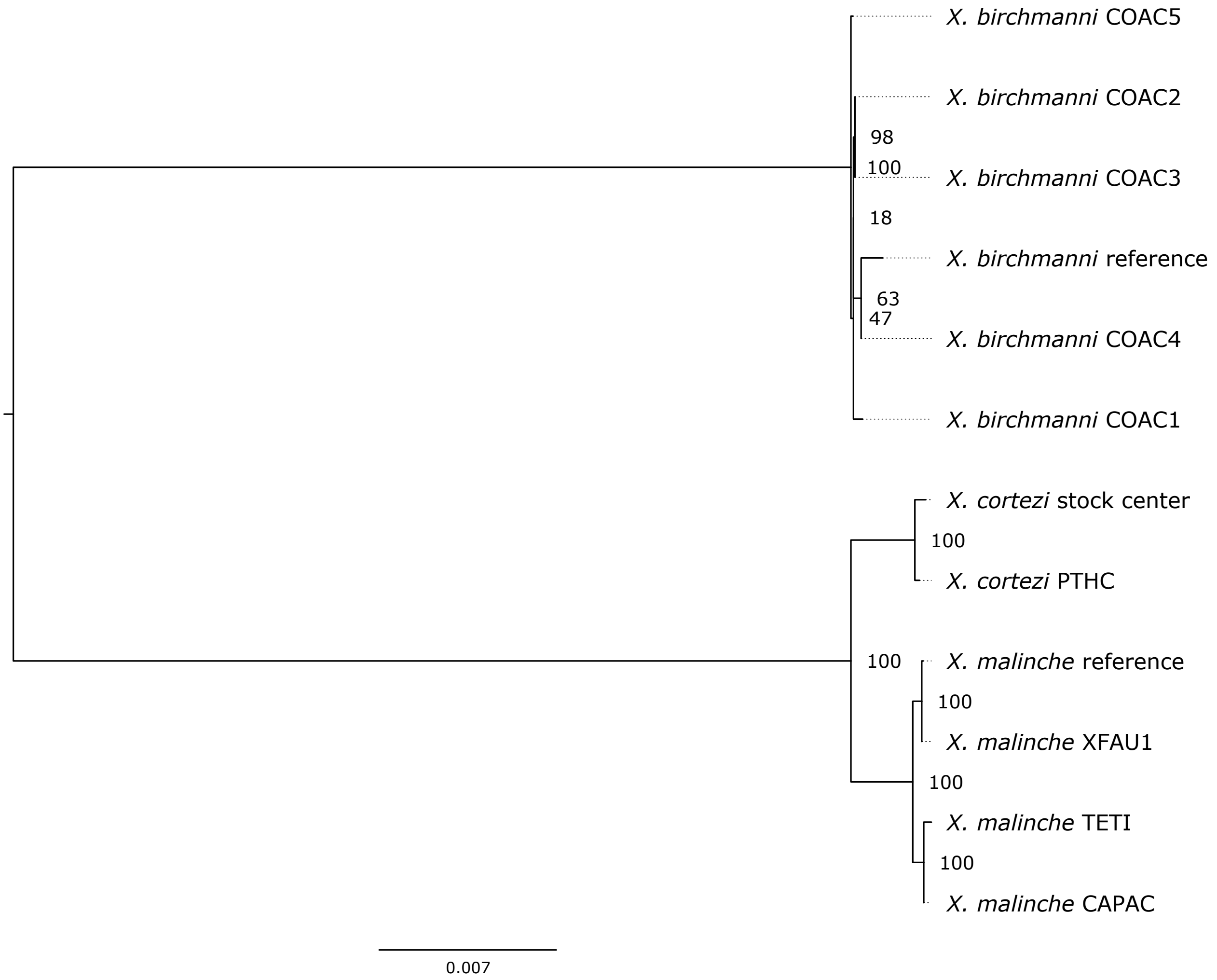

**Fig. S46**. RAxML phylogeny of whole mitochondrial sequences from PacBio HiFi long amplicon sequencing and the *X. birchmanni* and *X. malinche* reference sequences. *X. birchmanni* individuals were sampled from Coacuilco (COAC), *X. malinche* individuals were sampled from Upper Xontla Falls (XFAU), Tetipanchalco (TETI), and CAPAC populations (CAPAC), while *X. cortezi* individuals were sampled from (PTHC) and the *Xiphophorus* Genetic stock center. Numbers at nodes represent support from 100 rapid bootstraps, and branch length is proportional to the number of substitutions per basepair.

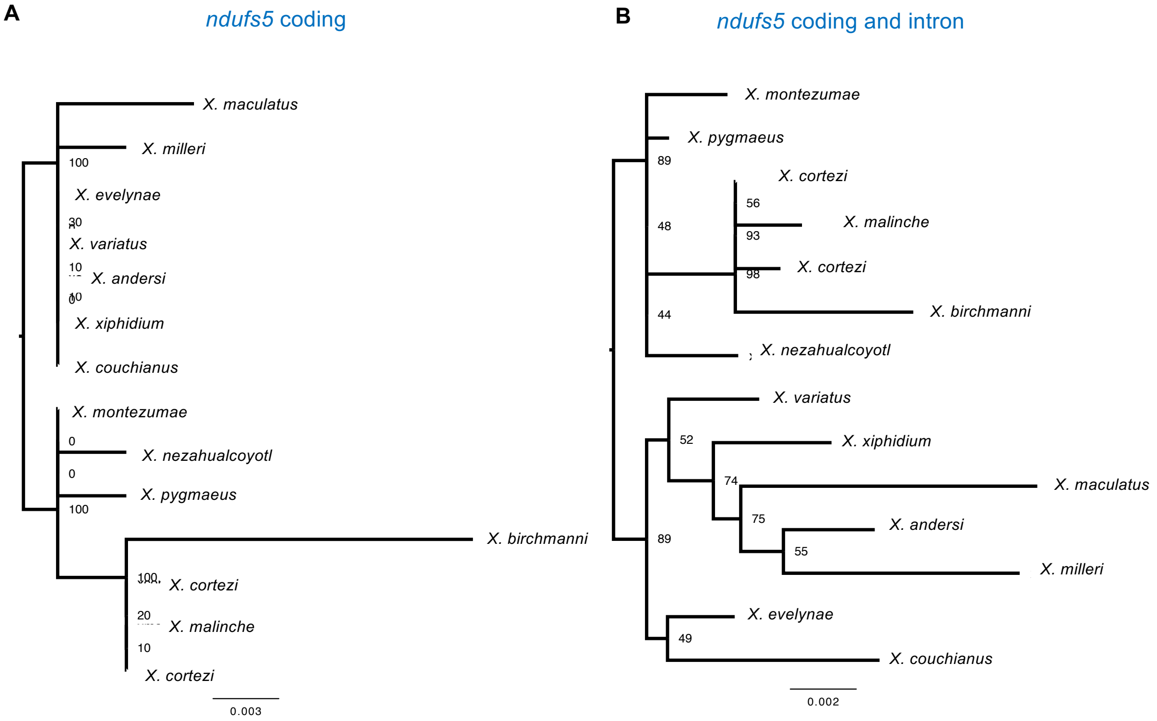

**Fig. S47**. RAxML phylogenies constructed using (**A**) cDNA only and (**B**) cDNA and introns for *Xiphophorus ndufs5*. Bootstrap support from 100 replicates displayed at nodes.

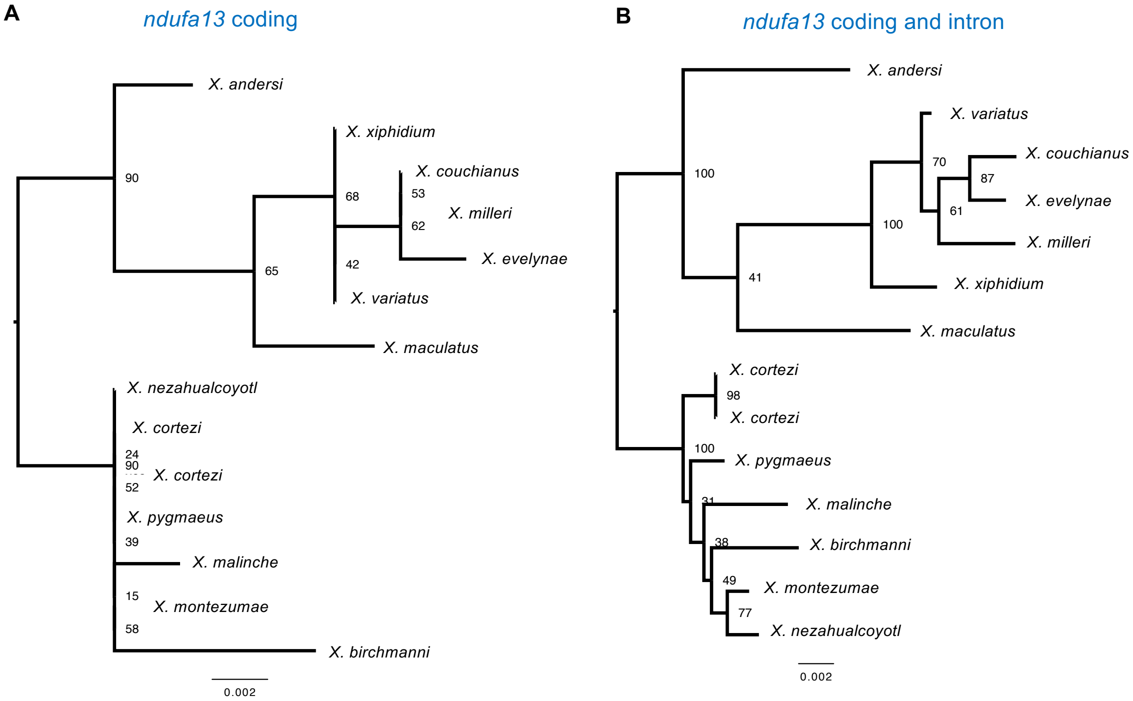

**Fig. S48**. RAxML phylogenies constructed using (**A**) cDNA only and (**B**) cDNA and introns for *Xiphophorus ndufa13*. Bootstrap support from 100 replicates displayed at nodes.

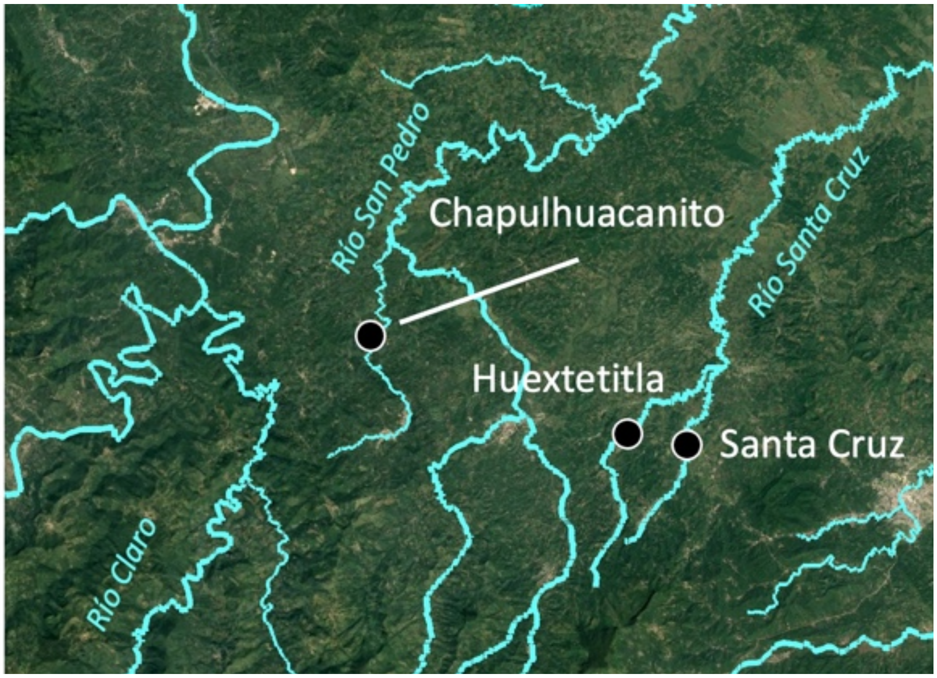

**Fig. S49.** Map of *X. cortezi* × *X. malinche* populations genotyped for mitonuclear incompatibility depletion analysis.

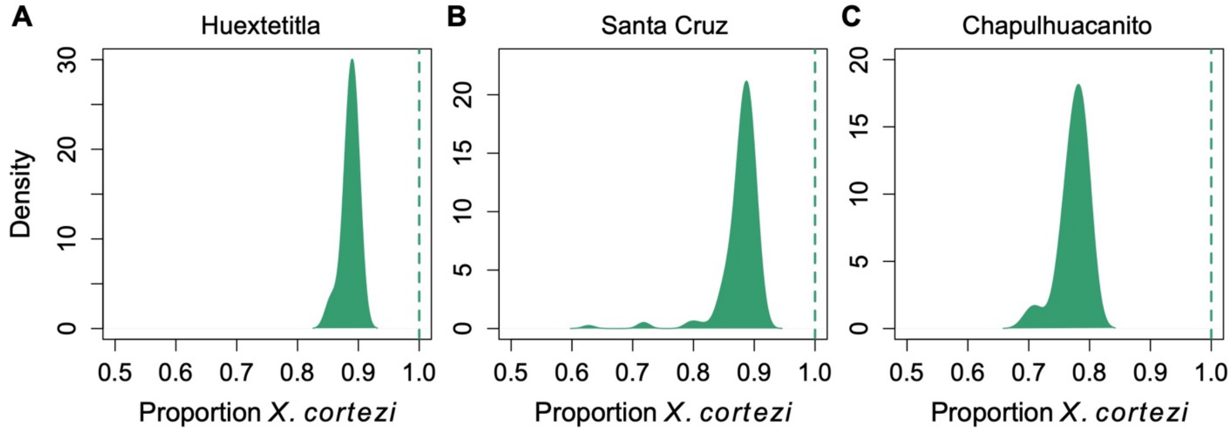

**Fig. S50**. Average genome-wide ancestry in three hybrid populations formed between *X. birchmanni* × *X. cortezi* that are fixed for *X. cortezi* mitochondrial ancestry. Curves represent distribution of ancestry fractions across individuals within the dataset. Dashed lines indicate the mitochondrial ancestry fixed in that population.

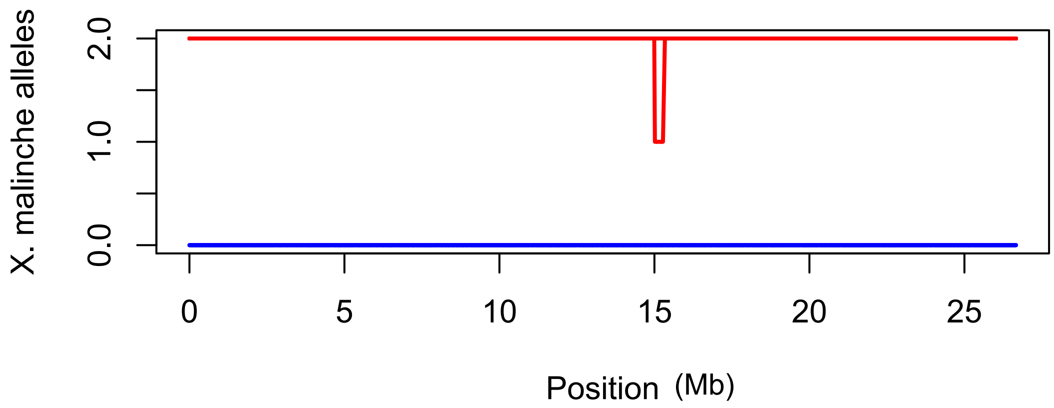

**Fig. S51**. Local ancestry plot for 10 *X. birchmanni* (blue lines) and 10 *X. malinche* individuals (red lines) on chromosome 13 show no signs of erroneous ancestry switching in the region of peak mitonuclear ancestry association (1.7-2.1 Mb).

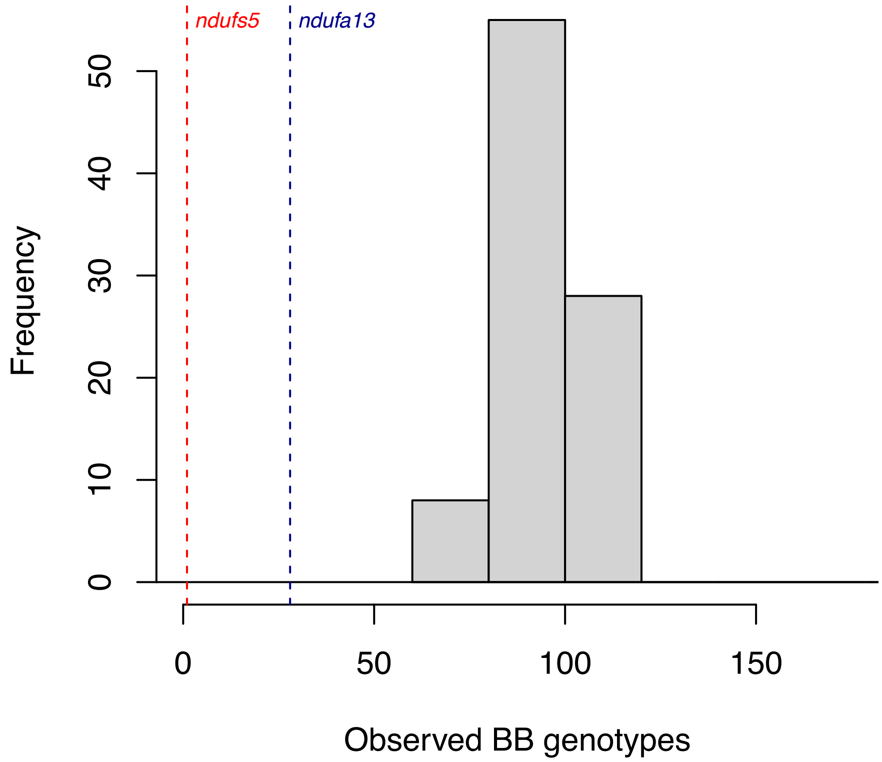

**Fig. S52.** Results of simulations of a dominant lethal nuclear-nuclear hybrid incompatibility using the simulator admix'em. Plotted here are the minimum observed number of homozygous *X. birchmanni* genotypes (BB) for each simulation at either of the two loci involved in the hybrid incompatibility. These simulations indicate that the observation of 0.1% (N=1) and 3% (N=28) BB genotypes at the chromosome 13 and chromosome 6 regions in $F_2$ hybrids is unexpected in the case of a nuclear-nuclear hybrid incompatibility. Dashed vertical lines highlight empirically observed counts of BB genotypes at *ndufs5* (red) and *ndufa13* (blue). Plotted here are the results from 100 simulations.

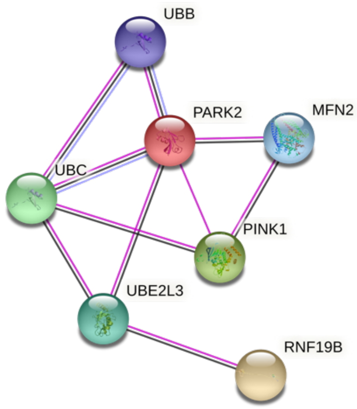

**Fig. S53.** STRING network of another candidate gene evaluated in the chromosome 13 interval. Of the three genes in the interval, *ndufs5* is the only one known to directly interact with any mitochondrial protein. However, another gene in the interval, *rnf19b,* is annotated as interacting with a mitonuclear gene, *park2*, via an intermediary, *ube2l3*. The STRING interaction network for *rnf19b* is shown here. Pink connections show experimentally verified interactions between proteins, purple shows interactions inferred from protein homology, and black lines show interactions inferred by coexpression data.

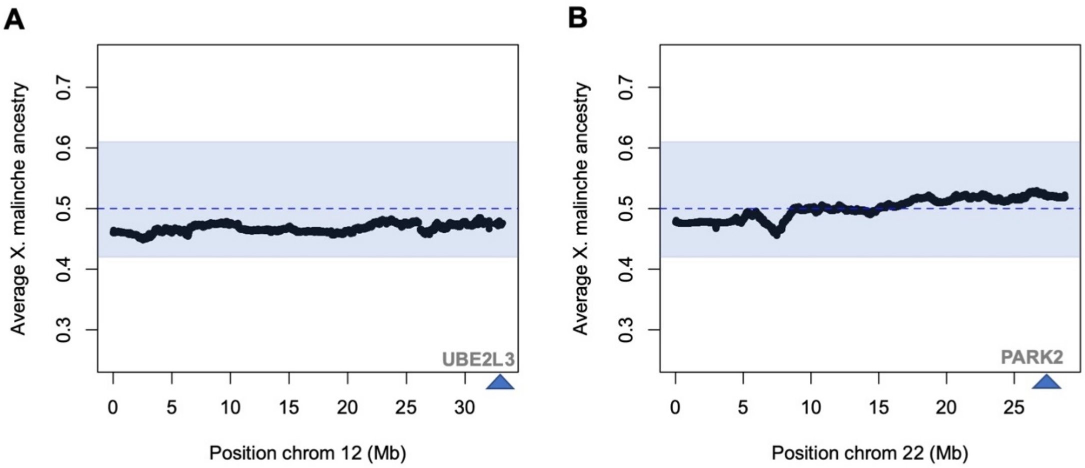

**Fig. S54.** Investigation of possible interaction partners of *rnf19b*. *rnf19b* is physically linked to *ndufs5*, resulting in ancestry and mapping patterns that are nearly identical across the two genes. However, if *rnf19b* interacts with the mitochondria, it is likely through its interactions with *ube2l3* and *park2* (only *park2* is known to be a mitonuclear gene; Fig. S53). Thus, if these genes mediated the incompatibility, we would expect to see segregation distortion in $F_2$ hybrids with an *X. malinche* mitochondria surrounding these genes, as we do at *ndufa13*. However, we see no evidence of segregation distortion at either of these genes. Black points show average *X. malinche* ancestry across 943 individuals at each ancestry informative site. The dotted blue line shows expected ancestry given the cross design. The blue envelop shows the 99% quantiles of *X. malinche* ancestry at all ancestry informative sites genome wide. Blue triangles show the approximate location of each gene.

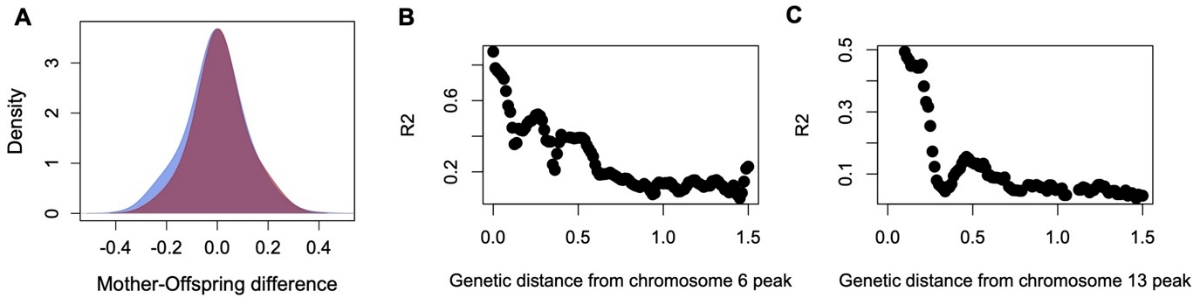

**Fig. S55.** Evaluation of possible confounding factors in our admixture mapping dataset. **A.** Ancestry assortative mating in hybrid populations can drive population structure that elevates background admixture LD and increases false positive rates in admixture mapping studies. We evaluated whether mother-embryo ancestry patterns were suggestive of assortative mating by comparing observed mother-embryo ancestry differences (red) to those expected under random mating (blue). Our data indicates that the observed mother-embryo ancestry differences are consistent with random mating by ancestry in our admixture mapping hybrid population. **B & C** Assembly errors are another potential source of LD between physically unlinked markers. Here we show that the peak markers in the chromosome 6 and chromosome 13 peaks are also in LD with nearby markers and that LD decays predictably over genetic distance (plotted here in cMs). This pattern is inconsistent with an assembly error driving associations between markers.

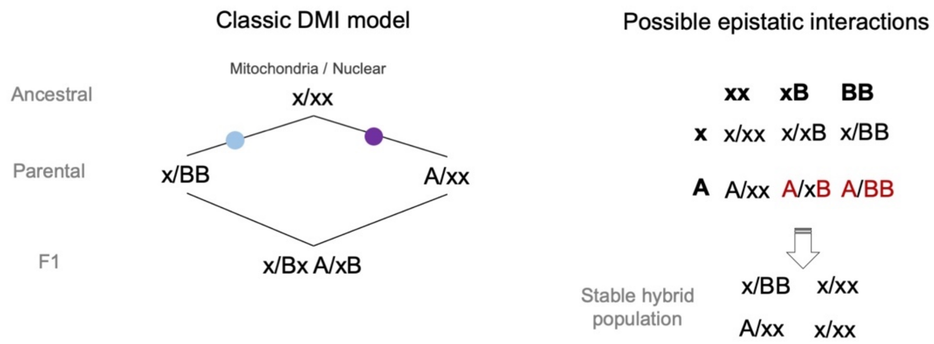

**Fig. S56.** Classic DMI model for the evolution of a mitonuclear hybrid incompatibility. A substitution in the nuclear lineage of one species (x → B) and an interacting mitochondrial gene in another species (x → A) leads to an incompatibility in hybrids when the species with the A mitochondrial genotype is the mother in a cross (right panel). However, this type of incompatibility is quickly resolved in hybrid populations which are predicted to revert to ancestral or transitional genotype combinations. In contrast, we see bidirectional incompatibility between the nuclear gene *ndufs5* and mitochondria derived from either *X. birchmanni* or *X. malinche* (Fig. 1C), suggesting that these incompatibilities should form stable barriers to gene flow between species.

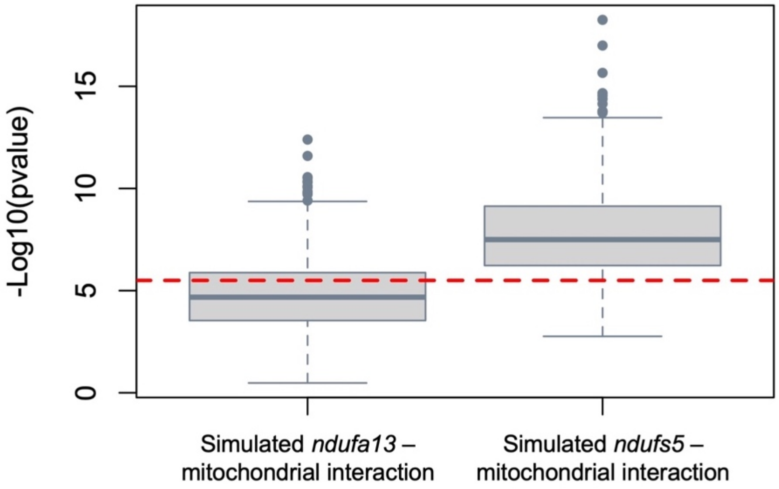

**Fig. S57.** Results of power simulations evaluating our ability to detect genome-wide significant signals in admixture mapping. Based on the results of ABC simulations (Fig. 1, S3) and inference about the genetic architecture of the chromosome 6 (*ndufa13*) and chromosome 13 (*ndufa13*) incompatibilities (Fig. 1) we performed simulations mimicking selection against these hybrid incompatibilities in our admixture mapping dataset (see Supplementary Information 1.2.3). We found that based on these simulations (N=1,000) we had only marginal power to detect the interaction involving *ndufa13* at our genome-wide significance threshold (red dashed line). However, we inferred that our power to detect the interaction involving *ndufs5* was much better. This result is notable because it highlights that the architecture of the incompatibility involving *ndufa13* (Fig. 1) makes it more difficult to detect.

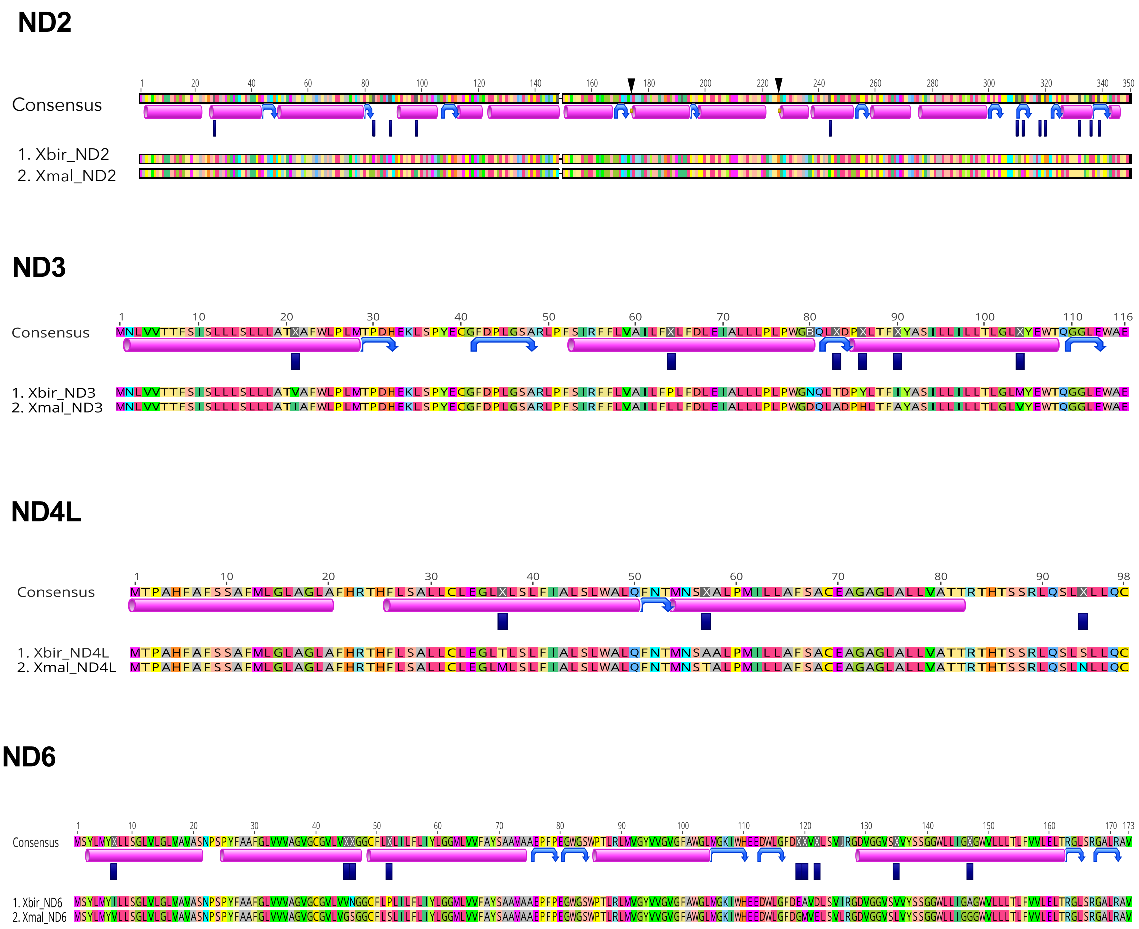

**Fig. S58.** Alignments of *X. birchmanni* and *X. malinche* mitochondrial genes modeled in RaptorX analyses with secondary structure inferred with STRIDE annotated on the consensus sequence. Magenta cylinders indicate the locations of alpha helices, blue curved arrows indicate turns, and black triangles indicate beta bridges. Dark blue rectangles indicate the locations of substitutions that differentiate *X. birchmanni* and *X. malinche.* Alignments were plotted using Geneious.

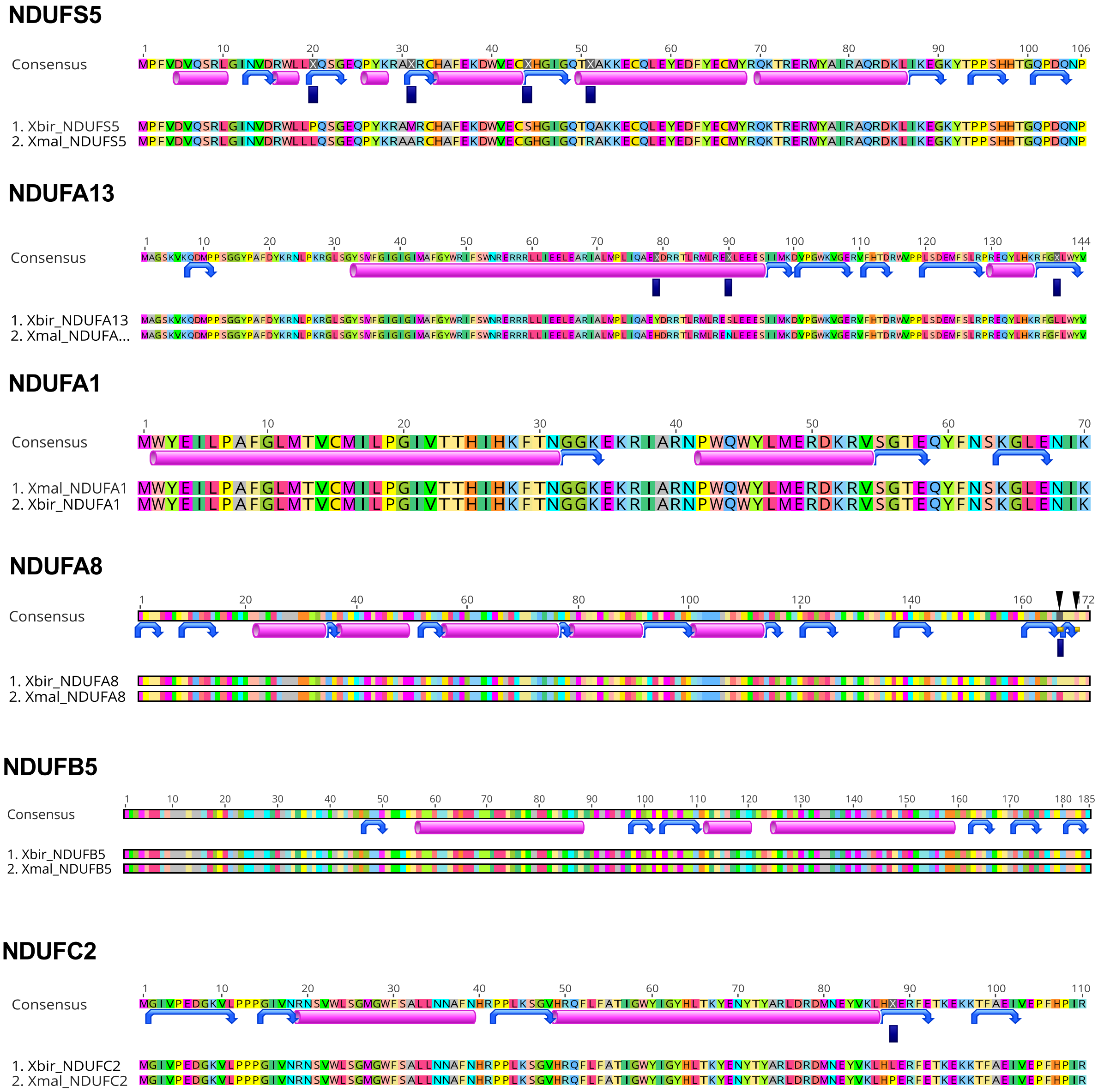

**Fig. S59.** Alignments of *X. birchmanni* and *X. malinche* nuclear genes modeled in RaptorX analyses with secondary structure inferred with STRIDE annotated on the consensus sequence. Magenta cylinders indicate the locations of alpha helices, blue curved arrows indicate turns, and black triangles indicate beta bridges. Dark blue rectangles indicate the locations of substitutions that differentiate *X. birchmanni* and *X. malinche.* Alignments were plotted using Geneious.

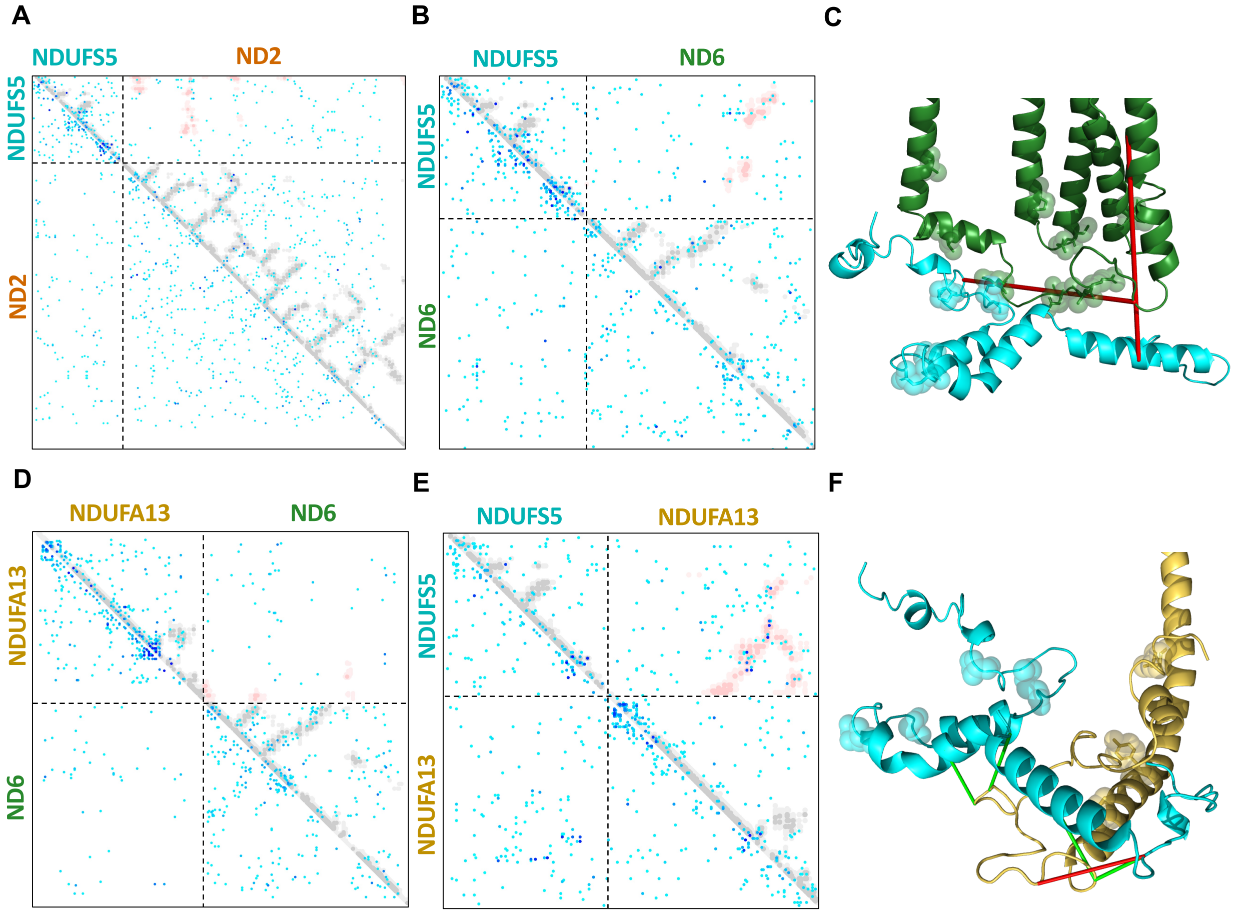

**Fig. S60.** Results from GREMLIN analyses of coevolution signals between *ndufs5* and *nd2* (A), *ndufs5* and *nd6* (B,C), *ndufa13* and *nd6* (D) and *ndufs5* and *ndufa13* (E,F). (A,B,D,E) Dotplots show the full results of GREMLIN, with X and Y coordinates displaying pairwise GREMLIN covariance scores. Blue dots indicate that covariance between the two sites is higher than average, and darker blue indicates stronger covariance. The darkest blue dots highlight GREMLIN scores 2.5 times greater than average. Dashed lines delineate interactions within (upper left and lower right quadrants) and between (upper right and lower left quadrants) proteins. Superimposed grey points indicate intraprotein contacts in the RaptorX *Xiphophorus* Complex I model, while red points indicate interprotein contacts. (C,F) RaptorX structures for the interaction of *ndufs5* and *nd6* (C) and *ndufs5* and *ndufa13* (F) with GREMLIN pairs scoring higher than 2.5 indicated by colored lines. Green lines are used to indicate pairs separated by less than 8 Å, and red lines greater than 8 Å.

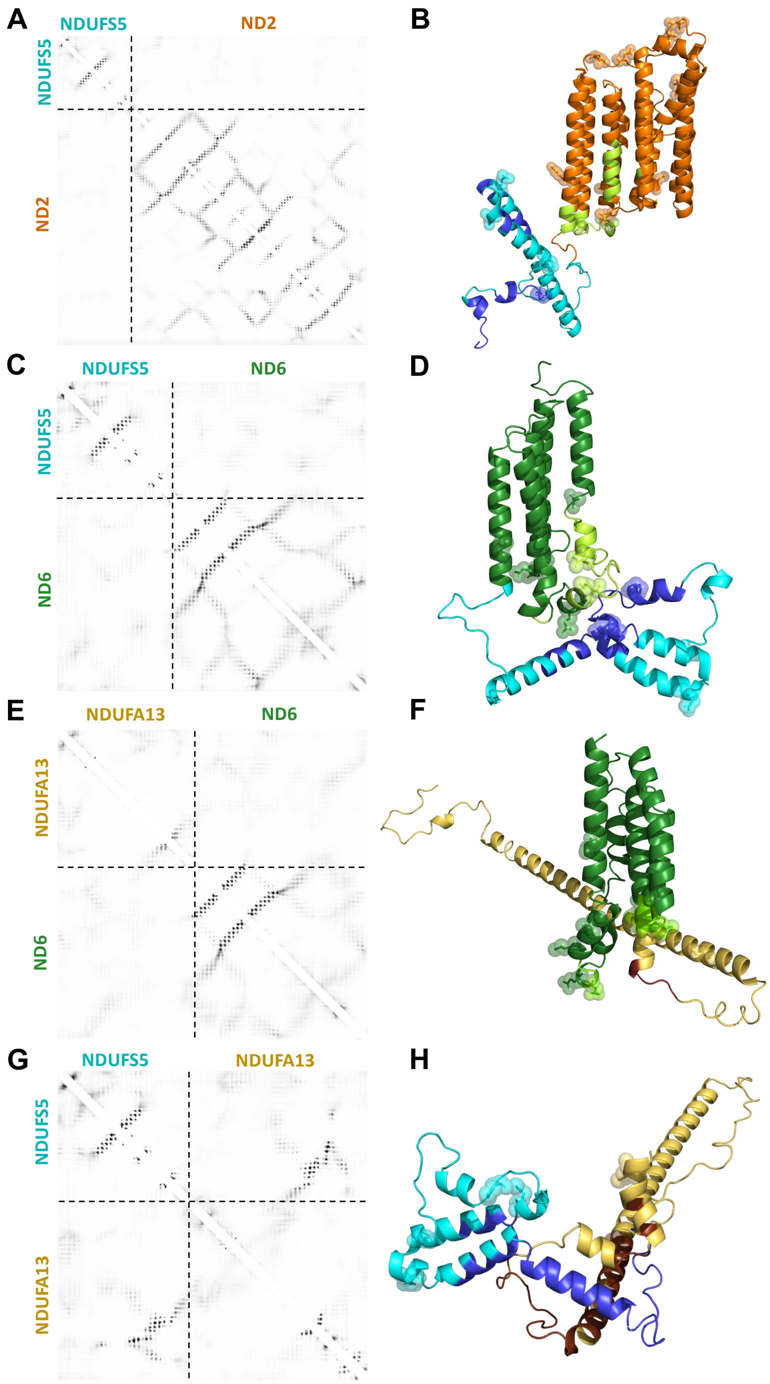

**Fig. S61.** RaptorX-Contact analysis between (A-B) *ndufs5* and *nd2*, (C-D) *ndufs5* and *nd6*, (E-F) *ndufa13* and *nd6*, and (G-H) *ndufs5* and *ndufa13*. Left panels display probabilities of contact (defined as inter-residue distance ≤ 8 Å) within and between proteins based on analysis of alignments. Darker shading indicates higher probability of contact. Dashed lines delineate interactions within (upper left and lower right quadrants) and between (upper right and lower left) proteins. Right panels display the highest-scoring model generated by RaptorX-Contact for each alignment. Residues in contact (≤ 8 Å) between these proteins in template-based models (Fig. 3) are shown in blue (*ndufs5*), light green (*nd2* and *nd6*), and brown (*ndufa13*).

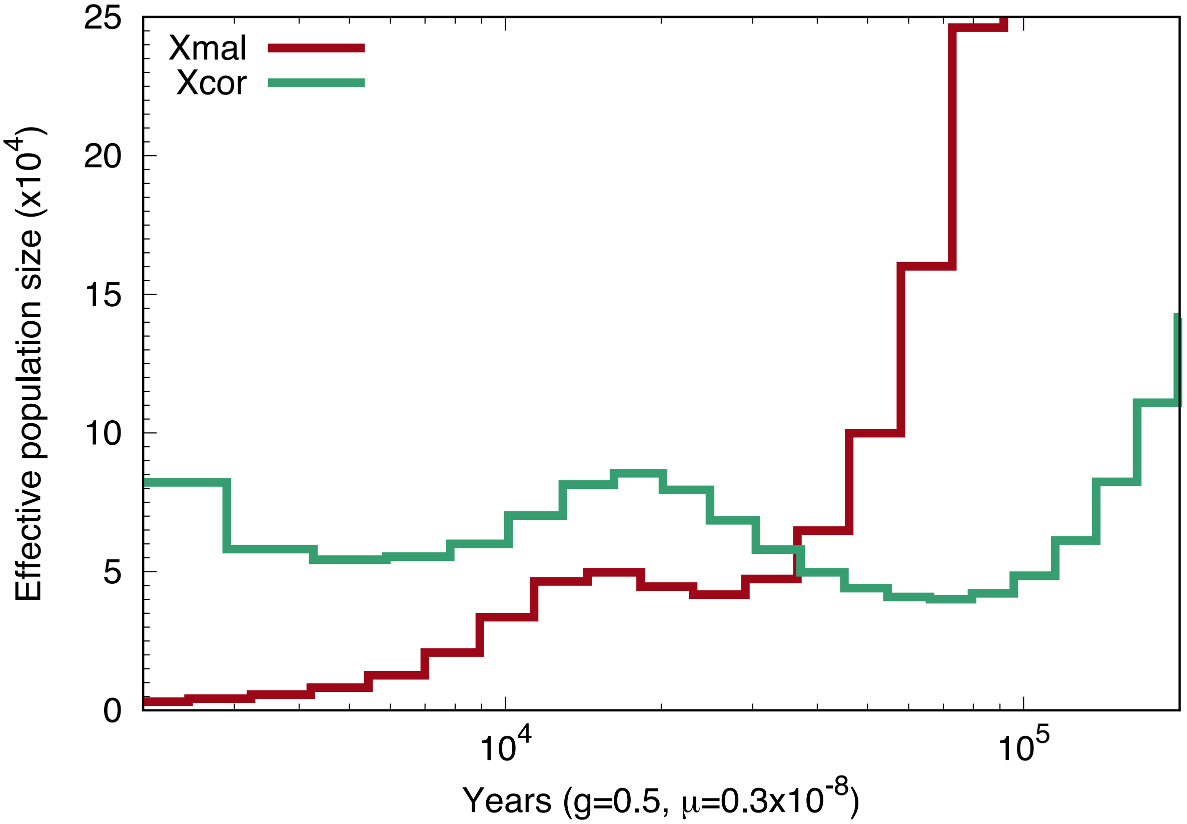

**Fig. S62.** PSMC results analyzing population history from a single whole-genome sample of *X. malinche* (red) and *X. cortezi* (green). Analysis was conducted with a ρ/θ ratio of 2, generation time of two generations per year, and mutation rate of $3.5 \times 10^{-9}$. These inferred population histories were used as the basis for simulations described in *Supplementary Information 1.5.4 Introgression of an X. malinche mitochondrial haplotype incompatible with X. birchmanni ndufs5 and ndufa13*.

### 3 Supplementary Tables

**Tables S1 – S3 are provided as Excel files**

**Table S1.** Locations of annotated Complex I genes in the *X. birchmanni* genome assembly.

**Table S2.** Genes found in the admixture mapping QTL region on chromosome 6.

**Table S3.** Genes found in the admixture mapping QTL region on chromosome 15.

**Table S4.** Survival of individuals from $F_1$ hybrid broods born prematurely (developmental stage <12, following Haynes 1995) in the lab. Note that $F_1$ hybrids are not impacted by the mitonuclear hybrid incompatibility. These results are consistent with findings in other poecilid species, see Supplementary Information 1.3.1.

| **Brood size** | **Developmental stage at birth** | **Mortality rate** | **Number of days survived past birth** |
|---|---|---|---|
| 3 | 9 | 100% | 1-2 |
| 7 | 8 | 100% | 1-7 |
| 2 | 9 | 100% | 1-2 |
| 13 | 9 | 92% (N=1 survived) | 1-2 |

**Table S5.** Results of linear regression analysis of individual length as a function of *ndufs5* genotype, *ndufa13* genotype, and an estimate of the expected length for the focal brood (see Supplementary Information 1.3.5). Brood was included as a blocking variable in the analysis and physically damaged individuals were excluded. Linear regression model: length_mm ~ ndufs5 * ndufa13 * brood_length + Mother

| | D.F. | Sum Sq | Mean Sq | *F* value | *P value* |
|---|---|---|---|---|---|
| *ndufs5* | 2 | 118.9 | 59.5 | 40.622 | 4.68e-15 *** |
| *ndufa13* | 2 | 13.3 | 6.7 | 4.547 | 0.012 * |
| brood_length | 1 | 360.9 | 360.9 | 246.58 | < 2e-16 *** |
| Mother | 8 | 9.8 | 1.2 | 0.835 | 0.573 |
| *ndufs5: ndufa13* | 4 | 1.4 | 0.4 | 0.24 | 0.916 |
| *ndufs5:* brood_length | 2 | 85.4 | 42.7 | 29.184 | 1.44e-11 *** |
| *ndufa13:* brood_length | 2 | 0.6 | 0.3 | 0.209 | 0.811 |
| *ndufs5: ndufa13:* brood_length | 4 | 4.2 | 1.1 | 0.72 | 0.579 |
| Residuals | 164 | 240 | 1.5 | | |
| Signif. Codes: 0 ‘***’ 0.001 ‘**’ 0.01 ‘*’ 0.05 ‘.’ 0.1 ‘ ‘ 1 | | | | | |
| **31 observations deleted due to missingness** | | | | | |

**Table S6.** Results of linear regression analysis of heart rate as a function of *ndufs5* genotype, *ndufa13* genotype, and standard length, with brood as a blocking variable. Physically damaged embryos were excluded from the analysis. Linear regression model: heart_rate ~ ndufs5 * ndufa13 * length + Mother

| | D.F. | Sum Sq | Mean Sq | *F* value | *P value* |
|---|---|---|---|---|---|
| *ndufs5* | 2 | 7726 | 3863 | 38.592 | 3.65e-14 *** |
| *ndufa13* | 2 | 1795 | 897 | 8.965 | 0.000213 *** |
| length | 1 | 6158 | 6158 | 61.521 | 8.73e-13 *** |
| Mother | 9 | 2984 | 332 | 3.312 | 0.00105 ** |
| *ndufs5: ndufa13* | 4 | 346 | 86 | 0.863 | 0.487701 |
| *ndufs5:* length | 2 | 21 | 11 | 0.107 | 8.98e-01 |
| *ndufa13:* length | 2 | 1412 | 706 | 7.053 | 0.001193 ** |
| *ndufs5: ndufa13:* length | 4 | 231 | 58 | 0.578 | 0.679271 |
| Residuals | 145 | 14514 | 100 | | |
| Signif. Codes: 0 '***' 0.001 '**' 0.01 '*' 0.05 '.' 0.1 ' ' 1 | | | | | |
| **49 observations deleted due to missingness** | | | | | |

**Table S7.** Results of linear regression analysis of head width as a function of *ndufs5* genotype, *ndufa13* genotype, and standard length, with brood as a blocking variable. Physically damaged embryos were excluded from the analysis. Linear regression model: head_width ~ ndufs5 * ndufa13 * length + Mother

| | D.F. | Sum Sq | Mean Sq | *F* value | *P value* |
|---|---|---|---|---|---|
| *ndufs5* | 2 | 16.92 | 8.46 | 460.049 | < 2e-16 *** |
| *ndufa13* | 2 | 1.05 | 0.52 | 28.432 | 2.66e-11 *** |
| Length | 1 | 33.08 | 33.08 | 1799.245 | < 2e-16 *** |
| Mother | 9 | 2 | 0.22 | 12.073 | 1.83e-14 *** |
| *ndufs5: ndufa13* | 4 | 0.15 | 0.04 | 2.051 | 0.08977 **.** |
| *ndufs5:*Length | 2 | 0.25 | 0.13 | 6.916 | 1.31e-03 *** |
| *ndufa13:* Length | 2 | 0.03 | 0.01 | 0.746 | 0.47605 |
| *ndufs5: ndufa13:* Length | 4 | 0.06 | 0.02 | 0.879 | 0.47808 |
| Residuals | 161 | 2.96 | 0.02 | | |
| Signif. Codes: 0 '***' 0.001 '**' 0.01 '*' 0.05 '.' 0.1 ' ' 1 | | | | | |
| **33 observations deleted due to missingness** | | | | | |

**Table S8.** Results of linear regression analysis of sinu-atrium width as a function of *ndufs5* genotype, *ndufa13* genotype, and standard length, with brood as a blocking variable. Physically damaged embryos were excluded from the analysis. Linear regression model: sinuatrium_width ~ ndufs5 * ndufa13 * length + Mother

| | D.F. | Sum Sq | Mean Sq | *F* value | *P value* |
|---|---|---|---|---|---|
| *ndufs5* | 2 | 0.00293 | 0.001465 | 0.78 | 4.61e-01 |
| *ndufa13* | 2 | 0.01703 | 0.008515 | 4.534 | 0.0125 * |
| length | 1 | 0.00072 | 0.000718 | 0.383 | 5.37e-01 |
| Mother | 9 | 0.08137 | 0.009042 | 4.814 | 1.53e-05 *** |
| *ndufs5: ndufa13* | 4 | 0.00067 | 0.000167 | 0.089 | 0.9858 |
| *ndufs5:*length | 2 | 0.00356 | 0.00178 | 0.948 | 3.90e-01 |
| *ndufa13:* length | 2 | 0.00327 | 0.001634 | 0.87 | 0.4214 |
| *ndufs5: ndufa13:* length | 4 | 0.00628 | 0.001571 | 0.836 | 0.5045 |
| Residuals | 128 | 0.24038 | 0.001878 | | |
| Signif. Codes: 0 ‘***’ 0.001 ‘**’ 0.01 ‘*’ 0.05 ‘.’ 0.1 ‘ ‘ 1 | | | | | |
| **66 observations deleted due to missingness** | | | | | |

**Table S9.** Results of linear regression analysis of yolk diameter as a function of *ndufs5* genotype, *ndufa13* genotype, and standard length, with brood as a blocking variable. Physically damaged embryos were excluded from the analysis. Linear regression model: yolk_diam ~ ndufs5 * ndufa13 * length + Mother

| | D.F. | Sum Sq | Mean Sq | *F* value | *P value* |
|---|---|---|---|---|---|
| *ndufs5* | 2 | 0.365 | 0.182 | 4.214 | 1.65e-02 * |
| *ndufa13* | 2 | 0.058 | 0.029 | 0.674 | 5.11e-01 |
| length | 1 | 13.911 | 13.911 | 321.289 | <2e-16 *** |
| Mother | 9 | 6.211 | 0.69 | 15.938 | <2e-16 *** |
| *ndufs5: ndufa13* | 4 | 0.117 | 0.029 | 0.676 | 0.6099 |
| *ndufs5:*length | 2 | 0.073 | 0.036 | 0.841 | 4.33e-01 |
| *ndufa13:* length | 2 | 0.019 | 0.009 | 0.215 | 0.8064 |
| *ndufs5: ndufa13:* length | 4 | 0.156 | 0.039 | 0.901 | 0.4647 |
| Residuals | 155 | 6.711 | 0.043 | | |
| Signif. Codes: 0 ‘***’ 0.001 ‘**’ 0.01 ‘*’ 0.05 ‘.’ 0.1 ‘ ‘ 1 | | | | | |
| **39 observations deleted due to missingness** | | | | | |

**Table S10.** Results of linear regression analysis of blank-corrected oxygen consumption rate as a function of *ndufs5* genotype, *ndufa13* genotype, and standard length, with respirometry batch as a blocking variable. Physically damaged embryos were excluded from the analysis. Linear regression model: mO2 ~ ndufs5 * ndufa13 * length + Batch

| | D.F. | Sum Sq | Mean Sq | *F* value | *P value* |
|---|---|---|---|---|---|
| *ndufs5* | 2 | 351696 | 175848 | 31.108 | 5.58e-12 *** |
| *ndufa13* | 2 | 3739 | 1870 | 0.331 | 7.19e-01 |
| length | 1 | 519260 | 519260 | 91.857 | < 2e-16 *** |
| Batch | 10 | 505436 | 50544 | 8.941 | 2.20e-11 *** |
| *ndufs5: ndufa13* | 4 | 25340 | 6335 | 1.121 | 0.34909 |
| *ndufs5:*length | 2 | 38141 | 19071 | 3.374 | 3.70e-02 * |
| *ndufa13:* length | 2 | 55533 | 27766 | 4.912 | 0.00862 ** |
| *ndufs5: ndufa13:* length | 4 | 98747 | 24687 | 4.367 | 0.0023 ** |
| Residuals | 146 | 825323 | 5653 | | |
| Signif. Codes: 0 ‘***’ 0.001 ‘**’ 0.01 ‘*’ 0.05 ‘.’ 0.1 ‘ ‘ 1 | | | | | |
| **17 observations deleted due to missingness** | | | | | |

**Table S11.** Results of linear regression analysis of the proportion of respiration lost after exposure to rotenone as a function of *ndufs5* genotype, *ndufa13* genotype, and standard length, with respirometry batch as a blocking variable. Physically damaged embryos were excluded from the analysis as were outliers in post-rotenone relative respiration (see Supplementary Information 1.3.5). Linear regression model: prop_rotenone_sensitive ~ ndufs5 * ndufa13 * length + Mother

| | D.F. | Sum Sq | Mean Sq | *F* value | *P value* |
|---|---|---|---|---|---|
| *ndufs5* | 2 | 0.109 | 0.05475 | 1.651 | 1.96e-01 |
| *ndufa13* | 2 | 0.009 | 0.00465 | 0.14 | 8.69e-01 |
| length | 1 | 0.28 | 0.2798 | 8.438 | 4.41e-03 ** |
| Mother | 10 | 2.076 | 0.20762 | 6.261 | 1.33e-07 *** |
| *ndufs5: ndufa13* | 4 | 0.403 | 0.10073 | 3.038 | 0.02016 * |
| *ndufs5:*length | 2 | 0.089 | 0.04468 | 1.347 | 2.64e-01 |
| *ndufa13:* length | 2 | 0 | 0.00022 | 0.007 | 0.99324 |
| *ndufs5: ndufa13:* length | 4 | 0.218 | 0.0545 | 1.643 | 0.16812 |
| Residuals | 115 | 3.814 | 0.03316 | | |
| Signif. Codes: 0 ‘***’ 0.001 ‘**’ 0.01 ‘*’ 0.05 ‘.’ 0.1 ‘ ‘ 1 | | | | | |
| **1 observations deleted due to missingness** | | | | | |

**Table S12.** Percent identity and percent similarity between *Mus* proteins used in RaptorX modeling and corresponding swordtail proteins.

| | Percent identity with *Mus* | Percent similarity with *Mus* (BLOSUM 45) |
|---|---|---|
| ND2 | 47% | 75% |
| ND3 | 63% | 78% |
| ND4L | 47% | 80% |
| ND6 | 40% | 74% |
| NDUFA8 | 79% | 92% |
| NDUFA13 | 72% | 90% |
| NDUFC2 | 44% | 77% |
| NDUFS5 | 75% | 89% |
| NDUFA1 | 64% | 93% |
| NDUFB5 | 57% | 84% |

**Table S13.** Amino acid length in swordtails and mammalian species of Complex I subunits modelled in RaptorX. Mammalian sequence lengths are based on Uniprot entries.

| | Length in swordtails (insertions or deletions relative to *Mus*) | Length in *Mus* | Length in humans | Length in *Ovis* | Length in *Bos* |
|---|---|---|---|---|---|
| ND2 | 349 (4) | 345 | 347 | 347 | 347 |
| ND3 | 116 (1) | 115 | 115 | 115 | 115 |
| ND4L | 98 (0) | 98 | 98 | 98 | 98 |
| ND6 | 173 (11) | 172 | 174 | 175 | 175 |
| NDUFA8 | 172 (0) | 172 | 172 | 172 | 172 |
| NDUFA13 | 144 (0) | 144 | 144 | 144 | 144 |
| NDUFC2 | 110 (10) | 120 | 119 | 120 | 122 |
| NDUFS5 | 106 (0) | 106 | 106 | 106 | 106 |
| NDUFA1 | 70 (0) | 70 | 70 | 70 | 70 |
| NDUFB5 | 185 (4) | 189 | 189 | 189 | 189 |

**Table S14.** Effects of structural model approach (program and template) on calculated distances between interspecific substitutions in *X. birchmanni* nuclear genes and four nearby substitutions in *nd6*. The mean ± SD is listed across five replicate MODELLER runs. It is absent for RaptorX modelling results, where one model was constructed. Distance was calculated between alpha carbons except in the case of *ndufa13* tyrosine at position 79, where the side chain's OH group was chosen as a landmark due to its large size and stable orientation within an alpha helix.

| ***ndufs5 M31*** | | | | |
|---|---|---|---|---|
| **Model** | **119E** | **120A** | **122D** | **126I** |
| **RaptorX** | 23.79 | 20.44 | 13.76 | 3.48 |
| **5LNK** | 8.93 ± 1.23 | 9.53 ± 2.66 | 12.42 ± 1.95 | 14.61 ± 1.19 |
| **5LDW** | 10.22 ± 1.30 | 9.95 ± 1.00 | 11.75 ± 1.43 | 15.21 ± 1.55 |
| **5XTC** | 8.38 ± 1.96 | 10.64 ± 1.60 | 11.60 ± 1.13 | 14.80 ± 0.48 |
| **6G2J** | 10.68 ± 3.24 | 10.32 ± 1.48 | 8.71 ± 3.37 | 11.23 ± 1.94 |
| ***ndufa13 Y79*** | | | | |
| **Model** | **119E** | **120A** | **122D** | **126I** |
| **RaptorX** | 27.68 | 23.86 | 16.81 | 13.08 |
| **5LNK** | 7.03 ±1.97 | 6.42 ± 1.79 | 8.48 ± 2.01 | 16.25 ± 1.64 |
| **5LDW** | 10.88 ± 1.33 | 7.61 ± 1.72 | 5.29 ± 0.75 | 12.69 ± 1.28 |
| **5XTC** | 9.30 ± 1.29 | 7.81 ± 0.82 | 5.86 ± 0.36 | 15.08 ± 0.58 |
| **6G2J** | 6.92 ± 1.77 | 7.58 ± 0.95 | 6.28 ± 2.64 | 13.44 ± 0.45 |
| ***ndufa13 L140*** | | | | |
| **Model** | **119E** | **120A** | **122D** | **126I** |
| **RaptorX** | 28.51 | 25.70 | 19.26 | 16.95 |
| **5LNK** | 12.58 ± 2.42 | 11.94 ± 2.74 | 11.14 ± 2.28 | 13.08 ± 0.55 |
| **5LDW** | 13.73 ± 1.84 | 13.73 ± 2.07 | 10.94 ± 1.95 | 13.63 ± 0.63 |
| **5XTC** | 14.62 ± 1.48 | 11.06 ± 1.38 | 10.29 ± 1.10 | 12.26 ± 0.24 |
| **6G2J** | 13.70 ± 2.09 | 11.36 ± 1.00 | 14.6 ± 1.43 | 12.97 ± 1.08 |

**Table S15**. Results for CodeML branch tests for NDUFS5, NDUFA13, and all mitochondrial Complex I genes. The variable vs. constant rate *P*-value listed is from a likelihood ratio test comparing a model with a constant rate to an alternative model of a distinct substitution rate on the *X. birchmanni* branch. The *P*-value in rightmost column is from testing two models: one where dN/dS on the *X. birchmanni* branch is allowed to vary versus a null model the *X. birchmanni* branch dN/dS is fixed at 1.

| **Gene** | **dN/dS Estimate, variable rate** | | **dN/dS Estimate, constant rate** | ***P*-value, variable vs. constant rate** | ***P*-value, *X. birchmanni* free vs. = 1** |
|---|---|---|---|---|---|
| | ***X. birchmanni*** | **Others** | **All branches** | | |
| **NDUFS5** | 999 | 0.08 | 0.26 | 0.005 | 0.186 |
| **NDUFA13** | 999 | 0.04 | 0.17 | 0.002 | 0.157 |
| **ND1** | 0.10 | 0.07 | 0.07 | 0.39 | 4.064e-8 |
| **ND2** | 0.11 | 0.10 | 0.10 | 0.91 | 1.550e-8 |
| **ND3** | 0.24 | 0.11 | 0.12 | 0.22 | 0.021 |
| **ND4** | 0.18 | 0.08 | 0.09 | 0.031 | 3.227e-8 |
| **ND4L** | 0.13 | 0.12 | 0.12 | 0.95 | 0.020 |
| **ND5** | 0.12 | 0.08 | 0.09 | 0.35 | 1.022e-8 |
| **ND6** | 10.5 | 0.09 | 0.10 | 0.005 | 0.126 |

**Table S16**. SIFT analyses of all amino acid substitutions that differ between *X. birchmanni* and *X. malinche* at *ndufs5, ndufa13* and the mt-DNA encoded proteins *ndufs5* contacts. All substitutions listed in both proteins are inferred to be derived in *X. birchmanni*. Substitutions are listed with the single letter amino acid code for the *X. malinche* allele first, followed by the *X. birchmanni* allele, separated by the residue number.

| Gene | Substitutions predicted tolerated | Substitutions predicted not tolerated |
|---|---|---|
| *ndufs5* | L20P<br>G44S | A31M<br>R51Q |
| *ndufa13* | H79Y | N90S<br>F140L |
| *nd2* | Y27F<br>G83E<br>T89M<br>I98T<br>I191L<br>V243I<br>T309A<br>M311T<br>N317K<br>R319H<br>T331M<br>F335L<br>A338I | N148D |
| *nd3* | I21V<br>D80N<br>A83T<br>H86Y<br>A90I<br>V104M | L64P |
| *nd4l* | T57A<br>N94S | M37T |
| *nd6* | V7I<br>G45V<br>S46N<br>G119E<br>M120A<br>E122D<br>L126I<br>L135V<br>G147A | S52P |

**Table S17.** RNAseq data derived from liver tissue used to evaluate expression of mitochondrial and nuclear genes of interest.

| **Group** | **Data source** |
|---|---|
| *X. birchmanni* | Pending, PRJNA746324; SUB9964665 |
| $F_1$ hybrids | Pending, PRJNA746324; SUB9964665 |
| *X. malinche* | Pending, PRJNA746324; SUB9964665 |

**Table S18**. High resolution mitochondrial respiration protocol and associated flux states.

| State (Fig. S17) | Titration (mM) | Abbreviation | Sites of electron entry | Explanation |
|---|---|---|---|---|
| A | Malate (0.5)<br>Pyruvate (10)<br>Glutamate (10) | M<br>P<br>G | CI | LEAK-associated (non-coupled) respiration supported by saturating concentrations of substrates for electron supply to Complex I in the absence of ADP |
| B | ADP (3) | ADP | CI | OXPHOS-associated respiration supported by electrons from Complex I |
| C | Succinate (40) | S | CI + CII | Maximal OXPHOS-associated respiration supported by electrons from CI and CII |
| D | Oligomycin (5 nM) | Omy | CI + CII | LEAK-associated respiration when ATP synthase is inhibited |
| E | Carbonyl cyanide m-chlorophenyl hydrazone (0.01) | CCCP | CI + CII | Uncoupled maximum electron transport capacity |
| F | Rotenone (0.01) | ROT | CII | Respiration supported by electrons from CII while CI is inhibited |
| G | Malonate (5) | MAL | None | LEAK state when CI and CII are inhibited |
| H | Ascorbate (10)<br>N,N,N′, N′-tetramethyl-p-phenylenediamine (0.3) | ASC<br>TMPD | CIV | Maximal capacity of CIV-mediated respiration supported by artificial electron donors |

**Table S19**. Respiratory flux control factors

| Flux control factor | Calculation[a] | Explanation |
|---|---|---|
| CI efficiency | 1-(A/B) | Extent of ADP control over respiration driven by CI, relating to the coupling efficiency of oxidative phosphorylation ranging from zero (completely non-coupled or no ADP control) to 1.0 (maximally coupled or 100% ADP control) |
| CII-A flux control | 1-(B/C) | Proportion of maximal (NADH + Succinate-supported) OXPHOS-linked respiration contributed by CII |
| CI + CII efficiency | 1-(D/C) | Extent of ADP control over respiration driven by CI and CII |
| ETS capacity | 1-(D/E) | Maximum capacity of the ETS to oxidize substrates in an uncoupled state |
| CI flux control | 1-(F/E) | Proportion of maximum ETS capacity driven by CI |
| CII-B flux control | 1-(G/F) | Proportion of maximum ETS capacity driven by CII |
| Excess CIV capacity | (H/G)-1 | Apparent excess respiratory capacity of cytochrome c oxidase over a “LEAK” state |

[a]Letters refer to the respiration states in Table S11 and Fig. S17

**Table S20.** Diagnostic heavy-labeled peptides synthesized for parallel reaction monitoring experiments. Asterix symbol indicate amino acids synthesized with heavy C and N. All cysteines were carbamoylmethylated.

| **Amino acid Sequence** | **Gene** | **Species** | **Molecular weight (Daltons)** |
|---|---|---|---|
| PFVDVQSR* | NDUFS5 | Shared | 956.9756 |
| WLLLQSGEQPYK* | NDUFS5 | *X. malinche* | 1469.60261 |
| WLLPQSGEQPYK* | NDUFS5 | *X. birchmanni* | 1453.56015 |
| WLLLQSGEQPYKR* | NDUFS5 | *X. malinche* | 1627.7751 |
| WLLPQSGEQPYKR* | NDUFS5 | *X. birchmanni* | 1611.73264 |
| DWVECGHGIGQTR* | NDUFS5 | *X. malinche* | 1524.55106 |
| DWVECGHGIGQTQAK* | NDUFS5 | *X. birchmanni* | 1723.78395 |
| LLIEELEAR* | NDUFA13 | Shared | 1095.18092 |
| ENLEEESIIMK* | NDUFA13 | *X. malinche* | 1342.43515 |
| ESLEEESIIMK* | NDUFA13 | *X. birchmanni* | 1315.40981 |
| IALMPLIQAEHDR* | NDUFA13 | *X. malinche* | 1516.698 |
| IALMPLIQAEYDR* | NDUFA13 | *X. birchmanni* | 1542.73198 |

**Table S21**. Data links and SRA accessions for previously sequenced individuals used in mitochondrial phylogenetic analysis.

| **Species** | **Data link or accession** |
|---|---|
| *X. cortezi* | Dryad: https://doi.org/10.5061/dryad.z8w9ghx82 |
| *X. malinche* | Dryad: https://doi.org/10.5061/dryad.z8w9ghx82 |
| *X. birchmanni* | Dryad: https://doi.org/10.5061/dryad.z8w9ghx82 |
| *X. couchianus* | SRA: SRR2127230 |
| *X. maculatus* | GenBank: AP005982.1 |
| *X. hellerii* | SRA: SRR7532852 |

**Table S22.** Pairwise mitochondrial divergence between *Xiphophorus* sequences generated by HiFi PacBio amplicon sequencing or 10X linked read sequencing.

| | ***X. birchmanni*** | | | | | | ***X. malinche*** | | | | ***X. cortezi*** | |
|---|---|---|---|---|---|---|---|---|---|---|---|---|
| | **Ref.** | **COAC1** | **COAC2** | **COAC3** | **COAC4** | **COAC5** | **Ref.** | **CAPAC** | **TETI** | **XFAU** | **PTHC** | **Stock** |
| **Ref.** | - | 0.0013 | 0.0012 | 0.0012 | 0.0008 | 0.0013 | 0.0539 | 0.0540 | 0.0543 | 0.0538 | 0.0540 | 0.0539 |
| **COAC1** | | - | 0.0004 | 0.0004 | 0.0007 | 0.0005 | 0.0537 | 0.0539 | 0.0542 | 0.0536 | 0.0539 | 0.0540 |
| **COAC2** | | | - | 0.0000 | 0.0004 | 0.0002 | 0.0537 | 0.0539 | 0.0542 | 0.0536 | 0.0539 | 0.0540 |
| **COAC3** | | | | - | 0.0004 | 0.0002 | 0.0537 | 0.0539 | 0.0542 | 0.0536 | 0.0539 | 0.0540 |
| **COAC4** | | | | | - | 0.0005 | 0.0538 | 0.0540 | 0.0543 | 0.0537 | 0.0540 | 0.0541 |
| **COAC5** | | | | | | - | 0.0535 | 0.0538 | 0.0541 | 0.0535 | 0.0538 | 0.0538 |
| **Ref.** | | | | | | | - | 0.0008 | 0.0011 | 0.0001 | 0.0055 | 0.0057 |
| **CAPAC** | | | | | | | | - | 0.0003 | 0.0008 | 0.0055 | 0.0057 |
| **TETI** | | | | | | | | | - | 0.0011 | 0.0058 | 0.0060 |
| **XFAU** | | | | | | | | | | - | 0.0054 | 0.0056 |
| **PTHC** | | | | | | | | | | | - | 0.0006 |
| **Stock** | | | | | | | | | | | | - |

**Table S23.** Pairwise mitochondrial and nuclear divergence between pairs of *Xiphophorus* species with no known history of introgression. Above the diagonal shows mitochondrial divergence, below the diagonal shows nuclear divergence.

| | *X. malinche* | *X. hellerii* | *X. couchianus* | *X. maculatus* |
|---|---|---|---|---|
| *X. malinche* | - | 0.075 | 0.074 | 0.072 |
| *X. hellerii* | 0.016 | - | 0.055 | 0.053 |
| *X. couchianus* | 0.015 | 0.017 | - | 0.046 |
| *X. maculatus* | 0.016 | 0.017 | 0.013 | - |

### 4 Supplementary Information References

16. Marlow, F. L. *Maternal Control of Development in Vertebrates: My Mother Made Me Do It!* (Morgan & Claypool Life Sciences, 2010).

39. Tavolga, W. N. Embryonic development of the platyfish (Platypoecilus), the swordtail (Xiphophorus), and their hybrids. Bulletin of the AMNH ; v. 94, article 4. *Embryonic development in fish* (1949).
