## Supplementary Data 1 for "A Lethal Genetic Incompatibility between Naturally Hybridizing Species in Mitochondrial Complex I"

**ND2**

>xbir_ND2

MAPFVCSTLIISLGLGTTMTFASTHWFLAWMGIEINTLAIIPLMTQNHNPRTIEATTKYFFAQATASATLLFAAVSNAFL

TGEWDILQMNHPLTSTLTTLALAMKIGLAPLHSWMPEVMQGVSLLTGLTLSTWQKLAPLCLIYQIQPD-SPSVFTTLGLL

SVIVGGWGGLNQVQLRKILAYSSIAHLGWMILILPFSPPLTLLTLFTYLMMTFSLFSSFMLIRTTHINSLSTSWAKIPAL

TASIPLILLSLGGLPPLTGFLPKWLILQELTKQDLAPIATLAALSSLFSLYFYLRLSYTMTLTMPPNNPAGTLPWRLKPH

HNTLPLALTTTMTIYLLPITPAILALFTL

>xmal_ND2

MAPFVCSTLIISLGLGTTMTFASTHWYLAWMGIEINTLAIIPLMTQNHNPRTIEATTKYFFAQATASATLLFAAVSNAFL

TGGWDILQTNHPLTSTLITLALAMKIGLAPLHSWMPEVMQGVSLLTGLTLSTWQKLAPLCLIYQIQPN-SPSVFTTLGLL

SVIVGGWGGLNQVQLRKILAYSSIAHLGWMIIILPFSPPLTLLTLFTYLMMTFSLFSSFMLIRTTHINSLSTSWAKIPAL

TASVPLILLSLGGLPPLTGFLPKWLILQELTKQDLAPIATLAALSSLFSLYFYLRLSYTMTLTMPPNNPTGMLPWRLNPR

HNTLPLALTTTTTIYFLPATPAILALFTL

>Mus_ND2

MNPITLAIIYFTIFLGPVITMSSTNLMLMWVGLEFSLLAIIPMLINKKNPRSTEAATKYFVTQATASMIILLAIVLNYKQ

LGTWMFQQQTNGLILNMTLMALSMKLGLAPFHFWLPEVTQGIPLHMGLILLTWQKIAPLSILIQIYPLLNSTIILMLAIT

SIFMGAWGGLNQTQMRKIMAYSSIAHMGWMLAILPYNPSLTLLNLMIYIILTAPMFMALMLNNSMTINSISLLWNKTPAM

LTMISLMLLSLGGLPPLTGFLPKWIIITELMKNNCLIMATLMAMMALLNLFFYTRLIYSTSLTMFPTNNNSKMMTHQTKT

KPNLMFSTLAIMSTMTLPLAPQLIT----

>Ovis_ND2

MNPIILIIILMTVMLGTIIVMISTHWLLIWIGFEMNMLAIIPIMMKKHNPRATEASTKYFLTQSTASMLLMMAIIINLMF

SGQWTVMKLFNPMASMLMTMALAMKLGMAPFHFWVPEVTQGIPLSSGLILLTWQKLAPMSVLYQILPSINLDLILTLSIL

SITIGGWGGLNQTQLRKIMAYSSIAHMGWMTAVLLYNPTMTLLNLIIYIIMTSTMFTLFMANSTTTTLSLSHTWNKAPIM

TILVLITLLSMGGLPPLSGFMPKWMIIQEMTKNDSIILPTLMAITALLNLYFYMRLTYSTALTMFPSTNNMKMKWQFPTT

KRMTLLPTMTVLSTMLLPLTPILSILE--

>Bos_ND2

MNPIIFIIILLTIMLGTIIVMISSHWLLVWIGFEMNMLAIIPIMMKNHNPRATEASTKYFLTQSTASMLLMMAVIINLMF

SGQWTVMKLFNPMASMLMTMALAMKLGMAPFHFWVPEVTQGIPLSSGLILLTWQKLAPMSVLYQIFPSINLNLILTLSVL

SILIGGWGGLNQTQLRKIMAYSSIAHMGWMTAVLPYNPTMTLLNLIIYIIMTSTMFTMFMANSTTTTLSLSHTWNKTPIM

TVLILATLLSMGGLPPLSGFMPKWMIIQEMTKNNSIILPTFMAITALLNLYFYMRLTYSTTLTMFPSTNNMKMKWQFPLM

KKMTFLPTMVVLSTMMLPLTPMLSVLE--

>Human_ND2

MNPLAQPVIYSTIFAGTLITALSSHWFFTWVGLEMNMLAFIPVLTKKMNPRSTEAAIKYFLTQATASMILLMAILFNNML

SGQWTMTNTTNQYSSLMIMMAMAMKLGMAPFHFWVPEVTQGTPLTSGLLLLTWQKLAPISIMYQISPSLNVNLLLTLSIL

SIMAGSWGGLNQTQLRKILAYSSITHMGWMMAVLPYNPNMTILNLTIYIILTTTAFLLLNLNSSTTTLLLSRTWNKLTWL

TPLIPSTLLSLGGLPPLTGFLPKWAIIEEFTKNNSLIIPTIMATITLLNLYFYLRLIYSTSITLLPMSNNVKMKWQFEHT

KPTPFLPTLIALTTLLLPISPFMLMIL--

**ND3**

>Xbir_ND3

MNLVVTTFSISLLLSLLLATVAFWLPLMTPDHEKLSPYECGFDPLGSARLPFSIRFFLVAILFPLFDLEIALLLPLPWGN

QLTDPYLTFIYASILLILLTLGLMYEWTQGGLEWAE

>Xmal_ND3

MNLVVTTFSISLLLSLLLATIAFWLPLMTPDHEKLSPYECGFDPLGSARLPFSIRFFLVAILFLLFDLEIALLLPLPWGD

QLADPHLTFAYASILLILLTLGLVYEWTQGGLEWAE

>Mus_ND3

MNLYTVIFI-NILLSLTLILVAFWLPQMNLYSEKANPYECGFDPTSSARLPFSMKFFLVAITFLLFDLEIALLLPLPWAI

QTIKTSTMMIMAFILVTILSLGLAYEWTQKGLEWTE

>Human_ND3

MNFALILMT-NTLLALLLMIITFWLPQLNGYMEKSTPYECGFDPMSPARVPFSMKFFLVAITFLLFDLEIALLLPLPWAL

QTTNLPLMVMSSLLLIIILALSLAYEWLQKGLDWAE

>Ovis_ND3

MNLMITLLT-NFTLATLLVTIAFWLPQLNVYSEKTSPYECGFDPMGSARLPFSMKFFLVAITFLLFDLEIALLLPLPWAS

QTTNLNTMLTMALLLIFLLAVSLAYEWTQKGLEWTE

>Bos_ND3

MNLMLALLT-NFTLATLLVIIAFWLPQLNVYSEKTSPYECGFDPMGSARLPFSMKFFLVAITFLLFDLEIALLLPLPWAS

QTANLNTMLTMALFLIILLAVSLAYEWTQKGLEWTE

**ND4L**

>Xbir_ND4L

MTPAHFAFSSAFMLGLAGLAFHRTHFLSALLCLEGLTLSLFIALSLWALQFNTMNSAALPMILLAFSACEAGAGLALLVA

TTRTHTSSRLQSLSLLQC

>Xmal_ND4L

MTPAHFAFSSAFMLGLAGLAFHRTHFLSALLCLEGLMLSLFIALSLWALQFNTMNSTALPMILLAFSACEAGAGLALLVA

TTRTHTSSRLQSLNLLQC

>Mus_ND4L

MPSTFFNLTMAFSLSLLGTLMFRSHLMSTLLCLEGMVLSLFIMTSVTSLNSNSMSSMPIPITILVFAACEAAVGLALLVK

VSNTYGTDYVQNLNLLQC

>Human_ND4L

MPLIYMNIMLAFTISLLGMLVYRSHLMSSLLCLEGMMLSLFIMATLMTLNTHSLLANIMPIVMLVFAACEAAVGLALLVS

ISNTYGLDYVHNLNLLQ-

>Ovis_ND4L

MSLVYMNIMMAFTVSLTGLLMYRSHLMSSLLCLEGMMLSLFILATLMILNSHFTLASMMPIILLVFAACEAALGLSLLVM

VSNTYGTDYVQNLNLLQC

>Bos_ND4L

MSMVYMNIMMAFTVSLVGLLMYRSHLMSSLLCLEGMMLSLFVMAALTILNSHFTLASMMPIILLVFAACEAALGLSLLVM

VSNTYGTDYVQNLNLLQC

**ND6**

>Xbir_ND6

-MSYLMYILLSGLVLGLVAVASNPSPYFAAFGLVVVAGVGCGVLVVNGGCFLPLILFLIYLGGMLVVFAYSAAMAAEPFP

EGWGSWPTL--RLMVGYVVGVGFAW--------GLMGKIWHEEDWLGFDEAVDLSVIRGDVGGVSVVYSSGGWLLIGAGW

VLLLTLFVVLELTRGLSRGALRAV

>Xmal_ND6

-MSYLMYVLLSGLVLGLVAVASNPSPYFAAFGLVVVAGVGCGVLVGSGGCFLSLILFLIYLGGMLVVFAYSAAMAAEPFP

EGWGSWPTL--RLMVGYVVGVGFAW--------GLMGKIWHEEDWLGFDGMVELSVLRGDVGGVSLVYSSGGWLLIGGGW

VLLLTLFVVLELTRGLSRGALRAV

>Ovis_ND6

MMTYIVFILSIIFVMGFVGFSSKPSPIYGGLGLIVSGGVGCGIVLNFGGSFLGLMVFLIYLGGMMVVFGYTTAMATEQYP

EVWVSNKVVLGTFITGLLMEFLMVYYVLKDKEVEIVFKFNGMGDWVIYDTG-DSGFFSEEAMGIAALYSYGTWLVIVTGW

SLLIGVVVIMEITRGN--------

>Bos_ND6

MMLYIVFILSVIFVMGFVGFSSKPSPIYGGLGLIVSGGVGCGIVLNFGGSFLGLMVFLIYLGGMMVVFGYTTAMATEQYP

EIWLSNKAVLGAFVTGLLMEFFMVYYVLKDKEVEVVFEFNGLGDWVIYDTG-DSGFFSEEAMGIAALYSYGTWLVIVTGW

SLLIGVVVIMEITRGN--------

>Human_ND6

-MMYALFLLSVGLVMGFVGFSSKPSPIYGGLVLIVSGVVGCVIILNFGGGYMGLMVFLIYLGGMMVVFGYTTAMAIEEYP

EAWGSGVEVLVSVLVGLAMEVGLVLWVKEYDGVVVVVNFNSVGSWMIYEGE-GSGFIREDPIGAGALYDYGRWLVVVTGW

TLFVGVYIVIEIARGN--------

>Mus_ND6

-MNNYIFVLSSLFLVGCLGLALKPSPIYGGLGLIVSGFVGCLMVLGFGGSFLGLMVFLIYLGGMLVVFGYTTAMATEEYP

ETWGSNWLILGFLVLGVIMEVFLICVLNYYDEVGVI-NLDGLGDWLMYEVD-DVGVMLEGGIGVAAMYSCATWMMVVAGW

SLFAGIFIIIEITRD---------

**NDUFA1**

>Xbir_NDUFA1

MWYEILPAFGLMTVCMILPGIVTTHIHKFTNGGKEKRIARNPWQWYLMERDKRVSGTEQYFNSKGLENIK

>Xmal_NDUFA1

MWYEILPAFGLMTVCMILPGIVTTHIHKFTNGGKEKRIARNPWQWYLMERDKRVSGTEQYFNSKGLENIK

>Mus_NDUFA1

MWFEILPGLAIMGVCLVIPGVSTAYIHKFTNGGKEKRVARVQYQWYLMERDRRISGVNRYYVSKGLE---

>Human_NDUFA1

MWFEILPGLSVMGVCLLIPGLATAYIHRFTNGGKEKRVAHFGYHWSLMERDRRISGVDRYYVSKGLENID

>Ovis_NDUFA1

MWFEVLPGIAVMGVCLFIPGMATARIHRFSNGGREKRVAHYSYQWYLMERDRRVSGVNRYYVSKGLENID

>Bos_NDUFA1

MWFEVLPGIAVMGVCLFIPGMATARIHRFSNGGKEKRVAHYPYQWYLMERDRRVSGVNRSYVSKGLENID

**NDUFA8**

>Xbir_NDUFA8

MPTTLEVPTLQDLKVDEVNVSSAVLKAAAHHYGSQCDKPNKEFMLCRWEEKDPRKCLQEGKKVNECALNFFRQIKGNCAE

SFTEYWTCLDYTNLTELRHCRKQQQAFDSCVLDKLGWVRPDLGDLSKVTKVSTSRPLPENPYHSRPRPEPNPTIEGNLEP

AKHGSRFFFWTW

>Xmal_NDUFA8

MPTTLEVPTLQDLKVDEVNVSSAVLKAAAHHYGSQCDKPNKEFMLCRWEEKDPRKCLQEGKKVNECALNFFRQIKGNCAE

SFTEYWTCLDYTNLTELRHCRKQQQAFDSCVLDKLGWVRPDLGDLSKVTKVSTSRPLPENPYHSRPRPEPNPTIEGNLEP

AKHGSRLFFWTW

>Mus_NDUFA8

MPGIVELPTLEELKVEEVKVSSAVLKAAAHHYGAQCDKTNKEFMLCRWEEKDPRRCLKEGKLVNGCALNFFRQIKSHCAE

PFTEYWTCLDYSNMQLFRHCRQQQAKFDQCVLDKLGWVRPDLGQLSKVTKVKTDRPLPENPYHSRARPEPNPVIEGDLKP

AKHGTRFFFWTV

>Human_NDUFA8

---IVELPTLEELKVDEVKISSAVLKAAAHHYGAQCDKPNKEFMLCRWEEKDPRRCLEEGKLVNKCALDFFRQIKRHCAE

PFTEYWTCIDYTGQQLFRHCRKQQAKFDECVLDKLGWVRPDLGELSKVTKVKTDRPLPENPYHSRPRPDPSPEIEGDLQP

ATHGSRFYFWTK

>Ovis_NDUFA8

-PGIVELPSLEDLKVQEVKVSSSVLKAAAHHYGAQCDKPNKEFMLCRWEEKDPRRCLEEGKLVNQCALEFFRQIKRHCAE

PFTEYWTCIDYSGLQLFRRCRKEQAQFDKCVLDKLGWVRPDLGELSKVTKVKTDRPLPENPYHSRARPEPNPEVEGDLKP

ARHGSRLFFWTM

>Bos_NDUFA8_PDB

MPGIVELPSLEDLKVQEVKVSSSVLKAAAHHYGAQCDKPNKEFMLCRWEEKDPRRCLEEGKLVNQCALEFFRQIKRHCAE

PFTEYWTCIDYSGLQLFRRCRKQQAQFDECVLDKXXXXXXXXXXXXXXXXXXXXXXXXXXXXXXXXXXXXXXXXXXXXXX

XXXXXXXXX---

>NP_787020.1 Bos_NDUFA8

MPGIVELPSLEDLKVQEVKVSSSVLKAAAHHYGAQCDKPNKEFMLCRWEEKDPRRCLEEGKLVNQCALEFFRQIKRHCAE

PFTEYWTCIDYSGLQLFRRCRKQQAQFDECVLDKLGWVRPDLGDLSKVTKVKTDRPLPENPYHSRARPEPNPEVEGDLKP

ARHGSRLFFWTM

**NDUFA13**

>Xbir_NDUFA13

MAGSKVKQDMPPSGGYPAFDYKRNLPKRGLSGYSMFGIGIGIMAFGYWRIFSWNRERRRLLIEELEARIALMPLIQAEYD

RRTLRMLRESLEEESIIMKDVPGWKVGERVFHTDRWVPPLSDEMFSLRPREQYLHKRFGLLWYV

>Xmal_NDUFA13

MAGSKVKQDMPPSGGYPAFDYKRNLPKRGLSGYSMFGIGIGIMAFGYWRIFSWNRERRRLLIEELEARIALMPLIQAEHD

RRTLRMLRENLEEESIIMKDVPGWKVGERVFHTDRWVPPLSDEMFSLRPREQYLHKRFGFLWYV

>Mus_NDUFA13

MAASKVKQDMPPPGGYGPIDYKRNLPRRGLSGYSMFAVGIGALIFGYWRMMRWNQERRRLLIEDLEARIALMPLFQAEKD

RRTLQILRENLEEEAIIMKDVPNWKVGESVFHTTRWVPPLIGEMYGLRTKEEMSNANFGFTWYT

>Human_NDUFA13_PDB

----------------------------GLSGYSMLAIGIGTLIYGHWSIMKWNRERRRLQIEDFEARIALLPLLQAETD

RRTLQMLRENLEEEAIIMKDVPDWKVGESVFHTTRWVPPLIGELYGLRTTEEALHASHGFMWYT

>AAH09189.1 Human_NDUFA13

MAASKVKQDMPPPGGYGPIDYKRNLPRRGLSGYSMLAIGIGTLIYGHWSIMKWNRERRRLQIEDFEARIALLPLLQAETD

RRTLQMLRENLEEEAIIMKDVPDWKVGESVFHTTRWVPPLIGELYGLRTTEEALHASHGFMWYT

>Ovis_NDUFA13

-AASKVKQDMPPVGGYGPIDYKRNLPRRGLSGYSMFAVGIGALLFGYWSMMRWNRERRRLQIEDFEARIALMPLLQAEKD

RRVLQMLRENLEEEATIMKDVPGWKVGESVFHTTRWVTPMMGELYGLRTGEEILSSTYGFIWYT

>Bos_NDUFA13_PDB

-----XXXXXXXXXXXXXXXXXXXXXXXXXXGYSMFAVGIGALLFGYWSMMKWNRERRRLQIEDFEARIALMPLLQAEKD

RRVLQMLRENLEEEATVMKDXXXXXXXXXXXXXXXXXXXXXXXXXXXXXXXXXXXXXXXXXXX-

>NP_788845.1 Bos_NDUFA13

MAASKVKQDMPPVGGYGPIDYKRNLPRRGLSGYSMFAVGIGALLFGYWSMMKWNRERRRLQIEDFEARIALMPLLQAEKD

RRVLQMLRENLEEEATVMKDVPGWKVGESVFHTTRWVTPMMGELYGLRASEEVLSATYGFIWYT

**NDUFB5**

>Xbir_NDUFB5

MVGMSLLRSAAAFA-ARLGPLKAGNNA--ANILARTIPRTNKVATRWG-HGKKMFVVNPTDYYDRRFLHLLRYYILLTGI

PVALLVTAVNIFIGEAELAEIPEGYEPEYWEYYKHPITRWIVRNIYDSPVKDYEKVMAAIQIEKEKADMRLTQLEVRRQM

RHHGDGPWFQVPTADKGLIDNSPKSSPDN

>Xmal_NDUFB5

MVGMSLLRSAAAFA-ARLGPLKAGNNA--ANILARTIPRTNKVATRWG-HGKKMFVVNPTDYYDRRFLHLLRYYILLTGI

PVALLVTAVNIFIGEAELAEIPEGYEPEYWEYYKHPITRWIVRNIYDSPVKDYEKVMAAIQIEKEKADMRLTQLEVRRQM

RHHGDGPWFQVPTADKGLIDNSPKSSPDN

>Mus_NDUFB5

MAAMSLLQRASVSALTALSCRRAGPRLGVGSFLTRSFPKTVAPVRHSGDHGKRLFVVKPSLYYDARFLRLMKFYLMLTGI

PVIIGITLVNIFIGEAELAEIPEGYIPEHWEYYKHPISRWIARNFYDGPEKNYEKTLAILQIESEKAELRLKEQEVRRLM

RARGDGPWYQFPTPEKEFIDHSPKATPDN

>Human_NDUFB5_PDB

---------------------------------------------------KRLFVIRPSRFYDRRFLKLLRFYIALTGI

PVAIFITLVNVFIGQAELAEIPEGYVPEHWEYYKHPISRWIARNFYDSPEKIYERTMAVLQIEAEKAELRVKELEVRKLM

HVRGDGPWYYYETIDKELIDHSPKATPDN

>NP_002483.1 Human_NDUFB5

MAAMSLLRRVSVTAVAALSGRPLGTRLGFGGFLTRGFPKAAAPVRHSGDHGKRLFVIRPSRFYDRRFLKLLRFYIALTGI

PVAIFITLVNVFIGQAELAEIPEGYVPEHWEYYKHPISRWIARNFYDSPEKIYERTMAVLQIEAEKAELRVKELEVRKLM

HVRGDGPWYYYETIDKELIDHSPKATPDN

>Ovis_NDUFB5_PDB

----------------------------------------------SGDHGKRLFIIKPSGFYDKRFLKLLRFYILLTGI

PVVIGITLINVFIGEAELAEIPEGYVPEHWEYFKHPISRWIARTFFDAPEKNYERTMAILQIESEKAELRLKELEVRRLM

RAKGDGPWFQYPTIDKALIDHSPKATPDN

>XP_004003178.1 Ovis_NDUFB5

MAAMSLLHRASVSAVVALSGRRLGTRLGFGGFLTRDFPKTVAPVRHSGDHGKRLFIIKPSGFYDKRFLKLLRFYILLTGI

PVVIGITLINVFIGEAELAEIPEGYVPEHWEYFKHPISRWIARTFFDAPEKNYERTMAILQIESEKAELRLKELEVRRLM

RAKGDGPWFQYPTIDKALIDHSPKATPDN

>Bos_NDUFB5_PDB

----------------------------------------------------RLFIIKPSGFYDKRFLKLLRFYILLTGI

PVAIGITLINVXXXXXXXXXXXXXXXXXXXXXXXXXXXXXXXXXXXXXXXXXXXXXXXXXXXXXXXXXXXXXXXXXXXXX

XXXXXXXXXXXXXXXXXXXXXXXXXX---

>NP_788829.1 Bos_NDUFB5

MAAMSLLHRASVSAVAALSGRRLGTRLGFGGFLTRDFPKTVAPVRHSGDHGKRLFIIKPSGFYDKRFLKLLRFYILLTGI

PVAIGITLINVFIGEAELAEIPEGYVPEHWEYFKHPISRWIARTFFDGPEKNYERTMAILQIEAEKAELRLKELEVRRLM

RARGDGPWFHYPTIDKELIDHSPKATPDN

**NDUFC2**

>Xbir_NDUFC2

----------MGIVPEDGKVLPPPGIVNRNSVWLSGMGWFSALLNNAFNHRPPLKSGVHRQFLFATIGWYIGYHLTKYEN

YTYARLDRDMNEYVKLHLERFETKEKKTFAEIVEPFHPIR

>Xmal_NDUFC2

----------MGIVPEDGKVLPPPGIVNRNSVWLSGMGWFSALLNNAFNHRPPLKSGVHRQFLFATIGWYIGYHLTKYEN

YTYARLDRDMNEYVKLHPERFETKEKKTFAEIVEPFHPIR

>Mus_NDUFC2

MMNGRPGHEPLKFLPDEARSLPPPKLNDPRLVYMGLLGYCTGLMDNMLRMRPVMRAGLHRQLLFVTSFVFAGYFYLKRQN

YLYAVKDHDMFGYIKLHPEDFPEKEKKTYAEILEPFHPVR

>Human_NDUFC2

-MIARRNPEPLRFLPDEARSLPPPKLTDPRLLYIGFLGYCSGLIDNLIRRRPIATAGLHRQLLYITAFFFAGYYLVKRED

YLYAVRDREMFGYMKLHPEDFPEEDKKTYGEIFEKFHPIR

>Ovis_NDUFC2

MMTGRQARAPLQFLPDEARSLPPPKLTDPRLAYIGFLGYCSGLIDNAIRRRPVLSAGLHRQFLYITSFVFVGYYLLKRQD

YMYAVRDHDMFSYIKSHPEDFPEKDKKTYREVFEEFHPVR

>Bos_NDUFC2_PDB

--XXXXXXXXXXXXXXXXXXXXXXXXXXPRLAFVGFLGYCSGLIDNAIRRRPVLLAGLHRQLLYITSFVFVGYYLLKRQD

YMYAVRDHDMFSYIKSHXXXXXXXXXXXXXXXXXXX----

>NP_788815.1 Bos_NDUFC2

MMTGRQGRATFQFLPDEARSLPPPKLTDPRLAFVGFLGYCSGLIDNAIRRRPVLLAGLHRQLLYITSFVFVGYYLLKRQD

YMYAVRDHDMFSYIKSHPEDFPEKDKKTYGEVFEEFHPVR

**NDUFS5**

>ref|NP_004543.1| Human_NDUFS5

MPFLDIQKRFGLNIDRWLTIQSGEQPYKMAGRCHAFEKEWIECAHGIGYTRAEKECKIEYDDFVECLLRQKTMRRAGTIR

KQRDKLIKEGKYTPPPHHIGKGEPRP

>ref|NP_001025445.1| Mus_NDUFS5

MPFLDIQKKLGISLDRHFMFLSAEQPYKNAARCHAFEKEWIECAHGIGGTRAKKECKIEFDDFEECLLRYKTMRRMHDIK

KQREKLMKEGKYTPPPHHSGREEPRP

>ref|NP_788825.1| Bos_NDUFS5

MPFFDVQKRLGVDLDRWMTIQSAEQPHKIPSRCHAFEKEWIECAHGIGSIRAEKECKIEFEDFRECLLRQKTMKRLHAIR

RQREKLIKEGKYTPPPHHSGQEEPRS

>ref|XP_004001873.1| Ovis_NDUFS5

MPFFDVQKKLGVDLDHWMTIQSAEQPHRIPARCHAFEKEWIECAHGIGSIRAEKECKIEFEDFRECLLRQKTMKRLNAIK

RQRDKLIKEGKYTPPPHHSGQEDLRP

>Xbir_NDUFS5

MPFVDVQSRLGINVDRWLLPQSGEQPYKRAMRCHAFEKDWVECSHGIGQTQAKKECQLEYEDFYECMYRQKTRERMYAIR

AQRDKLIKEGKYTPPSHHTGQPDQNP

>Xmal_NDUFS5

MPFVDVQSRLGINVDRWLLLQSGEQPYKRAARCHAFEKDWVECGHGIGQTRAKKECQLEYEDFYECMYRQKTRERMYAIR

AQRDKLIKEGKYTPPSHHTGQPDQNP
